# HHIP’s Dynamic Role in Epithelial Wound Healing Reveals a Potential Mechanism of COPD Susceptibility

**DOI:** 10.1101/2024.09.05.611545

**Authors:** Dávid Deritei, Wardatul Jannat Anamika, Xiaobo Zhou, Edwin K. Silverman, Erzsébet Ravasz Regan, Kimberly Glass

## Abstract

A genetic variant near *HHIP* has been consistently identified as associated with increased risk for Chronic Obstructive Pulmonary Disease (COPD), the third leading cause of death worldwide. However HHIP’s role in COPD pathogenesis remains elusive. Canonically, HHIP is a negative regulator of the hedgehog pathway and downstream GLI1 and GLI2 activation. The hedgehog pathway plays an important role in wound healing, specifically in activating transcription factors that drive the epithelial mesenchymal transition (EMT), which in its intermediate state (partial EMT) is necessary for the collective movement of cells closing the wound. Herein, we propose a mechanism to explain HHIP’s role in faulty epithelial wound healing, which could contribute to the development of emphysema, a key feature of COPD. Using two different Boolean models compiled from the literature, we show dysfunctional HHIP results in a lack of negative feedback on GLI, triggering a full EMT, where cells become mesenchymal and do not properly close the wound. We validate these Boolean models with experimental evidence gathered from published scientific literature. We also experimentally test if low HHIP expression is associated with EMT at the edge of wounds by using a scratch assay in a human lung epithelial cell line. Finally, we show evidence supporting our hypothesis in bulk and single cell RNA-Seq data from different COPD cohorts. Overall, our analyses suggest that aberrant wound healing due to dysfunctional HHIP, combined with chronic epithelial damage through cigarette smoke exposure, may be a primary cause of COPD-associated emphysema.

**Graphical abstract:** 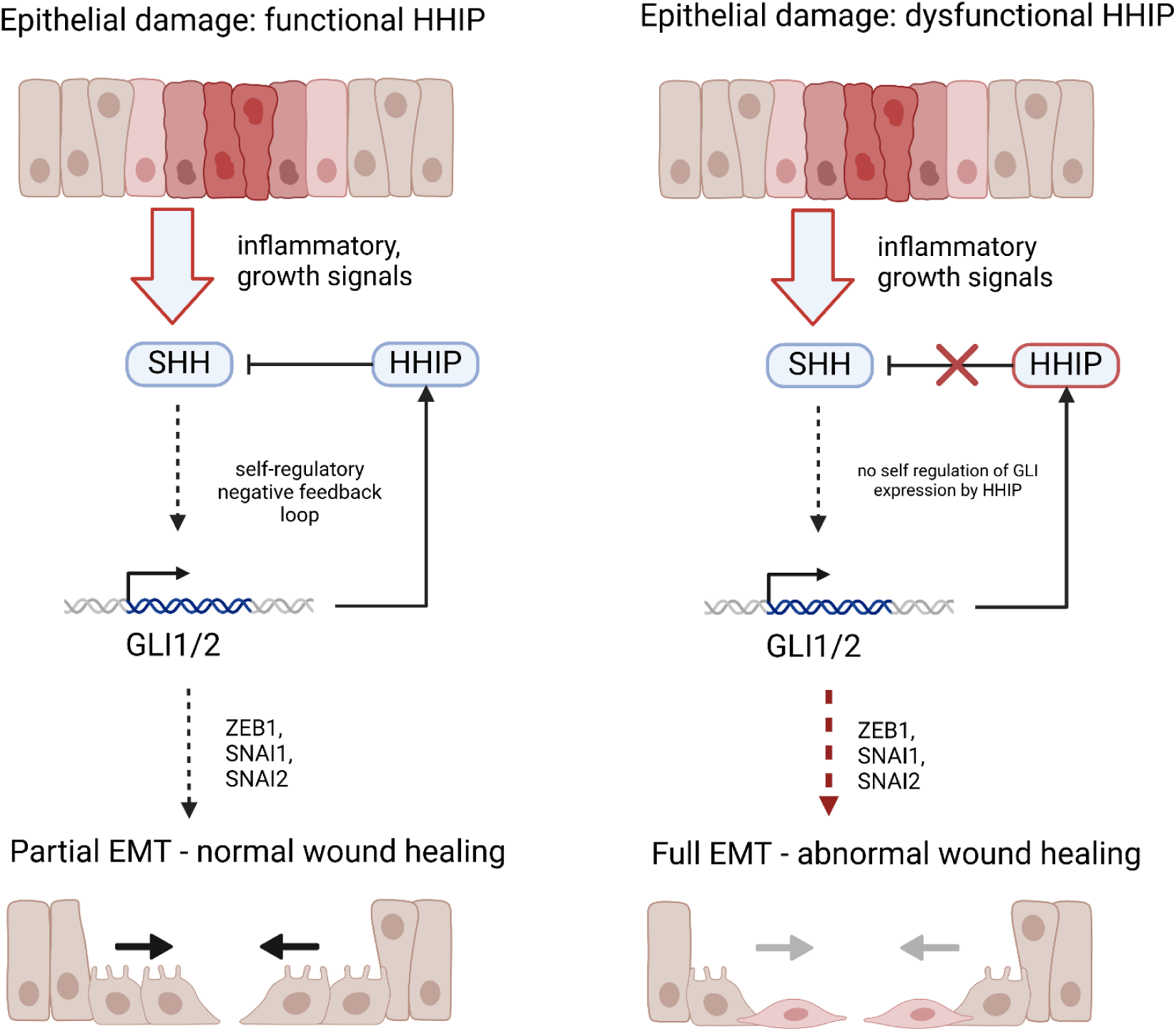

## Introduction

Chronic obstructive pulmonary disease (COPD) is one of the leading causes of death worldwide (*1*). COPD is a complex disease primarily caused by a combination of cigarette smoke and genetic predisposition. One of the primary pathological features of COPD is emphysema, which is the destruction of the alveolar structure and the surrounding tissue due to chronic inflammation and faulty wound healing (*2–4*). This process can be self-sustaining even years after a patient has quit smoking (*4*). Many of the biological details of COPD susceptibility and the mechanistic causes of emphysema are not well understood. Genome-wide association studies (GWAS) have identified more than 80 genomic regions that contribute to COPD susceptibility. Among COPD GWAS risk loci, a genomic region near Hedgehog interacting protein (*HHIP*), a regulator of the Hedgehog signaling pathway, is consistently associated with COPD risk (*5–8*). Furthermore, experiments in murine models have shown that *Hhip* haploinsufficiency in the presence of cigarette smoke contributes to emphysema (*9*, *10*). However, the mechanism through which *HHIP* risk variants lead to COPD susceptibility and specifically emphysema remains unclear.

Toxins such as cigarette smoke damage the alveolar epithelium, triggering a number of different processes, which mitigate the damage and initiate closure of a wound. One key process of re-epithelialization is the epithelial mesenchymal transition (EMT), a multi-step process during which cells downregulate the expression of cell-cell adhesion proteins (such as E-cadherin), increase motility, and transform into mesenchymal cells. The first step of EMT is *reversible* and is associated with more mobile but not fully detached cells capable of moving collectively (a state also termed hybrid E/M). In contrast, the second transition, where cells become mesenchymal and detach from their epithelial neighbors, is self-sustaining due to autocrine feedback signals such as TGFβ, secreted by mesenchymal cells (*11*). This multi-step process is governed by two regulatory switches created by different double-negative feedback loops (*12*). Partial or hybrid EMT is marked by SNAI1/2 upregulation, the downregulation of miR-34, and moderate TWIST and ZEB1 expression. At the same time, miR-200 remains expressed (inhibiting TGFβ protein expression) and E-Cadherin is also present, maintaining adherens junctions between cells (*13–15*).

Normal wound healing is associated with *partial, reversible* EMT. Epithelial cells at the edge of wounds partially differentiate and move collectively in order to close the wound, then reform the epithelium (*16–18*). Inflammatory cells recruited to the wound site secrete many of the signals necessary to induce the healing process. For example, macrophages promote proliferation and migration by secreting growth factors (FGF, EGF, TGF-β and PDGF) (*11*). EMT can be triggered by multiple signaling pathways, including TGFβ, Notch, and Wnt, as well as the Hedgehog (HH) pathway.

Signals on the edge of wounds can trigger Hedgehog signaling through both inflammatory and growth factor signaling pathways. For example, NF-Kβ, a target transcription factor of both growth factor and inflammatory signals, can contribute to the Hedgehog signaling cascade by transcribing SHH (*19*).The effector genes of the Hedgehog signaling cascade are GLI1 and GLI2. Studies have shown that among the transcription targets of GLI1/2 are EMT transcription factors, such as SNAIL1 (*20*, *21*) and ZEB1 (*22*). GLI1/2 also contributes to the nuclear localization of β-catenin (*23*), another process linked with EMT. To moderate their EMT-promoting effects, GLI1 and GLI2 also transcribe HHIP, a negative feedback regulator of the Hedgehog pathway and GLI1/2.

HHIP binds the membrane ligands SHH, IHH, and DHH (upstream receptor ligands of the HH pathway), and inhibits their ability to bind to Patch, quelling the signaling cascade (*24–26*). In this study we explore HHIP’s role in epithelial wound healing in a systems context, focusing on the way that the Hedgehog pathway interacts with the epithelial mesenchymal transition (EMT) pathway at the edges of wounds and linking its misregulation to aberrant wound healing in emphysema. We hypothesize that in the context of HHIP deficiency the Hedgehog pathway is hyper-activated. Specifically, HHIP deficiency leads to dysregulated GLI expression, which subsequently activates EMT drivers and leads to a higher likelihood of irreversible EMT (instead of the desired partial EMT) in the cell population around the damaged epithelium. As a consequence, fully mesenchymal cells do not reform the epithelium, divide less, and secrete factors that destroy the basement membrane (*27*, *28*), ultimately leading to a chronic state of alveolar breakdown, fibrosis, and inflammation (*11*, *29*).

To test our hypothesis we propose a Boolean network model partially based on previous EMT models that focuses on encapsulating the relevant interactions of Hedgehog signaling and EMT mechanisms. The model reproduces the wild-type partial EMT given in response to epithelial damage, as well as an irreversible mesenchymal transition in HHIP-deficient cells. We also reproduce some of the key phenotypes of emphysema in an expanded Boolean model of Sullivan et al. (*30*), which provides a higher-level regulatory context for wound healing, including regulatory modules such as the cell cycle and apoptosis. Guided by our model’s predictions, we explore our hypothesis experimentally in wound healing assays with the alveolar cancer cell line A549, and show that HHIP knockdown leads to the enrichment of EMT markers. We further support our hypothesis with single-cell RNA-Seq data from COPD lung epithelial cells.

## Results

### 1. Asynchronous model of EMT and Hedgehog signaling reproduces HHIP dependent partial vs. full EMT at the edge of a wound

#### Topological Features of the model

To better understand the relationship between HHIP and the different levels of EMT in the presence of epithelial damage, we constructed a simple Boolean dynamical system, as shown in Figure 1. The interactions of the network are drawn from the scientific literature and are all reported to be direct interactions, such as binding, transcription, or mRNA-degradation. See Supplementary Tables 1 and 2 for documentation on each node / edge. We note that we use a conservative approach that aims to explain the relevant phenotypes with a minimal set of preferably canonical interactions, however, this approach is vulnerable to ignoring some false negatives, and most interactions are drawn from a variety of experiments using many different cell types. We discuss the validation and limitations of the model in later sections of the manuscript.

**Figure 1:**
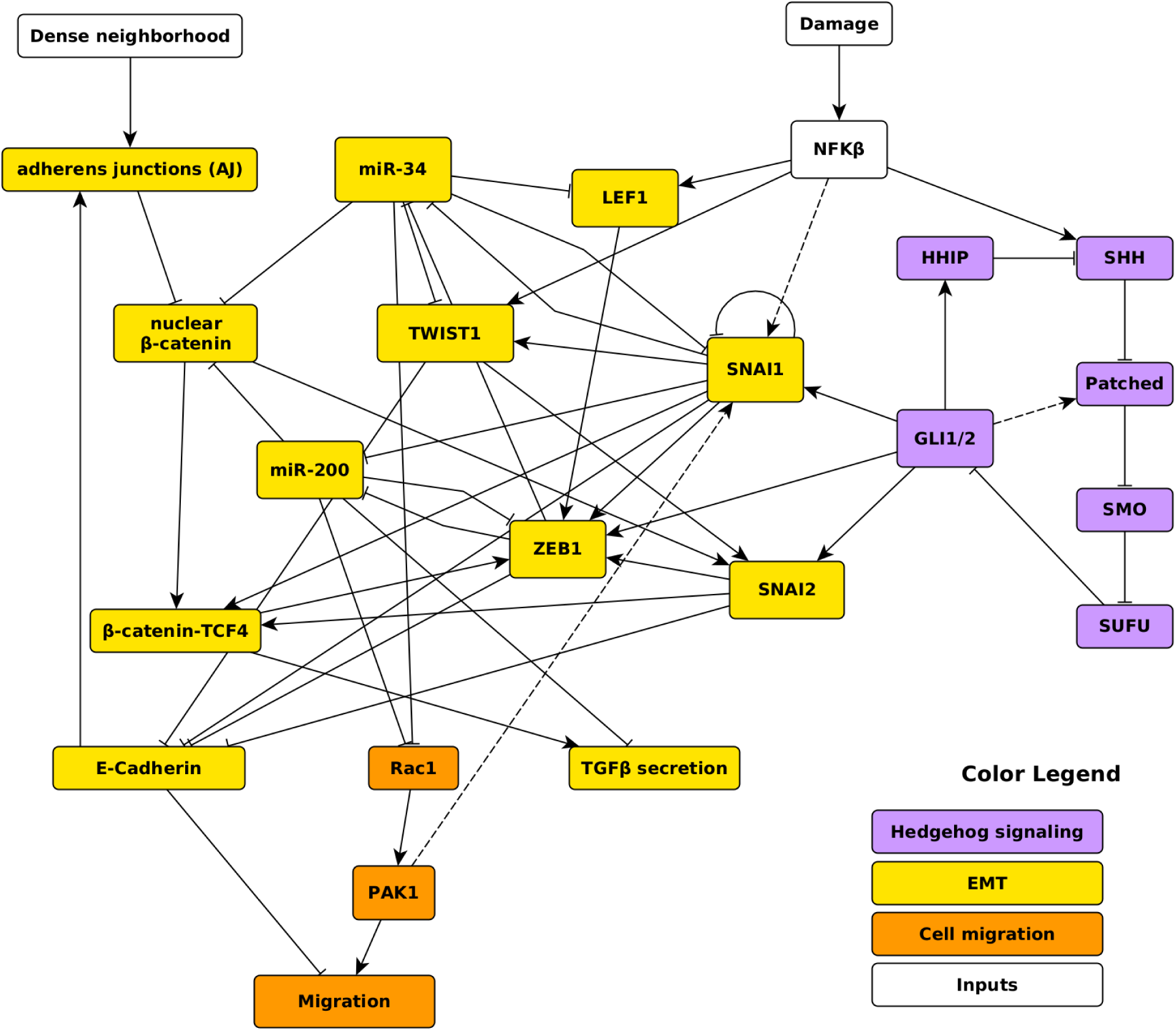
Network representation of the Boolean model of the interaction between Hedgehog signaling and EMT. Colors represent the different regulatory modules. Links ending in arrows represent directed activation (sufficient or necessary); links ending in T represent inhibition. The details of each interaction (literature source, interaction type, cell type, etc.) are documented in Supplementary Table 1. Dashed lines represent documented interactions that are not directly incorporated in the logical functions.

The network has three main modules and two inputs. The first module is the Hedgehog pathway, which is a linear signaling pathway with Sonic Hedgehog as its primary ligand (represented in the model by the node SHH). HHIP creates a negative feedback loop to the transcription factors GLI1 and GLI2, the effectors of the signaling pathway (GLI1 and GLI2 are merged into a single node). The second module is the larger EMT module, which is partly based on previously published dynamic models of EMT (*12*, *30–32*). We included the interactions discussed by (*12*), a previously published ODE model of EMT, which reproduced the two-step EMT (partial and full), interactions with adherens junction-mediated contact sensing from (*30*), and interactions with the Hedgehog pathway from (*31*). However, we reevaluated or fine-tuned the specifics of the model based on more recent literature and our hypothesis. For example, none of the previous models include HHIP, but many include ZEB2 and/or other microRNAs, which we decided to omit here because they were not critical to the phenotypes we are testing and tended to be downstream of pathways other than the Hedgehog pathway. We also removed nodes representing multiple concentration levels of the same gene/protein and chose to rely instead on dynamic equilibria in large ensembles of Boolean models, which can probabilistically generate moderate expression levels for nodes under conflicting regulatory constraints (for details, see Methods). The main link between Hedgehog signaling and EMT is GLI, which has been shown to promote multiple EMT transcription factors, such as SNAI1, SNAI2 and ZEB1 (*22*, *33*, *34*).

The third module contains nodes that are key readouts for cellular migration. While migration is a complex process involving a large ensemble of proteins and dynamics beyond our current scope (*35*), the two proteins included here – Rac1 and PAK1 – are not only essential to migration but are also regulated by nodes from the EMT module. Finally, the two inputs represent damage signals that act through the NFKβ inflammation pathway (“damage”) and the presence or absence of a dense neighborhood of cells (“Dense neighborhood”). For example, a scratch on an epithelium removes neighbors (Dense neighborhood=OFF or P(Dense neighborhood=ON)<1) and produces damage signals such as Damage-Associated Molecular Patterns (DAMPs) by the destruction of cells (*36*, *37*).

Notably, we exclude TGFβ signaling and related feedback mechanisms from this model. TGFβ is one of the main drivers of EMT, and the main focus of previous models (*31*, *38*). The reason for this is to show that misregulated Hedgehog signaling can cause a self-sustaining mesenchymal state even *without* TGFβ feedback. Nonetheless, we do include an output variable representing TGFβ secretion, which is required for irreversible EMT stabilization by TGFβ.

#### Dynamical Features of the model

Based on our hypothesis that partial EMT in the case of epithelial damage is sustained by the Hedgehog pathway, which in the absence of HHIP leads to full EMT, we expect three main biological phenotypes: a stable epithelial state, a reversible partial EMT state (in response to wound damage), and an irreversible fully mesenchymal state. Indeed the model has two steady states (fixed point attractors) corresponding to the epithelial and mesenchymal phenotypes and two complex attractors (oscillations), both of which correspond to the partial EMT in two different input configurations (see the attractor analysis in Supplementary File 1).

To simulate the dynamical behavior of the model, we use a general asynchronous update approach to run large ensembles of models from the same initial conditions. In Figure 2A we show the dynamic evolution of an ensemble of 5000 networks initiated in the epithelial state (phenotype). At time-step 300 we introduce a “wound” by changing the Damage input to 1 and the Dense neighborhood input to 0. This triggers the hedgehog pathway, but due to the negative feedback loop between GLI and HHIP we get an oscillating attractor where many variables reach a *dynamical equilibrium* other than 0 or 1 on average. This attractor corresponds to the hallmarks of the partial EMT associated with wild-type wound healing: SNAI1 and SNAI2 are upregulated, while ZEB1 settles at an intermediary concentration (*12*, *13*) (in this case mainly due to the oscillation of GLI). Among the antagonists of EMT, miR-34 is downregulated almost completely and allows the partial activation of migration, which is necessary for wound closure. E-cadherin is also in a dynamic equilibrium, representing the need for partial removal for cell-cell junctions necessary for motility and collective migration. Chiefly, the removal of the damage signals and the appearance of new neighbors (due to, for example, proliferation, not discussed here), shown in Figure 2A at time-step 1300, causes the system to return to the epithelial state, re-solidifying the junctions and forming a confluent monolayer.

**Figure 2:**
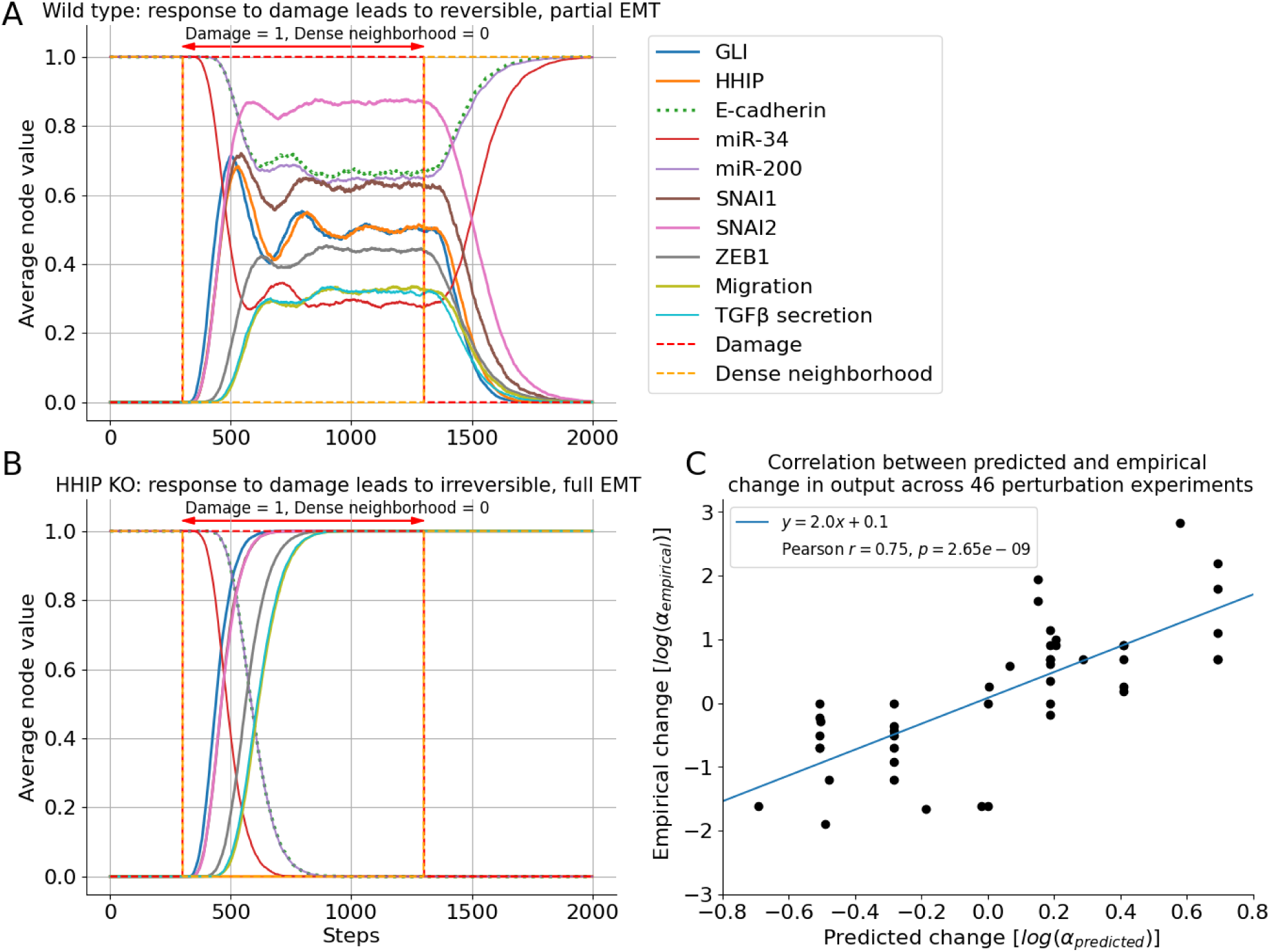
The dynamic behavior of the Boolean model reproduces the expected wild-type behavior and predicts full, irreversible EMT in the case of HHIP knock-out. Dynamic trajectories of relevant variables; average of 5000 simulations using general asynchronous update from the same initial conditions with the same perturbations. **A)** Wild-type wound healing behavior: with the introduction of the wound the GLI-HHIP negative feedback loop leads to the oscillation of the HH pathway and most EMT drivers, which results in intermediary concentrations corresponding to partial EMT. Once the wound input is removed the system reverts to the initial epithelial state. **B)** HHIP knock-out: the wound input results in a transition to an irreversible mesenchymal attractor, due to the lack of negative feedback from HHIP. **C)** Correlation between the predicted and empirical change in output for the perturbation experiments listed in Supplementary Table 3. The blue line is the fitted linear regression model.

In contrast, the same *in silico* experiment performed with a complete knock-down of HHIP (HHIP permanently set to 0) shows a dramatically different behavior, as shown in Figure 2B. As soon as the “wound” is introduced, GLI, which is lacking negative feedback from HHIP, stabilizes in the ON (1) state and induces full EMT. Remarkably, this state is not reversed by the removal of the perturbation input at time-step 1300. This mesenchymal state emerges *despite* not having TGFβ feedback loops, which would further stabilize the phenotype. Indeed, TGFβ secretion goes up due in part to the complete downregulation of miR-200, its principal antagonist.

We validated this model with 42 additional perturbation experiments from the literature that were not used to construct our network as well as 4 perturbation experiments performed in our own lab (see Supplementary Table 3). The network validation methods are described in detail in the Methods section and the experiments in a later section. Briefly, we examined whether our model reproduced each of these 46 perturbation experiments by permanently enforcing the state(s) of a (subset of) node(s) that corresponded to the gene(s)/molecule(s) perturbed in the experiment and determining the resulting stable states of the *other* nodes in the model. The output measured by the experiment (e.g., the expression level of a downstream gene) was then associated with the appropriate corresponding node in our model, and we recorded how much the measured output changed in the *in silico* perturbation experiment (as compared to a “wild-type” scenario or other baseline). As a first approximation, we determined whether the direction of change (or lack of change) in the corresponding node in our model agreed with the output observed in the perturbation experiment. In this context, our model obtained an accuracy of 91.3% (42 out of 46). Notably, 3 out of the 4 experiments where the prediction is inaccurate are experiments in which the measurement showed no significant change in the output compared to the baseline; we similarly find that our model predicts only minimal change for 2 out of these 3 experiments. Next, we quantified the average change in the associated output node in our model and compared it to the change in the experimentally measured output. The Pearson correlation between the experiments versus our simulations is r = 0.746 with p-value = 2.65x10^-9^ (See Figure 2C, analysis described in detail in Methods). These results indicate our model robustly recapitulates expected biological behavior in the context of node perturbation.

### 2. Hedgehog signaling linked to a mechanosensitive, TGFβ-responsive EMT model indicates that loss of HHIP compromises reepithelialization by destabilizing partial EMT

Next, we probed the effects of altered Hedgehog signaling on the behavior of lung epithelial cells during the wounding of a cellular monolayer by examining the interplay between EMT, mechanosensing, and proliferation in response to microenvironmental signals near the wound. To do this, we expanded a previously published model of mechanosensitive EMT, which examined the synergistic effect of low cell density, stiff extracellular matrix (ECM) and strong mitogens in driving EMT in the absence of transforming signals such as *TGFβ* or *SHH* (*30*) (Supplementary File 2). This model includes a detailed cell cycle control circuit driven by growth signaling (MAPK, PI3K/Akt, and mTORC), integrated with adhesion, contact inhibition, control of EMT, as well as apoptosis (all links and gate logic detailed/referenced in Supplementary Table 4, new nodes/links highlighted in blue). As Figure 3 shows, we first added a Hedgehog signaling module (*left, maroon module*), such that it impacts the rest of the network by promoting EMT and survival (*22*, *25*, *33*, *39*, *40*). The rest of the network, in turn, modulates the SHH signal primarily via C/EBPα, a transcription factor required for lung maturation and known to reduce GLI1 expression in lung epithelia (*41*, *42*). To model the effect of EMT on the underlying ECM, we next incorporated the feedback control of key lung alveolar ECM components (collagens, laminin, fibronectin and elastin) by matrix metalloproteinases MMP2/7/9 (*top, gray module*) (*43–45*). Together with fibronectin, these MMPs are induced by SNAI1/2, ZEB2, and/or β-catenin/TCF4/LEF1 (repressed by C/EBPα), and lead to basement membrane destruction and interstitial ECM stiffening (*28*, *46–50*). To validate this larger model and show that the output of a synchronous update approach matches key experimental data, we first identified the model’s stable cell phenotypes (attractors) (Supplementary File 2, visualized in Supplementary Figure 2). Next, we created detailed *in silico* experimental protocols to mimic the 49 validation experiments listed in Supplementary Table 5, testing whether partial knockdown/overexpression can also match these experiments (Supplementary Table 5; protocols in Supplementary File 2). Our model matched the direction of change in 47 of 49 experiments (two experiments disagree; results summarized in Supplementary Table 5, figures in Supplementary File 3; script for automated comparison, statistical tests and figure generation in Supplementary File 2).

**Figure 3.**
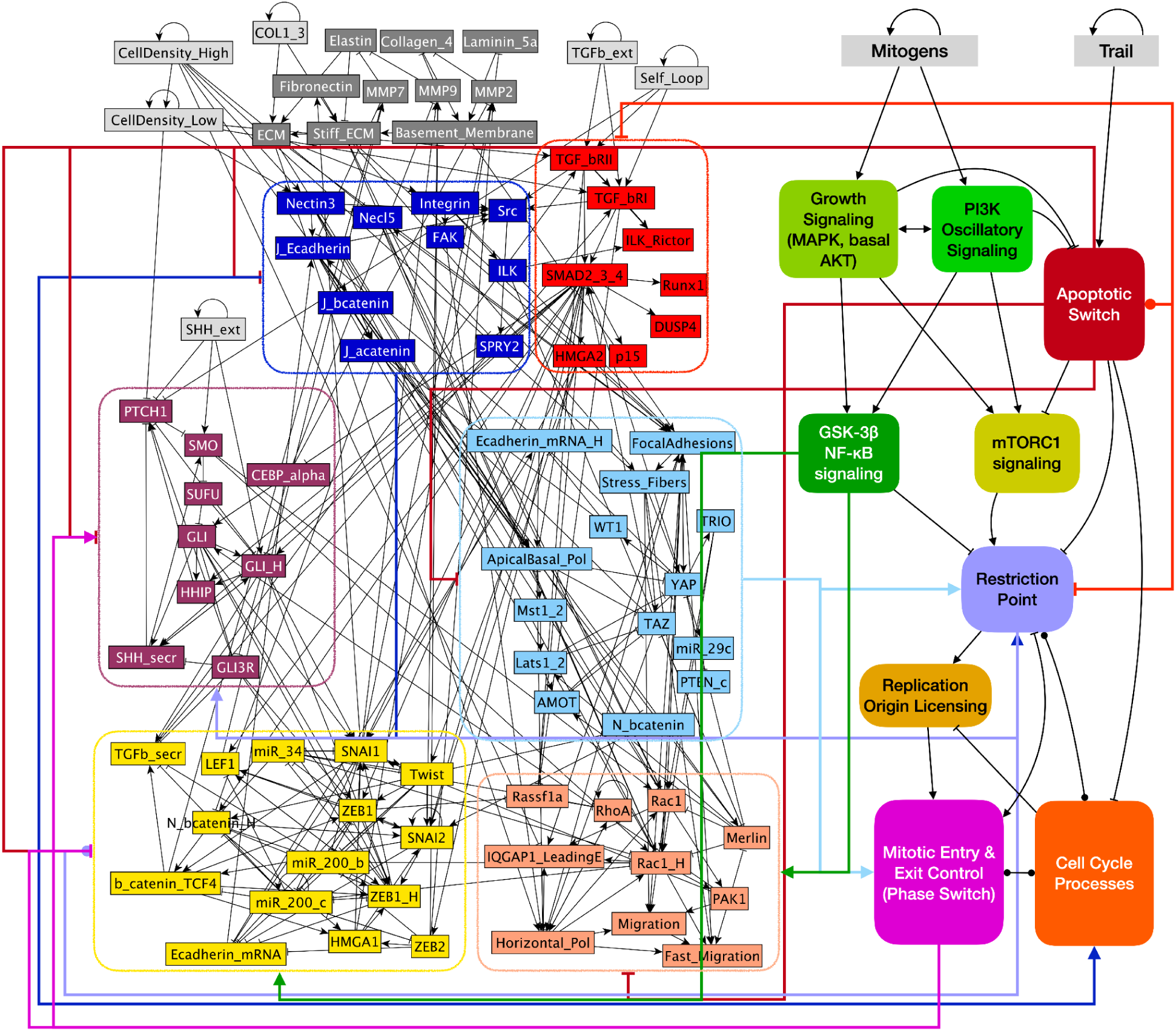
Boolean model of SHH signaling linked to mechanosensitive EMT regulation and cell cycle progression. Modular network representation of our Boolean model expanded from (*30*). *light gray*: inputs representing factors in the microenvironment of the modeled cell; *gray*: Matrix (feedback control of ECM composition; new); *blue*: Adhesion signals; *red*: TGFβ signaling; *maroon*: SHH signaling (new); *light blue*: Contact Inhibition; *yellow*: EMT switch; *light orange*: Migration; remaining modules noted in filled rounded squares are shown in detail in Supplementary Figure 1; *black links between molecules:* → : activation; –| : inhibition; *links between modules:* color: source module; → : activation; –| : inhibition; –● : complex influence. (See Supplementary Table 4 for details on all nodes and links, and Supplementary File 2 for Boolean model in .dmms (*53*), .SBML (*54*) and .BooleanNet (*55*) formats.)

To simulate the appearance of a gap or wound, we chose an initial condition that matches the state of cells prior to wounding (epithelial cell with apical-basal polarity and strong junctions, in an intact monolayer, exposed to mitogens but contact-inhibited and quiescent) (Figure 4A, *left interval;* full dynamics in Supplementary Figure 3A). The appearance of a gap was modeled by a change in the model cell’s environment from high to moderate density, mimicking a wound’s edge (Figure 4A, *middle interval*). The resulting model dynamics indicates that cells at a monolayer’s edge lose their contact inhibition, start migrating and enter the cell cycle, but also undergo partial EMT to a hybrid E/M state (as seen in (*30*)). As the wound closes and density goes up, all of these phenotypic transitions are reversed; namely the cell re-establishes its original epithelial phenotype, stops migrating and exits the cell cycle (Figure 4A, *right interval*).

**Figure 4.**
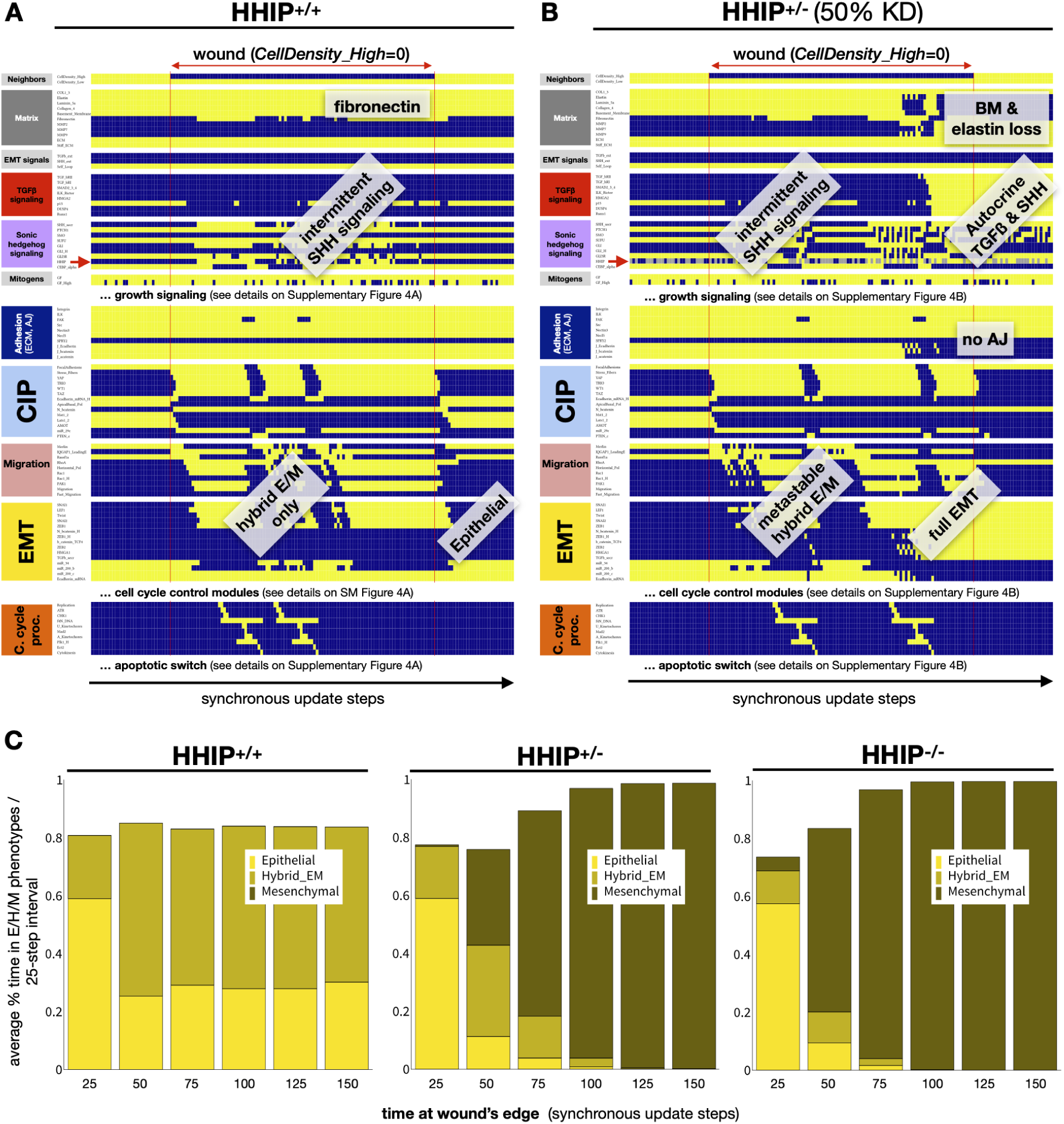
Model linking EMT to contact inhibition and cell cycle predicts that HHIP haploinsufficiency upregulates full EMT during lung re-epithelialization, leading to the loss of adherens junctions and ECM remodeling. A-B) Dynamics of the expression/activity of a select set of regulatory molecules in a cell at high confluency (first interval), exposed to the edge of a wound (middle interval), then transitioning back to high confluency (last interval) (A) wild-type (*HHIP*+/+) cells and (B) heterozygous (*HHIP*+/-) cells (HHIP is inactive in 50% of time-steps, modeling the stochastic weakening of its effects at reduced levels) exposed to 75% saturating mitogenic signals but no external Shh or TGFβ. *X axis*: time-steps (synchronous update); *y axis*: nodes organized in regulatory modules; *red circle*: HHIP. *yellow/blue/gray*: ON/OFF/forced OFF; *vertical red line:* start / end of wound; *labels*: relevant phenotype changes (Supplementary Figures 4A-B show dynamics of all modules). **C)** Stacked bar charts showing the average % time cells exposed to a gap (at a monolayer’s edge) spend in the Epithelial (*yellow*), Hybrid E/M (dark yellow) and Mesenchymal (mustard) states in consecutive 25-minute intervals (75% of saturating mitogen level, no external Shh or TGFβ). *Sample size:* 1000 independent runs; *Update:* synchronous (Supplementary Figure 3C shows results with biased asynchronous update).

Our model predicts that the loss of C/EBPα activation during the hybrid E/M phase triggers moderate GLI activation, which oscillates along with the cell cycle. GLI, in turn, upregulates HHIP, reducing the availability of secreted SHH to activate Hedgehog signaling (*24*, *51*, *52*).

The main effect of external SHH exposure on our model, as expected, is to trigger full EMT (Supplementary Figure 4). In the context of wound healing, excess SHH would lead to cells that migrate alone, lose their ability to adhere to neighbors upon wound closure, proliferate significantly less than their hybrid E/M counterparts, secrete MMPs that break down the basal lamina of alveoli, and promote interstitial ECM stiffening due to elastin loss and excess fibronectin. To test whether the loss of HHIP has a similar effect on our model, we compared the wound healing behavior of simulated wild-type cells (Figure 4A) to those with varying levels of HHIP knockdown. To mimic a cell with one functional copy of *HHIP* (heterozygous), we assumed that lower HHIP protein levels manifest as unreliable HHIP function; an effect we modeled by randomly setting the HHIP node to the off state in 50% of the time steps (Figure 4B, Supplementary Figure 3B). Our simulations predict that partial loss of HHIP triggers full EMT in a subset of cells (Figure 4B, *right*). This occurs when SHH signaling fluctuations and bursts of GLI activity allow the EMT module to commit, then remain self-sustaining due to autocrine TGFβ and SHH signaling. Moreover, the longer a single cell remains near a monolayer’s edge, the more likely it is to undergo full EMT; a process accelerated by full HHIP knockdown (Figure 4C, Supplementary Figure 3C). Overall, the predicted behavior of cells with partial loss of HHIP reproduces several epithelial pathologies observed in emphysema, including weakened barrier function, loss of junctions, reduced or arrested proliferation, and loss of the alveolar basement membrane paired with an increase in interstitial ECM stiffness due elastin loss and fibronectin accumulation (Supplementary Figure 5).

### 3. HHIP knock-down in alveolar cell line promotes EMT

To test whether the loss of HHIP results in the predicted dysregulation of epithelial wound healing, we performed an *in vitro* scratch assay with A549 cells, an immortalized alveolar cancer cell line. We examined wound closure rate and the expression of epithelial and mesenchymal marker genes (see Methods). First, we verified that GLI activity promotes wound healing in the absence of exogenous SHH, as predicted by our models. Indeed, cells treated with the GLI inhibitor GANT61 were much slower in closing the wound than control cells (Supplementary Figure 6). These results are congruent with similar experiments conducted in the same cell line published in (*56*).

Next, we repeated the scratch assays in the context of *HHIP* knockdown. We observed that cells transfected with HHIP siRNA migrated at a higher rate compared to control cells (Figure 5A), although we note that these results were not conclusive across all replicates (Supplementary Figure 7). On the other hand, gene expression of EMT markers (measured with RT-PCR) showed a clear and significant upregulation of GLI2, along with CDH2 (N-cadherin) upregulation, indicating the activation of EMT (Figure 5B). At the same time, CDH1 (the gene encoding E-cadherin) was downregulated. Interestingly, GLI1 was lower in HHIP deficient cells than in controls, suggesting that the Hedgehog pathway in this cell-type may be acting through GLI2. Remarkably, cells that were transfected with HHIP siRNA but were *not scratched* showed no significant change in the expression of the genes highlighted above, indicating that the damage is necessary for the induction of Hedgehog signaling (Supplementary Figure 8).

**Figure 5:**
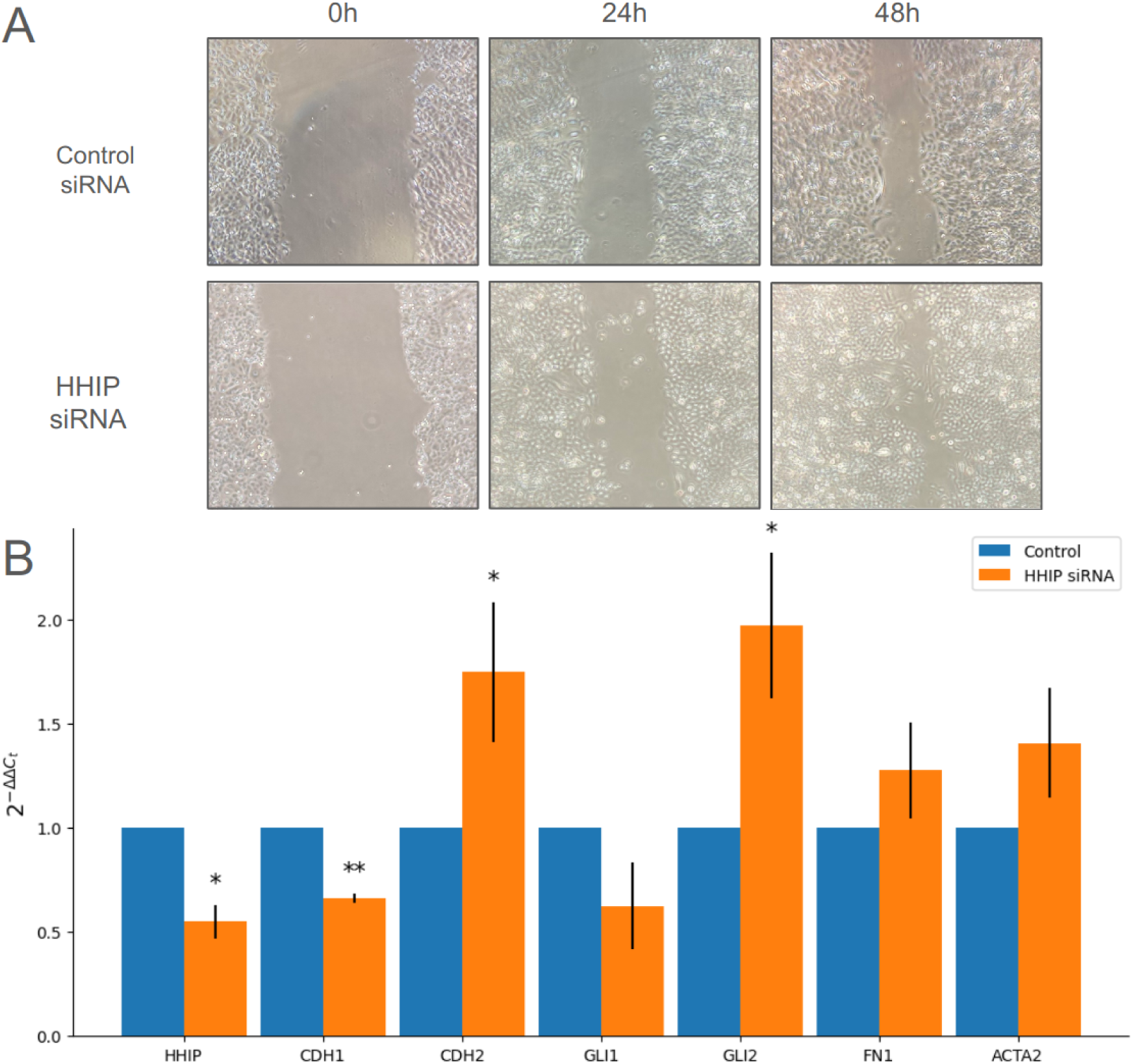
*HHIP* knock-down in A549 pulmonary epithelial cell line derived from a human alveolar cell carcinoma shows enrichment for EMT markers. **A)** representation of migration assays over time in *HHIP* KD and control (see Supplemental Figure 8), **B)** Gene expression in the cells collected from the scratch assays (3 repeats) measured by RT-PCR, represented as 2^−ΔΔ*Ct*^ values (control values normalized to 1). Significance determined by paired t-test (“ * ” : p<0.05 ; “ ** “: p<0.01)

### 4. Lung tissue and single cell gene expression data support our EMT hypothesis

We next examined whether our hypothesis is consistent with transcriptomics data collected from human COPD lung samples. First, we analyzed gene expression obtained with bulk RNA-Seq on lung tissues from the Lung Tissue Research Consortium (LTRC) (*57*). We split subjects into three groups: COPD with emphysema (135 patients), smoker controls (no emphysema, 88 patients) and never smokers (44 patients; specific criteria detailed in Methods), and ran a Mann–Whitney U test for 8 of the relevant genes shown in Figure 6A. The expression of GLI1 and GLI2 show agreement with our *in vitro* results, with GLI1 lower and GLI2 higher in COPD patients than in both control groups, respectively. Moreover, both SNAI1 and SNAI2 (GLI targets) are higher in COPD, suggesting a shift towards EMT. However, CDH1 (E-cadherin) does not show significant differences across the groups, while HHIP is upregulated in patients with emphysema compared to smoker and non-smoker controls, which is consistent with results published by Ghosh et al (*58*). It is also important to note that this data is bulk RNA-Seq on lung tissue, and thus represents a cumulative signal of multiple cell types; therefore, heterogeneity in cell type proportions across samples may be a source of bias in the results shown here.

**Figure 6:**
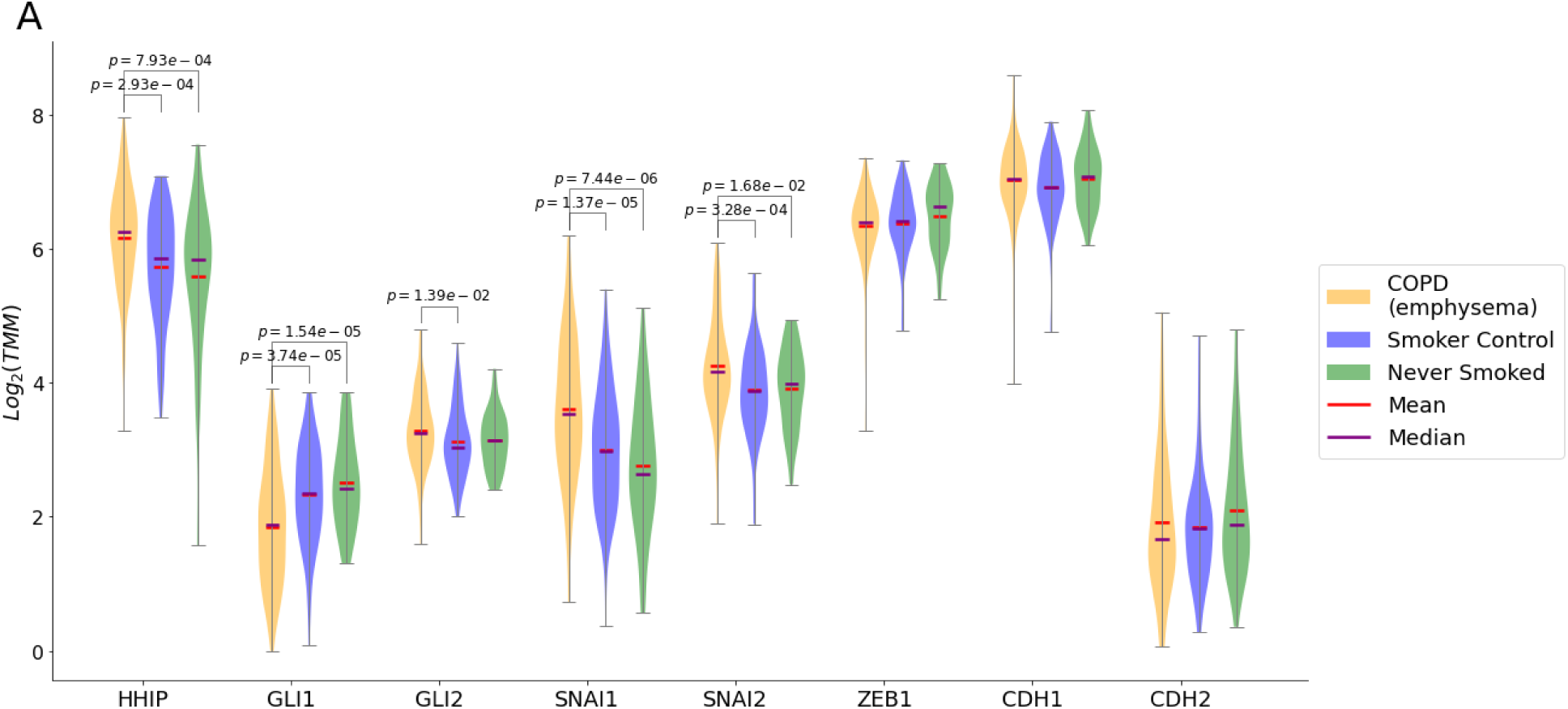
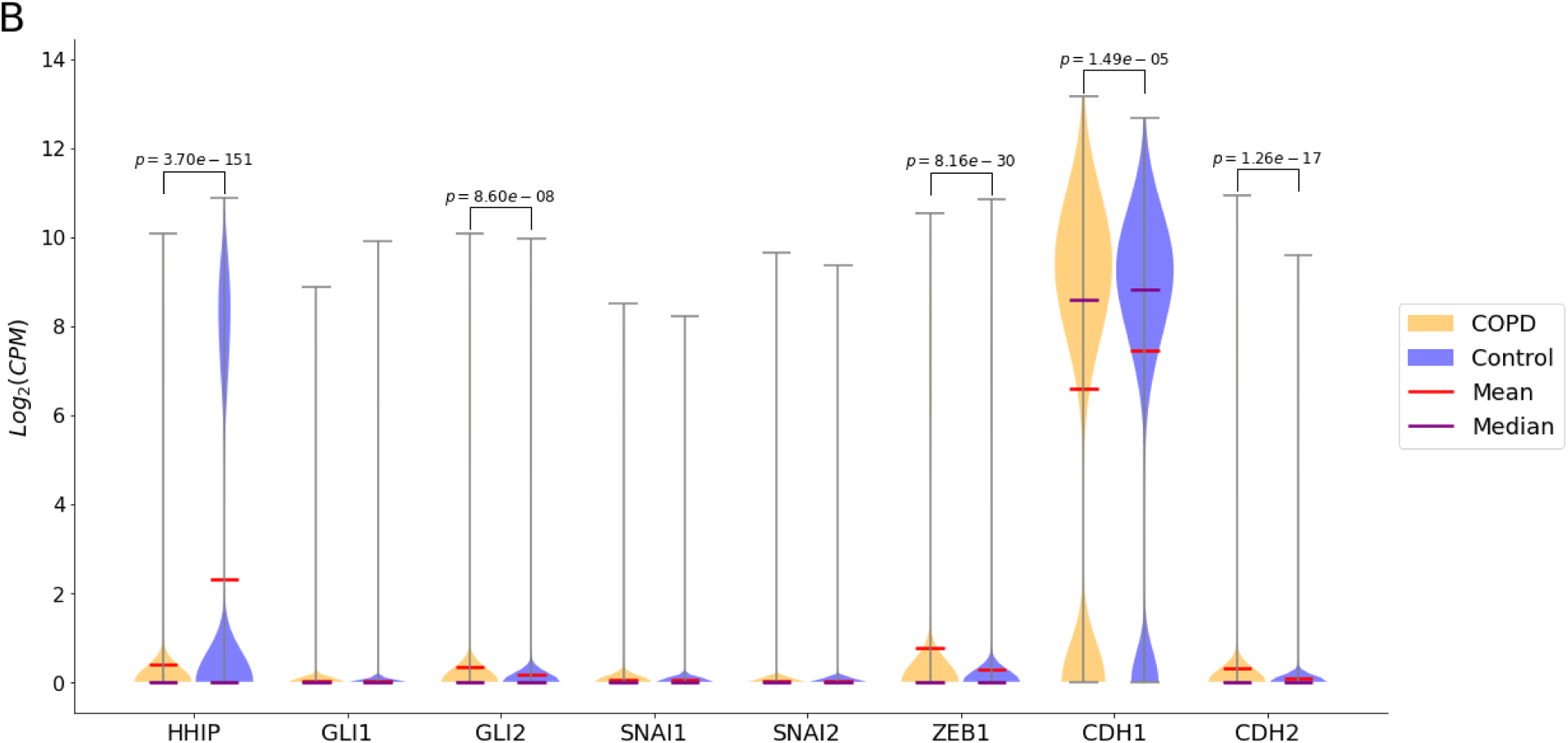
Bulk RNA-seq from lung tissue and scRNA-Seq in epithelial cells shows enrichment for EMT markers. **A)** Violin plots of bulk RNA-Seq data from the LTRC for case and two control groups. The selection criteria for the COPD/emphysema and two control groups are detailed in methods. The significance of the differences between expression levels was estimated using a Mann–Whitney U test. **B)** Violin plots of single cell RNA-Seq data of *epithelial* cells in COPD and controls. Significance levels were estimated using nonparametric Mann–Whitney U test.

To address this issue, we next looked at single cell RNA-Seq data from (*59*), analyzing gene expression of the same 8 genes in cells annotated as *epithelial*. In this case the results are clearer: HHIP expression is significantly lower in COPD epithelial cells (p=3.7e-151), while GLI2 (p=8.6e-08), CDH2 (p=1.26e-17), and ZEB1 (p=8.16e-30) expression are slightly but significantly higher. It is worth noting that the overall low expression of these genes can be explained by the fact the Hedgehog pathway is only activated in cells around acute damage. Interestingly, CDH1 shows a bimodal distribution where the nonzero peaks are relatively similar between COPD and control groups, but there are significantly more cells with zero or close to zero E-Cadherin expression in cells from COPD patients.

The combined picture from bulk lung tissue data together with single cell RNA-Seq suggests that on the tissue level there is some EMT gene upregulation likely associated with the tissue remodeling observed in the lungs of COPD patients. Focusing on epithelial cells reveals an active Hedgehog pathway with a reversed GLI2-HHIP balance in COPD compared to controls, along with the enrichment of other EMT markers. These results strongly support our proposed mechanism and modeling predictions.

## Discussion

In this work we presented a hypothesis for a mechanism that links *HHIP*, a well-established COPD susceptibility gene, and emphysema. We proposed that in the case of epithelial damage in the lungs, lack of negative feedback from HHIP allows GLI to shift the balance from partial EMT to full, irreversible EMT. Mesenchymal cells, in turn, cause slow re-epithelialization, damage the epithelial basement membrane, and stiffen the interstitial ECM, leading to tissue remodeling. Following this, we proposed a Boolean dynamic model based on canonical interactions that reproduces the phenotypes associated with normal wound healing, as well as the irreversible mesenchymal transition in response to *HHIP* knockdown. We validated this model using a series of perturbation experiments from literature and our own lab. We further examined the role of HHIP deficiency in the larger context of cell behaviors using a Boolean network expanded from Sullivan et al., and showed that misregulation of the Hedgehog pathway leads to cell cycle arrest and fibrosis at the edges of wounds. We confirmed the hypothesis *in vitro* using scratch assays on A549 cells, showing a significant enrichment of EMT markers in HHIP deficient cells attempting to close an epithelial gap. Finally, we have shown that single cell RNA-Seq data of COPD lung epithelial cells is also consistent with our model predictions and *in vitro* experiments.

EMT has been linked to COPD in multiple studies, however the role of HHIP in driving this process is not fully understood (*60*, *61*). Nonetheless, previous work has provided evidence for the involvement of the Hedgehog pathway in tissue remodeling: Li et al. (*56*) showed increased EMT in lung cancer cell lines with SHH and GLI overexpression, consistent with our dysfunctional HHIP hypothesis; Omenetti et al (*62*) showed that the EMT is regulated through the Hh pathway during biliary fibrosis, partly by the downregulation of HHIP by cholangiocytes; and Guo et al. (*63*) showed that low HHIP levels promote TGFβ secretion and the upregulation of EMT markers in human airway epithelial cells. These results are consistent with our findings. Our contribution is to offer a mechanistic explanation for the relationship between lung epithelial damage, wound healing, and an unhealthy increase in EMT due to HHIP dysfunction, leading to aberrant alveolar healing and ECM remodeling. Previous models of SHH-driven EMT do not take into account the mechanosensitive, density dependent response of epithelial cells to SHH (*31*). Moreover, the highly successful EMT model of Steinway et al. does not include important links from GLI to EMT drivers such as ZEB1 and SNAI2, which are critical in our model (see comparison to (*31*) in Supplementary Table 2). Conversely, the mechanosensitive EMT model our larger network was built upon only accounts for TGFβ as an external EMT driver, and thus does not address the aberrant behavior of cells with dysfunctional HHIP (*30*). By focusing on a microenvironment that mimics a wounded monolayer, here we not only model the effects of hyperactive HHIP on EMT, but also offer testable, mechanistic predictions for: *i)* the context (i.e., wound)-dependent need for HHIP function, and *ii)* increased GLI activation in response to tissue damage (first model) and/or proliferative signals (second model). Interestingly, our second model predicts that the faster migration of cells that commit to a mesenchymal phenotype, while beneficial to moving into a wounded area, also results in a slow or halted cell cycle. This creates difficulties in quantifying the effect of HHIP knockdown in standard scratch assays, as control cells are predicted to proliferate more to fill the gap, while HHIP knockdown cells would migrate into it faster. We posit that this prediction, while requiring careful future validation, explains our difficulty in seeing statistically significant differences in gap closure rate in our migration assays (Figure 5A).

Our study has several limitations. First, COPD is a complex disease, with multiple different mechanisms contributing to it, such as Alpha-1 antitrypsin deficiency causing lack of neutrophil elastase inhibition, chronic inflammation (*64*), oxidative stress (*65*), and tissue aging by increased rate of senescence (*66*, *67*) . Moreover, GWAS studies point to other genes besides *HHIP* that are also strongly associated with COPD susceptibility, including *IREB2*, *FAM13A*, and *MFAP2*. While the full picture of how these genes contribute to COPD is yet to emerge, this study is an important step because it provides a mechanistic explanation for a statistical association. Second, Boolean models have well-recognized limitations, such as difficulties in correctly approximating the complexity of biochemical interactions with Boolean logic and the fact that predictions are more qualitative than quantitative. Yet, the quantitative predictions of our models show high correlation with perturbation experiments from the literature, and further automated fine-tuning is possible (*68*, *69*). Time-scales of such *in silico* simulations are also difficult to match with reality because interaction times in cells vary greatly (e.g., signaling reactions vs. transcription). Asynchronous and synchronous Boolean updates both have drawbacks; while interaction times in our first, asynchronous model vary within the simulated ensemble due to the noise of asynchronous update, our second synchronous model assumes lock-step propagation of all parallel signals. However, the robustness of this second model’s behavior, regardless of update procedure, indicates that these inaccuracies do not influence the key outcomes of perturbed Hedgehog signaling in HHIP deficient cells.

In our models we treated several genes and proteins as functionally equivalent, such as the *GLI1* and *GLI2* transcription factors (merged into GLI) or the different Hedgehog signaling proteins IHH, DHH, and SHH (merged into SHH). This may be a plausible approximation in our modeling context, yet there is evidence for the distinct roles of these regulators (*70*, *71*), which we may need to account for in follow-up models. Finally the interactions in our models are collected from studies that use different animal models, cell lines and cell types (see Supplementary Tables 1-4). We limited these interactions to mammals; however the cell-type and disease specificity of interactions may play a role in *in vivo* settings that our models do not account for (e.g., many interactions were identified in cancer cells).

Our *in vitro* experiments have limitations too, for instance A549 cells are an immortalized cancer cell line, which is not ideal, because by their faulty regulatory mechanisms these cells can introduce biases into our experiments, such as abnormal levels of proliferation. In follow-up studies these experiments could be repeated in primary alveolar cells as well as other relevant cell types. Finally, most of our results rely on perturbation more akin to cutaneous wounds, while cigarette smoke damages the alveolar epithelium more by toxicity and by reactive oxygen species. Yet damage caused by cigarette smoke does lead to wound-like damage in the epithelium via necrosis, apoptosis (*72*), and ferroptosis (*73*). Thus, our wound model presents a plausible approximation of epithelial responses to damage from cigarette smoke.

A significant advantage of building mechanistic *in silico* models of targeted biological systems is that we can use methods from control theory to provide targeted interventions for reversing a pathological phenotype. These types of minimal interventions then can be tested experimentally as targeted potential treatments. An important next step is to examine and model the role of HHIP, as well as other GWAS-derived COPD genes such as IREB2 or DSP in the context of aging tissue microenvironments. COPD affects over 10% of US adults older than 65 (*74*), and its death rate rises sharply with age (8 per 100,000 for 45-55 year-olds to above 300 per 100,000 for individuals over 75) (*75*). A critical feature of aging tissues is an abundance of senescent cells, which promote oxidative stress and tissue inflammation above and beyond potential contributions from damage due to cigarette smoke. Understanding the effect of neighboring senescent cells as well as the damaging agents themselves on epithelial cells attempting to heal a wound requires an integration of our mechanistic models with models of cellular senescence (*53*, *76*, *77*). Such a model will help us map the cellular decision between entering senescence or undergoing EMT, two mutually exclusive cell fates (*78*, *79*) that are more common in COPD lung epithelia, but compromise the process of re-epithelialization in different ways. By expanding our model to integrate DNA damage response and senescence, we will approach the type of synthesis required to reproduce most cell-autonomous responses observed in a COPD context. As COPD is an independent risk factor for lung cancer (*80*), mutant versions of these models have the potential to link COPD-driven cell behaviors to the emergence of cancer hallmarks involving EMT/MET, proliferation, resistance to growth suppressors (e.g., TGFβ) or apoptosis. Finally, using our models as internal drivers of the behavior of cells in a multi-scale model of alveolar tissue homeostasis, its breakdown in COPD and the effect of aging on COPD progression could help uncover effective interventions to halt the disease.

## Methods

### Boolean models

A *Boolean network* can be represented by a graph *G* consisting of *N* nodes and *E* directed edges. Each node has a Boolean *state* (1 or 0, also referred to as ON or OFF), which can change in time. The state of each node is determined by a unique logical function (rule), which determines how the node responds to the states of its regulators (parent nodes). The state of the *system* at a given time point is the collective state of all of its constituent nodes.

*Update schemes* determine the order in which the states of the nodes are updated. Using a synchronous update scheme, all nodes are updated at the same time, such that the state of the system at time *t* fully determines the next state of the system at time *t+1*. This leads to deterministic trajectories, where the emergent dynamics of the system only depend on the initial configuration. Using a general asynchronous update scheme randomly selects a single node to update at each time step, allowing repetition.

The *attractors* of a Boolean system represent the long-term equilibrium states of the system dynamics. Fixed-point attractors (also known as steady states) are states in which all node functions are satisfied and updating nodes no longer changes the state of the system. Attractors can also be limit cycles or *complex attractors*, which, instead of a single state, are a set of states that the system keeps visiting in a loop indefinitely. Complex attractors emerge due to negative feedback loops, and represent a subset of all possible states, in which the *state transition graph* (STG) is strongly connected (all states can be reached from all other states). When in a complex attractor, a system simulated with general asynchronous update follows a random walk in this strongly connected subset of the STG. In practice this means that nodes under conflicting regulatory constraints never settle to a single state but oscillate at rates determined by their regulators and the update scheme. Due to this stochasticity, it is difficult to map a complex attractor to a biological phenotype. However, we can estimate the expected value of each node by averaging the states of a large ensemble of Boolean networks simulated with the same set of initial conditions (and perturbations). When the average value of a node settles in time to a stable expected value between 1 and 0 we call this a *dynamic equilibrium*. The average value of a node can be interpreted as an intermediary concentration of the corresponding biological variable. Different perturbations to the system can change the average node values, and we can use this to validate models against perturbation experiments.

### Simulation of Boolean models and attractor analysis

The ensemble simulations of the Boolean models were carried out using the following packages:

● The BooleanNet python library(*55*): https://github.com/ialbert/booleannet
● Boolean2PEW(*68*), a modified version of BooleanNet capable of simulating probabilistic edge weights and more advanced noise perturbations: https://github.com/deriteidavid/boolean2pew
● Cubewalkers, a cuda (GPU) based python library which makes the simulation of large ensembles of Boolean models possible within a short time(*81*): https://github.com/jcrozum/cubewalkers
● *dynmod*, a Haskell package that simulates synchronous as well as asynchronous Boolean dynamics in networks with many inputs, focusing on automated, modular attactor-to-phenotype translation: https://github.com/Ravasz-Regan-Group/dynmod (*53*).

In addition:

● AEON.py(*82*) was used for the attractor detection: https://github.com/sybila/biodivine-aeon-py
● pystablemotifs(*83*) for the stable motif analysis: https://github.com/jcrozum/pystablemotifs

Data for Figures 3-4, Supplementary Figures 2-7 and Latex Source file for Supplementary Table 4 can be generated using the files in Supplementary File 2 as follows:

dynmod COPD_EMT_CellCycle_Apoptosis.dmms -g -t -s

dynmod COPD_EMT_CellCycle_Apoptosis.dmms -e COPD_Figures.vex

- ● To re-run dynmod’s attractor detection (provided as ‘COPD_EMT_CellCycle_Apoptosis_attractors_a200_25_2.0e-2.csv’ in Supplementary File 2), remove the comments in front of the ‘Sample:’ line in ‘COPD_Figures COPD_Figures.vex’ and place them in front of ‘Read:’ .

#### Automated Validation of the first Boolean model

We developed a method for quantifying the performance of Boolean models that is also capable of handling oscillations and complex attractors. This validation approach is based on the method used in Maheshwari et al. (*84*), where a perturbation experiment is reproduced *in silico*, using the Boolean model and the resulting emergent attractor is compared with the experimental phenotype. In our case, if a change occurs in an output variable as a result of the perturbation we compare the direction of the change to the “wild-type” phenotype (or other baseline phenotype) and verify that the direction of the change agrees with the experiment.

To begin, we organize a set of perturbation experiments, as shown in Supplementary Table 3, and use these as an input to our validation algorithm. Our validation algorithm then calculates a score by performing an *in silico* simulation of each of the experiments described in this table and comparing the output of the simulation against the expected result:, i.e. the direction of change

in the output node is compared to the direction of change in the empirical measurement. Supplementary Table 3 includes the following information. The square brackets note the column name in the table:

1. experiment ID for internal use [exp_id]
2. pubmed link [pubmed link]
3. reference [reference]
4. a relevant quote from the reference (for human verification, often citing figures in the reference) [relevant quote]
5. the name(s) of the *perturbed node(s) in the reference* [perturbed molecule in ref]
6. the corresponding node name(s) in our model [perturbed node in model]
7. the *direction* of the perturbation(s) (1 - overexpression, 0 - knock-down). If there are multiple perturbed nodes, the format will be a list of 1s and 0s separated with commas and ordered in the same way as the nodes specified in column 6. **[perturbation type]**
8. the name(s) of the output variable(s) in the reference (can be a gene or a phenomenon like migration) [downstream change in (ref)]
9. **the name of the corresponding output node(s) in our model** [downstream node in model]
10. **the direction of change of the output variable in the experiment** (1 - increased, 0 - decreased) [change direction]
11. the amount of change (descriptive) [degree (descriptive)]
12. change multiplier α_*empirical*_ (=1 - no change, <1 - decrease; >1 - increase) [**degree multiplier]**
13. **experiment ID** of the baseline - this can be another experiment or a hard-coded initial condition, like the epithelial state or the wild-type wound-healing state. [comparison exp_id]
14. cell type that was used in the experiment [comparison exp_id]
15. comments [comments]

The items highlighted with **bold** font above are used to perform the validation based on the following steps:

1. We initiate a Boolean ensemble simulation with the same initial conditions as the *baseline* experiment (column 13). This simulation should already exist in memory. States such as the epithelial and mesenchymal phenotypes or wild-type wound healing are pre-computed.
2. We permanently fix the value(s) of the node(s) specified in column 6 to the corresponding state(s) specified in column 7.
3. We run the simulation of an ensemble of *n* networks for *T* steps (in our case we set n=3000, T=2500). We run the simulation until all nodes have reached a steady state or an oscillating dynamic equilibrium.
4. We generate a histogram of the output node(s) (column 9) values across the last 1000 time steps i.e. the tail of the simulation (averaged across the ensemble). This is a normal distribution centered on the average node value.
5. We perform a standard t-test between the equilibrium distribution of the output node(s) in the perturbed simulation vs. the baseline simulation.
6. We calculate the *predicted* change multiplier by the following formula: 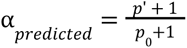 where α_*predicted*_ is the predicted change multiplier, *p*’ is the output-node average from the perturbed system, and *p*_0_ is the baseline output-node average. We add 1 to both the numerator and the denominator, so the in the case where there is no change, then α_*predicted*_ will equal 1, and if *p*_0_ = 0 the denominator is not 0.
7. If the difference between the baseline and the perturbed output average is significant (p<0.05), it’s in the same direction as in the experiment (increase or decrease; column 10) we count it as a qualitatively accurate prediction. It also counts as an accurate prediction if there is no change in the output node in the real experiment and the difference in the simulated outputs is not significant.
8. Finally we calculate the Pearson correlation between the logarithm of the empirical change multipliers (*log*(α_*empirical*_ + ε)) and the predicted change multipliers *log*_*pred*_(α + ε). One may set ε << 1 if there are cases of α = 0. Our data has no such cases, so in the calculations ε = 0 .

### Validation of the second (large) Boolean model against partial knockdown/over-expression

To test whether the model can reproduce the effects of partial knockdown/overexpression (which is a more realistic scenario for most perturbation experiments), and to gain deeper insight into the effects of each perturbation on the larger model, we created a “*in silico* protocol” for the *dynmod* program (in Supplemental File 2). These validation experiments start from an epithelial cell state, and compare the average behavior of 1000 cells at a wound’s edge over a period of 200 time-steps to cells affected by the node perturbations listed in Supplementary Table 5. As both knockdown/over-expression and the external conditions (such as growth factor levels) are non-saturating, the simulations produce a heterogeneous mix of responses; we use t-tests to compare control vs. perturbed cell responses.

To run the validation experiments for Supplementary File 3, run (∼1 hour on 2022 Mac Studio; 20-core CPU): dynmod COPD_EMT_CellCycle_Apoptosis.dmms -e Validation_Table_COPD_EMT_CellCycle_Apoptosis.vex

The python script ‘Large Model Validation Script.py’ included in Supplementary File 2, placed in the same folder as ST5 - Large model validation.csv as well as the _EXP directory where all simulation results are saved, will perform t-tests on all model output nodes and phenotypes listed in Supplemental Table 5. It also auto-generates the figures in Supplementary File 3.

### HHIP siRNA transfection, Scratch Assay and Quantitative PCR

A549 cells were cultured in 1X DMEM and seeded in a 6-well plate at a density of 4.5 x 10^5^ cells per well and allowed to reach 70% confluency before transfection. siRNA knockdown of HHIP was performed using ON-TARGET plus SMARTpool Human HHIP siRNA (Dharmacon, Cat#J-013018-09-0010) for the experimental group, alongside control siRNA (ON-TARGETplus Non-targeting Control pool, Dharmacon, Cat# D-001810-10- 05) for the control group with Lipofectamine RNAimax transfection reagent. Transfection was conducted on 60-70% confluent A549 cells using 50 pmol per well of siRNA. Scratching was performed after 48 hrs of transfection by scraping the cell layer in a straight line using a 1mm pipette tip perpendicular to the surface, maintaining contact with the bottom of the well to remove the cell layer without applying excessive pressure. After scratching, washing with 1X PBS was conducted to eliminate detached cells, followed by replenishing with 1X DMEM medium containing 10% FBS. Images of the same area were captured at 0 hrs, 24 hrs, and 48 hrs post-scratching and treatment using Olympus microscope at 4X magnification. After 48 hrs, cells were harvested for RNA extraction using the Zymo Research Quick RNA mini prep kit (Zymo Research, #R2052), followed by RT-qPCR analysis utilizing gene probes (Supplementary Table 6) to assess the expression of target genes. Reverse transcription was done using extracted RNA from cells by High-Capacity cDNA Reverse Transcription Kit (4368813, Thermo Fisher Scientific) as per manufacturer’s protocol. Real-time qPCR was performed to validate expression change on potential targeted genes upon knockdown by Applied Biosystems TaqMan Fast Advanced Master Mix (Thermo Fisher Scientific, #4444557).

### qPCR data analysis

We used an in-house python script for the calculation of the gene expression based on the Ct values measured in the different repeats. In all cases we used GAPDH as the baseline housekeeping gene. We calculated the delta-delta Ct values for each gene in case and control experiments and estimated the statistical significance of the differences using a paired t-test.

### Migration rate calculation

We measured the gap distances on scratch images manually at three or four points (depending on the shape of the scratch) using GIMP (GNU Image Manipulation Program) 2.1 and normalized the pixel counts with the diameter of the eyepiece circle in the image. The *migration rates* are calculated for the different time-points (24 or 48 hrs) as the gap distance at t=0, the moment of scratching, (d0) minus the gap distance at time t (dt), divided by d0. The difference in migration rates between case and control plates, measured at different points along the scratch were then compared and the statistical significance of the differences estimated using a standard t-test.

### Lung tissue bulk RNA-Seq data analysis

We selected the COPD group and the two control groups from the LTRC cohort based on the following criteria:

- We excluded subjects (N=35) that were under ongoing chemotherapy or radiation therapy within one year of time of the sample collection.

#### COPD with emphysema group (N=135)

- Post-bronchodilator GOLD spirometry stage>= 3
- Indicated as having COPD as defined by (*58*)
- Percent of lung low attenuation areas (LLA) less than -950 Hounsfield units (HU) >= 10
- Smoker (current or former)
- Pack years > 5

#### Smoking controls (N=88)

- Smoker (current or former)
- Indicated as not having COPD as defined by (*58*)
- Percent of lung low attenuation areas (LLA) less than -950 Hounsfield units (HU) < 10
- Pack years > 5

#### Non-smoking controls (N=44)

- Never smoker
- Indicated as not having COPD as defined by (*58*)
- Percent of lung low attenuation areas (LLA) less than -950 Hounsfield units (HU) < 10

### Single-cell RNA-Seq data analysis

We downloaded publicly available COPD single cell RNA-Seq data from Adams et al. (<u>GSE136831</u>). We selected the subset of cells annotated as *Epithelial* (CellType_Category) and conducted simple CPM (counts per million) normalization of the raw RNA counts in the epithelial subset using the *bioinfokit* [https://github.com/reneshbedre/bioinfokit] python package.

The case (COPD, n=17) and control (Control, n=15) groups were selected based on the Disease_Identity variable. We compared the difference in expression between the two groups for the genes of interest using the nonparametric Mann–Whitney U test.

## Code availability

Code reproducing the simulations, validation, and data analysis is available at:

https://github.com/deriteidavid/models_of_hhip_driven_emt_in_copd https://github.com/Ravasz-Regan-Group/dynmod

## Supplementary Material Legends

**Supplementary Table 1** - node list of the smaller EMT model with the description of node states and Boolean rules

**Supplementary Table 2** - edge list of the smaller EMT model, documented with references, interaction type, and cell type used in the reference

**Supplementary Table 3** - perturbation list used for validation, see Methods for a detailed description of the columns

**Supplementary Table 4** - Large (107-page), formatted and referenced table describing the biological evidence behind each node, link and logic gate of the large Boolean model, organized by regulatory module. *Black text:* previously published node and link descriptions; *blue text:* new to in the current model; *olivegreen text:* minor modification from previous model.

**Supplementary Table 5** - Manual validation summary for larger Boolean model, including references to each perturbation experiment, key elements of these experiments, agreement / disagreement with the simulation result and the names of *in silico* experiments that simulate them, as listed in the *in silico* protocols included in Supplementary File 2).

**Supplementary Table 6:** All the probes were ordered from IDT (Integrated DNA Technologies) From PrimeTime™ Predesigned qPCR Assays.

**Supplementary File 1** - attractor analysis for the smaller model

**Supplementary File 2 -** Files read by *dynmod* to simulate the large model’s behaviors. These include the model in .dmms format (as well as .BooleanNet and .SBML formats, namely COPD_EMT_CellCycle_Apoptosis.dmms, COPD_EMT_CellCycle_Apoptosis_Fine.booleannet, the “in silico protocol” files for reproducing all figures (COPD_Figures.vex), re-running the validation experiments (Validation_Table_COPD_EMT_CellCycle_Apoptosis.vex) and running a python script that performs all validation statistical tests, generating a results list and regenerating Supplementary File 3 (Large Model Validation Script.py, ST5 - Large model validation.csv).

**Supplementary File 3 -** Figures showing the results of manual validation summarized in Supplemental Table 5.

## Author Contributions

D.D. - conceptualization, first (smaller) Boolean model development, Boolean simulations, validation of the models, processing of RT-PCR data, processing of scratch assays images, RNA-Seq data analysis (bulk and single cell), visualization (graphical abstract, figures 1, 2, 5, 6, S7, S8), writing the manuscript.

W.J.A - scratch assays, RT-PCR experiments, processing of RT-PCR data, visualization (figure S6) writing the manuscript.

X.Z. - supporting W.J.A., writing the manuscript.

K.G. - supporting D.D., validation of the models, writing the manuscript.

E.R. - conceptualization, second (larger) Boolean model development, Boolean simulations, validation of the models, visualization (figures 3, S1 to S5), writing the manuscript.

E.S. - supporting D.D., sponsoring the experiments, writing the manuscript.

## Grant Support

D.D. and K.G. were supported by R01HL152728 and R01HL155749.

X.Z. was supported by R01HL148667 and R01HL127200.

E.R.R. was supported by the College of Wooster’s Endowed Fund for the Life Sciences.

E.K.S. was supported by R01HL133135, P01HL114501, and R01HL152728.

## Conflicts of interest

In the past three years, EKS has received grant support from Bayer and Northpond Laboratories.

## Acknowledgements

The authors would like to thank Leonardo Martini, Margherita De Marzio, Enrico Maiorino, Pete Regan, Camila Lopes-Ramos, and Min Hyung Ryu for their valued contributions and advice.

## Supplementary Figures

**Supplementary Figure 1:**
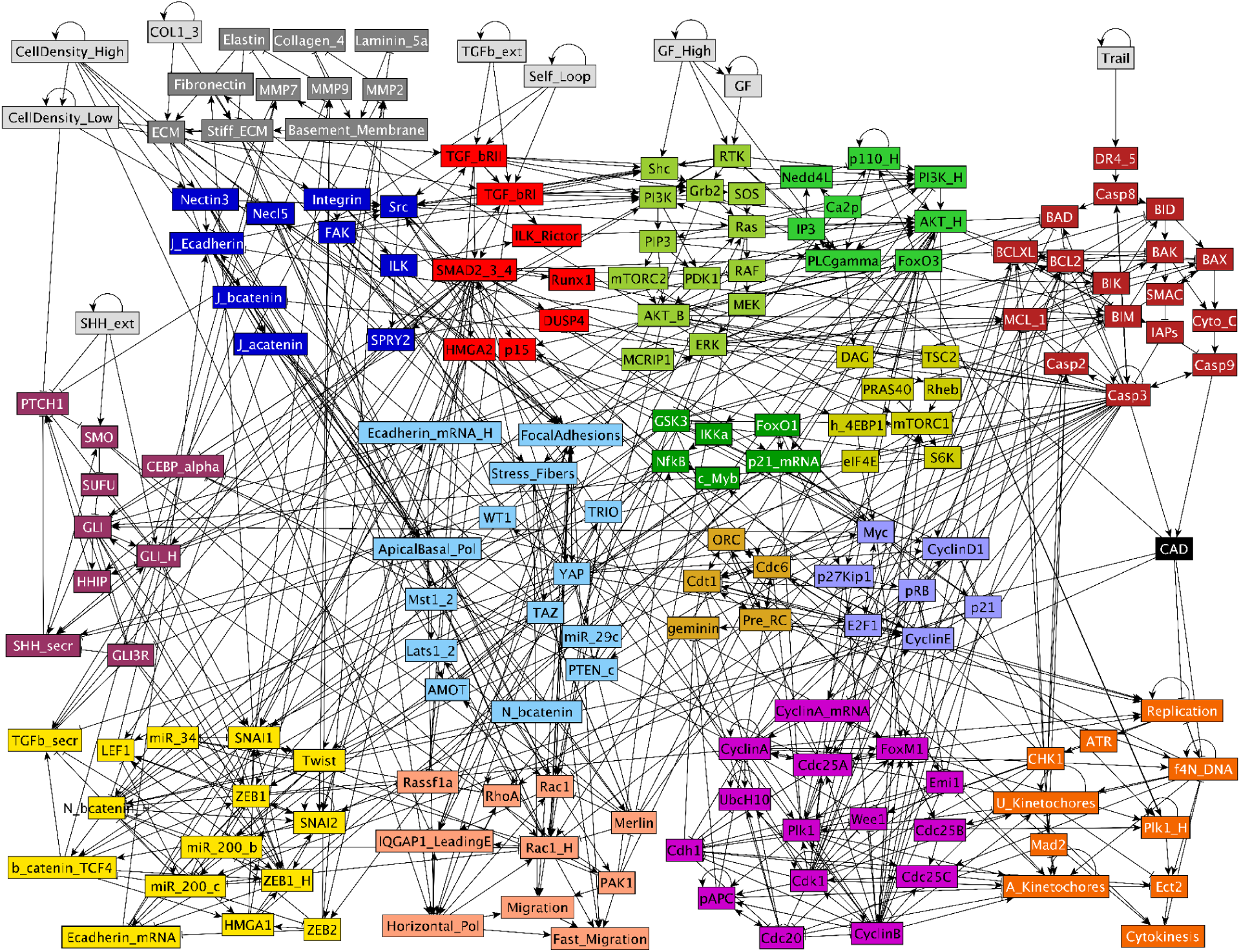
Full Boolean network model of SHH signaling linked to mechanosensitive EMT regulation and cell cycle progression. Full network representation of our Boolean model expanded from (*30*). *light gray*: inputs representing factors in the microenvironment of the modeled cell; *dark gray*: Matrix (feedback control of ECM composition; new); *dark blue*: Adhesion signals; *red*: TGFb signaling; *soft maroon*: Shh signaling (new); *light blue*: Contact Inhibition; *yellow*: EMT switch; *pink/light orange*: Migration; *light/dark green*: Growth factor signaling & NF-κB; *mustard*: mTORC1 signaling; *brown*: Origin Licensing; *lilac*: Restriction Switch; *magenta*: cell cycle phase switch; *orange*: cell cycle control; *dark red*: Apoptosis; *black links between molecules:* → : activation; –| : inhibition (see Supplementary Table 4 for details on all nodes and links, and Supplementary File 2 for Boolean model in .dmms, .SBML and .BooleanNet formats.)

**Supplementary Figure 2:**
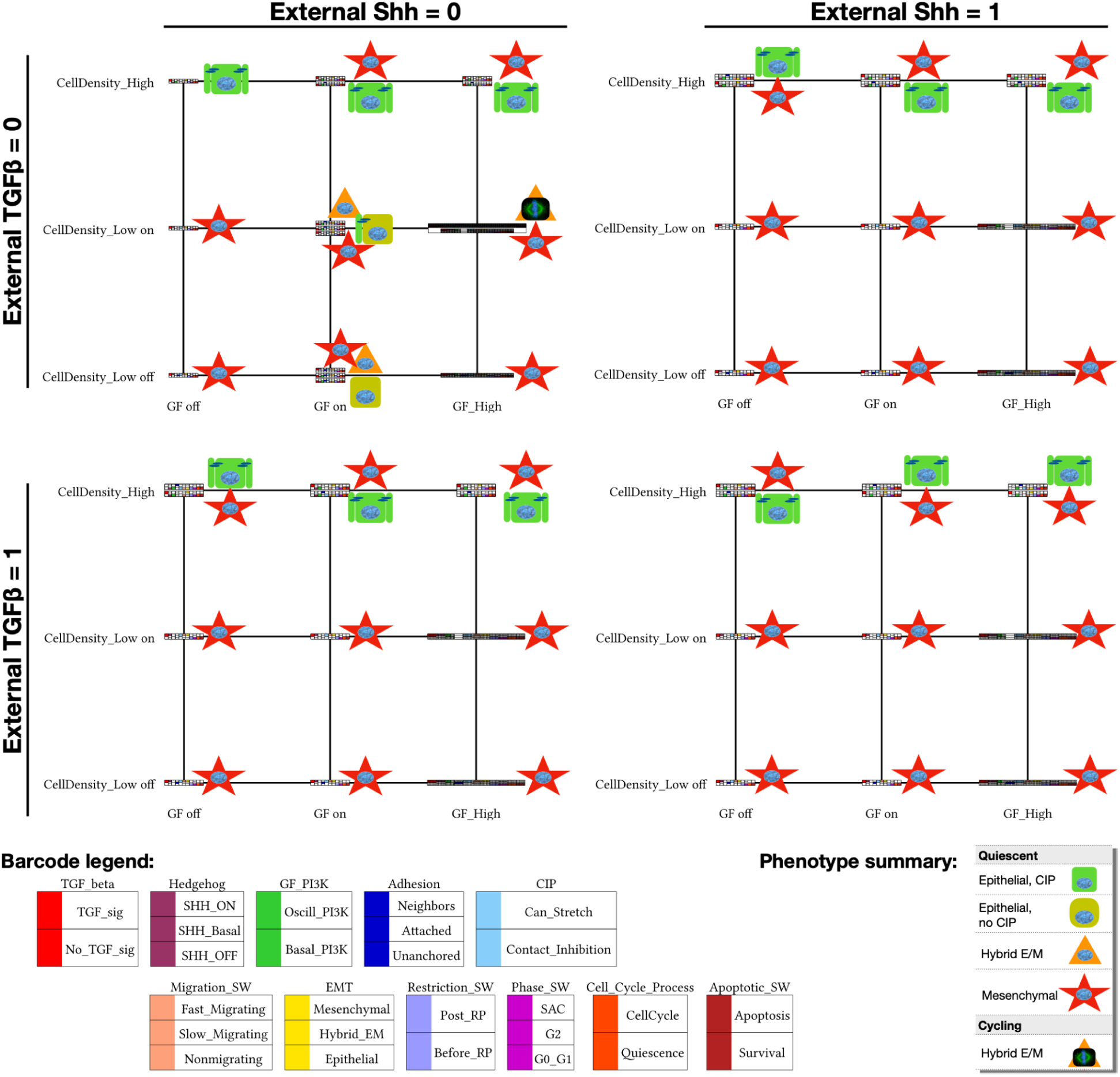
Relevant attractors of the large Boolean model, organized by external environment. Relevant model cell states (synchronous attractors) detected in every combination of no/low/high growth-factor (*x* axis), no/low/high cell density (alone / monolayer’s edge / full confluence with no room to stretch; *y* axis), absence/presence of external Shh (*left/right*), and absence/presence of external TGFβ (*top/bottom*). Barcodes representing each attractor were derived by comparing the expression of nodes in each relevant module to a predetermined molecular signature known to represent a cell phenotype (e.g., apoptosis vs. survival) and encoded in the model’s .dmms file (barcode legend, *bottom left*). Oscillatory phenotypes have expanded barcodes that mark the transitions their regulatory switches undergo during the cycle. ***Note:*** attractors representing apoptotic cells, stable in every environment-combination, were filtered out to reduce clutter; as were quiescent polyploid cell states (reachable due to cell cycle errors only, but stable in quiescence). Visual summaries of overall cell states indicate epithelial (*green*) vs. hybrid E/M (*orange*) vs. mesenchymal (*red*) phenotypes, as well as quiescence (*round nucleus*) vs. cell cycle (*mitotic spindle image*). ON/OFF state of the network in every detected attractor included as Supplementary File 4.

**Supplementary Figure 3:**
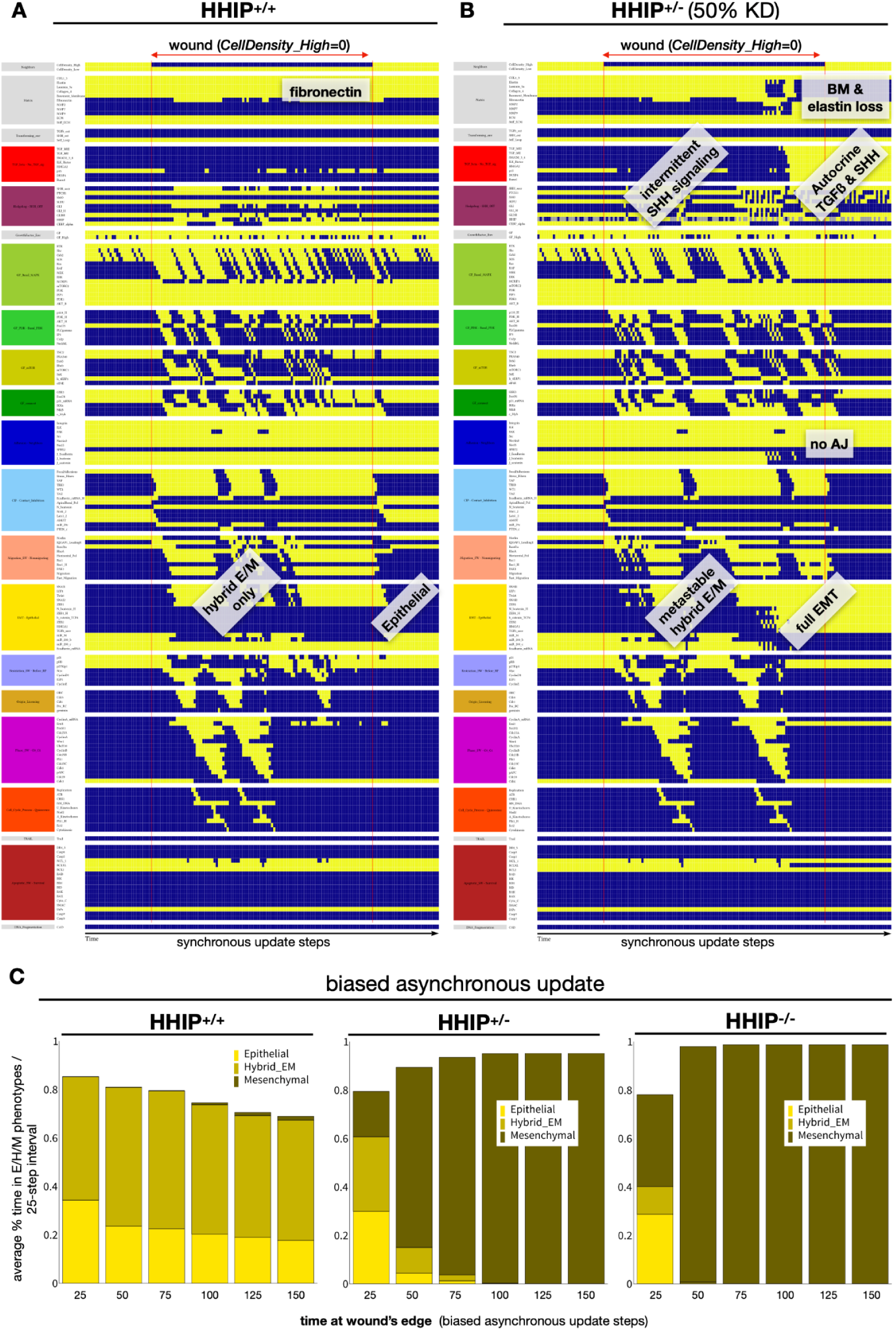
Full network dynamics for Figure 4A-B (Boolean model predicts that HHIP haploinsufficiency upregulates full EMT during lung re-epithelialization). **A-B)** Dynamics of the expression/activity of all regulatory molecules in a cell at high confluency (first interval), exposed to the edge of a wound in (middle interval), then transitioning to high confluency (last interval) (A) wild-type (HHIP+/+) and (B) heterozygous HHIP+/- cells (HHIP is inactive in 50% of time-steps, modeling the stochastic weakening of its effects at reduced levels) exposed to 75% saturating mitogenic signals but no external Shh or TGFβ. *X axis*: time-steps (synchronous update); *y axis*: nodes organized in regulatory modules; *yellow/blue/gray*: ON/OFF/forced OFF; *vertical red line:* start / end of wound; *labels*: relevant phenotype changes. **C)** Figure 4C ***results with biased asynchronous update:*** Stacked bar charts showing the average % time cells exposed to a gap (at a monolayer’s edge) spend in Epithelial (*yellow*), Hybrid E/M (dark yellow) and Mesenchymal (mustard) states in consecutive 25-minute intervals (75% of saturating mitogen level, no external Shh or TGFβ). *Sample size:* 1000 independent runs; *Update:* biased asynchronous.

**Supplementary Figure 4:**
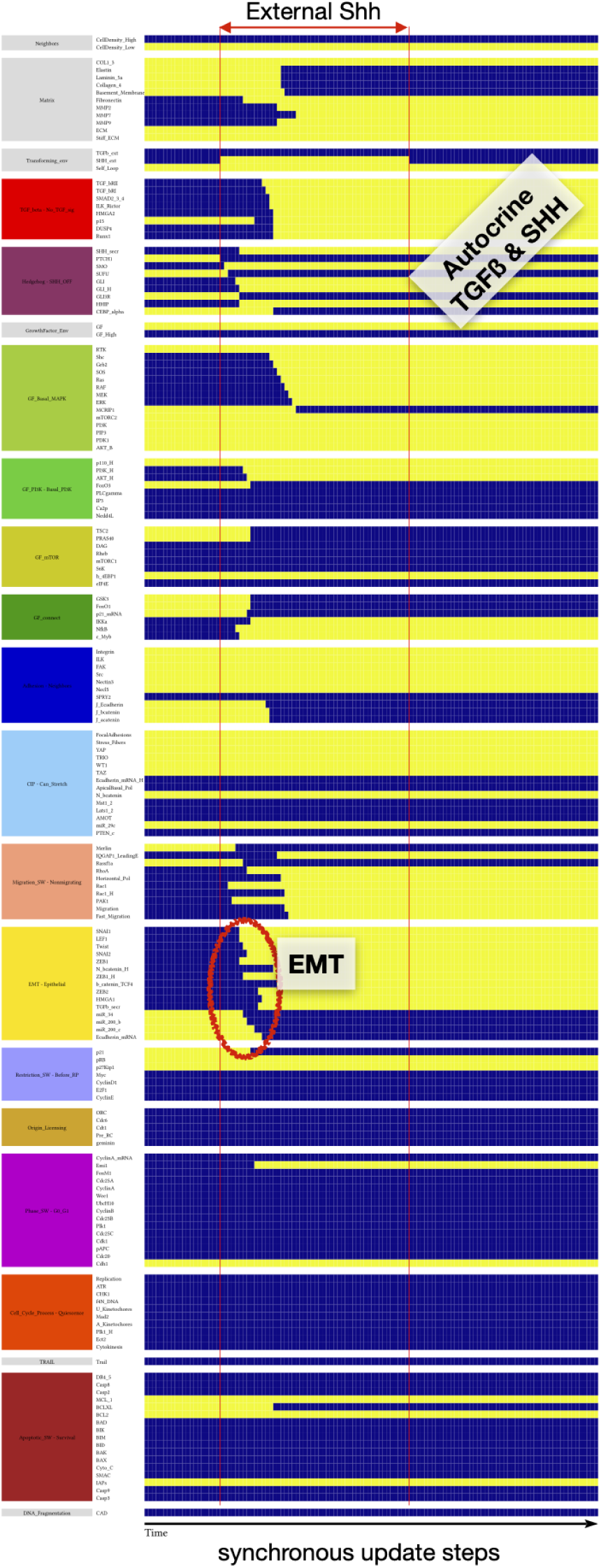
External Shh triggers EMT. Dynamics of the expression/activity of all regulatory molecules in a cell at a monolayer’s edge, exposed external Shh for 100 time-steps (mitogenic signals are sufficient for survival but not cell cycl;e entry; no external TGFβ). *X axis*: time-steps (synchronous update); *y axis*: nodes organized in regulatory modules; *yellow/blue*: ON/OFF; *vertical red line:* start / end of Shh pulse; *red oval:* timing of EMT; *labels*: relevant phenotype changes.

**Supplementary Figure 5:**
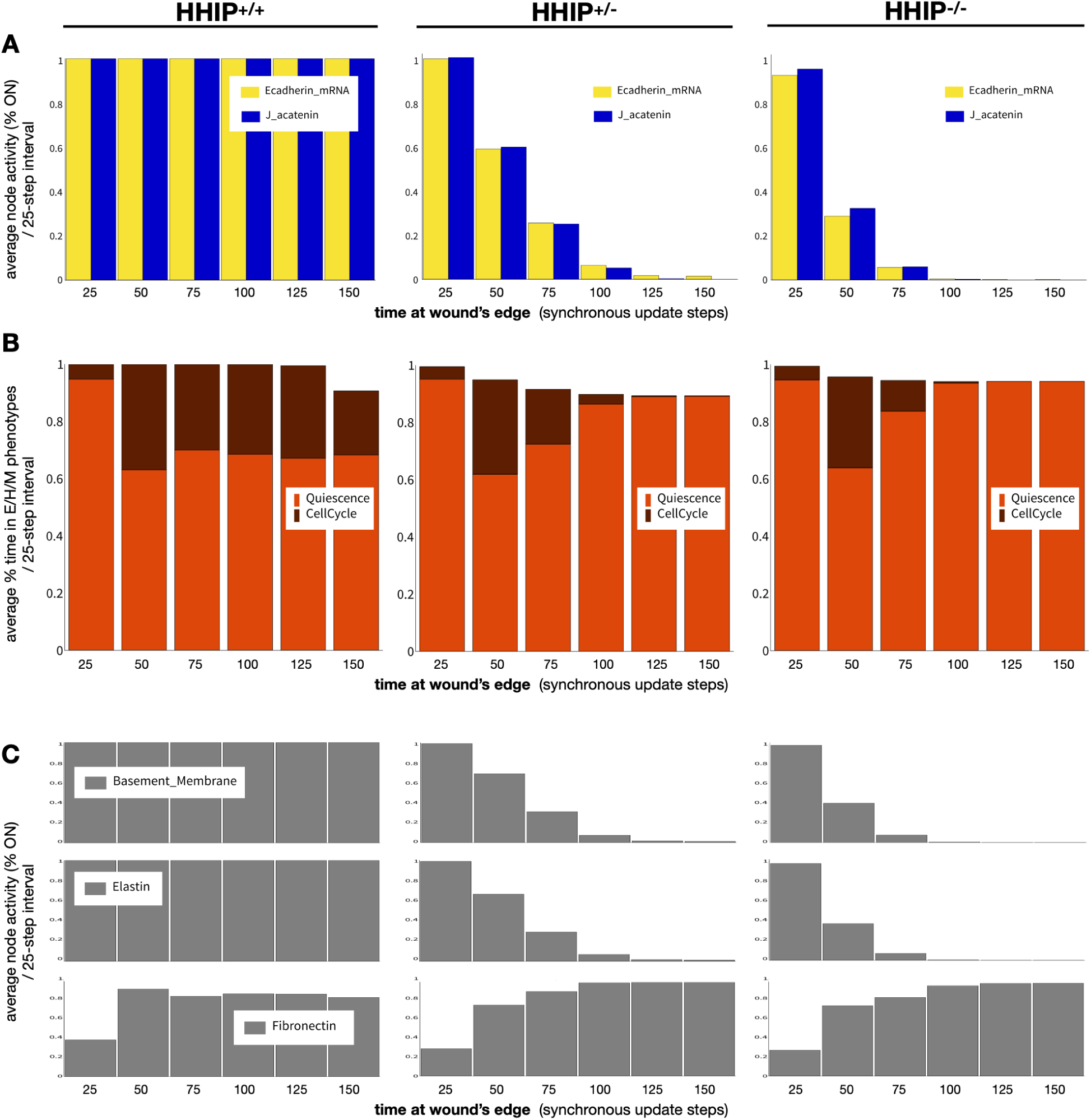
HHIP knockdown reduces cell-cell adhesion, proliferation, and stiffens the ECM while degrading the alveolar basement membrane. **A-C)** Wild-type (HHIP^+/+^; *left*), heterozygous HHIP^+/-^ (*middle*) and HHIP-null (HHIP^-/-^; *right*) cells exposed to 75% saturating mitogenic signals but no external Shh or TGFβ. **A)** Average activity (% time ON) of the E-cadherin mRNA node (yellow) and junctional ɑ-catenin (*blue*); **B)** stacked bar charts showing the average % time cells spend in Quiescence (*red*) vs. in cell cycle (*dark red*); and **C)** average levels (% time ON) of the Basement Membrane (*top*), Elastin (middle) and Fibronectin (bottom) nodes in consecutive 25-minute intervals. *Sample size:* 1000 independent runs; *Update:* synchronous.

**Supplementary Figure 6:**
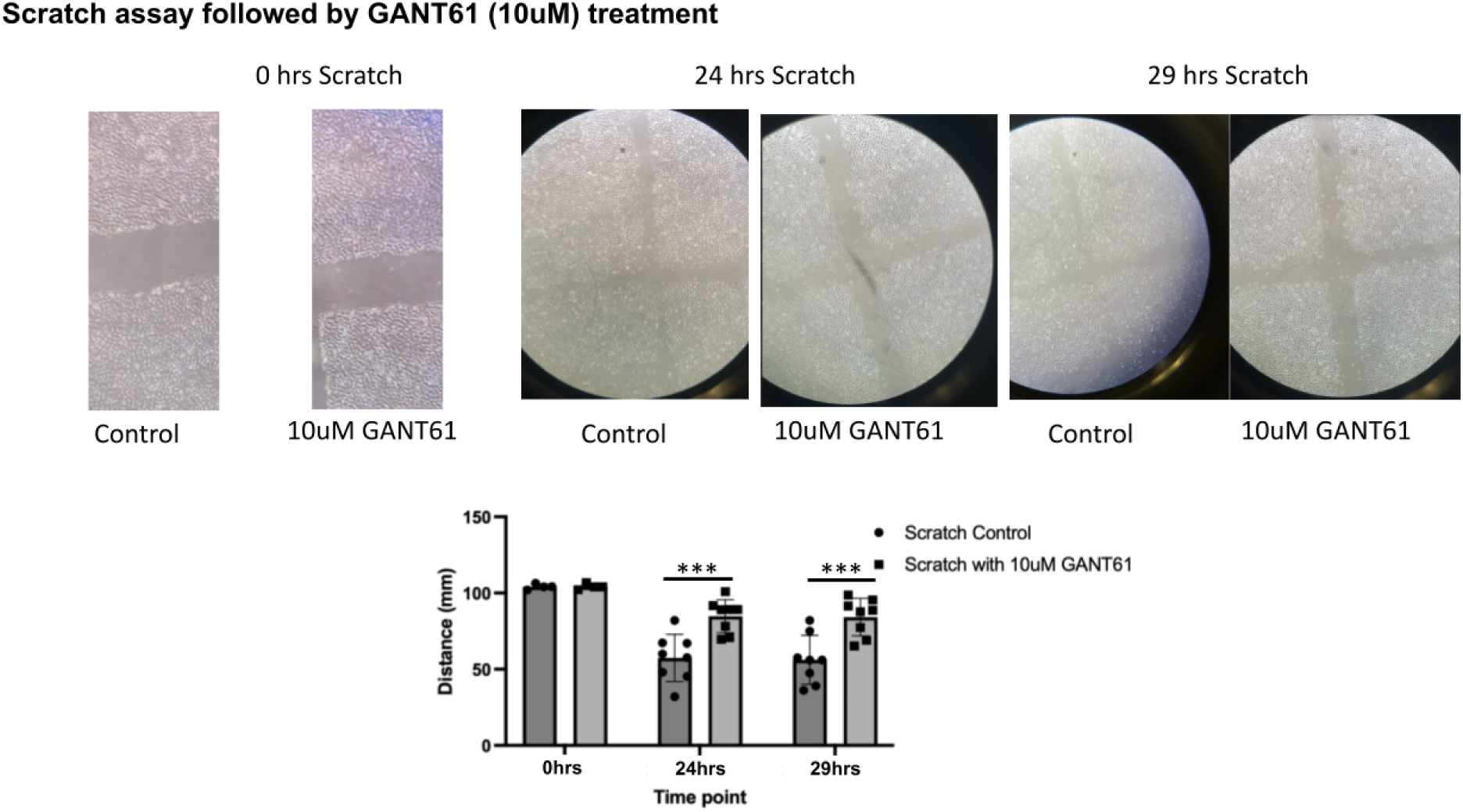
Wound healing assay with GLI inhibition GANT61.

**Supplementary Figure 7:**
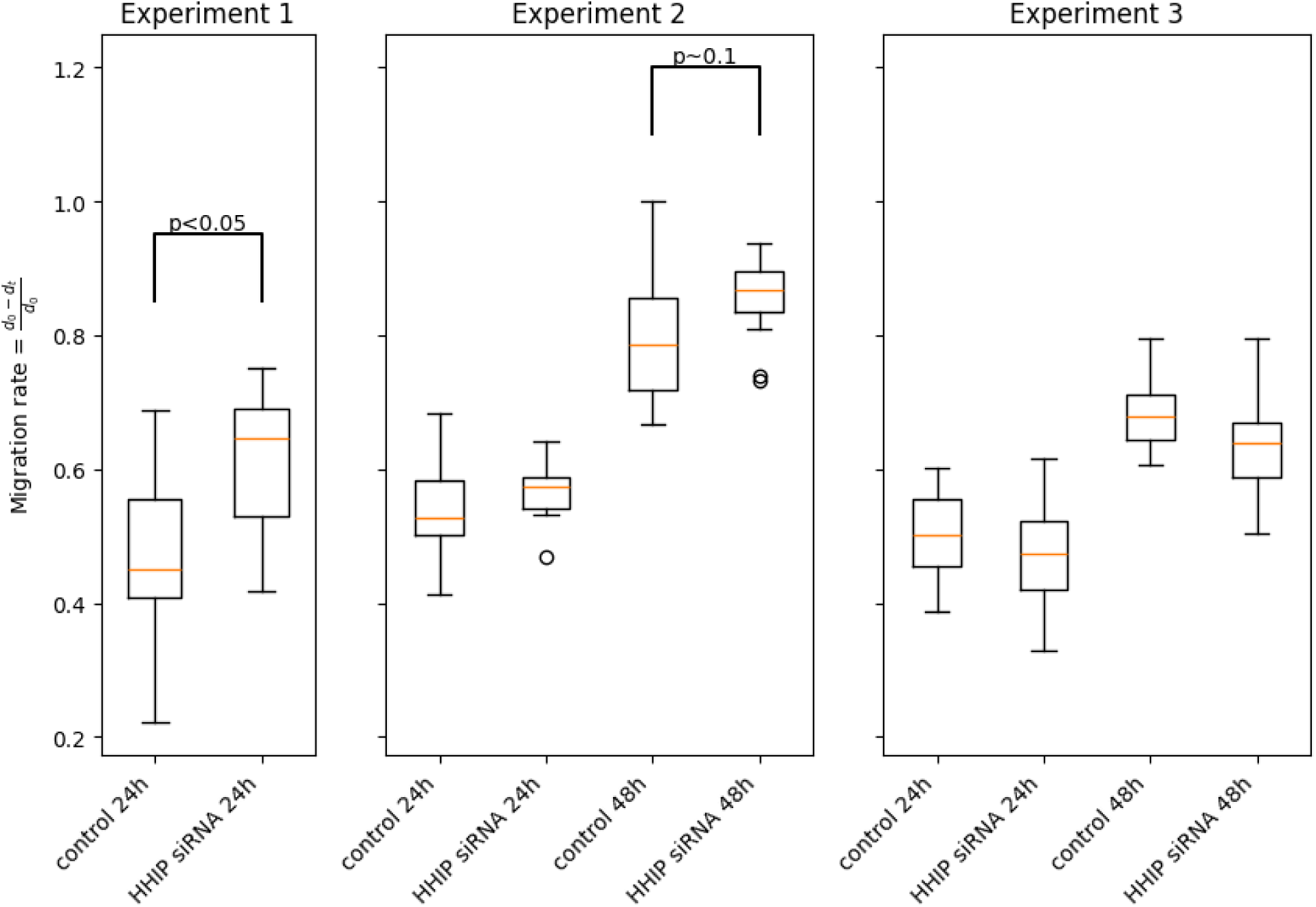
Migration rates in HHIP knockdown cells in 3 repeats.

**Supplementary Figure 8:**
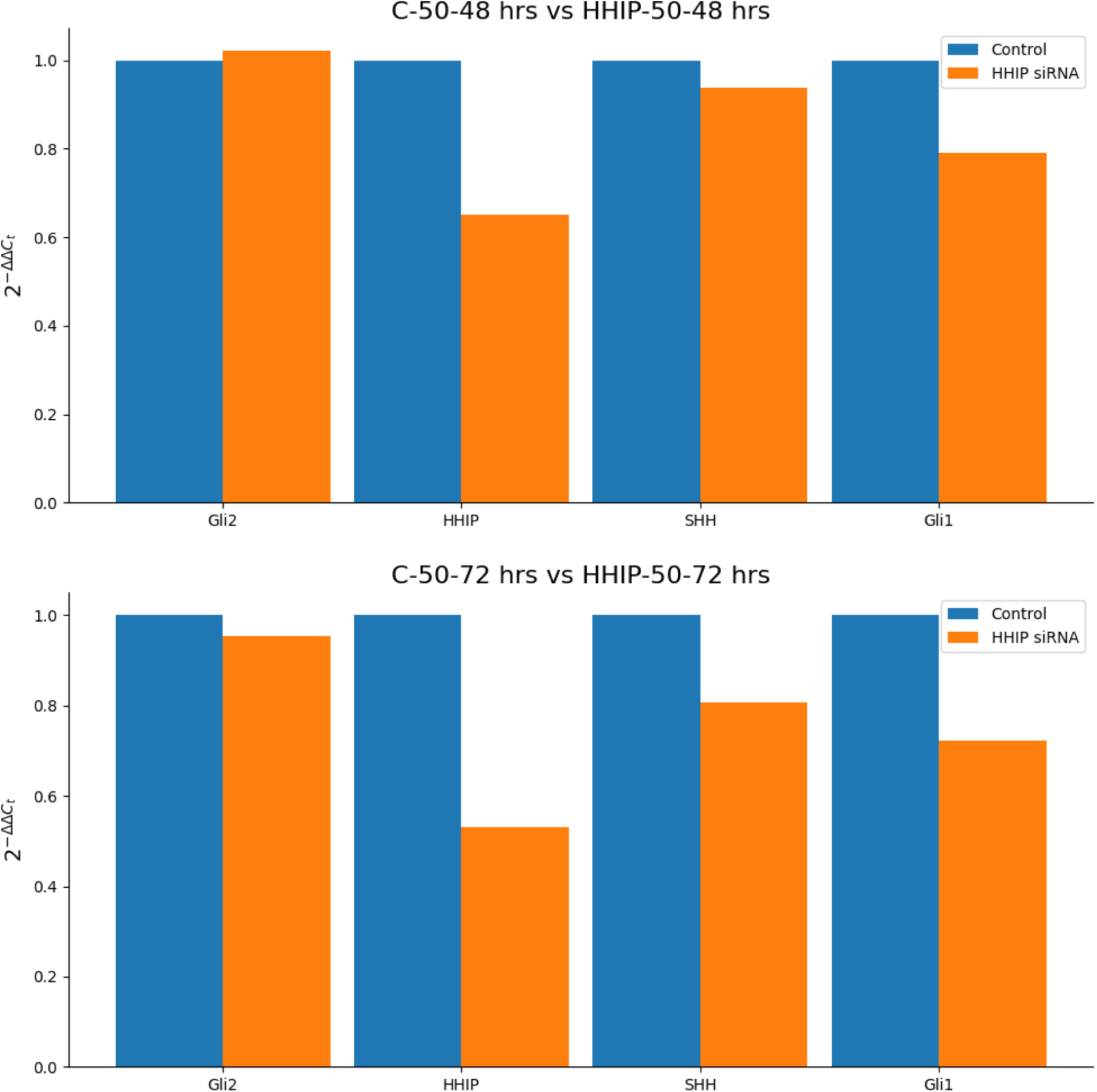
qPCR results of HHIP KO cells with *no scratch* at 48 and 72 hours. (single repeat measured in two time-points)

## Notes

https://github.com/deriteidavid/models_of_hhip_driven_emt_in_copd

### Summary of Updates

- Updated Figure 2 (added the former Suppl Figure 1 as a third panel) - Updated validation info on the second Boolean model - Added grant support info

