## Supplementary material for "HHIP’s Dynamic Role in Epithelial Wound Healing Reveals a Potential Mechanism of COPD Susceptibility": Channing Review HHIP EMT - 2024.10.02 - final.docx

### Supplementary Figures

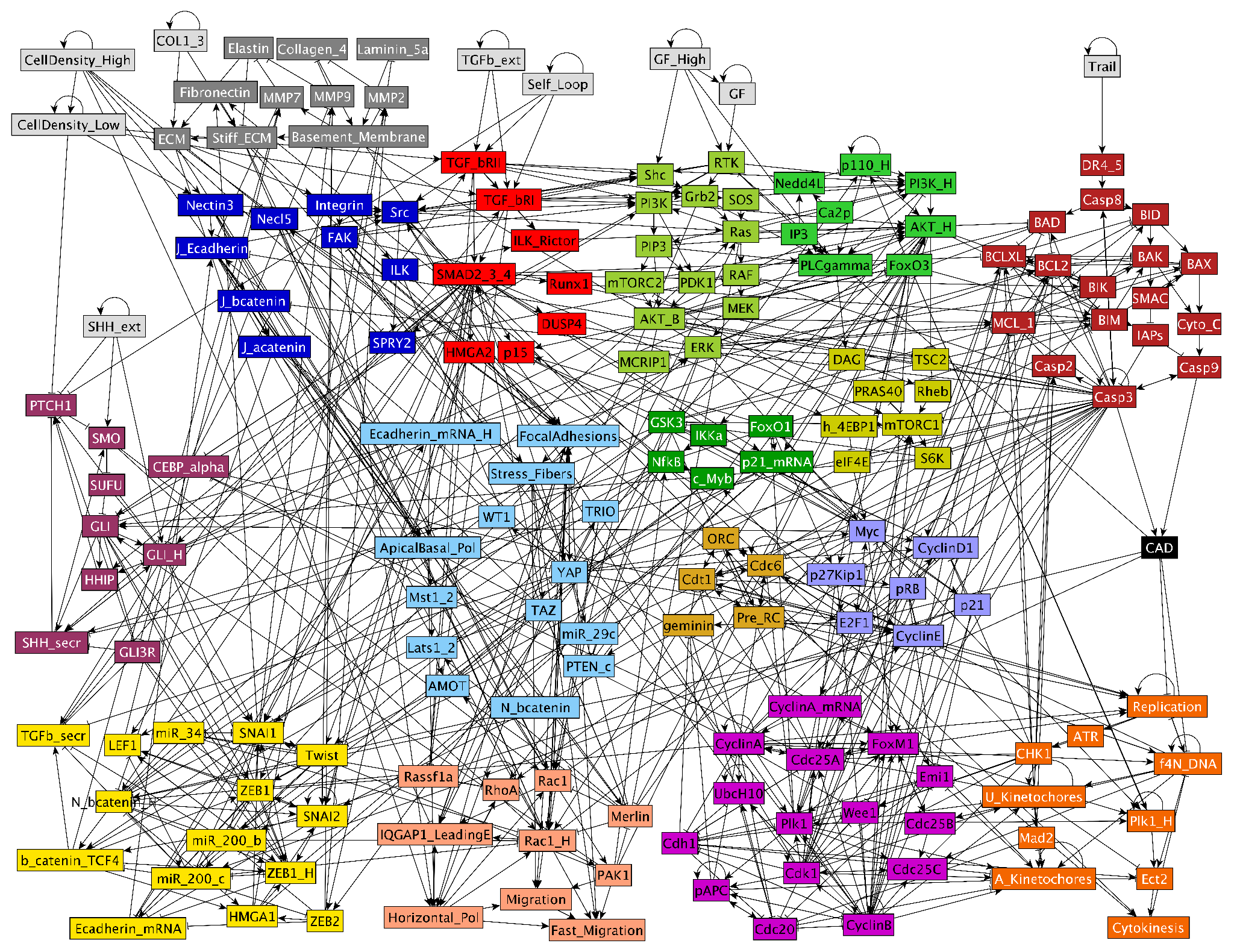

**Supplementary Figure 1: Full Boolean network model of SHH signaling linked to mechanosensitive EMT regulation and cell cycle progression.** Full network representation of our Boolean model expanded from (*30*). *light gray*: inputs representing factors in the microenvironment of the modeled cell; *dark gray*: Matrix (feedback control of ECM composition; new); *dark blue*: Adhesion signals; *red*: TGFb signaling; *soft maroon*: Shh signaling (new); *light blue*: Contact Inhibition; *yellow*: EMT switch; *pink/light orange*: Migration; *light/dark green*: Growth factor signaling & NF-κB; *mustard*: mTORC1 signaling; *brown*: Origin Licensing; *lilac*: Restriction Switch; *magenta*: cell cycle phase switch; *orange*: cell cycle control; *dark red*: Apoptosis; *black links between molecules:* → : activation; –| : inhibition (see Supplementary Table 4 for details on all nodes and links, and Supplementary File 2 for Boolean model in .dmms, .SBML and .BooleanNet formats.)

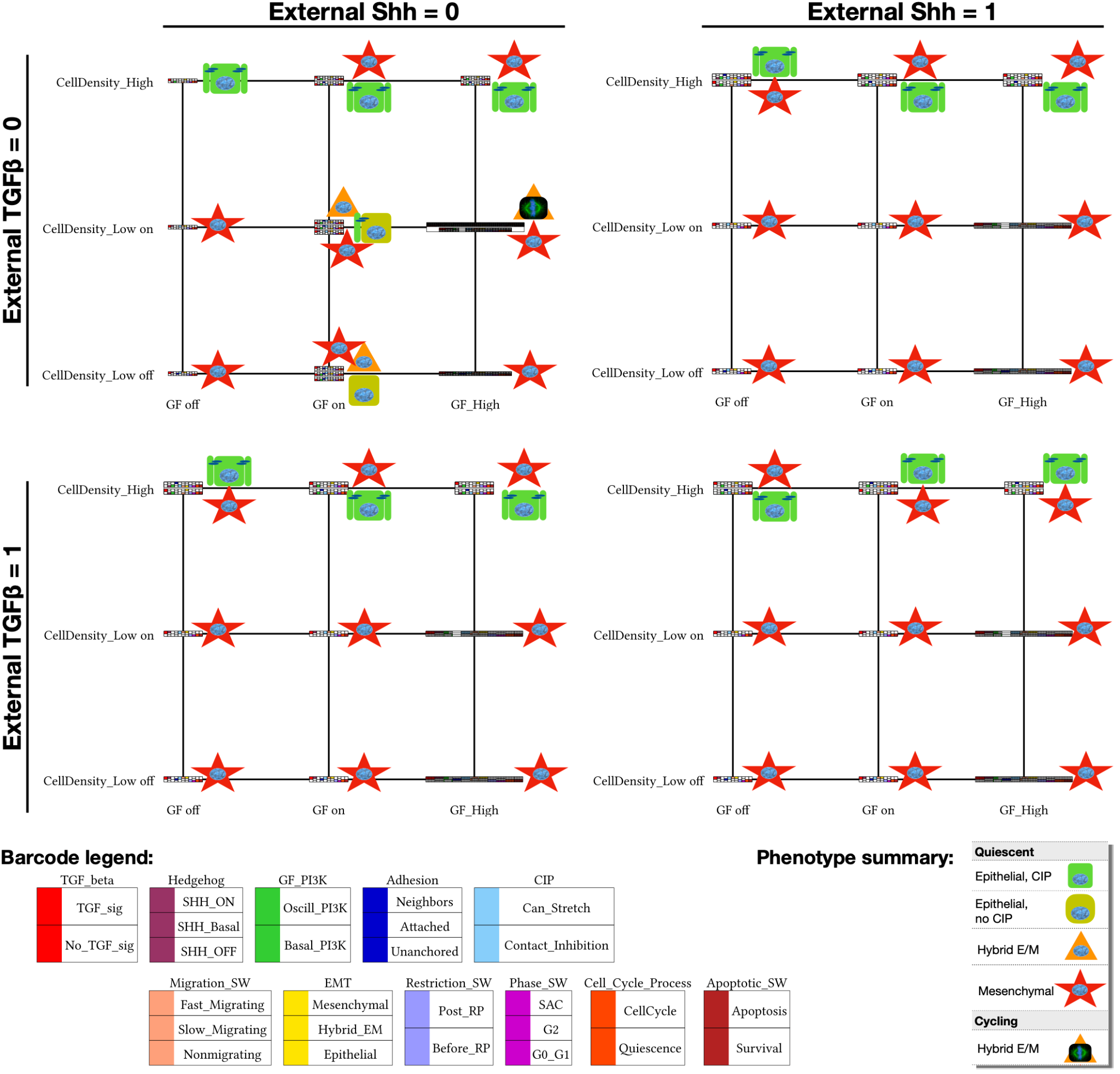

**Supplementary Figure 2: Relevant attractors of the large Boolean model, organized by external environment.** Relevant model cell states (synchronous attractors) detected in every combination of no/low/high growth-factor (*x* axis), no/low/high cell density (alone / monolayer’s edge / full confluence with no room to stretch; *y* axis), absence/presence of external Shh (*left/right*), and absence/presence of external TGFβ (*top/bottom*). Barcodes representing each attractor were derived by comparing the expression of nodes in each relevant module to a predetermined molecular signature known to represent a cell phenotype (e.g., apoptosis vs. survival) and encoded in the model’s .dmms file (barcode legend, *bottom left*). Oscillatory phenotypes have expanded barcodes that mark the transitions their regulatory switches undergo during the cycle. ***Note:*** attractors representing apoptotic cells, stable in every environment-combination, were filtered out to reduce clutter; as were quiescent polyploid cell states (reachable due to cell cycle errors only, but stable in quiescence). Visual summaries of overall cell states indicate epithelial (*green*) vs. hybrid E/M (*orange*) vs. mesenchymal (*red*) phenotypes, as well as quiescence (*round nucleus*) vs. cell cycle (*mitotic spindle image*). ON/OFF state of the network in every detected attractor included as Supplementary File 4.

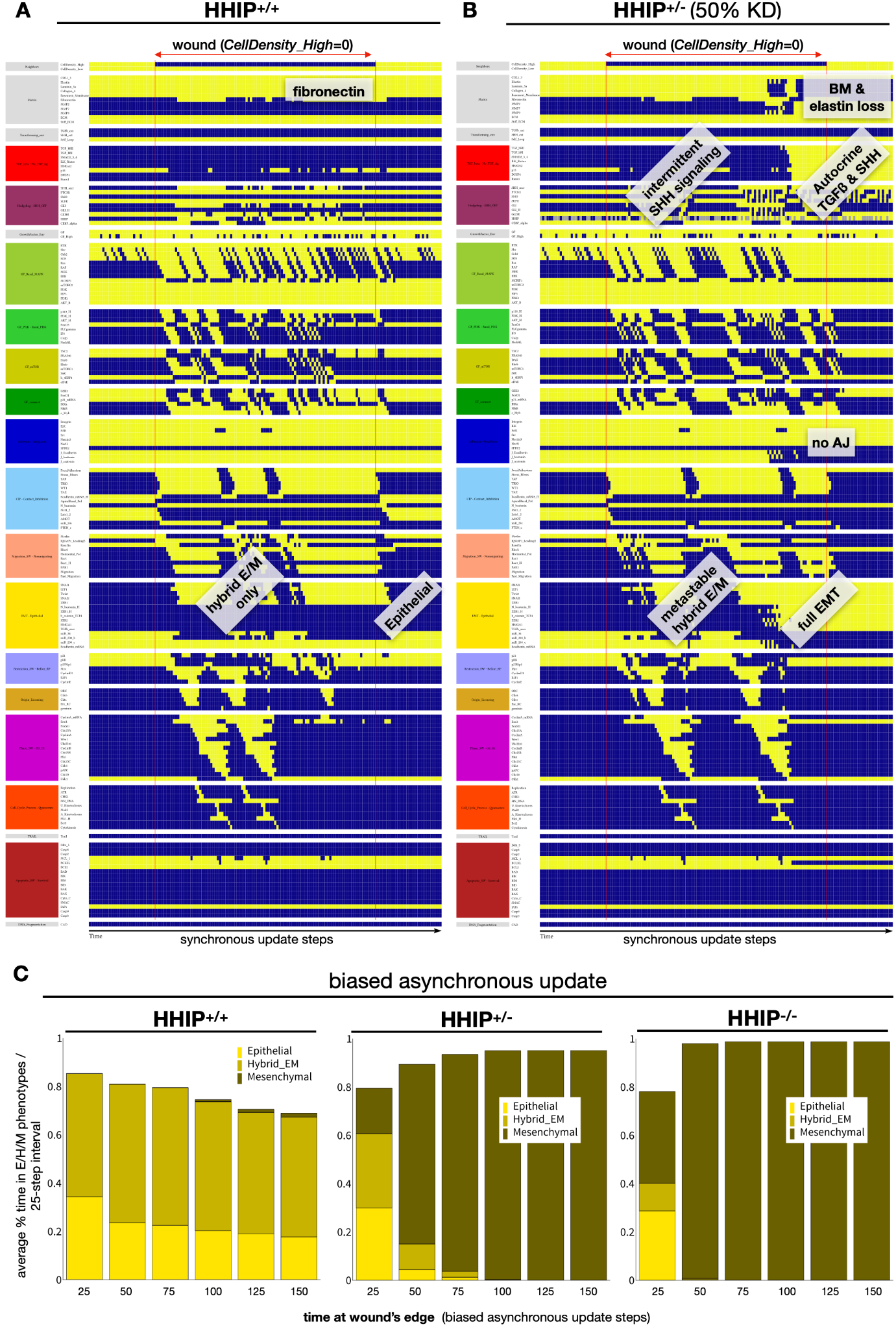

**Supplementary Figure 3: Full network dynamics for Figure 4A-B (Boolean model predicts that HHIP haploinsufficiency upregulates full EMT during lung re-epithelialization). A-B)** Dynamics of the expression/activity of all regulatory molecules in a cell at high confluency (first interval), exposed to the edge of a wound in (middle interval), then transitioning to high confluency (last interval) (A) wild-type (HHIP+/+) and (B) heterozygous HHIP+/- cells (HHIP is inactive in 50% of time-steps, modeling the stochastic weakening of its effects at reduced levels) exposed to 75% saturating mitogenic signals but no external Shh or TGFβ. *X axis*: time-steps (synchronous update); *y axis*: nodes organized in regulatory modules; *yellow/blue/gray*: ON/OFF/forced OFF; *vertical red line:* start / end of wound; *labels*: relevant phenotype changes. **C) *Figure 4C results with biased asynchronous update:*** Stacked bar charts showing the average % time cells exposed to a gap (at a monolayer’s edge) spend in Epithelial (*yellow*), Hybrid E/M (dark yellow) and Mesenchymal (mustard) states in consecutive 25-minute intervals (75% of saturating mitogen level, no external Shh or TGFβ). *Sample size:* 1000 independent runs; *Update:* biased asynchronous.

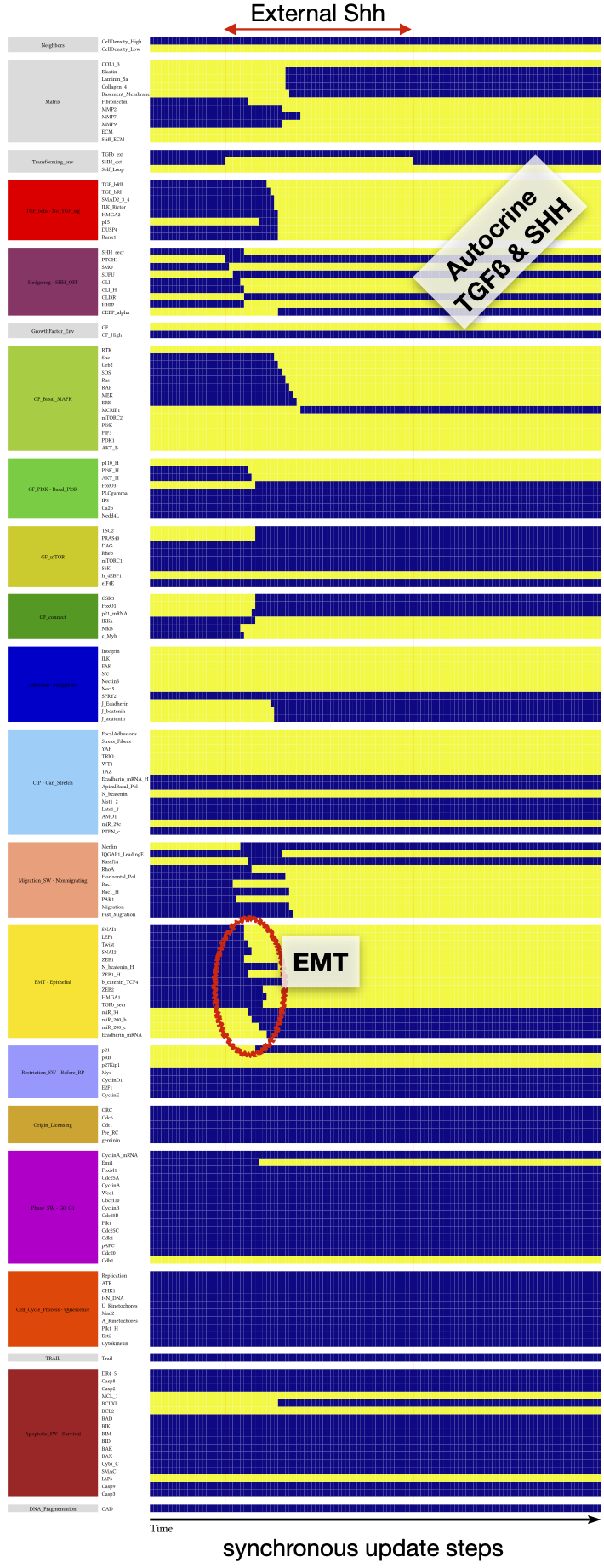

**Supplementary Figure 4: External Shh triggers EMT.** Dynamics of the expression/activity of all regulatory molecules in a cell at a monolayer’s edge, exposed external Shh for 100 time-steps (mitogenic signals are sufficient for survival but not cell cycl;e entry; no external TGFβ). *X axis*: time-steps (synchronous update); *y axis*: nodes organized in regulatory modules; *yellow/blue*: ON/OFF; *vertical red line:* start / end of Shh pulse; *red oval:* timing of EMT; *labels*: relevant phenotype changes.

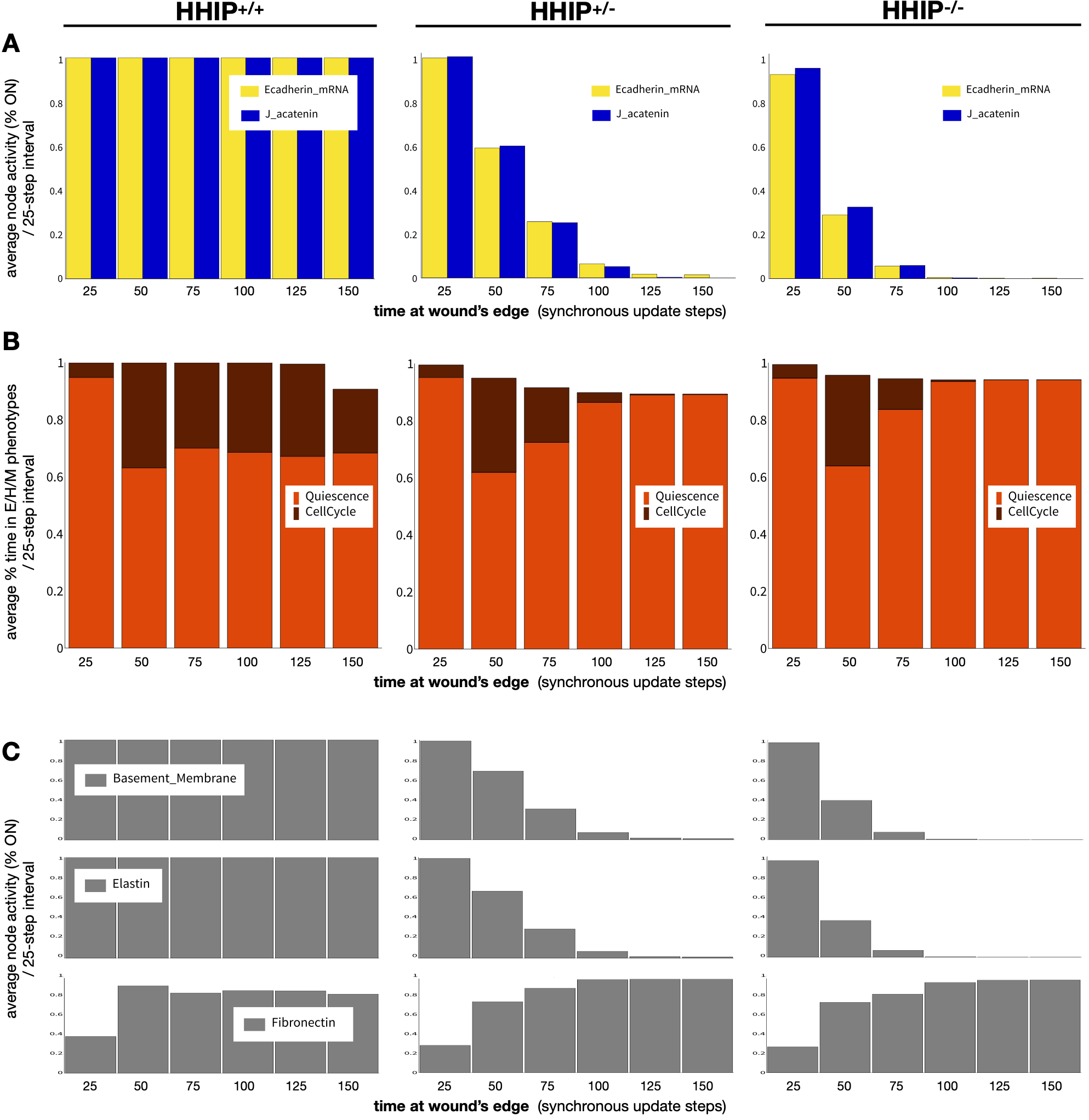

**Supplementary Figure 5: HHIP knockdown reduces cell-cell adhesion, proliferation, and stiffens the ECM while degrading the alveolar basement membrane. A-C)** Wild-type (HHIP^+/+^; *left*), heterozygous HHIP^+/-^ (*middle*) and HHIP-null (HHIP^-/-^; *right*) cells exposed to 75% saturating mitogenic signals but no external Shh or TGFβ. **A)** Average activity (% time ON) of the E-cadherin mRNA node (yellow) and junctional ɑ-catenin (*blue*); **B)** stacked bar charts showing the average % time cells spend in Quiescence (*red*) vs. in cell cycle (*dark red*); and **C)** average levels (% time ON) of the Basement Membrane (*top*), Elastin (middle) and Fibronectin (bottom) nodes in consecutive 25-minute intervals. *Sample size:* 1000 independent runs; *Update:* synchronous.

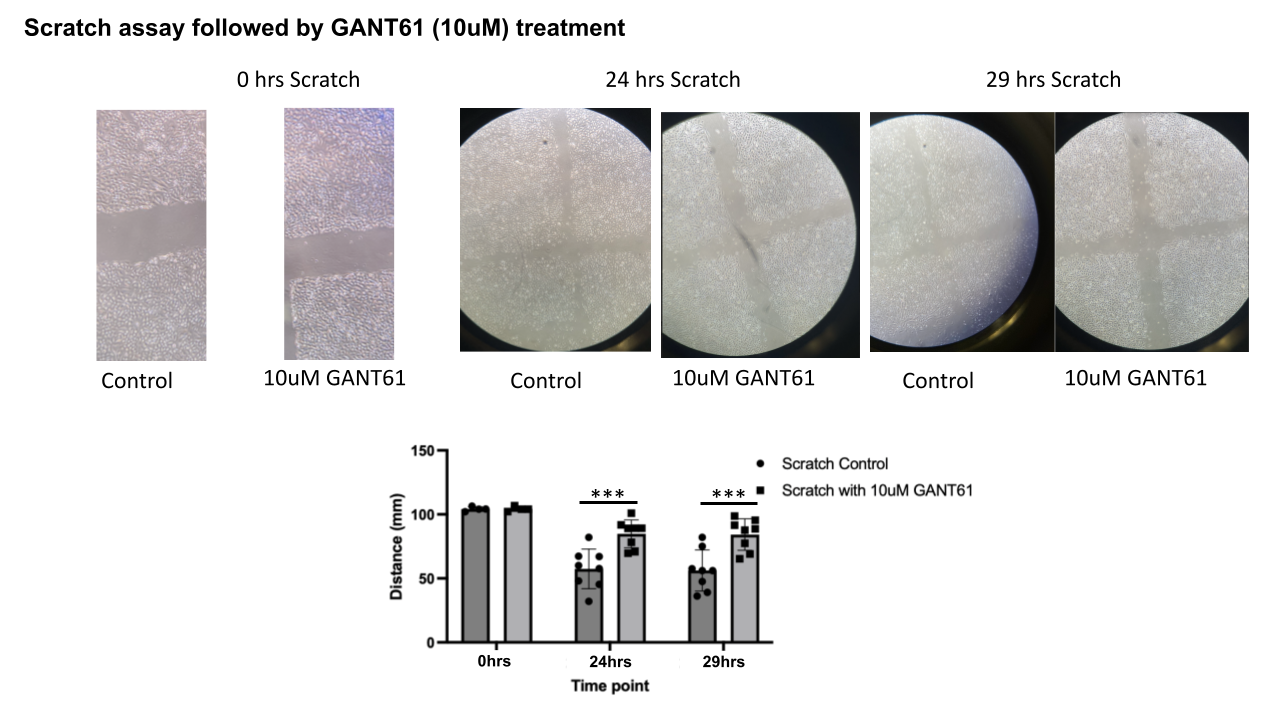

**Supplementary Figure 6: Wound healing assay with GLI inhibition GANT61**

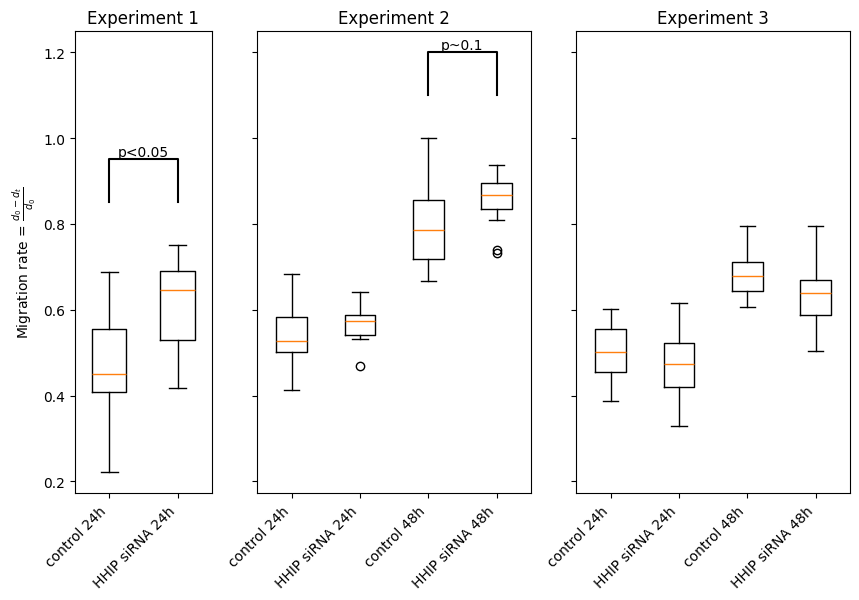

**Supplementary Figure 7: Migration rates in HHIP knockdown cells in 3 repeats**

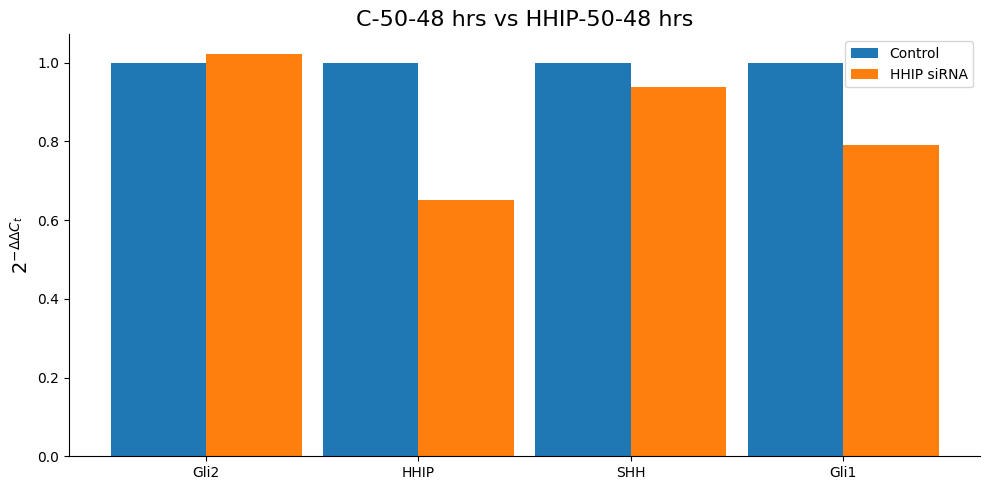

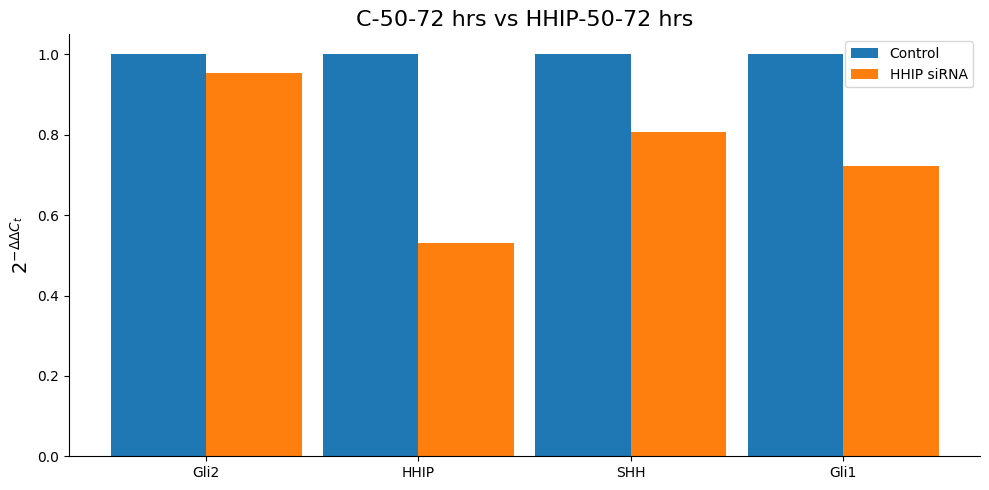

**Supplementary Figure 8: qPCR results of HHIP KO cells with *no scratch* at 48 and 72 hours** (single repeat measured in two time-points)
