## Supplementary material for "HHIP’s Dynamic Role in Epithelial Wound Healing Reveals a Potential Mechanism of COPD Susceptibility": Supplementary File 1 - attractor_analysis_new_model.html


### This Document contains the attractor analysis of the Hedghehog-EMT model¶

We use AEON.py for to find the attractors, to install the exact version we use run:  
`pip install biodivine_aeon==1.0.0a6`

In [1]:

```
import biodivine_aeon as ba
from pathlib import Path
import os.path
import numpy as np
import pandas as pd
```

```
Detected IPython (`ZMQInteractiveShell`). Log level set to `LOG_ESSENTIAL`.
```

In [2]:

```
model_name = 'HH_EMT'
```

convert boolean net format to bnet

In [3]:

```
import pystablemotifs as sm
with open(model_name + '.booleannet','r') as f:
    rules = f.read()
rules=sm.format.booleannet2bnet(rules)
with open(model_name + '.bnet','w') as f:
    f.write(rules)
```

In [4]:

```
bn = ba.BooleanNetwork.from_file(model_name + ".bnet")
```

In [5]:

```
bn = bn.inline_inputs(infer_inputs=True, repair_graph=True)
print(bn)

ctx = ba.SymbolicSpaceContext(bn)
graph = ba.AsynchronousGraph(bn, ctx)
```

```
BooleanNetwork(variables=22, regulations=50, explicit_parameters=2, implicit_parameters=0)
```

### Steady states¶

In [6]:

```
fixed_points = ba.FixedPoints.symbolic(graph, graph.mk_unit_colored_vertices())
```

```
Start symbolic fixed-point search with 16777216[nodes:2] candidates.
Found 2[nodes:27] fixed-points.
```

In [7]:

```
#Export into dictionary
all_steady_states = []
for param, vertex in fixed_points:
    steady_state_dict = {}
    for param_name in bn.explicit_parameter_names():
        steady_state_dict[param_name] = param[param_name].as_const()
        
    for variable_name in bn.variable_names():
        steady_state_dict[variable_name] = vertex[variable_name]

    all_steady_states.append(steady_state_dict)
```

In [8]:

```
all_steady_states
```

Out[8]:

```
[{'Damage': False,
  'neighbor': False,
  'AJ': False,
  'B_catenin_TCF4': False,
  'B_catenin_nuc': False,
  'E_cadherin': True,
  'GLI': False,
  'HHIP': False,
  'LEF1': False,
  'Migration': False,
  'NFKB': False,
  'PAK1': False,
  'Patched': True,
  'Rac1': False,
  'SHH': False,
  'SMO': False,
  'SNAI1': False,
  'SNAI2': False,
  'SUFU': True,
  'TGFb_secr': False,
  'TWIST1': False,
  'ZEB1': False,
  'miR_200': True,
  'miR_34': True},
 {'Damage': False,
  'neighbor': True,
  'AJ': True,
  'B_catenin_TCF4': False,
  'B_catenin_nuc': False,
  'E_cadherin': True,
  'GLI': False,
  'HHIP': False,
  'LEF1': False,
  'Migration': False,
  'NFKB': False,
  'PAK1': False,
  'Patched': True,
  'Rac1': False,
  'SHH': False,
  'SMO': False,
  'SNAI1': False,
  'SNAI2': False,
  'SUFU': True,
  'TGFb_secr': False,
  'TWIST1': False,
  'ZEB1': False,
  'miR_200': True,
  'miR_34': True}]
```

In [9]:

```
#plot_state_succession(newly_found_steady_states,state_labels=None,title=None, nodes=[i.strip() for i in df_attr['node']], fontsizex=10)
```

### Complex attractors¶

In [10]:

```
complex_attractor_mean_states = []

# Next, search for all minimal trap spaces.

if not os.path.isfile('minimal-traps.bdd'):
    # Compute the minimal trap spaces, but skip anything that contains a fixed-point, because
    # we already reported those. This will take a few hours, but should finish 
    # and will report progress.
    #
    # This could be faster if we first pre-computed some kindidates on a reduced network,
    # but this should be "good enough" for this case.
    #
    candidate_spaces = ctx.mk_unit_colored_spaces(graph)
    contains_fixed_point = ctx.mk_super_spaces(fixed_points.to_singleton_spaces(ctx))
    candidate_spaces = candidate_spaces.minus(contains_fixed_point)
    minimal_traps = ba.TrapSpaces.minimal_symbolic(ctx, graph, candidate_spaces)
    print("Computed minimal traps: ", minimal_traps)

    # Save for later.
    Path('minimal-traps.bdd').write_text(minimal_traps.to_bdd().data_string())
else:
    # Result exists, reload it from file.
    bdd = ba.Bdd(ctx.bdd_variable_set(), Path('minimal-traps.bdd').read_text())
    minimal_traps = ba.ColoredSpaceSet(ctx, bdd)
    print("Loaded minimal traps: ", minimal_traps)
```

```
Loaded minimal traps:  ColoredSpaceSet(cardinality=2, symbolic_size=91)
```

In [11]:

```
print(" >>>>> MINIMAL TRAP SPACES <<<<<")
for param, space in minimal_traps:
    print(f"Minimap trap space: inputs({param}) variables({space})")

complex_attractor_mean_states = []

# Now we can search for attractors *inside* the minimal trap spaces.
# There are slightly more efficient ways to do this (ideally, we just want to process
# each minimal trap independently), but here it seems that this is fast enough.

trap_states = minimal_traps.to_colored_vertices(ctx)
# We can just use this simpler algorithm, because the minimal trap spaces are rather small already.
attractor_sets = ba.Attractors.xie_beerel(graph, trap_states)
instantiated_attractors = []
for attractor in attractor_sets:
    for input_valuation in attractor.colors():
        # This is still a relation of vertices and colors, but it only contains a single color (input valuation), 
        # hence a single attractor.
        attractor_states = attractor.intersect_colors(input_valuation.to_symbolic())
        instantiated_attractors.append(attractor_states)
        print(f"Found complex attractor for {input_valuation} with {attractor_states.cardinality()} vertices.")	

        complex_attractor_mean_state = {}

        # This prints per-variable statistics about variable values.		
        for var in graph.network_variables():
            # Splits the attractor states based on the value of `var` and prints statistics about the split.
            attr_true = graph.mk_subspace({ var: 1 }).intersect(attractor_states)
            attr_false = graph.mk_subspace({ var: 0 }).intersect(attractor_states)
            #print(f"Variable {graph.get_network_variable_name(var)}: {attr_true.cardinality()} one / {attr_false.cardinality()} zero | {float(attr_true.cardinality())/attractor_states.cardinality()}")
            
            for param_name in bn.explicit_parameter_names():            	
            	complex_attractor_mean_state[param_name]=float(input_valuation[param_name].as_const())
            complex_attractor_mean_state[graph.get_network_variable_name(var)] = float(attr_true.cardinality())/attractor_states.cardinality()        
        complex_attractor_mean_states.append(complex_attractor_mean_state)
```

```
 >>>>> MINIMAL TRAP SPACES <<<<<
Minimap trap space: inputs(ColorModel({'Damage': 'true', 'neighbor': 'false'})) variables(SpaceModel({'AJ': 0, 'B_catenin_TCF4': *, 'B_catenin_nuc': *, 'E_cadherin': *, 'GLI': *, 'HHIP': *, 'LEF1': 1, 'Migration': *, 'NFKB': 1, 'PAK1': *, 'Patched': *, 'Rac1': *, 'SHH': *, 'SMO': *, 'SNAI1': *, 'SNAI2': *, 'SUFU': *, 'TGFb_secr': *, 'TWIST1': *, 'ZEB1': *, 'miR_200': *, 'miR_34': *}))
Minimap trap space: inputs(ColorModel({'Damage': 'true', 'neighbor': 'true'})) variables(SpaceModel({'AJ': 1, 'B_catenin_TCF4': 0, 'B_catenin_nuc': 0, 'E_cadherin': 1, 'GLI': *, 'HHIP': *, 'LEF1': 1, 'Migration': 0, 'NFKB': 1, 'PAK1': *, 'Patched': *, 'Rac1': *, 'SHH': *, 'SMO': *, 'SNAI1': *, 'SNAI2': *, 'SUFU': *, 'TGFb_secr': 0, 'TWIST1': *, 'ZEB1': 0, 'miR_200': 1, 'miR_34': *}))
Start Xie-Beerel attractor detection on 528384[nodes:18] candidates.
Found complex attractor for ColorModel({'Damage': 'true', 'neighbor': 'true'}) with 768 vertices.
Found complex attractor for ColorModel({'Damage': 'true', 'neighbor': 'false'}) with 73728 vertices.
 > Found a bottom SCC: 768x1[nodes:43].
 > Found a bottom SCC: 73728x1[nodes:35].
Attractor detection finished with 2 results.
```

In [12]:

```
len(complex_attractor_mean_states)
```

Out[12]:

```
2
```

In [13]:

```
print(input_valuation)
```

```
ColorModel({'Damage': 'true', 'neighbor': 'false'})
```

In [14]:

```
for param_name in bn.explicit_parameter_names():
    print(param_name, input_valuation[param_name].as_const())
```

```
Damage True
neighbor False
```

In [15]:

```
df = pd.DataFrame(all_steady_states+complex_attractor_mean_states).astype(float)
node_order = list(all_steady_states[0].keys())
df = df[node_order]
df=df.transpose()
```

In [16]:

```
import matplotlib.pyplot as plt
plt.figure(figsize = (15,30))
plt.imshow(df,interpolation='none')
plt.yticks(range(len(df)),df.index)
plt.colorbar()
```

Out[16]:

```
<matplotlib.colorbar.Colorbar at 0x7b7f119c2460>
```

In [ ]:
