## Supplementary material for "HHIP’s Dynamic Role in Epithelial Wound Healing Reveals a Potential Mechanism of COPD Susceptibility": Large Model Validation Script.py

```
import csv
import pandas as pd
import matplotlib.pyplot as plt
from mpl_toolkits import mplot3d
import seaborn as sns
from statannotations.Annotator import Annotator
from matplotlib.backends.backend_pdf import PdfPages
from mpl_toolkits.axes_grid1 import make_axes_locatable
import textwrap


model_folder = "/Users/eregan/Dropbox/__RESEARCH/MODEL_Folders/BooleanModels_dmms/__VALIDATION/4_5 - EMT and all using module/7a - COPD - DEV/"
validation_csv_folder = "/Users/eregan/Dropbox/__RESEARCH/MODEL_Folders/BooleanModels_dmms/__VALIDATION/4_5 - EMT and all using module/7a - COPD - DEV/"
#with open(validation_csv_folder + 'ST3 - Large model valid read by ER Python script.csv', newline='\n') as csvfile:

#pp = PdfPages('SFxx - Large model validaton figures.pdf')

with open(validation_csv_folder + 'ST5 - Large model validation.csv', newline='\n') as csvfile:
	ST3 = csv.DictReader(csvfile, delimiter=',')
	csv_cols = []
	Stat_tests_output = []
	Stat_tests_output.append(['perturbed node in model', 'perturbation type','downstream node in model','downstream phenotype in model','expected change direction','direction match','p-value'])

	BAD_tests_output = []
	BAD_tests_output.append(['perturbed node in model', 'perturbation type','downstream node in model','downstream phenotype in model','expected change direction','direction match','p-value'])
	
	line_OK = 0
	some_in_line_OK = 0
	tests_OK = 0
	N_lines = 0
	N_tests =0
	
	for exp_ID, col in enumerate(ST3):
		paper = col['paper']
		node_st = col['perturbed node in model'].replace(" ", "")
		nodes_perturbed = [str(n) for n in node_st.split(',')]
		#print(nodes_perturbed)
		pert_type_str = col['perturbation type'].replace(" ", "")
		pert_type = [int(n) for n in pert_type_str.split(',')]
		change_dir = col['change direction']
		#print(pert_type)
		exp_name = ""
		if (len(nodes_perturbed) == len(pert_type)):
			for i in range(0, len(nodes_perturbed)):
				if(pert_type[i] == 0):
					exp_name = exp_name + nodes_perturbed[i] + "_KD "
				else: exp_name = exp_name + nodes_perturbed[i] + "_OE "
		else: print("Line: " + col +"\n\t number of nodes perturbed does not match length of \'perturbation type\' list!")
		#print("Experiment name: " + exp_name)
		
		output_nodes_str = col['downstream node in model'].replace(" ", "")
		if(len(output_nodes_str)>0):
			output_nodes = [s for s in output_nodes_str.split(',')]
		else:
			output_nodes = []
		#print("output nodes string:" + output_nodes_str)
		#print(output_nodes)
					
		output_phen_str = col['downstream phenotype in model'].replace(" ", "")
		if(len(output_phen_str)>0):
			output_phen = [s for s in output_phen_str.split(',')]
		else:
			output_phen = []
		#print("output phenotypes string:" + output_phen_str)
		#print(output_phen)
			
		expected = col['change direction']
		ctr = col['In silico control name']
		exp = col['In silico experiment name']
		times_str = col['Time windows to compare'].replace(" ", "")
		times = [int(n) for n in times_str.split(',')]
		#print(times)
		attr_code = col['Attractor_code']
		csv_cols.append(col)
			
		if(output_nodes):
			N_lines = N_lines + 1
			print(output_nodes)
			print("N_lines = " + str(N_lines))
			fig, axes = plt.subplots(1, 1, figsize=(2.5*len(output_nodes)+len(exp_name)*0.2, 5), sharex=False, sharey=False)		
			df_nodes_list = []
			ctr_node_file = model_folder + "_EXP/General_Time_Series/"+ ctr + "/NodeBCh/" + attr_code + "_" + ctr + ".csv"
			#print(ctr_node_file)
			ok=0
			with open(ctr_node_file, newline='\n') as ct_file:
				CT = csv.reader(ct_file, delimiter=',', quotechar='"')
				#CT_cols = []
				for i in CT:
					#print(i[0])
					#print("ok = " + str(ok))
					if(ok==1 and (str(i[0]) in output_nodes)):
						#print("adding line:" + i[0])
						#CT_cols.append(i[0:10])
						for e in i[1:]:
							df_nodes_list.append(['Control',i[0],float(e)])
							#print("\t\tadding:" + i[0] + e)
					window=i[0][:-1]
					windows=window.split()
					#print(len(windows), windows)
					if(len(windows)>1):
						if(int(windows[1]) == times[0]-1): 
							ok=1
						else: ok=0
			#print(CT_cols)
			exp_node_file = model_folder + "_EXP/General_Time_Series/"+ exp + "/NodeBCh/" + attr_code + "_" + exp + ".csv"
			#print(ctr_node_file)
			ok=0
			with open(exp_node_file, newline='\n') as ct_file:
				EXP = csv.reader(ct_file, delimiter=',', quotechar='"')
				#EXP_cols = []
				for i in EXP:
					#print(i[0])
					#print(output_nodes)
					#print("ok = " + str(ok))
					if(ok==1 and (str(i[0]) in output_nodes)):
						#EXP_cols.append(i[0:10])
						#print("adding line:" + i[0])
						for e in i[1:]:
							df_nodes_list.append([exp_name,i[0],float(e)])
					window=i[0][:-1]
					windows=window.split()
					#print(len(windows), windows)
					if(len(windows)>1):
						if(int(windows[1]) == times[1]-1): 
							ok=1
						else: ok=0
			#print("df_nodes_list: ")
			#print(df_nodes_list)
			
			x = "Node output"
			y = "Av. molecule expression/activity"
			hue = "Experiment"
			df_nodes = pd.DataFrame(df_nodes_list, columns=[hue, x, y])
			hue_order=['Control', exp_name]
			pairs = []
			for gname in output_nodes:
				pairs.append(((gname, 'Control'), (gname, exp_name)))
				N_tests = N_tests + 1
			axes = sns.barplot(data=df_nodes, x=x, y=y, hue=hue, hue_order=hue_order, seed=2021, ci="sd", edgecolor="black", errcolor="black", errwidth=1.5, capsize = 0.1, alpha=0.5)
			#sns.stripplot(x=x, y=y, hue=hue, data=df_nodes, dodge=True, alpha=0.6, ax=ax)
				
			annot = Annotator(None, pairs)
			annot.new_plot(axes, pairs, plot='barplot', data=df_nodes, x=x, y=y, hue=hue, hue_order=hue_order, seed=2021)
			annot.configure(test='t-test_ind', text_format='star', loc='inside', verbose=2)
			_, test_results = annot.apply_test().annotate()
			
			axes.set_title("\n".join(textwrap.wrap(paper, 30*len(output_nodes))), fontsize = 8)
			plt.text(x=len(output_nodes), y=-0.1, s="\n".join(textwrap.wrap("Control sim.: " + ctr, 100)), fontsize=6, color='gray',rotation=90)
			plt.text(x=len(output_nodes)+0.1, y=-0.1, s="\n".join(textwrap.wrap("Perturb. sim.: " + exp, 100)), fontsize=6, color='gray',rotation=90)
			plt.legend(loc='upper left', bbox_to_anchor=(1.03, 1), title=hue)
			plt.tight_layout()
			
			ii = []
			for result in test_results:
				ii.append(output_nodes.index(result.data.group1[0]))
			
			stat_results = []
			for indnow in ii:
				stat_results.append(test_results[indnow].data.pvalue)


			dir_ok =[]
#			print(axes.containers)
			this_line_OK = 1
			some_OK =0

			for indnow, gname in enumerate(output_nodes):
 				ct_av = axes.containers[0][indnow].get_height()
 				exp_av = axes.containers[1][indnow].get_height()
 				#print(str(ct_av) + " vs. " + str(exp_av))
 				if((exp_av-ct_av > 0) and change_dir == '1'):
 					if(stat_results[indnow]<= 5.00e-02):
 						dir_ok.append("Significant")
 						some_OK = 1
 						tests_OK = tests_OK + 1
 					else:
 						dir_ok.append("Dir. OK but Not Sig.")
 						this_line_OK = 0
 						
 				if((exp_av-ct_av > 0) and change_dir == '0'):
 					if(stat_results[indnow]<= 5.00e-02):
 						dir_ok.append("Mismatch (sign.)")
 						this_line_OK = 0
 					else:
 						dir_ok.append("Mismatch (model ns.)")
 						this_line_OK = 0
 						
 				if((exp_av-ct_av > 0) and change_dir == 'no change'):
 					if(stat_results[indnow]<= 5.00e-02):
 						dir_ok.append("Mismatch, increase")
 						this_line_OK = 0
 					else:
 						dir_ok.append("Match (ns increase)")
 						some_OK = 1
 						tests_OK = tests_OK + 1

 				if((exp_av-ct_av < 0) and change_dir == '0'):
 					if(stat_results[indnow]<= 5.00e-02):
 						dir_ok.append("Significant")
 						some_OK = 1
 						tests_OK = tests_OK + 1
 					else:
 						dir_ok.append("Dir. OK but Not Sig.")
 						this_line_OK = 0
 						
 				if((exp_av-ct_av < 0) and change_dir == '1'):
 					if(stat_results[indnow]<= 5.00e-02):
 						dir_ok.append("Mismatch (sign.)")
 						this_line_OK = 0
 					else:
 						dir_ok.append("Mismatch (model ns.)")
 						this_line_OK = 0

 				if((exp_av-ct_av < 0) and change_dir == 'no change'):
 					if(stat_results[indnow]<= 5.00e-02):
 						dir_ok.append("Mismatch, decrease")
 						this_line_OK = 0
 					else:
 						dir_ok.append("Match (ns decrease)")
 						some_OK = 1
 						tests_OK = tests_OK + 1
 						
 				if((exp_av-ct_av == 0) and change_dir == 'no change'):
  					dir_ok.append("Match (no change)")
  					some_OK = 1
  					tests_OK = tests_OK + 1
 				if((exp_av-ct_av == 0) and change_dir == '1'):
 					dir_ok.append("Mismatch (inc. not reproduced)")
 					this_line_OK = 0
 				if((exp_av-ct_av == 0) and change_dir == '0'):
 					dir_ok.append("Mismatch (dec. not reproduced)")
 					this_line_OK = 0

			line_OK = line_OK + this_line_OK
			some_in_line_OK = some_in_line_OK + some_OK
			
			#print(change_dir)
			#print(stat_results)
			#print(dir_ok)
			#print(output_nodes)
			for ind in range(len(output_nodes)):
				Stat_tests_output.append([node_st, pert_type_str, output_nodes[ind],'',change_dir,dir_ok[ind],stat_results[ind]])
				if(dir_ok[ind] == "Dir. OK but Not Sig." or dir_ok[ind] == "Mismatch, increase" or dir_ok[ind] == "Mismatch, decrease" or dir_ok[ind] == "Mismatch (inc. not reproduced)" or dir_ok[ind] == "Mismatch (inc. not reproduced)" or dir_ok[ind] == "Mismatch (model ns.)"):
					BAD_tests_output.append([node_st, pert_type_str, output_nodes[ind],'',change_dir,dir_ok[ind],stat_results[ind]])
				
# 			plt.show()
# 			exit()	
			
		if(output_phen):
			N_lines = N_lines + 1
			print(output_phen)
			print("N_lines = " + str(N_lines))
			
			fig, axes = plt.subplots(1, 1, figsize=(2.5*len(output_phen)+len(exp_name)*0.4, 5), sharex=False, sharey=False)		
#			fig, axes = plt.subplots(1, 1, figsize=(2.5*len(output_nodes)+len(exp_name)*0.2, 5), sharex=False, sharey=False)		
			df_phen_list = []
			ctr_phen_file = model_folder + "_EXP/General_Time_Series/"+ ctr + "/PhBCh/" + attr_code + "_" + ctr + ".csv"
			#print(ctr_node_file)
			ok=0
			with open(ctr_phen_file, newline='\n') as ct_file:
				CT = csv.reader(ct_file, delimiter=',', quotechar='"')
				#CT_cols = []
				for i in CT:
					#print(i[0])
					#print("ok = " + str(ok))
					if(ok==1 and (str(i[0]) in output_phen)):
						#print("adding line:" + i[0])
						#CT_cols.append(i[0:10])
						for e in i[1:]:
							df_phen_list.append(['Control',i[0],float(e)])
							#print("\t\tadding:" + i[0] + e)
					window=i[0][:-1]
					windows=window.split()
					#print(len(windows), windows)
					if(len(windows)>1):
						if(int(windows[1]) == times[0]-1): 
							ok=1
						else: ok=0
			#print(CT_cols)
			ctr_phen_file = model_folder + "_EXP/General_Time_Series/"+ exp + "/PhBCh/" + attr_code + "_" + exp + ".csv"
			#print(ctr_node_file)
			ok=0
			with open(ctr_phen_file, newline='\n') as ct_file:
				EXP = csv.reader(ct_file, delimiter=',', quotechar='"')
				#EXP_cols = []
				for i in EXP:
					#print(i[0])
					#print(output_phen)
					#print("ok = " + str(ok))
					if(ok==1 and (str(i[0]) in output_phen)):
						#EXP_cols.append(i[0:10])
						#print("adding line:" + i[0])
						for e in i[1:]:
							df_phen_list.append([exp_name,i[0],float(e)])
							#print("\t\tadding:" + i[0] + e)
					window=i[0][:-1]
					windows=window.split()
					#print(len(windows), windows)
					if(len(windows)>1):
						if(int(windows[1]) == times[1]-1): 
							ok=1
						else: ok=0
			#print("df_nodes_list: ")
			#print(df_nodes_list)
			
			x = "Phenotype"
			y = "Average time in phenotype"
			hue = "Experiment"
			df_phen = pd.DataFrame(df_phen_list, columns=[hue, x, y])
			hue_order=['Control', exp_name]
			pairs = []
			for gname in output_phen:
				pairs.append(((gname, 'Control'), (gname, exp_name)))
				N_tests = N_tests + 1

			axes = sns.barplot(data=df_phen, x=x, y=y, hue=hue, hue_order=hue_order, seed=2021, ci="sd", edgecolor="black", errcolor="black", errwidth=1.5, capsize = 0.1, alpha=0.5, dodge=True)
			#sns.stripplot(x=x, y=y, hue=hue, data=df_nodes, dodge=True, alpha=0.6, ax=ax)
			annot = Annotator(None, pairs)
			annot.new_plot(axes, pairs, plot='barplot', data=df_phen, x=x, y=y, hue=hue, hue_order=hue_order, seed=2021)
			annot.configure(test='t-test_ind', text_format='star', loc='inside', verbose=2)
			_, test_results = annot.apply_test().annotate()

			#annot.apply_test().annotate()
			axes.set_title("\n".join(textwrap.wrap(paper, 30*len(output_phen))), fontsize = 8)
			plt.text(x=len(output_phen),     y=-0.1, s="\n".join(textwrap.wrap("Control sim.: "  + ctr, 100)), fontsize=6, color='gray',rotation=90)
			plt.text(x=len(output_phen)+0.1, y=-0.1, s="\n".join(textwrap.wrap("Perturb. sim.: " + exp, 100)), fontsize=6, color='gray',rotation=90)
			plt.legend(loc='upper left', bbox_to_anchor=(1.03, 1), title=hue)
			plt.tight_layout()
			#plt.show()
			
			ii = []
			for result in test_results:
				ii.append(output_phen.index(result.data.group1[0]))
			
			stat_results = []
			for indnow in ii:
				stat_results.append(test_results[indnow].data.pvalue)


			dir_ok =[]
#			print(axes.containers)
			this_line_OK = 1
			some_OK =0

			for indnow, gname in enumerate(output_phen):
 				ct_av  = axes.containers[0][indnow].get_height()
 				exp_av = axes.containers[1][indnow].get_height()
 				#print(str(ct_av) + " vs. " + str(exp_av))
 				if((exp_av-ct_av > 0) and change_dir == '1'):
 					if(stat_results[indnow]<= 5.00e-02):
 						dir_ok.append("Significant")
 						some_OK = 1
 						tests_OK = tests_OK + 1
 					else:
 						dir_ok.append("Dir. OK but Not Sig.")
 						this_line_OK = 0
 						
 				if((exp_av-ct_av > 0) and change_dir == '0'):
 					if(stat_results[indnow]<= 5.00e-02):
 						dir_ok.append("Mismatch (sign.)")
 						this_line_OK = 0
 					else:
 						dir_ok.append("Mismatch (model ns.)")
 						this_line_OK = 0
 						
 				if((exp_av-ct_av > 0) and change_dir == 'no change'):
 					if(stat_results[indnow]<= 5.00e-02):
 						dir_ok.append("Mismatch, increase")
 						this_line_OK = 0
 					else:
 						dir_ok.append("Match (ns increase)")
 						some_OK = 1
 						tests_OK = tests_OK + 1

 				if((exp_av-ct_av < 0) and change_dir == '0'):
 					if(stat_results[indnow]<= 5.00e-02):
 						dir_ok.append("Significant")
 						some_OK = 1
 						tests_OK = tests_OK + 1
 					else:
 						dir_ok.append("Dir. OK but Not Sig.")
 						this_line_OK = 0
 						
 				if((exp_av-ct_av < 0) and change_dir == '1'):
 					if(stat_results[indnow]<= 5.00e-02):
 						dir_ok.append("Mismatch (sign.)")
 						this_line_OK = 0
 					else:
 						dir_ok.append("Mismatch (model ns.)")
 						this_line_OK = 0
 
 				if((exp_av-ct_av < 0) and change_dir == 'no change'):
 					if(stat_results[indnow]<= 5.00e-02):
 						dir_ok.append("Mismatch, decrease")
 						this_line_OK = 0
 					else:
 						dir_ok.append("Match (ns decrease)")
 						some_OK = 1
 						tests_OK = tests_OK + 1
 						
 				if((exp_av-ct_av == 0) and change_dir == 'no change'):
  					dir_ok.append("Match (no change)")
  					some_OK = 1
  					tests_OK = tests_OK + 1
 				if((exp_av-ct_av == 0) and change_dir == '1'):
 					dir_ok.append("Mismatch (inc. not reproduced)")
 					this_line_OK = 0
 				if((exp_av-ct_av == 0) and change_dir == '0'):
 					dir_ok.append("Mismatch (dec. not reproduced)")
 					this_line_OK = 0

			line_OK = line_OK + this_line_OK
			some_in_line_OK = some_in_line_OK + some_OK
			
			#print(change_dir)
			#print(stat_results)
			#print(dir_ok)
			#print(output_phen)
			for ind in range(len(output_phen)):
				Stat_tests_output.append([node_st, pert_type_str, output_phen[ind],'',change_dir,dir_ok[ind],stat_results[ind]])
				if(dir_ok[ind] == "Dir. OK but Not Sig." or dir_ok[ind] == "Mismatch, increase" or dir_ok[ind] == "Mismatch, decrease" or dir_ok[ind] == "Mismatch (inc. not reproduced)" or dir_ok[ind] == "Mismatch (inc. not reproduced)" or dir_ok[ind] == "Mismatch (model ns.)"):
					BAD_tests_output.append([node_st, pert_type_str, output_phen[ind],'',change_dir,dir_ok[ind],stat_results[ind]])
	


def save_multi_image(filename):
	pp = PdfPages(filename)
	fig_nums = plt.get_fignums()
	figs = [plt.figure(n) for n in fig_nums]
	for fig in figs:
		fig.savefig(pp, format='pdf')
	pp.close()

save_multi_image(model_folder + "Suppl File 3 - Large Model Validation Figures.pdf")
df = pd.DataFrame(Stat_tests_output)
print(df.to_string())
df.to_csv('Large Model Validation Statistics.csv')

print("\n\nLines that match for all output nodes/phenotypes: " + str(line_OK) + " of " + str(N_lines) + " => " + str("%.2f" % float(line_OK/N_lines))+ "% match.")
print("Lines in which at least one output node/phenotype matches: " + str(some_in_line_OK) + " of " + str(N_lines) + " => " + str("%.2f" % float(some_in_line_OK/N_lines))+ "% match.")	
print("Number of unique statistical tests that match: " + str(tests_OK) + " of " + str(N_tests) + " => " + str("%.2f" % float(tests_OK/N_tests))+ "% match.")	

print("\n\nFailed tests:")
df2 = pd.DataFrame(BAD_tests_output)
print(df2.to_string())
```
