## Supplementary material for "HHIP’s Dynamic Role in Epithelial Wound Healing Reveals a Potential Mechanism of COPD Susceptibility": Suppl File 3 - Large Model Validation Figures.pdf

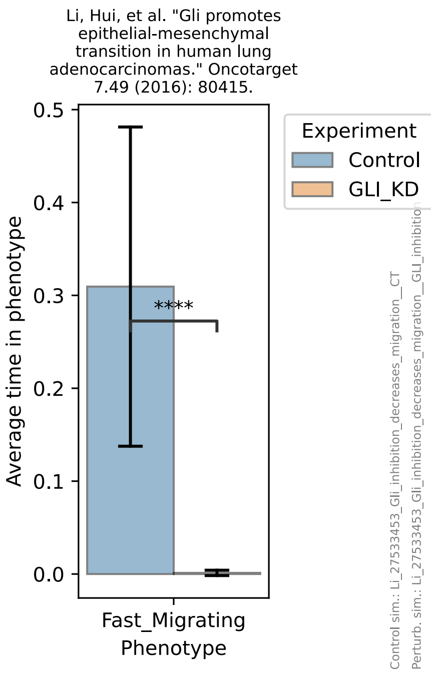

Li, Hui, et al. "Gli promotes epithelial-mesenchymal transition in human lung adenocarcinomas." Oncotarget 7.49 (2016): 80415.
Average time in phenotype
0.0
0.1
0.2
0.3
0.4
0.5
****
Fast_Migrating
Phenotype
Experiment
Control
GLI_KD
Control sim.: Li_27533453_Gli_inhibition_decreases_migration__CT
Perturb. sim.: Li_27533453_Gli_inhibition_decreases_migration__GLI_inhibitio


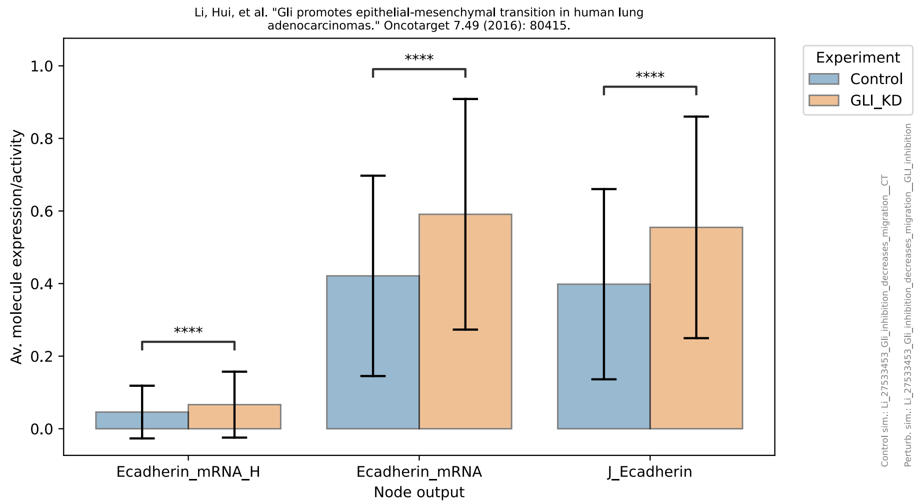

Li, Hui, et al. "Gli promotes epithelial-mesenchymal transition in human lung adenocarcinomas." Oncotarget 7.49 (2016): 80415.
Av. molecule expression/activity
0.0
0.2
0.4
0.6
0.8
1.0
****
****
****
Ecadherin_mRNA_H
Ecadherin_mRNA
J_Ecadherin
Node output
Experiment
Control
GLI_KD
Control sim.: Li_27533453_Gli_inhibition_decreases_migration__CT
Perturb. sim.: Li_27533453_Gli_inhibition_decreases_migration__GLI_inhibition


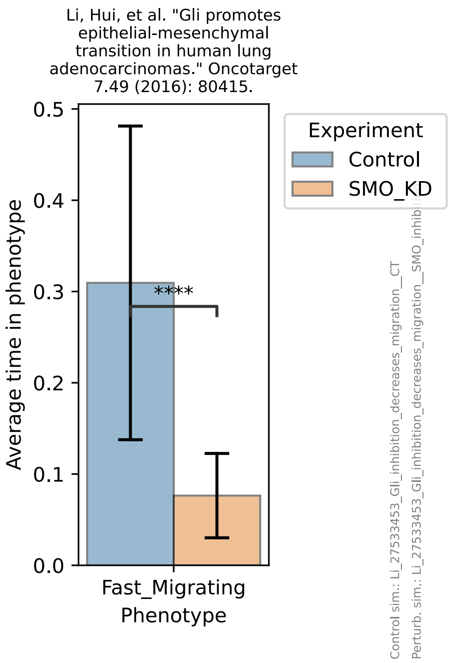

Li, Hui, et al. "Gli promotes epithelial-mesenchymal transition in human lung adenocarcinomas." Oncotarget 7.49 (2016): 80415.
Average time in phenotype
0.0
0.1
0.2
0.3
0.4
0.5
****
Fast_Migrating
Phenotype
Experiment
Control
SMO_KD
Control sim.: Li_27533453_Gli_inhibition_decreases_migration__CT
Perturb. sim.: Li_27533453_Gli_inhibition_decreases_migration__SMO_inhibition


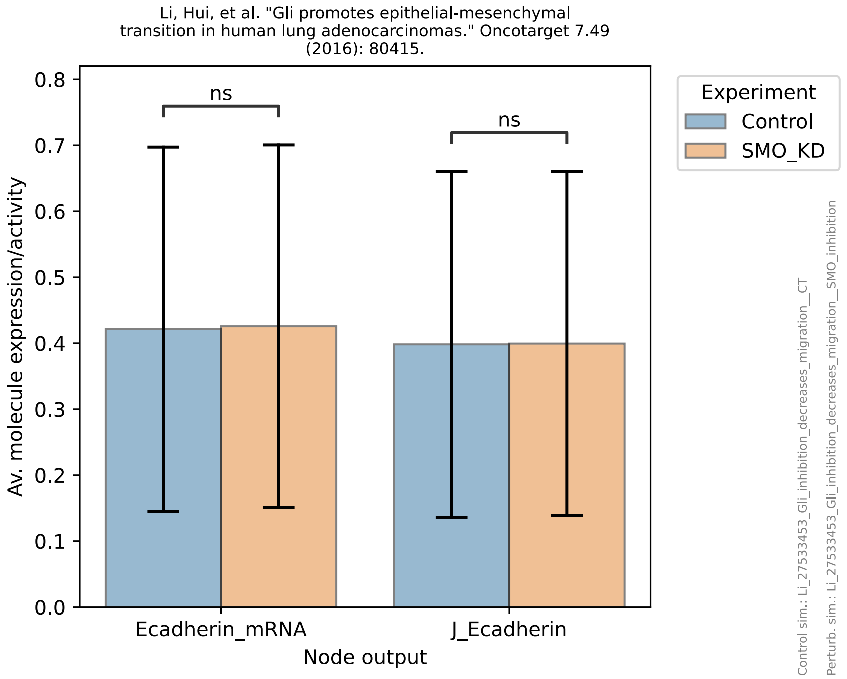

Li, Hui, et al. "Gli promotes epithelial-mesenchymal transition in human lung adenocarcinomas." Oncotarget 7.49 (2016): 80415.
Av. molecule expression/activity
0.0
0.1
0.2
0.3
0.4
0.5
0.6
0.7
0.8
ns
ns
Ecadherin_mRNA
J_Ecadherin
Node output
Experiment
Control
SMO_KD
Control sim.: Li_27533453_Gli_inhibition_decreases_migration__CT
Perturb. sim.: Li_27533453_Gli_inhibition_decreases_migration__SMO_inhibition

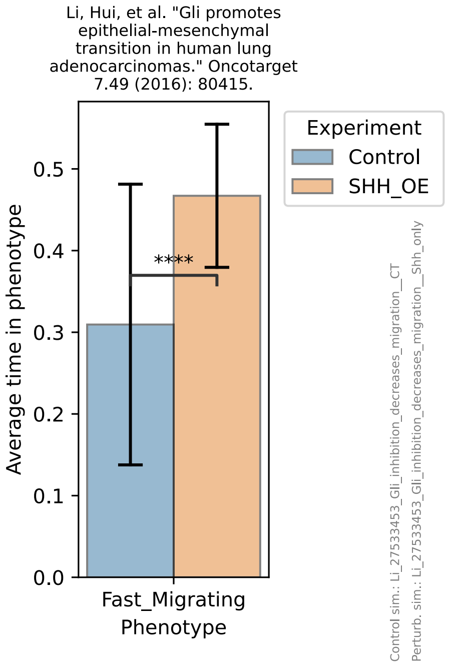
Li, Hui, et al. "Gli promotes epithelial-mesenchymal transition in human lung adenocarcinomas." Oncotarget 7.49 (2016): 80415.
Average time in phenotype
****
Fast_Migrating Phenotype
Experiment
Control
SHH_OE
Control sim.: Li_27533453_Gli_inhibition_decreases_migration__CT
Perturb. sim.: Li_27533453_Gli_inhibition_decreases_migration__Shh_only

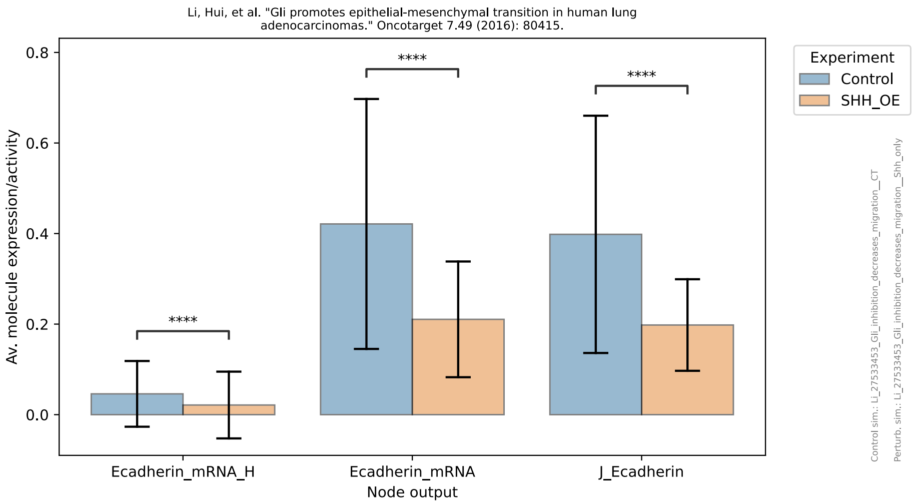
Li, Hui, et al. "Gli promotes epithelial-mesenchymal transition in human lung adenocarcinomas." Oncotarget 7.49 (2016): 80415.
Av. molecule expression/activity
****
****
****
Ecadherin_mRNA_H
Ecadherin_mRNA
J_Ecadherin
Node output
Experiment
Control
SHH_OE
Control sim.: Li_27533453_Gli_inhibition_decreases_migration__CT
Perturb. sim.: Li_27533453_Gli_inhibition_decreases_migration__Shh_only

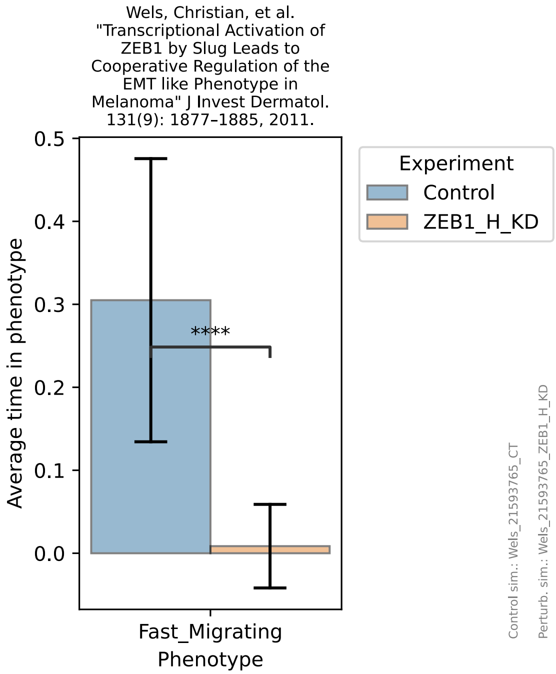
Wels, Christian, et al. "Transcriptional Activation of ZEB1 by Slug Leads to Cooperative Regulation of the EMT like Phenotype in Melanoma" J Invest Dermatol. 131(9): 1877–1885, 2011.
Average time in phenotype
****
Fast_Migrating Phenotype
Experiment
Control
ZEB1_H_KD
Control sim.: Wels_21593765_CT
Perturb. sim.: Wels_21593765_ZEB1_H_KD

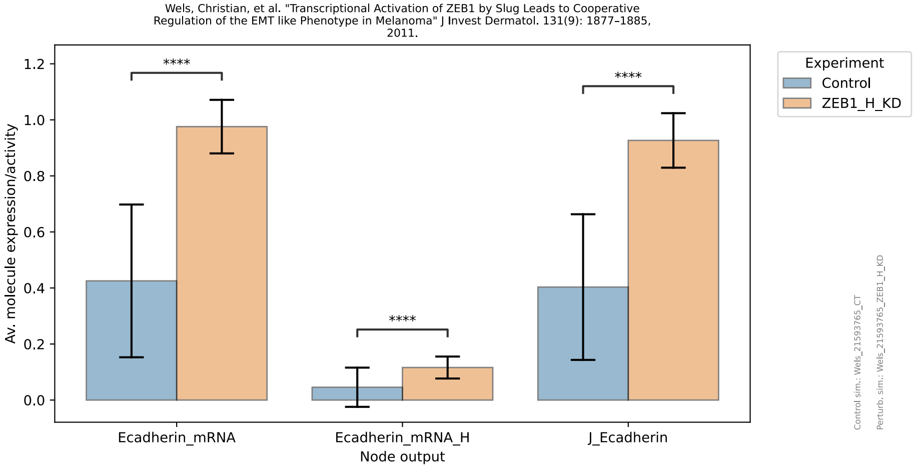
Wels, Christian, et al. "Transcriptional Activation of ZEB1 by Slug Leads to Cooperative Regulation of the EMT like Phenotype in Melanoma" J Invest Dermatol. 131(9): 1877–1885, 2011.
Av. molecule expression/activity
****
****
****
Ecadherin_mRNA
Ecadherin_mRNA_H
J_Ecadherin
Node output
Experiment
Control
ZEB1_H_KD
Control sim.: Wels_21593765_CT
Perturb. sim.: Wels_21593765_ZEB1_H_KD

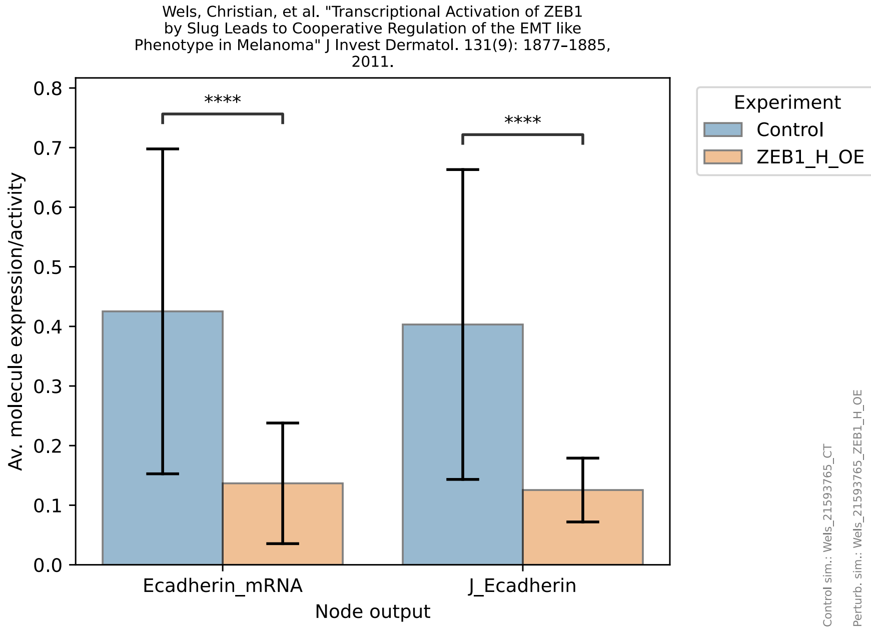
Wels, Christian, et al. "Transcriptional Activation of ZEB1 by Slug Leads to Cooperative Regulation of the EMT like Phenotype in Melanoma" J Invest Dermatol. 131(9): 1877–1885, 2011.
Av. molecule expression/activity
0.0
0.1
0.2
0.3
0.4
0.5
0.6
0.7
0.8
****
****
Ecadherin_mRNA
J_Ecadherin
Node output
Experiment
Control
ZEB1_H_OE
Control sim.: Wels_21593765_CT
Perturb. sim.: Wels_21593765_ZEB1_H_OE

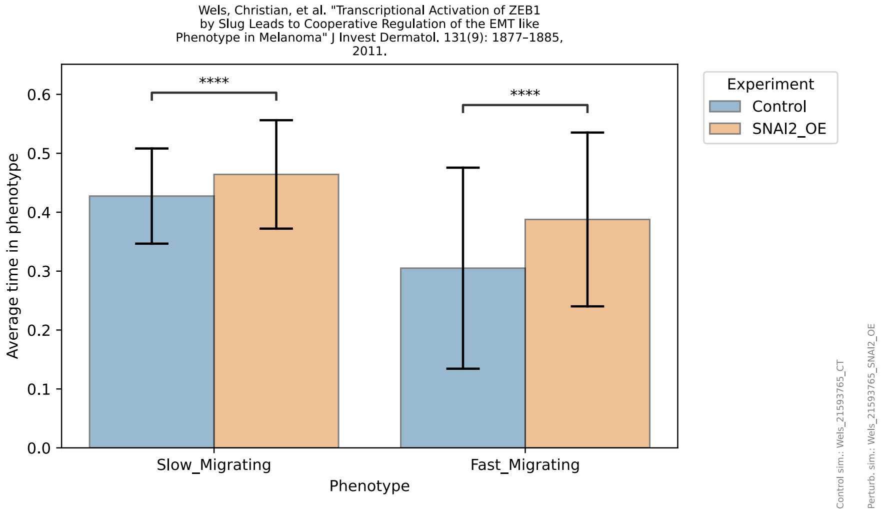
Wels, Christian, et al. "Transcriptional Activation of ZEB1 by Slug Leads to Cooperative Regulation of the EMT like Phenotype in Melanoma" J Invest Dermatol. 131(9): 1877–1885, 2011.
Average time in phenotype
0.0
0.1
0.2
0.3
0.4
0.5
0.6
****
****
Slow_Migrating
Fast_Migrating
Phenotype
Experiment
Control
SNAI2_OE
Control sim.: Wels_21593765_CT
Perturb. sim.: Wels_21593765_SNAI2_OE

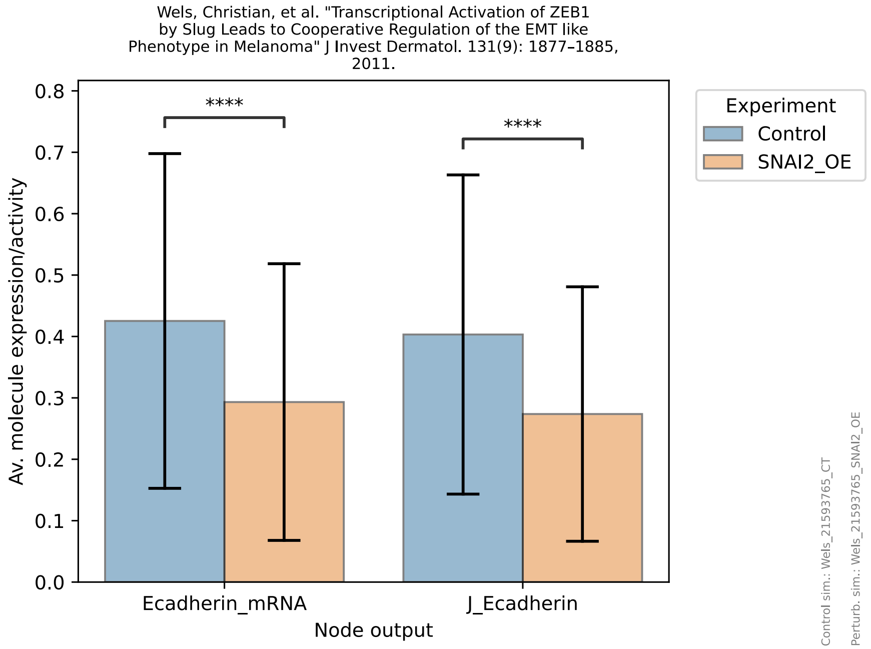
Wels, Christian, et al. "Transcriptional Activation of ZEB1 by Slug Leads to Cooperative Regulation of the EMT like Phenotype in Melanoma" J Invest Dermatol. 131(9): 1877–1885, 2011.
Av. molecule expression/activity
0.0
0.1
0.2
0.3
0.4
0.5
0.6
0.7
0.8
****
****
Ecadherin_mRNA
J_Ecadherin
Node output
Experiment
Control
SNAI2_OE
Control sim.: Wels_21593765_CT
Perturb. sim.: Wels_21593765_SNAI2_OE

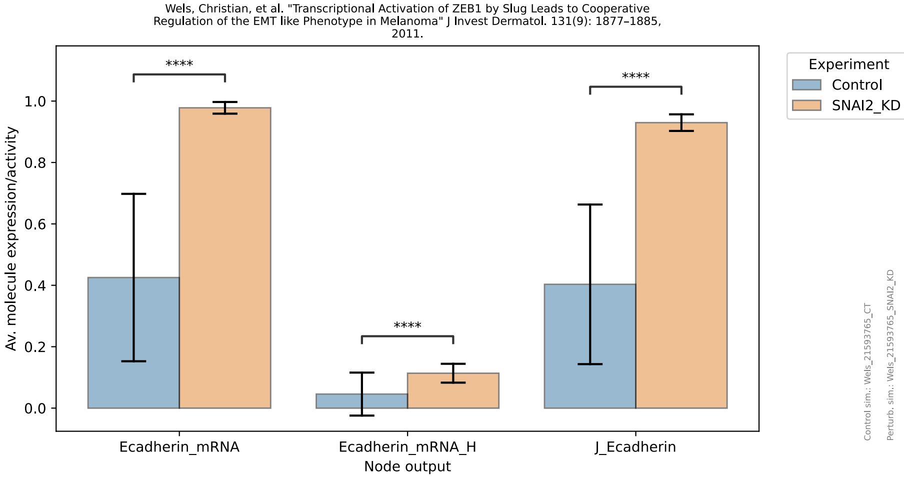
Wels, Christian, et al. "Transcriptional Activation of ZEB1 by Slug Leads to Cooperative Regulation of the EMT like Phenotype in Melanoma" J Invest Dermatol. 131(9): 1877–1885, 2011.
Av. molecule expression/activity
0.0
0.2
0.4
0.6
0.8
1.0
****
****
****
Ecadherin_mRNA
Ecadherin_mRNA_H
J_Ecadherin
Node output
Experiment
Control
SNAI2_KD
Control sim.: Wels_21593765_CT
Perturb. sim.: Wels_21593765_SNAI2_KD

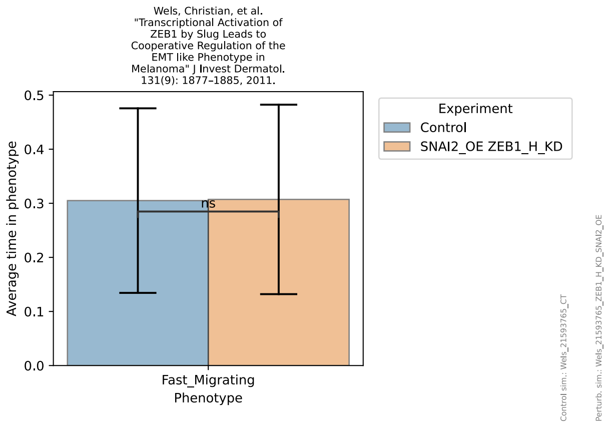
Wels, Christian, et al. "Transcriptional Activation of ZEB1 by Slug Leads to Cooperative Regulation of the EMT like Phenotype in Melanoma" J Invest Dermatol. 131(9): 1877–1885, 2011.
Average time in phenotype
0.0
0.1
0.2
0.3
0.4
0.5
ns
Fast_Migrating
Phenotype
Experiment
Control
SNAI2_OE ZEB1_H_KD
Control sim.: Wels_21593765_CT
Perturb. sim.: Wels_21593765_ZEB1_H_KD_SNAI2_OE

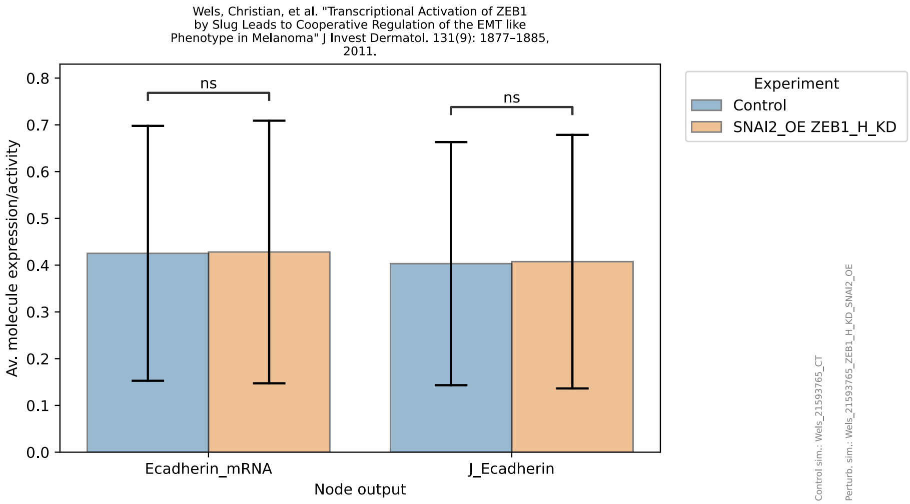
Wels, Christian, et al. "Transcriptional Activation of ZEB1 by Slug Leads to Cooperative Regulation of the EMT like Phenotype in Melanoma" J Invest Dermatol. 131(9): 1877–1885, 2011.
Av. molecule expression/activity
0.0
0.1
0.2
0.3
0.4
0.5
0.6
0.7
0.8
ns
ns
Ecadherin_mRNA
J_Ecadherin
Node output
Experiment
Control
SNAI2_OE ZEB1_H_KD
Control sim.: Wels_21593765_CT
Perturb. sim.: Wels_21593765_ZEB1_H_KD_SNAI2_OE

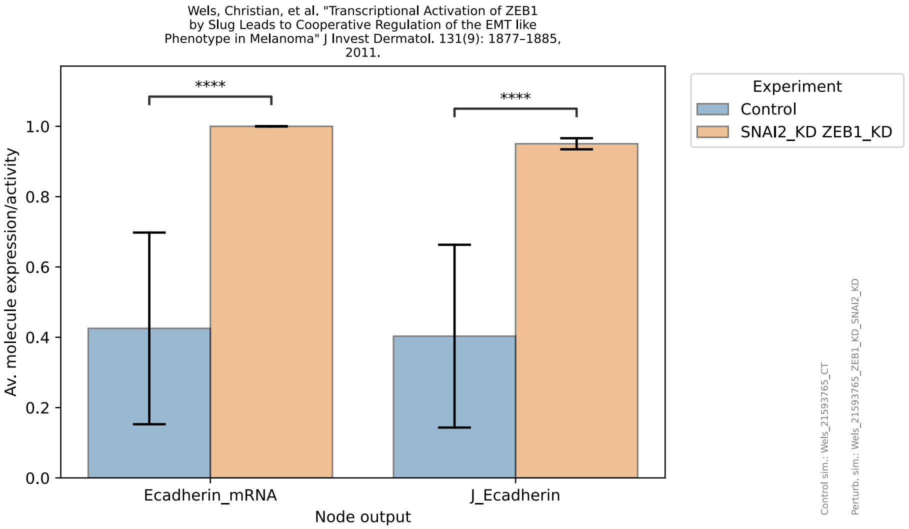
Wels, Christian, et al. "Transcriptional Activation of ZEB1 by Slug Leads to Cooperative Regulation of the EMT like Phenotype in Melanoma" J Invest Dermatol. 131(9): 1877–1885, 2011.
Av. molecule expression/activity
0.0
0.2
0.4
0.6
0.8
1.0
****
****
Ecadherin_mRNA
J_Ecadherin
Node output
Experiment
Control
SNAI2_KD ZEB1_KD
Control sim.: Wels_21593765_CT
Perturb. sim.: Wels_21593765_ZEB1_KD_SNAI2_KD

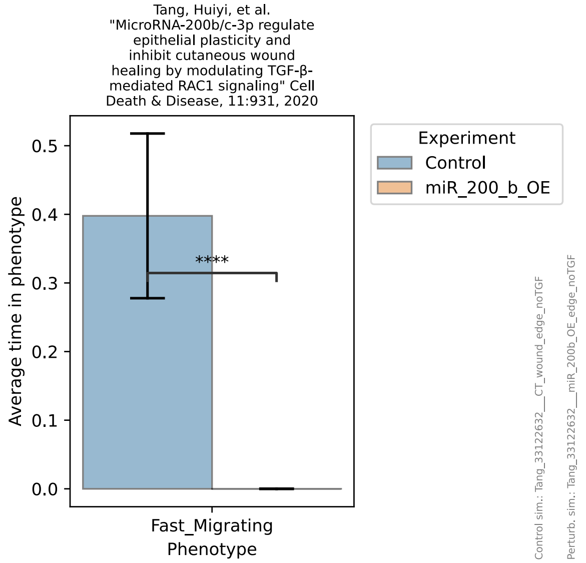
Tang, Huiyi, et al. "MicroRNA-200b/c-3p regulate epithelial plasticity and inhibit cutaneous wound healing by modulating TGF-β-mediated RAC1 signaling" Cell Death & Disease, 11:931, 2020
Average time in phenotype
0.0
0.1
0.2
0.3
0.4
0.5
****
Fast_Migrating
Phenotype
Experiment
Control
miR_200_b_OE
Control sim.: Tang_33122632__CT_wound_edge_noTGF
Perturb. sim.: Tang_33122632__miR_200b_OE_edge_noTGF

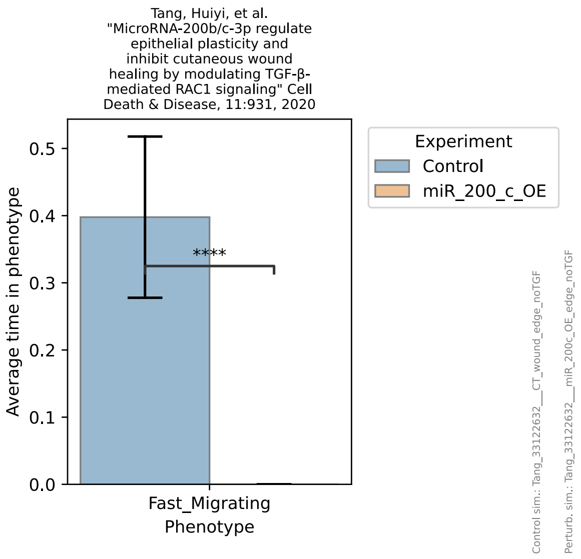
Tang, Huiyi, et al. "MicroRNA-200b/c-3p regulate epithelial plasticity and inhibit cutaneous wound healing by modulating TGF-β-mediated RAC1 signaling" Cell Death & Disease, 11:931, 2020
Experiment
Control
miR_200_c_OE
****
Average time in phenotype
0.0
0.1
0.2
0.3
0.4
0.5
Fast_Migrating
Phenotype
Control sim.: Tang_33122632___CT_wound_edge_noTGF
Perturb. sim.: Tang_33122632___miR_200c_OE_edge_noTGF

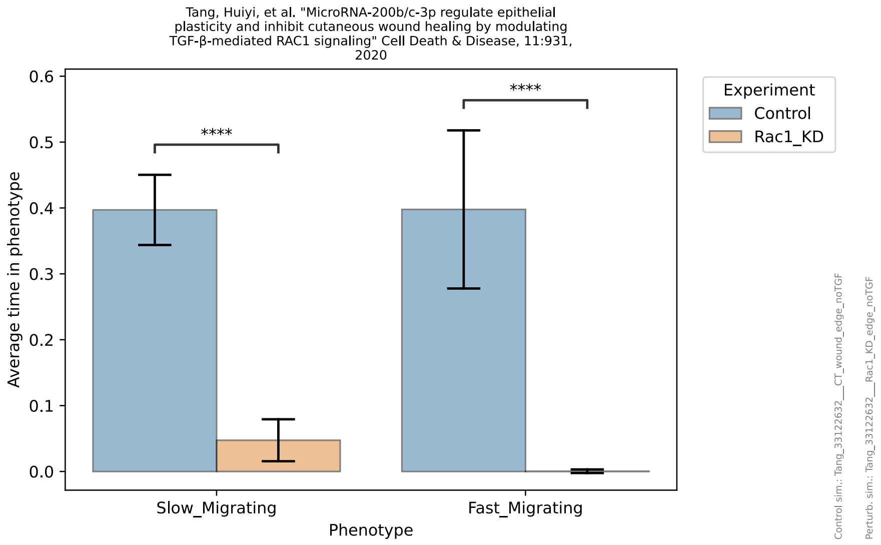
Tang, Huiyi, et al. "MicroRNA-200b/c-3p regulate epithelial plasticity and inhibit cutaneous wound healing by modulating TGF-β-mediated RAC1 signaling" Cell Death & Disease, 11:931, 2020
Experiment
Control
Rac1_KD
****
****
Average time in phenotype
0.0
0.1
0.2
0.3
0.4
0.5
0.6
Slow_Migrating
Fast_Migrating
Phenotype
Control sim.: Tang_33122632___CT_wound_edge_noTGF
Perturb. sim.: Tang_33122632___Rac1_KD_edge_noTGF

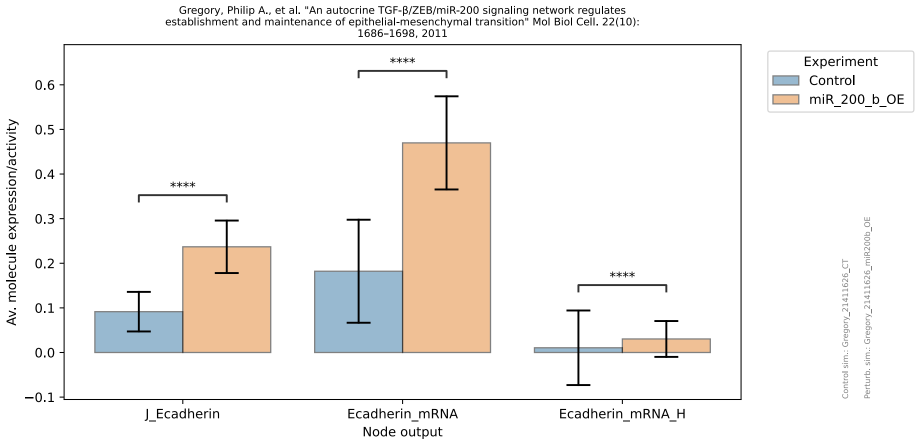
Gregory, Philip A., et al. "An autocrine TGF-β/ZEB/miR-200 signaling network regulates establishment and maintenance of epithelial-mesenchymal transition" Mol Biol Cell. 22(10): 1686–1698, 2011
Experiment
Control
miR_200_b_OE
****
****
****
Av. molecule expression/activity
−0.1
0.0
0.1
0.2
0.3
0.4
0.5
0.6
J_Ecadherin
Ecadherin_mRNA
Ecadherin_mRNA_H
Node output
Control sim.: Gregory_21411626_CT
Perturb. sim.: Gregory_21411626_miR200b_OE

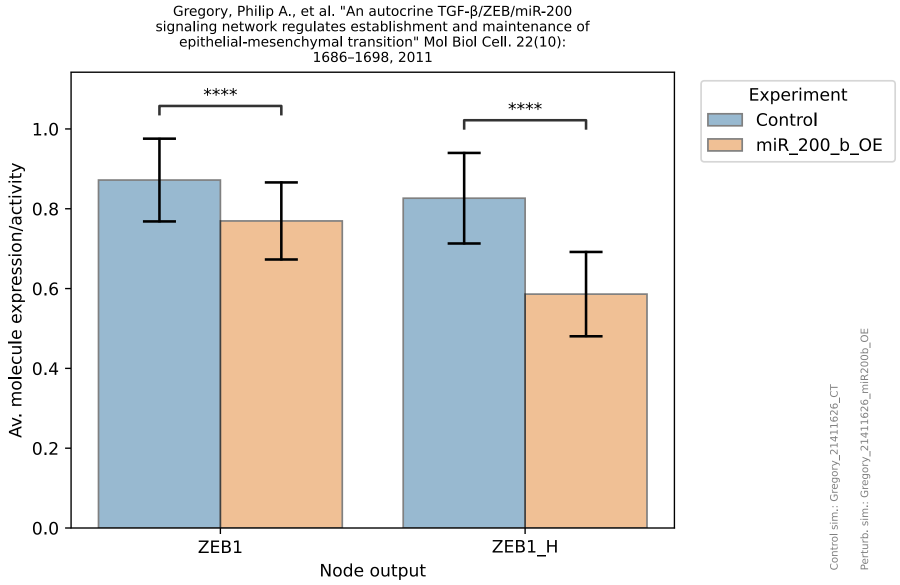
Gregory, Philip A., et al. "An autocrine TGF-β/ZEB/miR-200 signaling network regulates establishment and maintenance of epithelial-mesenchymal transition" Mol Biol Cell. 22(10): 1686–1698, 2011
Experiment
Control
miR_200_b_OE
****
****
Av. molecule expression/activity
0.0
0.2
0.4
0.6
0.8
1.0
ZEB1
ZEB1_H
Node output
Control sim.: Gregory_21411626_CT
Perturb. sim.: Gregory_21411626_miR200b_OE

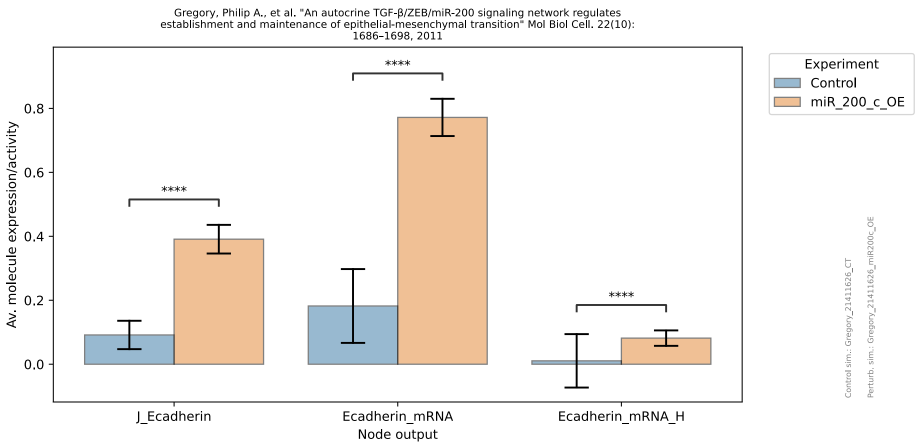
Gregory, Philip A., et al. "An autocrine TGF-β/ZEB/miR-200 signaling network regulates establishment and maintenance of epithelial-mesenchymal transition" Mol Biol Cell. 22(10): 1686–1698, 2011
Av. molecule expression/activity
Node output
J_Ecadherin
Ecadherin_mRNA
Ecadherin_mRNA_H
****
Experiment
Control
miR_200_c_OE
Control sim.: Gregory_21411626_CT
Perturb. sim.: Gregory_21411626_miR200c_OE

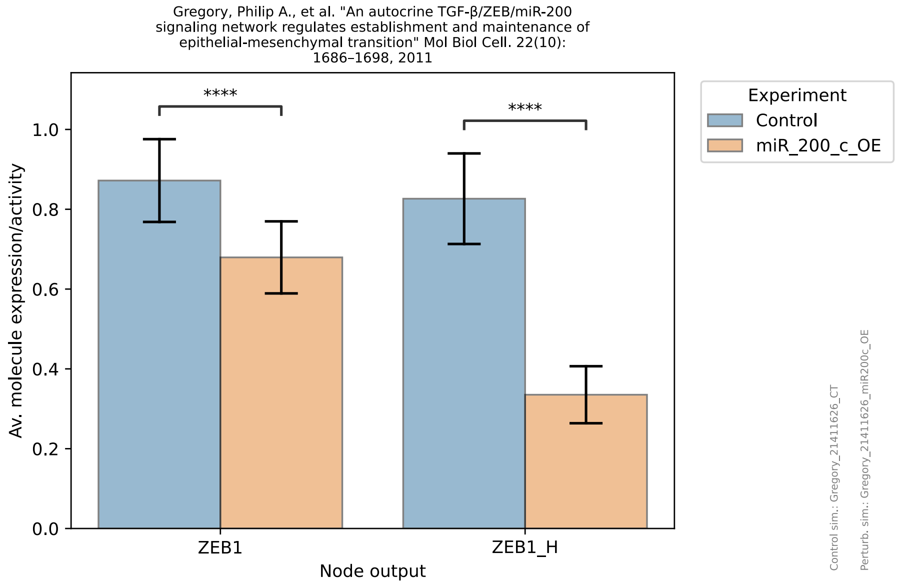
Gregory, Philip A., et al. "An autocrine TGF-β/ZEB/miR-200 signaling network regulates establishment and maintenance of epithelial-mesenchymal transition" Mol Biol Cell. 22(10): 1686–1698, 2011
Av. molecule expression/activity
Node output
ZEB1
ZEB1_H
****
Experiment
Control
miR_200_c_OE
Control sim.: Gregory_21411626_CT
Perturb. sim.: Gregory_21411626_miR200c_OE

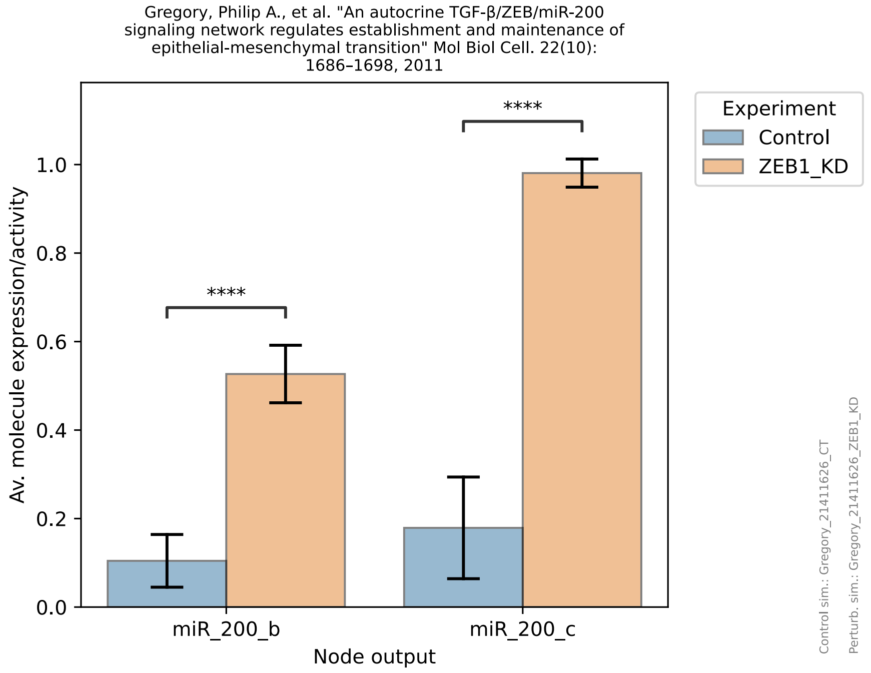
Gregory, Philip A., et al. "An autocrine TGF-β/ZEB/miR-200 signaling network regulates establishment and maintenance of epithelial-mesenchymal transition" Mol Biol Cell. 22(10): 1686–1698, 2011
Av. molecule expression/activity
Node output
miR_200_b
miR_200_c
****
Experiment
Control
ZEB1_KD
Control sim.: Gregory_21411626_CT
Perturb. sim.: Gregory_21411626_ZEB1_KD

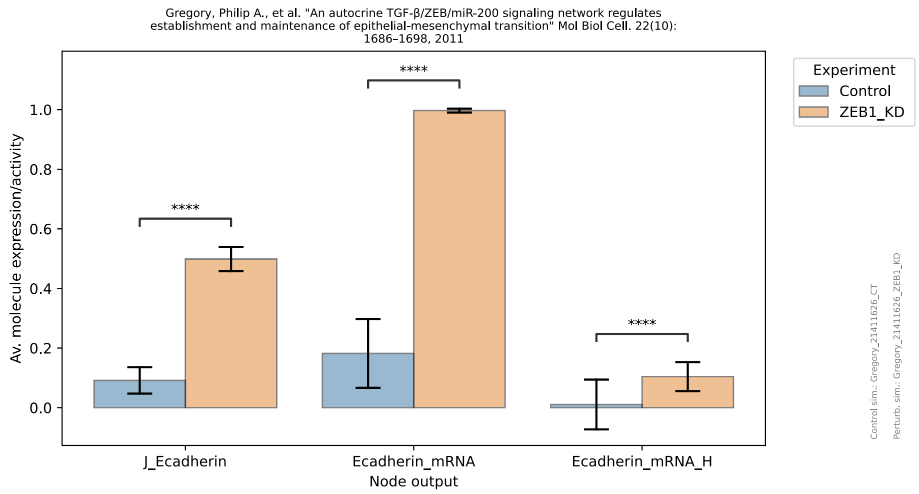
Gregory, Philip A., et al. "An autocrine TGF-β/ZEB/miR-200 signaling network regulates establishment and maintenance of epithelial-mesenchymal transition" Mol Biol Cell. 22(10): 1686–1698, 2011
Av. molecule expression/activity
Node output
J_Ecadherin
Ecadherin_mRNA
Ecadherin_mRNA_H
****
Experiment
Control
ZEB1_KD
Control sim.: Gregory_21411626_CT
Perturb. sim.: Gregory_21411626_ZEB1_KD

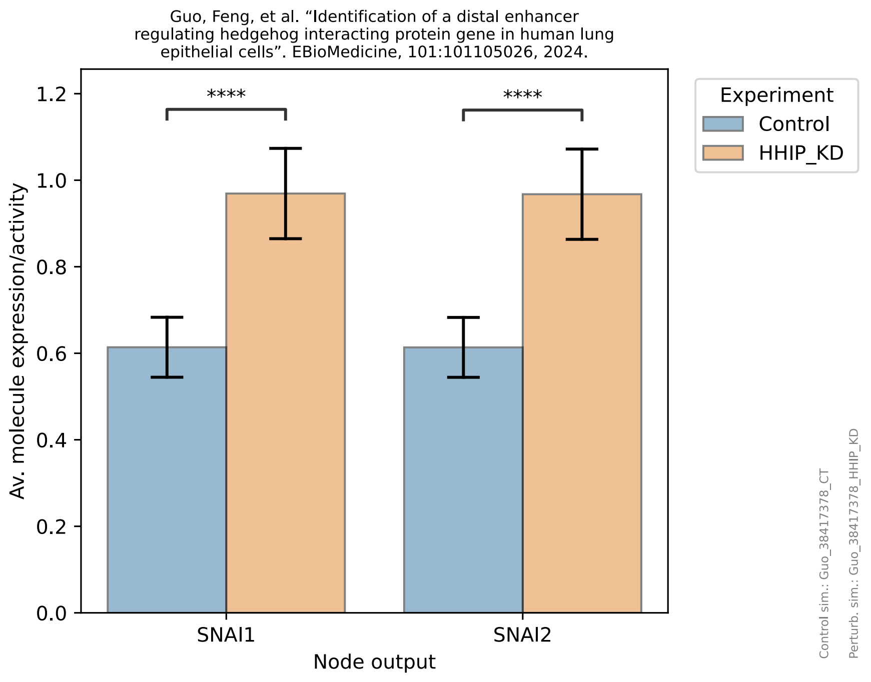
Guo, Feng, et al. "Identification of a distal enhancer regulating hedgehog interacting protein gene in human lung epithelial cells". EBioMedicine, 101:101105026, 2024.
Experiment
Control
HHIP_KD
****
****
Av. molecule expression/activity
SNAI1
SNAI2
Node output
Control sim.: Guo_38417378_CT
Perturb. sim.: Guo_38417378_HHIP_KD

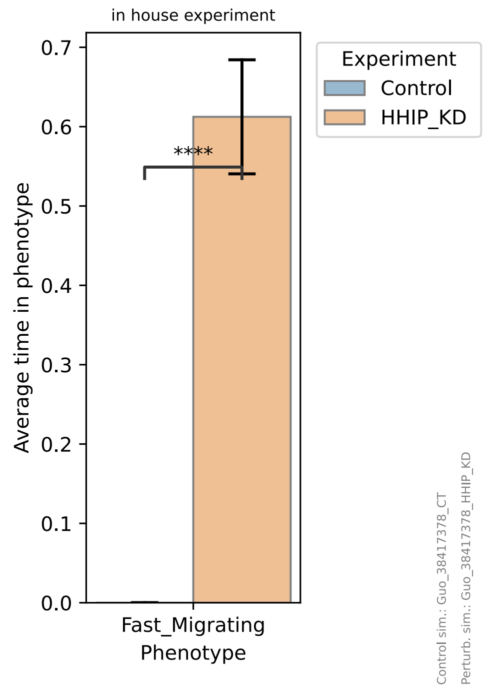
in house experiment
Experiment
Control
HHIP_KD
****
Average time in phenotype
Fast_Migrating
Phenotype
Control sim.: Guo_38417378_CT
Perturb. sim.: Guo_38417378_HHIP_KD

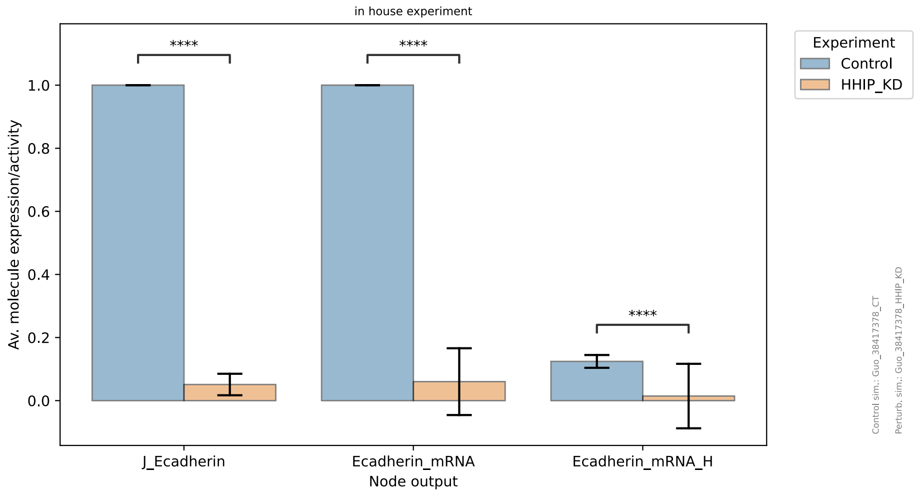
in house experiment
Experiment
Control
HHIP_KD
****
****
****
Av. molecule expression/activity
J_Ecadherin
Ecadherin_mRNA
Ecadherin_mRNA_H
Node output
Control sim.: Guo_38417378_CT
Perturb. sim.: Guo_38417378_HHIP_KD

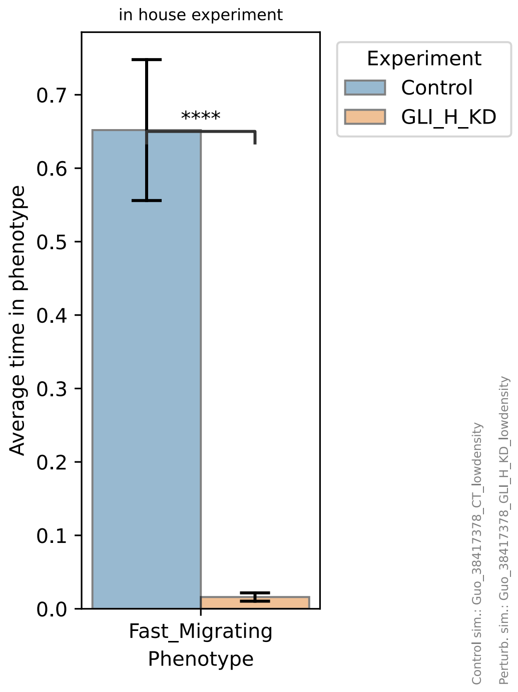
in house experiment
Experiment
Control
GLI_H_KD
****
Average time in phenotype
Fast_Migrating
Phenotype
Control sim.: Guo_38417378_CT_lowdensity
Perturb. sim.: Guo_38417378_GLI_H_KD_lowdensity

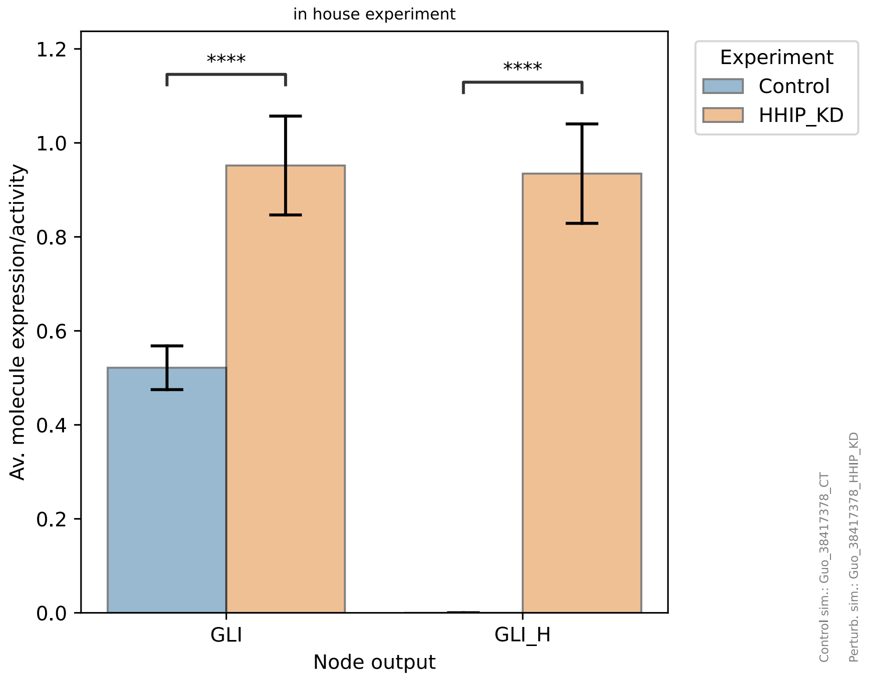
in house experiment
****
****
Experiment
Control
HHIP_KD
Av. molecule expression/activity
0.0
0.2
0.4
0.6
0.8
1.0
1.2
GLI
GLI_H
Node output
Control sim.: Guo_38417378_CT
Perturb. sim.: Guo_38417378_HHIP_KD

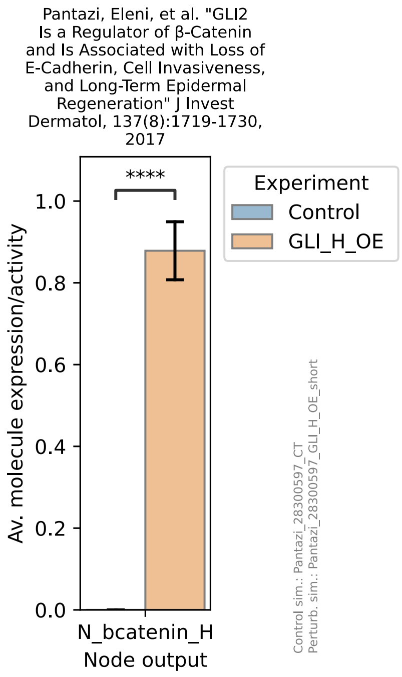
Pantazi, Eleni, et al. "GLI2 Is a Regulator of β-Catenin and Is Associated with Loss of E-Cadherin, Cell Invasiveness, and Long-Term Epidermal Regeneration" J Invest Dermatol, 137(8):1719-1730, 2017
****
Experiment
Control
GLI_H_OE
Av. molecule expression/activity
0.0
0.2
0.4
0.6
0.8
1.0
N_bcatenin_H
Node output
Control sim.: Pantazi_28300597_CT
Perturb. sim.: Pantazi_28300597_GLI_H_OE_short

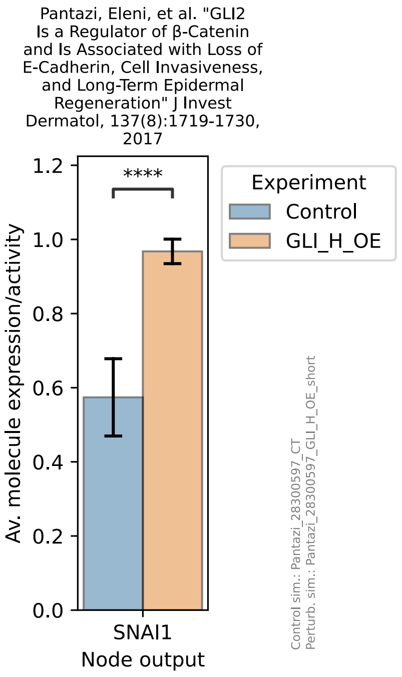
Pantazi, Eleni, et al. "GLI2 Is a Regulator of β-Catenin and Is Associated with Loss of E-Cadherin, Cell Invasiveness, and Long-Term Epidermal Regeneration" J Invest Dermatol, 137(8):1719-1730, 2017
****
Experiment
Control
GLI_H_OE
Av. molecule expression/activity
0.0
0.2
0.4
0.6
0.8
1.0
1.2
SNAI1
Node output
Control sim.: Pantazi_28300597_CT
Perturb. sim.: Pantazi_28300597_GLI_H_OE_short

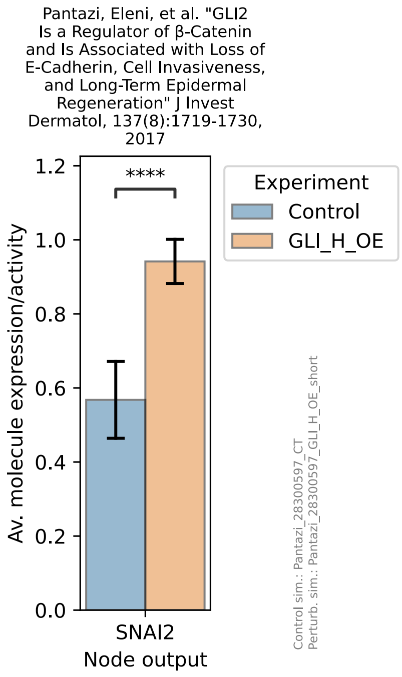
Pantazi, Eleni, et al. "GLI2 Is a Regulator of β-Catenin and Is Associated with Loss of E-Cadherin, Cell Invasiveness, and Long-Term Epidermal Regeneration" J Invest Dermatol, 137(8):1719-1730, 2017
****
Experiment
Control
GLI_H_OE
Av. molecule expression/activity
0.0
0.2
0.4
0.6
0.8
1.0
1.2
SNAI2
Node output
Control sim.: Pantazi_28300597_CT
Perturb. sim.: Pantazi_28300597_GLI_H_OE_short

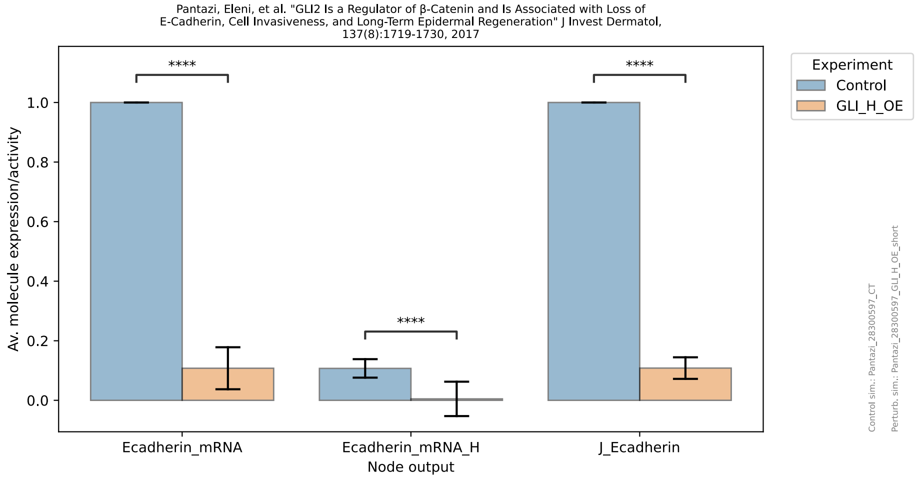
Pantazi, Eleni, et al. "GLI2 Is a Regulator of β-Catenin and Is Associated with Loss of E-Cadherin, Cell Invasiveness, and Long-Term Epidermal Regeneration" J Invest Dermatol, 137(8):1719-1730, 2017
Av. molecule expression/activity
Node output
Ecadherin_mRNA
Ecadherin_mRNA_H
J_Ecadherin
****
Experiment
Control
GLI_H_OE
Control sim.: Pantazi_28300597_CT
Perturb. sim.: Pantazi_28300597_GLI_H_OE_short

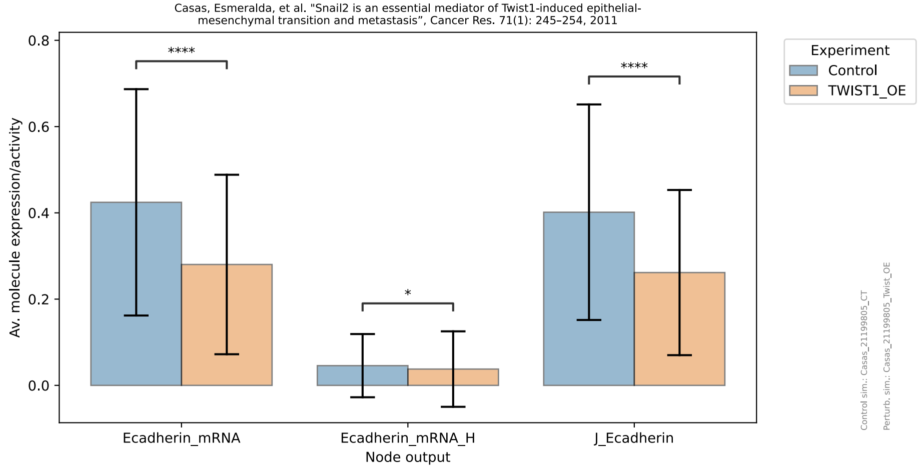
Casas, Esmeralda, et al. "Snail2 is an essential mediator of Twist1-induced epithelial-mesenchymal transition and metastasis", Cancer Res. 71(1): 245–254, 2011
Av. molecule expression/activity
Node output
Ecadherin_mRNA
Ecadherin_mRNA_H
J_Ecadherin
****
*
Experiment
Control
TWIST1_OE
Control sim.: Casas_21199805_CT
Perturb. sim.: Casas_21199805_Twist_OE

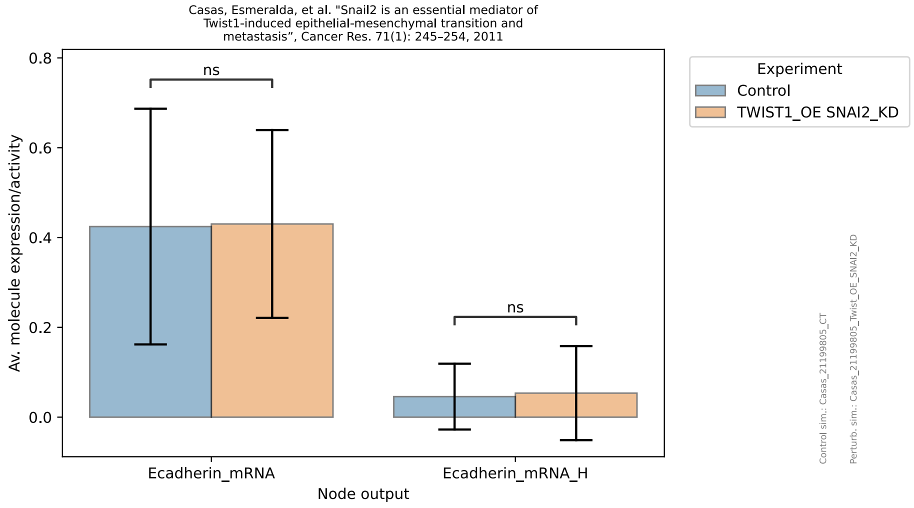
Casas, Esmeralda, et al. "Snail2 is an essential mediator of Twist1-induced epithelial-mesenchymal transition and metastasis", Cancer Res. 71(1): 245–254, 2011
Av. molecule expression/activity
Node output
Ecadherin_mRNA
Ecadherin_mRNA_H
ns
Experiment
Control
TWIST1_OE SNAI2_KD
Control sim.: Casas_21199805_CT
Perturb. sim.: Casas_21199805_Twist_OE_SNAI2_KD

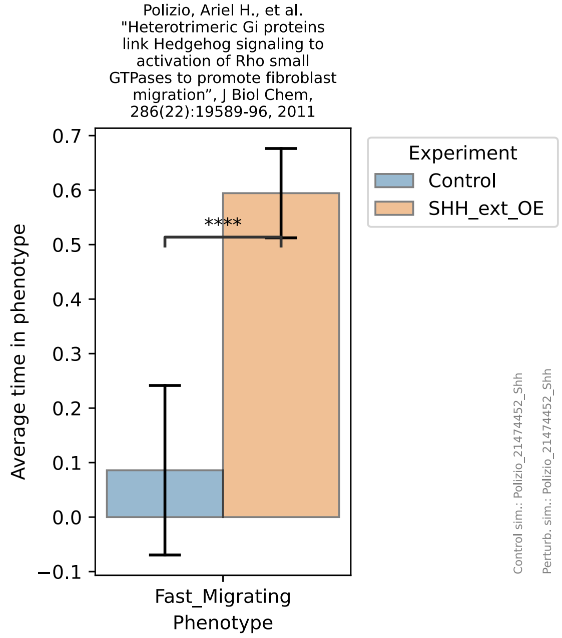
Polizio, Ariel H., et al. "Heterotrimeric Gi proteins link Hedgehog signaling to activation of Rho small GTPases to promote fibroblast migration", J Biol Chem, 286(22):19589-96, 2011
Average time in phenotype
Fast_Migrating Phenotype
****
Experiment
Control
SHH_ext_OE
Control sim.: Polizio_21474452_Shh
Perturb. sim.: Polizio_21474452_Shh

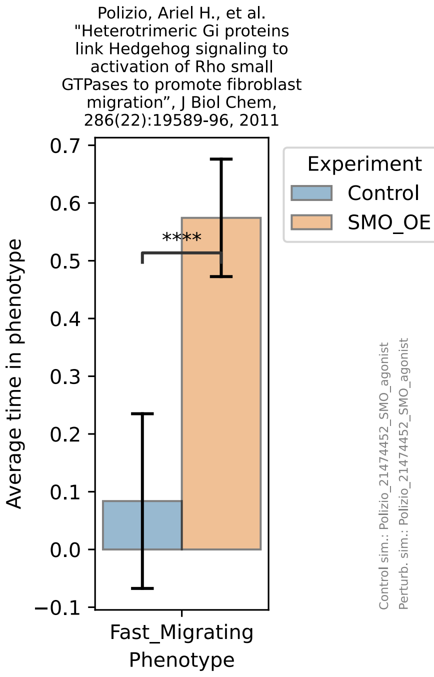
Polizio, Ariel H., et al. "Heterotrimeric Gi proteins link Hedgehog signaling to activation of Rho small GTPases to promote fibroblast migration", J Biol Chem, 286(22):19589-96, 2011
Average time in phenotype
0.7
0.6
0.5
0.4
0.3
0.2
0.1
0.0
−0.1
****
Fast_Migrating
Phenotype
Experiment
Control
SMO_OE
Control sim.: Polizio_21474452_SMO_agonist
Perturb. sim.: Polizio_21474452_SMO_agonist

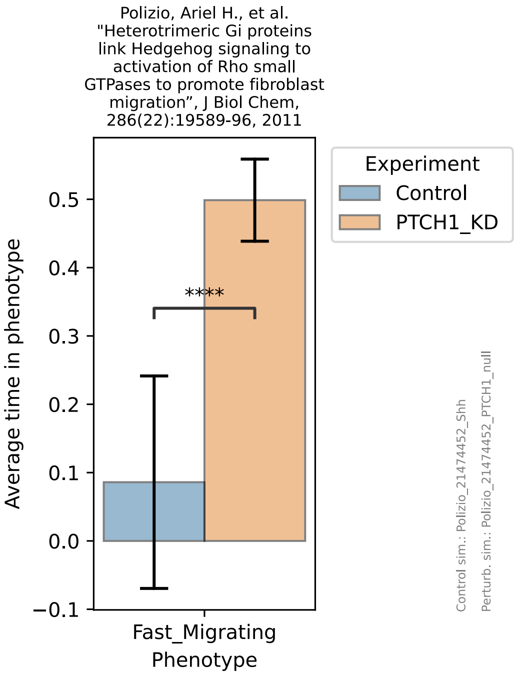
Polizio, Ariel H., et al. "Heterotrimeric Gi proteins link Hedgehog signaling to activation of Rho small GTPases to promote fibroblast migration", J Biol Chem, 286(22):19589-96, 2011
Average time in phenotype
0.5
0.4
0.3
0.2
0.1
0.0
−0.1
****
Fast_Migrating
Phenotype
Experiment
Control
PTCH1_KD
Control sim.: Polizio_21474452_Shh
Perturb. sim.: Polizio_21474452_PTCH1_null

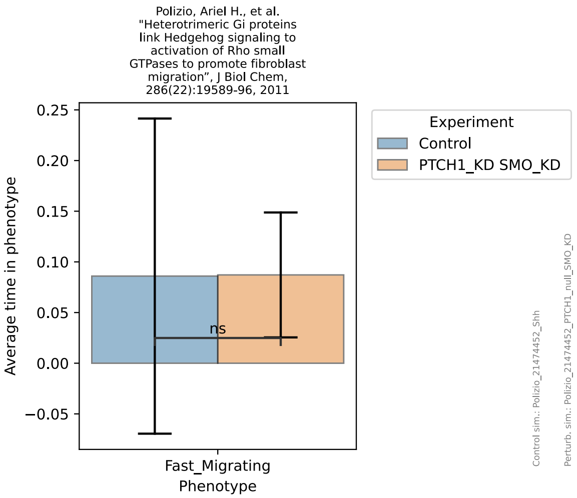
Polizio, Ariel H., et al. "Heterotrimeric Gi proteins link Hedgehog signaling to activation of Rho small GTPases to promote fibroblast migration", J Biol Chem, 286(22):19589-96, 2011
Average time in phenotype
0.25
0.20
0.15
0.10
0.05
0.00
−0.05
ns
Fast_Migrating
Phenotype
Experiment
Control
PTCH1_KD SMO_KD
Control sim.: Polizio_21474452_Shh
Perturb. sim.: Polizio_21474452_PTCH1_null_SMO_KD

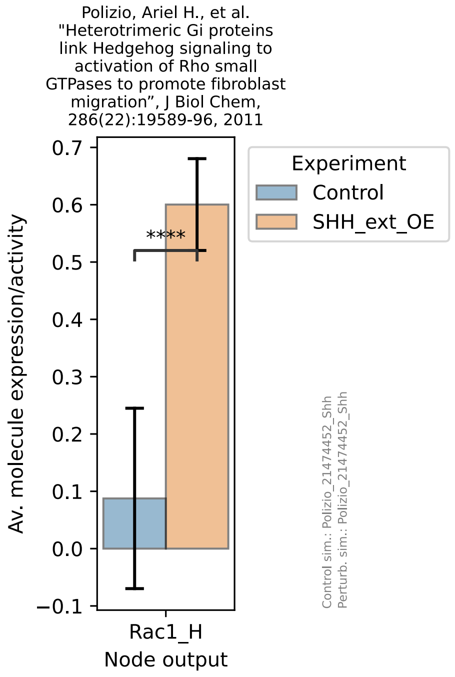
Polizio, Ariel H., et al. "Heterotrimeric Gi proteins link Hedgehog signaling to activation of Rho small GTPases to promote fibroblast migration", J Biol Chem, 286(22):19589-96, 2011
Av. molecule expression/activity
0.7
0.6
0.5
0.4
0.3
0.2
0.1
0.0
−0.1
****
Rac1_H
Node output
Experiment
Control
SHH_ext_OE
Control sim.: Polizio_21474452_Shh
Perturb. sim.: Polizio_21474452_Shh

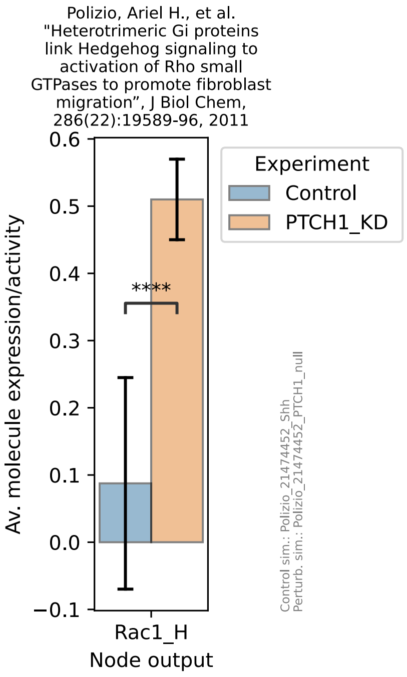
Polizio, Ariel H., et al. "Heterotrimeric Gi proteins link Hedgehog signaling to activation of Rho small GTPases to promote fibroblast migration", J Biol Chem, 286(22):19589-96, 2011
Av. molecule expression/activity
0.6
0.5
0.4
0.3
0.2
0.1
0.0
−0.1
****
Rac1_H
Node output
Experiment
Control
PTCH1_KD
Control sim.: Polizio_21474452_Shh
Perturb. sim.: Polizio_21474452_PTCH1_null

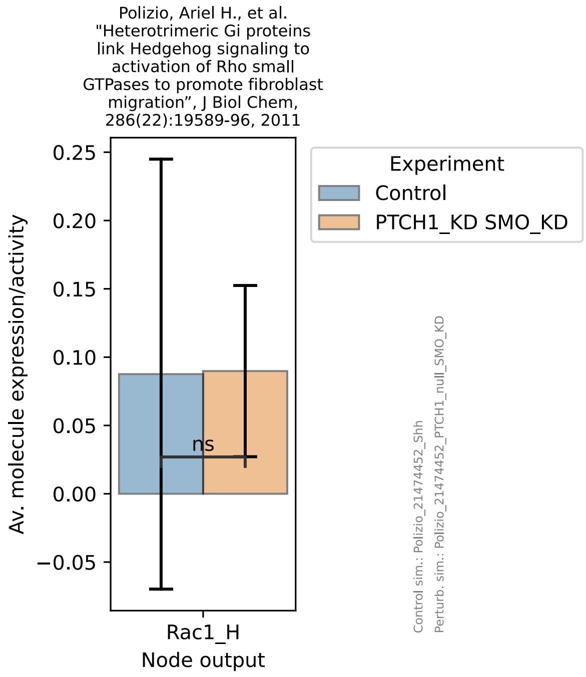
Polizio, Ariel H., et al. "Heterotrimeric Gi proteins link Hedgehog signaling to activation of Rho small GTPases to promote fibroblast migration", J Biol Chem, 286(22):19589-96, 2011
Av. molecule expression/activity
0.25
0.20
0.15
0.10
0.05
0.00
−0.05
ns
Rac1_H
Node output
Experiment
Control
PTCH1_KD SMO_KD
Control sim.: Polizio_21474452_Shh
Perturb. sim.: Polizio_21474452_PTCH1_null_SMO_KD

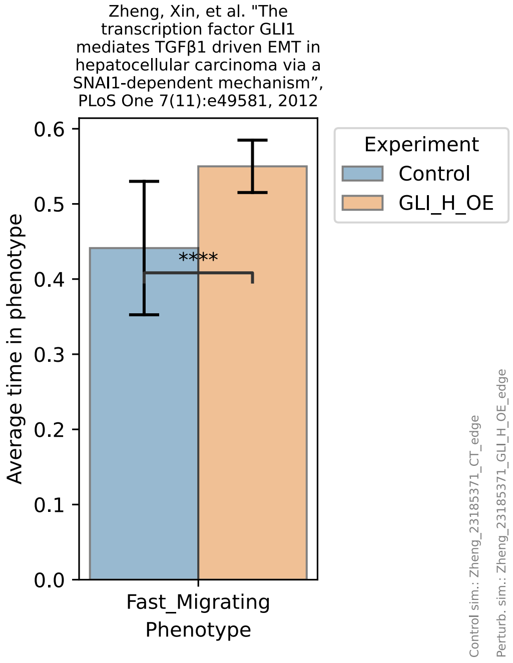
Zheng, Xin, et al. "The transcription factor GLI1 mediates TGFβ1 driven EMT in hepatocellular carcinoma via a SNAI1-dependent mechanism", PLoS One 7(11):e49581, 2012
Average time in phenotype
0.6
0.5
0.4
0.3
0.2
0.1
0.0
****
Fast_Migrating
Phenotype
Experiment
Control
GLI_H_OE
Control sim.: Zheng_23185371_CT_edge
Perturb. sim.: Zheng_23185371_GLI_H_OE_edge

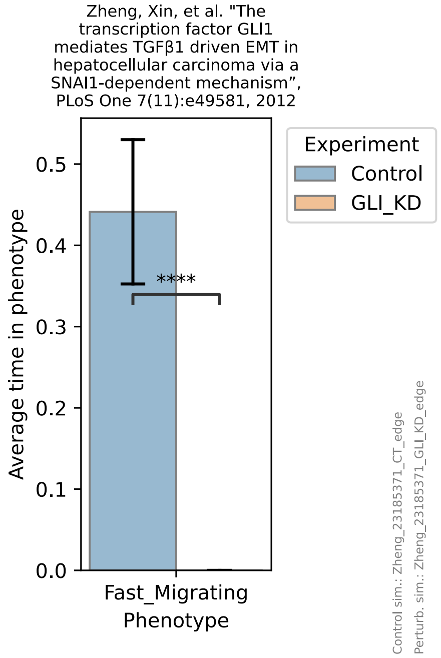
Zheng, Xin, et al. "The transcription factor GLI1 mediates TGFβ1 driven EMT in hepatocellular carcinoma via a SNAI1-dependent mechanism", PLoS One 7(11):e49581, 2012
Average time in phenotype
0.5
0.4
0.3
0.2
0.1
0.0
****
Fast_Migrating
Phenotype
Experiment
Control
GLI_KD
Control sim.: Zheng_23185371_CT_edge
Perturb. sim.: Zheng_23185371_GLI_KD_edge

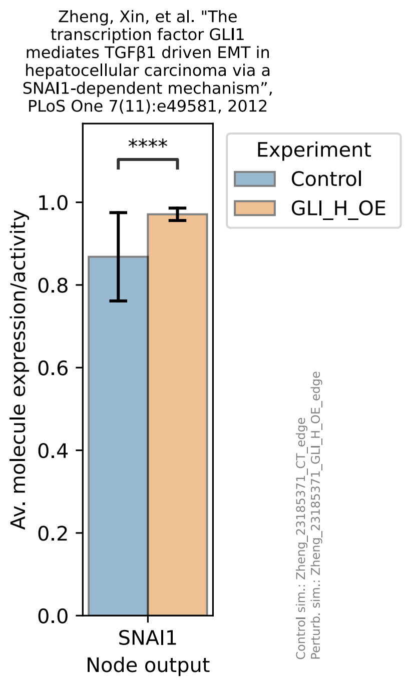
Zheng, Xin, et al. "The transcription factor GLI1 mediates TGFβ1 driven EMT in hepatocellular carcinoma via a SNAI1-dependent mechanism", PLoS One 7(11):e49581, 2012
****
Experiment
Control
GLI_H_OE
Av. molecule expression/activity
0.0
0.2
0.4
0.6
0.8
1.0
SNAI1
Node output
Control sim.: Zheng_23185371_CT_edge
Perturb. sim.: Zheng_23185371_GLI_H_OE_edge

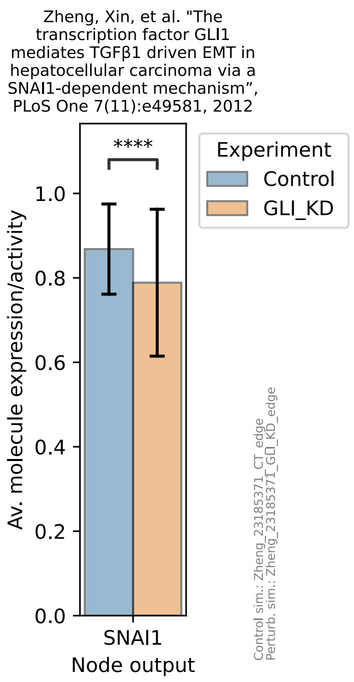
Zheng, Xin, et al. "The transcription factor GLI1 mediates TGFβ1 driven EMT in hepatocellular carcinoma via a SNAI1-dependent mechanism", PLoS One 7(11):e49581, 2012
****
Experiment
Control
GLI_KD
Av. molecule expression/activity
0.0
0.2
0.4
0.6
0.8
1.0
SNAI1
Node output
Control sim.: Zheng_23185371_CT_edge
Perturb. sim.: Zheng_23185371_GLI_KD_edge

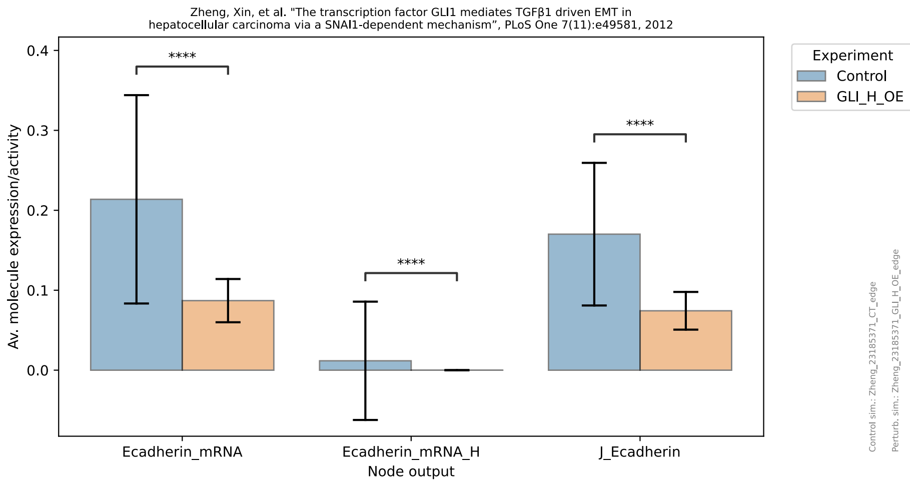
Zheng, Xin, et al. "The transcription factor GLI1 mediates TGFβ1 driven EMT in hepatocellular carcinoma via a SNAI1-dependent mechanism", PLoS One 7(11):e49581, 2012
Experiment
Control
GLI_H_OE
Av. molecule expression/activity
0.0
0.1
0.2
0.3
0.4
****
Ecadherin_mRNA
Ecadherin_mRNA_H
J_Ecadherin
Node output
Control sim.: Zheng_23185371_CT_edge
Perturb. sim.: Zheng_23185371_GLI_H_OE_edge

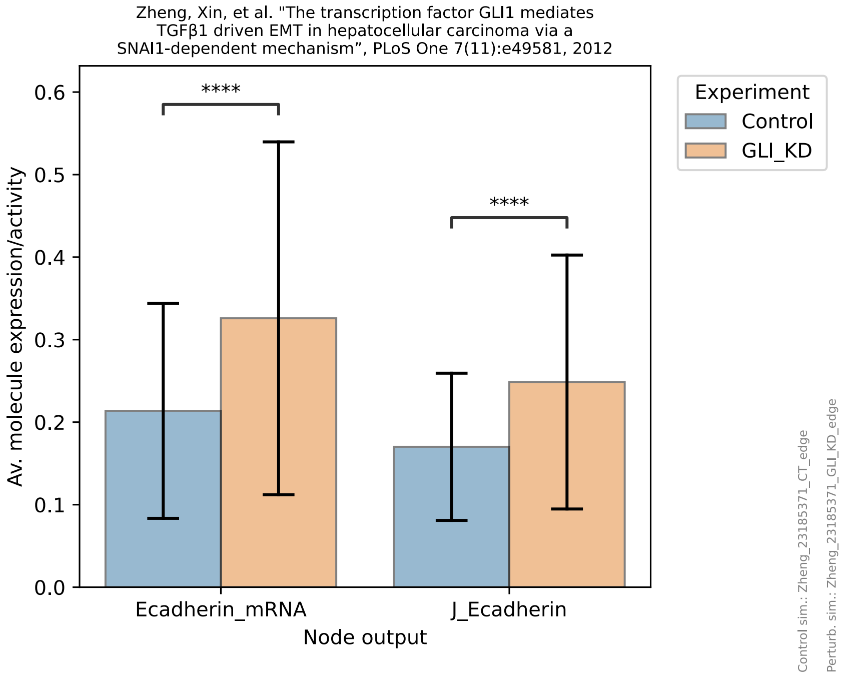
Zheng, Xin, et al. "The transcription factor GLI1 mediates TGFβ1 driven EMT in hepatocellular carcinoma via a SNAI1-dependent mechanism", PLoS One 7(11):e49581, 2012
Experiment
Control
GLI_KD
Av. molecule expression/activity
0.0
0.1
0.2
0.3
0.4
0.5
0.6
****
Ecadherin_mRNA
J_Ecadherin
Node output
Control sim.: Zheng_23185371_CT_edge
Perturb. sim.: Zheng_23185371_GLI_KD_edge

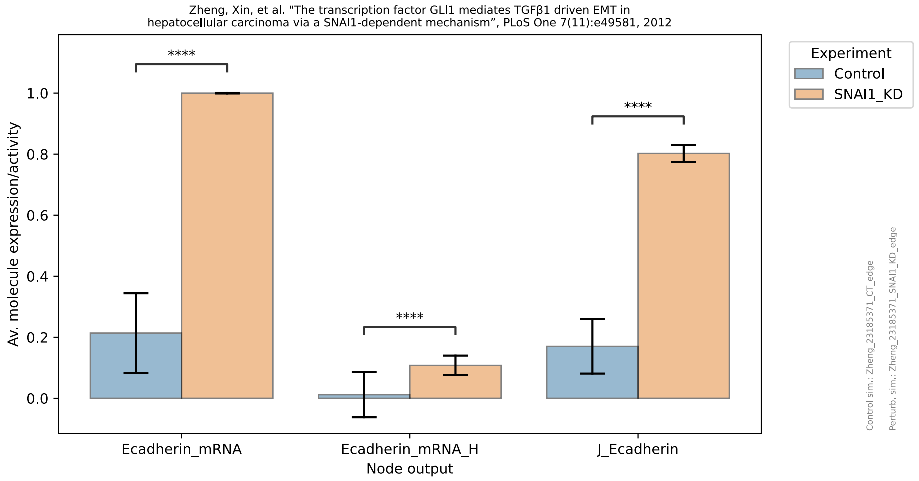

Zheng, Xin, et al. "The transcription factor GLI1 mediates TGFβ1 driven EMT in
hepatocellular carcinoma via a SNAI1-dependent mechanism", PLoS One 7(11):e49581, 2012
Experiment
Control
SNAI1_KD
Av. molecule expression/activity
1.0
0.8
0.6
0.4
0.2
0.0
****
****
****
Ecadherin_mRNA
Ecadherin_mRNA_H
J_Ecadherin
Node output
Control sim.: Zheng_23185371_CT_edge
Perturb. sim.: Zheng_23185371_SNAI1_KD_edge
