## Supplementary material for "HHIP’s Dynamic Role in Epithelial Wound Healing Reveals a Potential Mechanism of COPD Susceptibility": Supplementary Table 4 COPD_EMT_CellCycle_Apoptosis_SM.pdf

### S1 Description & experimental support for the modules of COPD_EMT_CellCycle_Apoptosis_Fine.

**Table S1a: Neighbors module**

| Target Node | Node Gate<br>Node Type<br>Link Type | Node Description<br>Input Node | Link Description |
|---|---|---|---|
| CellDensity _High | **CellDensity_High = CellDensity_High** | | |
| | Env | The *CellDensity_High* node in our model represents an extracellular environment with high enough cell density to block cell spreading. This input node is self-sustaining in the absence of *in silico* perturbation. | |
| | ← Env | CellDensity _High | *CellDensity_High* is self-sustaining in the absence of *in silico* perturbation. |
| CellDensity _Low | **CellDensity_Low = CellDensity_Low** or **CellDensity_High** | | |
| | Env | The *CellDensity_Low* node represents an environment with cell density comparable to the edge of a monolayer, where cells can maintain strong adhesions with each other but are also able to spread and polarize horizontally. | |
| | ← Env | CellDensity _High | *CellDensity_Low* is automatically ON at very high cell density. |
| | ← Env | CellDensity _Low | *CellDensity_Low* is self-sustaining in the absence of *in silico* perturbation. |

**Table S1b: Matrix module**

| Target Node | Node Gate<br>Node Type<br>Link Type | Node Description<br>Input Node | Link Description |
|---|---|---|---|
| COL1_3 | **COL1_3 = COL1_3** | | |
| | Secr | The interstitial ECM of alveoli is composed of a relaxed mesh, dominated by type I and III collagens and elastin [1]. Here we assume that these ECM components are secreted by tissue-resident fibroblasts into the connective tissue surrouding the alveolar basement membrane of alveolar type I and type II epithelial cells [1], and treat the *COL1_3* node as an environmental input. Complete absence of either the basement membrane (*Basement_Membrane* node) or *COL1_3* represents a scenario where the epithelial cells cannot form any ECM attachments (loss of anchorage; equivalent to the *ECM* = OFF and *Stiff_ECM* = OFF input in our [2] model). | |
| | ← Env | COL1_3 | *COL1_3* is self-sustaining in the absence of *in silico* perturbation. |

**Table S1b: Matrix module**

| | | | |
|---|---|---|---|
| Elastin | **Elastin** = not(**MMP7** or **MMP9**) | | |
| | Secr | In healthy lungs, the interstitial ECM, maitained by tissue-resident fibroblasts, is a loose meshwork of collagen and elastin fibers, cross-linked and connected to the basement membrane of the alveolar epithelial cell layer [1]. Elastin fibers are degraded by *MMP-7/9*, both *in vitro* and in lung tissue [3, 4, 5]. | |
| | $\dashv$ Deg | MMP7 | Elastin fibers are degraded by *MMP7*, both *in vitro* and in lung tissue [3, 4]. |
| | $\dashv$ Deg | MMP9 | Elastin fibers are cleaved by *MMP9*, both *in vitro* and in lung tissue from COPD and IPF patients [5]. |
| Laminin _5a | **Laminin_5a** = not **MMP2** | | |
| | Secr | Laminin $\alpha 5$ is a key component of the alveolar basement membrane in developing as well as adult lungs [6]. During EMT, laminins are cleaved by *MMP2* [7]. | |
| | $\dashv$ Deg | MMP2 | Cleavage of Laminin 5 by MMP2 was shown to aid epithelial cell migration [7]. |
| Collagen_4 | **Collagen_4** = not(**MMP9** or **MMP2**) | | |
| | Secr | Collagen IV is found primarily in the basal lamina, an exclusive member of basement membranes. Collagen IV fibers lack the tigh collagen helix structure of other collagens, and link to each other head-to-head (rather than in parallel). These two features allow the collagen IV molecules to form the sheet structure of basal lamina [8]. *MMP-2/9*, secreted by mesenchynal cells, can bind and degrade Collagen IV [9, 10]. | |
| | $\dashv$ Deg | MMP9 | *MMP9*, secreted by mesenchynal cells, can bind and degrade Collagen IV [9]. |
| | $\dashv$ Deg | MMP2 | *MMP2* binds and degrades Collagen IV [10]. |
| Basement _Membrane | **Basement_Membrane** = **Collagen_4** and **Laminin_5a** | | |
| | MSt | The alveolar basement membrane is a thin glycoproteins sheet that supports the basal side of the alveolar epithelium, separating it from the interstitial ECM and connective tissue. Its two critical components are Collagen IV and Laminin 5 [1]. | |
| | $\leftarrow$ Loc | Laminin _5a | Laminin 5 is a critical structural component of the basement membrane [1]. |
| | $\leftarrow$ Loc | Collagen_4 | Collagen IV is a critical structural component of the basement membrane [1]. |
| Fibronectin | **Fibronectin** = **SNAI1** and **NfkB** | | |

#### Table S1b: Matrix module

| | | | |
|---|---|---|---|
| | Secr | Basal levels of fibronectin serve to anchor the interstitial ECM to the basement membrane, and cross-link collagen and ellastin fibers. Increased fibronectin production and secretion, however, can stiffen the ECM and lead to fibrosis [1]. In our model, *Fibronectin* = OFF represnets basal levels, while the ON state reflects substantial epithelial cell-derived increase is fibronectin by *SNAI1*- and *NF-$\kappa$B*-induced transcription during partial of full EMT [11, 12]. | |
| | ← TR | SNAI1 | *SNAIL1* is a direct transcriptional inducer of *fibronectin* [11, 13]. |
| | ← TR | NfkB | *NF-$\kappa$B* is a direct transcriptional inducer of *fibronectin* [12]. |
| MMP2 | **MMP2** = ((not **CEBP_alpha**) and **SNAI1**) and (**ZEB2** or **HMGA1**) | | |
| | PTase | *MMP-2*, also known as gelatinase A, is a matrix metalloprotease that can degrade type IV collagen and thus compromise the basement membrane [9]. *MMP-2* is transcriptionally upregulated by *SNAI1* [14] and *HMGA1* [15], though *ZEB2* knockdown can significnatly reduce its levels as well [16]. In contrast, *C/EBP$\alpha$* can interfere with *MMP-2* mRNA induction [17]. | |
| | ⊢ TR | CEBP _alpha | *C/EBP$\alpha$* expression can interfere with TGF-$\beta$-mediated *MMP-2* induction [17]. |
| | ← TR | SNAI1 | *SNAI1* is a direct transcriptional inducer of *MMP-2*, binding to E-box element essential for its effect [18]. |
| | ← Ind | ZEB2 | *MMP-2* mRNA expression is significantly downregulated following *ZEB2* downregulation [16]. |
| | ← TR | HMGA1 | *MMP2* is a direct transcriptional target of *HMGA1* [15]. |
| MMP7 | **MMP7** = **LEF1** and **b_catenin_TCF4** and **ERK** | | |
| | PTase | *MMP-7* (matrilysin), is a potent elastase in humans, expressed by the mucosal epithelium and macrophages in the lung. It is upregulated in airway epithelial cells in response to injury [19], and it degrades ECM components such as elastin [20]. Elevated *MMP-7* levels are associated with impared lung function, and indeed its expression is increased in the blood of COPD patients [20]. *MMP-7* transcription is increased by *ERK* as well as *beta-catenin/TCF4/LEF1*[21, 22, 23]. | |
| | ← TR | LEF1 | *$\beta$-catenin/TCF-4/LEF1* complexes are direct transcriptional activators of *MMP7*[22]. |
| | ⊢ TR | b_catenin _TCF4 | *$\beta$-catenin/TCF-4* complexes are direct transcriptional activators of *MMP7* [23]. |
| | ← TR | ERK | *ERK1*, together with AP-1 (not explicitly modeled) are direct transcriptional activators of *MMP7* [21]. |
| MMP9 | **MMP9** = not **CEBP_alpha** and **SNAI1** and **SNAI2** and **ZEB2** | | |

**Table S1b: Matrix module**

| | | | |
|---|---|---|---|
| | PTase | | *MMP-9*, or Gelatinase B is a matrix metalloprotease with increased expression in COPD [20]. Among its targets are type IV collagen and elastin [20]. It's expression during EMT is induced by *SNAI1/2*, *ZEB2* and blocked by CEB/P-*alpha*. |
| | ← TR | SNAI1 | *SNAI1* is a direct transcriptional inducer of *MMP-9*; chromatin immunoprecipitation (ChIP) shows binding of *SNAI1* (together with EGR1 and SP1, not modeled) to the *MMP-9* promoter [24]. Moreover, *SNAI1* siRNA blocks *TGF-β1*-induced*MMP-9* expression [25]. |
| | ← TR | SNAI2 | *SNAI2* (*Slug*) increases *MMP-9* at both the protein and mRNA levels. Moreover, *Slug* siRNA decreases *TGF-β1*-induced *MMP-9* expression [26]. |
| | ← TR | ZEB2 | *MMP-9* mRNA expression is significantly upregulated by *ZEB2* overexpression, while siRNA against *ZEB2* strongly downregulates *MMP-9* expression as well as activity [27]. |
| | ⊣ Ind | CEBP _alpha | *C/EBP-α* can induce upregulation of *MMP-9* [28]. |
| ECM | **ECM** = (**COL1_3** and **Elastin**) or **Stiff_ECM** | | |
| | Env | | The *ECM* node represents access to at least a soft extracellular matrix that supports anchorage-dependent survival signaling, though it does not, by itself, support cell spreading or stress fiber formation ($< 0.5$ kPa) [29]. In out COPD model, this node requires the *COL1_3* input node along with *Elastin*, or it is overriden to an ON state by the *Stiff_ECM* node (meaning there is ECM). |
| | ← ComplProc | COL1_3 | The interstitial ECM of alveoli is composed of a relaxed mesh, dominated by type I and III collagens and elastin [1]. |
| | ← ComplProc | Elastin | The interstitial ECM of alveoli is composed of a relaxed mesh, dominated by type I and III collagens and elastin [1]. |
| | ← Per | Stiff_ECM | *ECM* node is automatically ON when cells have access to stiff *ECM*. |
| Stiff_ECM | **Stiff_ECM** = **Basement_Membrane** or (**COL1_3** and (**Fibronectin** or not **Elastin**)) | | |
| | Env | | The *Sfiff_ECM* node represents access to a very stiff extracellular matrix that promotes / supports stress fiber formation sufficiently to place no limitation on a cell's capacity to proliferate ( $> 100$ kPa) [29]. For alveolar epithelial cells, the basememnt membrane can serve this purpose, but so can a stiffened intersticial ECM outside of the basement membrane, containing excess fibronectin or lacking elastin. |
| | ← ComplProc | Basement _Membrane | The elastic or Young's modulus of adult BMs is in the 1-4 MPa range across a range of tissues, or ~1000 times greater than that of the epithelial cell layer (1-4 kPa) [30] or bulk lung tissue (0.44-7.5 kPa) [31]. |

**Table S1b: Matrix module**

| | | |
|---|---|---|
| ← ComplProc | COL1_3 | In the absence of a BM, we assume that our model cell experiences a stiff ECM if in contact with the interstitial ECM (mesh dominated by type I and III collagens) [1]. Thus, we require *COL1_3* = ON for all extracellular conditions with ECM contact. |
| ← ComplProc | Elastin | In the absence of a BM, we assume that our model cell experiences a stiff interstitial ECM, altered from its soft healthy mesh state by the loss of elastin [32]. |
| ← ComplProc | Fibronectin | Accumulation of excess fibronectin increases the density and fibrillar architecture of the ECM, increasing its stiffness [33, 34]. |

**Table S1c: Transforming_env module**

| Target Node | Node Gate<br>Node Type<br>Link Type | <br>Node Description<br>Input Node | <br><br>Link Description |
|---|---|---|---|
| TGFb_ext | **TGFb_ext = TGFb_ext** | | |
| | Env | The *TGFb_ext* node in our model represents an extracellular environment with saturating levels of *TGF-β*. This input node is self-sustaining in the absence of *in silico* perturbation. | |
| | ← Env | TGFb_ext | The *TGFb_ext* input node is self-sustaining in the absence of *in silico* perturbation. |
| SHH_ext | **SHH_ext = SHH_ext** | | |
| | Env | The *SHH_ext* node in our model represents an extracellular environment with saturating levels of *SHH*. This input node is self-sustaining in the absence of *in silico* perturbation. | |
| | ← Env | SHH_ext | The *SHH_ext* input node is self-sustaining in the absence of *in silico* perturbation. |
| Self_Loop | **Self_Loop = Self_Loop** | | |
| | Env | The *Self_Loop* input node qualtifies the extent to which *TGF-β* secreted by a single isolated cell can drive its own autocrine signaling to saturating levels. When tuned stochastically between 0 and 1, its level represents the fraction of saturating *TGF-β* signal driven by the autocrine loop. We expect the fraction that accurately characterises a given cell to be a funciton of both cell type (intrinsic ability to secrete *TGF-β*) and microenvironment, as the latter can influence diffusion and bio-availability of the secreted ligand. | |
| | ← Env | Self_Loop | The *Self_Loop* input node is self-sustaining in the absence of *in silico* perturbation. |

**Table S1d: TGF_beta module**

| Target Node | Node Gate<br>Node Type<br>Link Type | Node Description<br>Input Node | Link Description |
|---|---|---|---|
| TGF_bRII | **TGF_bRII** = **TGFb_ext** or (**TGFb_secr** and (**CellDensity_Low** or **Self_Loop**)) | | |
| | Rec | The ON state of the *TGF_bRII* node represents a *TGF-β*-bound *TβRII* receptor, and thus requires external *TGF-β* [35], or secreted *TGF-β* along with either strong autocrine signaling (*Self_Loop*=ON) or at least medium cell density (*CellDensity_Low*=ON). | |
| | ← ComplProc | CellDensity_Low | Though direct experimental evidence is hard to find, our model makes the assumption that autocrine signaling among cells that have at least medium density is strong enough to support autocrine *TGF-β* signaling, even if a single isolated cell's secretion is not sufficient. The assumption is indirecty supported by evidence that autocrine signaling is required for, and can indeed support the maintenance of a mesencymal state even after external *TGF-β* is no longer supplied [36, 37]. |
| | ← Ligand | TGFb_ext | Externally supplied *TGF-β* binds to and activates *TβRII* (*TGFBR2* gene) [35]. |
| | ← ComplProc | Self_Loop | In cells / environments where a single cell can secrete sufficient *TGF-β* to saturate its own *TGF-β* signaling, this input removes the requirement of neighboring cells to boost the availability of secreted *TGF-β*. |
| | ← Ligand | TGFb_secr | Mesenhymal cells secrete *TGF-β*, creating an autocrine signaling loop required to maintian their mesenchymal state [36, 37]. |
| TGF_bRI | **TGF_bRI** = **TGF_bRII** and (**TGFb_ext** or (**TGFb_secr** and (**CellDensity_Low** or **Self_Loop**))) | | |
| | Rec | The ON state of *TβRI* represents an active, ligand-bound receptor complex of *TGF-β*, *TβRII*, and *TβRI* [35]. Thus it requires *TGF-β*-bound *TβRII*, where *TGF-β* is either externally supplied or secreted *TGF-β* (the latter requiring strong autocrine signaling or medium cell density). | |
| | ← ComplProc | CellDensity_Low | Our model assumes that autocrine signaling among cells that have at least medium density is strong enough to support autocrine *TGF-β* signaling, even if a single isolated cell's secretion is not [36, 37]. |
| | ← Ligand | TGFb_ext | Externally supplied *TGF-β* binds to and activates *TβRII*, which in turn recruits and phosphorylates *TβRI* [35]. |
| | ← ComplProc | Self_Loop | In cells / environments where a single cell can secrete sufficient *TGF-β* to saturate its own *TGF-β* signaling, this input removes the requirement of neighboring cells to boost the availability of secreted *TGF-β*. |
| | ← P | TGF_bRII | When *TGF-β* binds to its type II receptor (*TβRII*), it recruits the type I receptor *TβRI* and activates it by phosphorylation [35]. |

**Table S1d: TGF_beta module**

| | | | |
|---|---|---|---|
| | ← Ligand | TGFb_secr | Mesenhymal cells secrete *TGF-β*, creating an autocrine signaling loop required to maintian their mesenchymal state [36, 37]. |
| SMAD2_3_4 | **SMAD2_3_4** = **TGF_bRI** and **TGF_bRII** and not **SPRY2** and ((**YAP** and **TAZ** and not **Rassf1a**) or not **ApicalBasal_Pol**) | | |
| | PC | | The *SMAD2_3_4* node represents transcriptionally active *Smad2/Smad3/Smad4* complexes. *Smad2* and *Smad3* are activated via phosphorylation by *TβRI*, which releases them from the receptor complex to aid their nuclear translocation in partnership with *Smad4* [35]. *SPRY2* reduces *Smad* phosphorylation in dense monolayers [38], which also sequester *TGF-β* receptors to their baso-lateral surface, hiding them from apically applied *TGF-β* [39]. In contrast, in the absence of *Rassf1a* *YAP* and *TAZ* bind to *Smads*, aid their nuclear localization, and potentiate *Smad*-induced transcription [40, 41]. Our model assumes that in order to respond to *TGF-β*, cells either need active *YAP/TAZ*, or no apical-basal polarizaiton. |
| | ⊣ DP | SPRY2 | *SPRY2* supresses *Smad2* phosphorylation and can block *TGF-β* induced EMT [38]. |
| | ← Loc | YAP | *YAP/TAZ* bind to *Smad2* and aid their nuclear accumulation and *Smad*-induced transcription. Fibroblast growth on soft gels with cytosolic *YAP/TAZ* or treated with chemical *YAP/TAZ* inhibitors show impaired *TGF-β*–induced *Smad2/3*-driven transcription [40]. |
| | ← Loc | TAZ | *YAP/TAZ* bind to *Smads* and aid their nuclear accumulation and *Smad*-induced transcription [40]. |
| | ⊣ Ind | Rassf1a | *Rassf1a* degradation is required for *YAP1* association with SMADs and the nuclear translocation of active *SMAD2* [41]. |
| | ⊣ Ind | ApicalBasal_Pol | Epithelial cell polarization was shown to block *TGF-β* signaling upstream and independently of cytoplasmic *YAP/TAZ* sequestration. In cells polarized along their apical-basal axis (represented in our model by *ApicalBasal_Pol* = ON), *TGF-β* receptors I and II are sequestered to the basolateral surface of the cell, depriving apically delivered *TGF-β* of access to its receptors and thus weakening the *TGF-β* response [39]. |
| | ← P | TGF_bRII | The actions of *TβRI* on *Smad2* and *Smad3* require an active receptor/ligand complex, and thus *TGF_bRII* [35]. |
| | ← P | TGF_bRI | The type I *TGF-β* receptor (*TβRI*) phosphorylates receptor-bound *Smad2* and *Smad3* at their carboxy-terminal, which releases them from the receptor complex and triggers their nuclear translocation. *Smad4* partners with activated *Smads* to help carry out their function [35]. |
| ILK_Rictor | **ILK_Rictor** = (**ILK** and **TGF_bRI**) and **TGF_bRII** | | |
| | PC | | *TGFβ-1* induces expression of the *mTORC2* component *Rictor*, *ILK* binding to *Rictor*, and *ILK*-dependent *Rictor* phosphorylation [42]. The complex, in turn, is known to activate *AKT1* [43]. |

**Table S1d: TGF_beta module**

| | | | |
|---|---|---|---|
| | ← Compl | ILK | *TGF$\beta$-1* induces *ILK* binding and phosphorylation of *Rictor* [42]; we assume this requres an active *ILK* kinase. |
| | ← Ind | TGF_bRII | *ILK/Rictor* binding is mediated by *TGF$\beta$* receptors I and II, responsive to *TGF$\beta$-1* [42]. |
| | ← Ind | TGF_bRI | *ILK/Rictor* binding is mediated by *TGF$\beta$* receptors I and II, responsive to *TGF$\beta$-1* [42]. |
| HMGA2 | **HMGA2** = **SMAD2_3_4** | | |
| | TF | | *HMGA2* is a transcription factor induced by *SMAD 3/4* in response to *TGF-$\beta$*. Once induced, it drives transcription of the master EMT switch by inducing *SNAI1/2* and *Twist* [44]. |
| | ← TR | SMAD2_3_4 | *HMGA2* is a direct transcriptonal target of *TGF-$\beta$* induced *SMAD 3/4* [44]. |
| p15 | **p15** = **SMAD2_3_4** or ((**FoxO1** or **FoxO3**) and not **Myc**) | | |
| | CDKI | | *p15* (*Ink4B*) is a cyclin-dependent kinase inhibitor that binds *cdk4* / *cdk6*, displacing *Cyclin D* and thus blocking G1/S progression [45]. *TGF$\beta$* induces both transcription [46] and stabilization of *p15* [47], while *Myc* represses its transcription [48]. In addition, *FoxO1/3* can also induce *p15* [49]. |
| | ← TR | FoxO3 | *FoxO3* is a direct transctiptional activator of *p15* [49]. |
| | ← TR | FoxO1 | *FoxO1* is a direct transctiptional activator of *p15* [49]. |
| | ← TR | SMAD2_3_4 | *Smad2*, *Smad3* and *Smad4* induce *p15* transcription and increase its protein stability in response to *TGF$\beta$* [46]. |
| | ⊢ TR | Myc | *Myc* is a direct transctiptional repressor of *p15* [48]. |
| DUSP4 | **DUSP4** = **SMAD2_3_4** | | |
| | Ph | | *TGF$\beta$* induces the transcription of the mitogen-activated protein kinase phosphatase *MKP2*, or *DUSP4*, through a *SMAD3*-dependent mechanism [50]. *DUSP4*, in turn, attenuates *ERK* and allows *BIM* accumulation, aiding *TGF$\beta$*-mediated apoptosis [50, 51]. |
| | ← TR | SMAD2_3_4 | *TGF$\beta$* induces *DUSP4* transcription through *SMAD3* [50]. |
| Runx1 | **Runx1** = **SMAD2_3_4** | | |
| | TF | | The *Runx1* transcription factor is transcriptionally induced by *TGF$\beta$*, leading to elevated mRNA and protein levels [52]. *Runx1* in turns binds to *FoxO3* to induce *BIM* and promote *TGF$\beta$*-mediated apoptosis [52]. |
| | ← TR | SMAD2_3_4 | Either *Smad2* or *Smad3*, plus *Smad4*, are required for induction of *Runx* transcription factors by *TGF$\beta$*1 [53]. |

**Table S1e: Hedgehog module**

| Target Node | Node Gate<br>Node Type<br>Link Type | Node Description<br>Input Node | Link Description |
|---|---|---|---|
| SHH_secr | **SHH_secr** = ((**GLI** and **NfkB**) or **SMAD2_3_4** or (**GLI_H** and not **GLI3R**)) and not **Casp3** | | |
| | Secr | *Shh* secretion by mesenchymal cell can aid a self-sustaining autocrine signal linked to *TGF-β* signaling. Namely, *Shh* signaling itself can boost *Shh* secretion via transcriptional activation by *GLI2* (*GLI* node; antagonized by the *GLI3R* repressor) [54, 55, 56], potentially aided by *NF-κB* [57]. In addition, *TGF-β* can stimulate *Shh* transcription (presumably via *SMAD2/3/4*) [58]. Finally, apoptotic *Caspase 3* activation blocks *Shh* secretion by dismantling vesicular transport between the ER and Golgi [59]. | |
| | ←<br>TR | GLI | *GLI2* is a direct transcriptional inducer of the *FoxA2* transcriptipn factor [54], which in turn induces *Shh* mRNa expression in several tissue contexts, including lung branching [55]. Moreover, *FoxA2* and *GLI2* bind to the same regulatory region in the *Shh* promoter, suggesting that they may work together to induce *Shh* [56]. |
| | ←<br>TR | GLI_H | *GLI2* is a direct transcriptional inducer of the *FoxA2* transcriptipn factor [54], which in turn induces *Shh* mRNa expression in several tissue contexts, including lung branching [55]. Moreover, *FoxA2* and *GLI2* bind to the same regulatory region in the *Shh* promoter, suggesting that they may work together to induce *Shh* [56]. |
| | ⊢<br>TR | GLI3R | Activator and repressor forms of the GLI transcription factors share the same DNA binding motifs. Thus, we assume that *GLI2* (the GLI node here) competes with *GLI3R* for access to the GLI binding site on their target genes. Here we assume that *Shh* transcription, secretion, and positive autocrine feedback requires this balance to facor the transcriptional activator *GLI2* [60]. |
| | ←<br>TR | NfkB | *NF-κB* is a direct transcriptional inducer of the *Shh* gene [61]. Inhibition of of*NF-κB* can suppress constitutive expression of *Shh* mRNA, whereas its stimulation by inflammatory signals boosts *Shh* production and secretion [57]. |
| | ←<br>Ind | SMAD2_3_4 | *TGF-β* stimulation induces *Shh* mRNA and protein expression [58]; here we assume that the mechanism involves *SMAD2/3/4* complexes. |
| | ⊢<br>Ind | Casp3 | During apopotsis, caspases cleave key components vesicular transport between the ER and Golgi, resulting in a complete block in ER-mediated protein secretion [59]. |
| PTCH1 | **PTCH1** = not(**SHH_ext**or(**SHH_secr**and(not **HHIP**or(**GLI_H**andnot **GLI3R**))and (**CellDensity_Low** or **Self_Loop**))) | | |

**Table S1e: Hedgehog module**

| | | | |
|---|---|---|---|
| | Rec | In the absence of Hh ligand binding, the *PTCH1* receptor represses the activity of Smoothened (*SMO*). When Hh ligands such as *Shh* bind to *PTCH1*, the receptor is trafficked to the lysosome for degradation [62]. Here we assume that exogenous, saturating levels of *Shh* (*SHH_ext* node) can repress *PTCH1*, while secreted *Shh* (*SHH_secr* node) can only fully repress it in the absence of *HHIP* or under high *Shh* expression conditions (balance between *GLI* activators and repressors strongly favors the former). In addition, we require either the assumption that cells secrete sufficient *Shh* to fully activate the hedgehog pathway (modeled by *Self_Loop* = ON), or that the cell is not isolated and may also receive *Shh* signals from its neighborhood. | |
| | ⊢ BLoc | SHH_ext | Hh ligands such as *Shh* bind to *PTCH1*, inducing its internalization and lysosomal degradation [62]. Here we assume that exogenous *Shh* can saturate the pathway. |
| | ⊢ BLoc | SHH_secr | Hh ligands such as *Shh* bind to *PTCH1*, inducing its internalization and lysosomal degradation [62]. Here we assume that secreted *Shh* can only saturate the pathway (block *PTCH1*) if not antagonized by *HHIP*, or produced at a higher rate by *GLI* [54, 55, 56]. |
| | ⊢ Ind | GLI_H | Here we assume that maximal *GLI2*-mediated transcription and secretion of *Shh* can saturate the pathway by fully blocking *PTCH1* [54, 55, 56]. |
| | ← Ind | GLI3R | Here we assume that the absence of the *GLI3R* repressor is required for *GLI2*-mediated transcription and secretion of sufficient *Shh* to saturate the autocrine signaling pathway [54, 55, 56, 60]. |
| | ← BLoc | HHIP | *HHIP* is a membrane glycoprotein that binds to all three mammalian Hedgehog proteins with an affinity comparable to that of *PTCH1* [63], and its loss upregulates *Hh* signaling during early lung branching [64]. |
| | ⊢ ComplProc | Self_Loop | In cells / environments where a single cell can secrete sufficient *SHH* to saturate its own *SHH* signaling (even in the dampening presence of *HHIP*), this input removes the requirement of neighboring cells to boost the availability of secreted *SHH*. |
| | ⊢ ComplProc | CellDensity_Low | Our model makes the assumption that autocrine signaling among cells that have at least medium density is strong enough to support autocrine *SHH* signaling, even if a single isolated cell's secretion is not sufficient. This matches our handling of *SHH* autocrine signaling, where the above assumption is indirecty supported by evidence that such autocrine signaling is required for, and can indeed support the maintenance of a mesencymal state even after external *SHH* is no longer supplied [36, 37]. |
| SMO | **SMO** = (not **PTCH1**) and (**SHH_secr** or **SHH_ext**) | | |

**Table S1e: Hedgehog module**

| | | | |
|---|---|---|---|
| | Rec | In the absence of *PTCH1*, the transmembrane signaling protein *SMO* is phosphorylated. This activates *SMO*, promotes its translocation into the primary cilia where they block phosphorylation and proteolytic cleavage of GLI2/3 (by cAMP-dependent *PKA* and *CK1*, not modeled) [62]. We assume that external *Shh* is required to keep *PTCH1* from blocking *SMO*. | |
| | ⊢ BLoc | PTCH1 | In the absence of *PTCH1*, *SMO* is phosphorylated and activated by *CK1α* and *GRK2* (kinases not modeled explicitly) [62]. |
| | ⊢ BLoc | SHH_ext | We assume that external or secreted *Shh* is required to keep *PTCH1* from blocking *SMO*. |
| | ⊢ BLoc | SHH_secr | We assume that external or secreted *Shh* is required to keep *PTCH1* from blocking *SMO*. |
| SUFU | **SUFU** = not **SMO** | | |
| | Prot | Suppressor of fused, or *SUFU*, is a major *GLI* regulator. It binds and sequesters these factorst to the primary cilia, where it aids their phosphorylation by *PKA*, *CK1*, and *GSK3β*, marking them for proteolytic cleavage [62]. | |
| | ← BLoc | SMO | Once activated and localized to prolary cilia, *SMO* inhibits the sequestration of *GLI* transcription factors by *SUFU*, sparing them from proteolytic cleavage [62]. |
| GLI | **GLI** = (not **SUFU** or not **CEBP_alpha** or **SMAD2_3_4** or **S6K** or **Myc**) and **PAK1** | | |
| | TF | *GLI1/2* are the main activator transcription factors that mediate *Hh* signaling. They are stabilized and released to the nucleus in the absence of *SUFU* [62] – overriden by phosphorylation by *S6K1* [65] –, and are induced in a *Hh*-independent manner by the loss of *C/EBPα* [66, 67] or transcription by *SMAD2/3/4* [68, 69]. Finally, *PAK1* activation helps relieve *SUFU*-mediated sequestration of *GLI1/2*, allowing them to accumulate in an active form [70]. | |
| | ⊢ BLoc | SUFU | *SUFU* binds and sequesters these factorst to the primary cilia, where it aids their phosphorylation by *PKA*, *CK1*, and *GSK3β*, marking them for proteolytic cleavage [62]. |
| | ⊢ Ind | CEBP_alpha | Loss of *C/EBPα* increases *GLI1*, including in alveolar cells [66, 67]. |
| | ← TR | S6K | *S6K1*-mediated *GLI1* phosphorylation at Ser84 releases *GLI1* from its endogenous inhibitor, *SUFU* [65]. |
| | ← TR | SMAD2_3_4 | *TGF-β* induces *GLI2* via *SMAD2/3/4*-mediated transcription [68, 69]. |
| | ← TR | Myc | *c-Myc* is a direct transcriptional inducer of *GLI1* [71]. |
| | ← Ind | PAK1 | *Rac1*-dependent active *PAK1* dampens *SUFU-GLI* complex formation and thus aids *GLI* activation [70]. |
| GLI_H | **GLI_H** = **GLI** and (not **SUFU** or **SMAD2_3_4**) and (not **CEBP_alpha** or **SHH_ext** or (**SHH_secr** and not **HHIP**)) | | |

**Table S1e: Hedgehog module**

| | | | |
|---|---|---|---|
| | TF | | *GLI1/2* are the main activator transcription factors that mediate *Hh* signaling. Here we model their activity levels with two Boolean nodes: *GLI* = ON and *GLI_H* = OFF represents moderate activation, whereas *GLI_H* = ON represents high levels of *GLI*-mediated transcription. For model consistency, this requires *GLI* = ON. In additon, we assume that saturating *GLI_H* activity is aided by ongoing *Shh* signaling (i.e., absence of *SUFU*, presence of a strong *Shh* signal), as well increased transcription by *SMAD2/3/4* [68, 69] or loss of *C/EBPα* [66, 67]. |
| | ← Per | GLI | *GLI_H* = ON requires *GLI* = ON, making sure that the two nodes together represent three distinct levels of *GLI* activity. |
| | ⊣ BLoc | SUFU | *SUFU* binds and sequesters these factorst to the primary cilia, where it aids their phosphorylation by *PKA*, *CK1*, and *GSK3β*, marking them for proteolytic cleavage [62]. |
| | ⊣ Ind | CEBP_alpha | Loss of *C/EBPα* increases *GLI1*, including in alveolar cells [66, 67]. |
| | ← TR | SMAD2_3_4 | *TGF-β* induces *GLI2* via *SMAD2/3/4*-mediated transcription [68, 69]. |
| | ← Ind | SHH_ext | We assume that saturating *GLI_H* activity is aided by ongoing expgenous *Shh* signaling. |
| | ← Ind | SHH_secr | We assume that saturating *GLI_H* activity is aided by ongoing secreted *Shh*, unrestrained by *HHIP* [63]. |
| | ⊣ Ind | HHIP | We assume that saturating *GLI_H* activity is aided by ongoing secreted *Shh*, unrestrained by *HHIP* [63]. |
| GLI3R | **GLI3R** = not(**GLI_H** or (**GLI** and **GSK3**)) | | |
| | TF | | The *GLI3* gene encodes a strong repressor, the proteolytically processed active form of which is termed *GLI3R*. This truncated protein antagonizes HH signalling, and it is dominant in the absence of HH ligands. Upon *SMO* activation, the activators *GLI1/2* are transcribed, while *GLI3R* formation is repressed [72]. Thus, we assume that *GLI3R* is inactive in the presence of high *GLI1/2* [60], or moderate *GLI* paired with a *GSK3* = ON cell state [73]. |
| | ⊣ TR | GLI_H | As activator and repressor forms of the GLI share the same DNA binding motifs, we assume that *GLI_H* = ON represents a state where the activators robustly outcompete the repressor forms [60]. |
| | ⊣ TR | GLI | We assume that moderate *GLI1/2* can outcompete *GLI3R* only with the aid of *Myc*-mediated repression of *GLI3*. |
| | ⊣ P | GSK3 | The Ser-9 phosphorylated, inactivated form of *GSK3β* (corresponding to *GSK3* = OFF in our model) can nevertheless phosphorylate the unprocessed form of *GLI3*, aiding its conversion into the repressor form *GLI3* [73]. |
| HHIP | **HHIP** = **GLI** or **GLI_H** or **SMAD2_3_4** | | |

**Table S1e: Hedgehog module**

| Target Node | Node Gate / Node Type / Link Type | Node Description / Input Node | Link Description |
|---|---|---|---|
| | Prot | *HHIP* is a membrane glycoprotein that binds to all three mammalian Hedgehog proteins with an affinity comparable to that of *PTCH1* [63], and its loss upregulates *Hh* signaling [64]. It is transcriptionally induced by *GLI* proteins as part of a homeostatic negative feedback loop, as well as by *TGF-β* via *SMAD2/3/4*. | |
| | ← TR | GLI | *HHIP* is a *GLI*-activated, Hh induced gene [74]; *GLI* binds to an enhancer region of the *HHIP* gene [75, 76]. |
| | ← TR | GLI_H | *HHIP* is a *GLI*-activated, Hh induced gene [74]; *GLI* binds to an enhancer region of the *HHIP* gene [75, 76]. |
| | ← TR | SMAD2_3_4 | *SMAD3* binds to a distant enhancer of the *HHIP* gene and upregulates it in response to *TGF-β* [77]. |
| CEBP_alpha | **CEBP_alpha** = not(**SMAD2_3_4** or **CyclinE** or (**CyclinA** and **Cdc25A**)) | | |
| | TF | *C/EBPα* is a transcriptipn factor that is required for the maturation of the respiratory epithelium in late gestation. It's loss can alter surfactant secretion (not modeled), and altered the biomechanical environment of the lung [67]. | |
| | ⊣ TR | SMAD2_3_4 | The *CEBPA* gene is a direct target of *SMAD3* represssion during *TGF-β*-mediated EMT [78]. |
| | ⊣ P | CyclinE | *Cdk2* (the activity of which is imlicitly modeled as *CyclinE* = ON or *CyclinA* and *Cdc25A* = ON) negatively regulates *C/EBPα* protein levels by ubiquitin-mediated proteasome degradation [79]. |
| | ⊣ Ind | Cdc25A | *Cdk2* (the activity of which is imlicitly modeled as *CyclinE* = ON or *CyclinA* and *Cdc25A* = ON) negatively regulates *C/EBPα* protein levels by ubiquitin-mediated proteasome degradation [79]. |
| | ⊣ P | CyclinA | *Cdk2* (the activity of which is imlicitly modeled as *CyclinE* = ON or *CyclinA* and *Cdc25A* = ON) negatively regulates *C/EBPα* protein levels by ubiquitin-mediated proteasome degradation [79]. |

**Table S1f: GrowthFactor_Env module**

| Target Node | Node Gate / Node Type / Link Type | Node Description / Input Node | Link Description |
|---|---|---|---|
| GF | **GF** = **GF** or **GF_High** | | |
| | Env | The *GF* node represents an extracellular environment with low levels of growth factors capable of sustaining survival signaling. Thus, the *GF* input node is self-sustaining in the absence of in silico perturbation. | |
| | ← Env | GF | The *GF* input node is self-sustaining in the absence of in silico perturbation. |

**Table S1f: GrowthFactor_Env module**

| | | | |
|---|---|---|---|
| | ← Env | GF_High | The *GF* node represents an extracellular environment with low levels of growth factors capable of sustaining survival signaling. Thus, this node is ON in high growth factor as well. |
| GF_High | **GF_High** = **GF_High** | | |
| | Env | The $GF_{High}$ node in our model represents an extracellular environment with saturating levels of growth factors; this input node is self-sustaining in the absence of *in silico* perturbation. | |
| | ← Env | GF_High | The $GF_{High}$ input node is self-sustaining in the absence of *in silico* perturbation. |

**Table S1g: GF_Basal_MAPK module**

| Target Node | Node Gate<br>Node Type<br>Link Type | Node Description<br>Input Node | Link Description |
|---|---|---|---|
| RTK | **RTK** = (not **CAD**) and (**GF_High** or **GF**) | | |
| | Rec | The ON state of the *RTK* node in our model represents basal growth receptor activation (required to keep a normal cell alive). Thus it requires the absence of *CAD* and at least low growth levels of growth factors in the extracellular environment [80]. | |
| | ← Ligand | GF | The ON state of the *RTK* node in our model represents basal growth receptor activation by low / medium growth factor availability, encoded by the *GF* node (required to keep a normal cell alive). |
| | ← Ligand | GF_High | Similarly, high growth factor availability also keeps *RTK* on. |
| | ⊢ Per | CAD | Caspase-activated DNase (*CAD*) inhibition of receptor tyrosine kinases ensures that apoptotic cells no longer maintain even basal levels of growth signaling. |
| Shc | **Shc** = ((**RTK** and **GF_High**) or (**TGF_bRI** and **TGF_bRII**)) and (**FAK** or **Src**) | | |
| | Adap | *Shc* is ON when *RTK*s are activated by high levels of extracellular growth factors (capable of driving proliferation) [81] or $TGF\beta$ activates $T\beta RI$-$T\beta RII$ [82], and its recruitment is aided by active *Src* kinase [83] or focal adhesion kinase (*FAK*) [83]. The two mediators of integrin signaling appears to be able to act independently, forming two parallel links between integrin signaling and full *RTK* activation [83]. | |
| | ← Compl | GF_High | The ON state of *Shc* in our model encode the change from basal *Shc* recruitment to weakly stimulated *RTKs* to the level of recruitment seen in high growth factor environments, capable of mediating *Ras* activation. |
| | ← Compl | RTK | *Shc* proteins are adaptors that binds to phosphor-tyrosine motifs, facilitating their recruitment to activated receptors such as receptor tyrosine kinases *RTKs* [81]. |

**Table S1g: GF_Basal_MAPK module**

| | | | |
|---|---|---|---|
| | ← Compl | FAK | *FAK* activated at sites of integrin-ECM attachments directly phosphorylates *Shc Tyr-317*, promoting its ability to assemble MAPK-inducing signaling scaffolds, including *Grb2* binding [83]. |
| | ← Compl | Src | *c-Src* recruited to and activated by integrin-ECM attachments directly phosphorylate *Shc*, promoting its ability to assemble MAPK-inducing signaling scaffolds, including *Grb2* binding [83]. |
| | ← Compl | TGF_bRII | *TGFβ* receptor II (*TβRII*) can be phosphorylated by *Src* on Y284, which provides a docking site for *Grb2* and *Shc*, and subsequent MAPK activation [82]. |
| | ← Compl | TGF_bRI | *TβRII* activation requires complex formation with *TβRI* upon ligand binding [82]. |
| Grb2 | **Grb2** = (**RTK** or (**TGF_bRI** and **TGF_bRII**)) and **Shc** | | |
| | Adap | | *Grb2* is recruited to *RTKs* or *TGFβ* receptors upon ligand binding and subsequent recruitment of *Shc* adaptors [84, 82]. |
| | ← Compl | RTK | The SH2 domain of *Grb2* binds to a phosphotyrosine residue in the activated *RTK*, where it functions as an adaptor protein [84]. |
| | ← Compl | Shc | *Shc* proteins phosphorylated by tyrosine kinases represent binding sites for *Grb2*, aiding its recruitment to active RTKs [81]. |
| | ← Compl | TGF_bRII | *TβRII* phosphorylated by *Src* on Y284, provides a docking site for *Grb2* and *Shc* [82]. |
| | ← Compl | TGF_bRI | *TβRII* works in complex with *TβRI* and their ligand [82]. |
| SOS | **SOS** = **Grb2** | | |
| | GEF | | *RTK*-bound *Grb2* recruits *SOS*, a guanine nucleotide–exchange protein (GEF) that converts inactive *Ras* to its active GTP-bound form [84]. |
| | ← Compl | Grb2 | *Grb2* recruits *SOS* to activated *RTK*s [84]. |
| Ras | **Ras** = ((**Grb2** and **SOS**) and **Src**) and ((**IQGAP1_LeadingE** or (not **Merlin**)) or **N_bcatenin_H**) | | |
| | GTPa | | *Ras* activation requires the GEF activity of *SOS* and the *RTK*-linked (active) adaptor protein *Grb2* [84] and *Src* [85, 86]. In addition to aiding sustained *Ras/Raf-1* signaling, *Src* may also physically link *IQGAP1* to *RTKs* such as *VEGFR2* [87]. *IQGAP1*, in turn, serves a scaffold for *MAPK* and *PI3K* signaling [88], leading us to link its activation at the leading edge. In contrast, *Merlin* blocks *Ras* activation at sites that link focal adhesions and actin filaments to *MAPK* signaling [89]. Here we assume that concentrated *IQGAP1* at the leading edge can override remaining *Merlin* activity in the rest of the cell. Alternatively, high levels of *β-catenin* can also sustain *Ras* by protecting it from lysosomal degradation [90]. |

**Table S1g: GF_Basal_MAPK module**

| | | | |
|---|---|---|---|
| | ← Compl | Grb2 | *RTK*-bound *Grb2* is required to recruits *SOS*, the GEF responsible for converting inactive *Ras* to its GTP-bound active form [84]. |
| | ← GEF | SOS | *SOS* is a GEF that is recruited to activate *Ras* near ligand-bound, active *RTKs* [84]. |
| | ← ComplProc | Src | Cellular Src (*c-Src*) is required for mitogenesis initiated by multiple growth factor receptors, including epidermal growth factor (*EGF*), platelet-derived growth factor (*PDGF*), colony stimulating factor-1 (*CSF-1*), and basic fibroblast growth factor (*bFGF*) [85]. In addition to aiding the formation of Shc/Grb2/SOS/Ras/Raf-1 cascade, *Src* may also increase the rate of receptor internalization and aid sustained *MAPK* signaling by internalized *Ras* on endosomes and Golgi [85, 86]. |
| | ← ComplProc | Merlin | *Merlin* uncouples *Ras* from growth factor signals by counteracting the ERM (ezrin, radixin, moesin)–dependent activation of *Ras*, which aids *Grb2*, *SOS*, *Ras* complex formation linked to filamentous actin [89]. |
| | ← Compl | IQGAP1 _LeadingE | *IQGAP1* acts as a scaffold for the *MAPK* cascade, binding directly to *B-Raf*, *MEK*, and *ERK* and regulating their activation [91]. The *IQGAP1-Leading-E* node in our model specifically links the availability of active *IQGAP1* recruited to lamellipodia and enhanced *MAPK* / *AKT* signaling. |
| | ← Ind | N_bcatenin _H | High levels of *β-catenin* protect *Ras* from lysosomal degradation [90]. Overall, *β-catenin* overexpression / silencing can activate / block *ERK* in a *MEK*-dependent way, like via *Ras/Raf* [92]. |
| RAF | **RAF** = ((not **Casp3**) and **Ras**) and (not **SPRY2**) | | |
| | K | | *Raf* is active in response to *Ras* activity in the absence of *Caspase 3*. As active *Raf-1* is continuously dephosphorylated and bound by *14-3-3*, which translocates it to the cytoplasm from the plasma membrane (not modeled explicitly), ongoing *Ras* activity [93] and lack of *SPRY2* inhibition [94, 95] are necessary to keep *Raf* ON. |
| | ← P | Ras | Active *Ras* phosphorylates *Raf*, enhancing its kinase activity [93]. |
| | ⊢ IBind | SPRY2 | Sprouty2 (*SPRY2*) blocks Raf activity and downstream MAPK signaling [94, 95]. |
| | ⊢ Lysis | Casp3 | *Raf-1* is cleaved and inhibited by *Caspase 3* [96]. |
| MEK | **MEK** = **RAF** | | |
| | K | | *Raf* phosphorylates and activates the *MEK* kinase [93]. |
| | ← P | RAF | *Raf* phosphorylates and activates the *MEK* kinase [93]. |
| ERK | **ERK** = (**MEK** and (not **BIK**)) and ((**FocalAdhesions** or (not **DUSP4**)) or **N_bcatenin_H**) | | |

**Table S1g: GF_Basal_MAPK module**

| | | | |
|---|---|---|---|
| | K | The *ERK* kinase is active when phoisphotylated by *MEK* [93] and allowed to translocate to the nucleus in the absence of *BIK* [97]. Here we assume that in the presence of *DUSP4* [50, 51], *ERK* activity can be maintained /compensated by either strong focal adhesion assembly (*FocalAdhesions* node) [98], or high levels of *β-catenin* [92]. | |
| | ← P | MEK | *MEK* phosphorylates and activates the *ERK* kinase [93]. |
| | ← ComplProc | FocalAdhesions | Focal adhesions recruit the MAPK scaffolding protein GIT1 and locally potentiate *ERK1/2* activation [98]. |
| | ⊢ DP | DUSP4 | *DUSP4* is a phosphatase that blocks *ERK* and allows *BIM* accumulation in response to *TGFβ*, aiding apoptosis [50, 51]. |
| | ← Ind | N_bcatenin _H | *β-catenin* overexpression / silencing can activate / block *ERK* in a *MEK*-dependent way [92]. |
| | ⊢ IBind | BIK | *BIK* binds to active, phosphorylated *ERK1/2* and suppresses its nuclear translocation [97]. |
| MCRIP1 | **MCRIP1** = not **ERK** | | |
| | Prot | *MCRIP1* is a binding partner and repressor of *CtBP*, a binding partner of *Zeb1* during transcription , and thus *MCRIP1* blocks *Zeb1*-mediated repression of *E-cadherin* unless inhibited by *ERK* phosphorylation [99]. | |
| | ⊢ P | ERK | Upon *ERK* phosphorylation *MCRIP1*, an inhibitor of EMT, dissociates and releases the *ZEB1* co-activator *CtBP* [99]. |
| mTORC2 | **mTORC2** = **PIP3** or (not **S6K**) | | |
| | PC | Our model assumes that *mTORC2* is active in quiescent cells with basal levels of *PI3K* activity leading to basal *PIP3* generation. Alternatively, the absence of high growth factor-stimulated *mTORC1* and *S6K1* can also increase *mTORC2* activity. | |
| | ← PBind | PIP3 | PtdIns(3,4,5)P3 (*PIP3*), interacts with the *mTORC2* component *Sin1* to release its inhibition on the *mTOR* kinase domain. Thus, *PIP3* is necessary for *mTORC2* activation [100]. |
| | ⊢ P | S6K | *Rictor*, a component of the *mTORC2 complex*, undergoes *S6K1*-mediated phosphorylation at T1135, dampening *mTORC2*-dependent phosphorylation of *Akt* [101, 102]. |
| PI3K | **PI3K** = ((**FAK** or **Src** or **GLI**) and (**Ras** or **RTK**)) or (**TGF_bRI** and **TGF_bRII**) | | |
| | K | In our model, basal *PI3K* activity can be maintained by active *RTK*s [80], active *Ras* [103, 104], or active *TβRI-TβRII* [105]. In addition, survival signalling via *PI3K* requires anchorage-dependent signals via active *FAK* [106], via *Src*-mediated blocking of basal *PTEN* activity (not modeled explicitly) [107], or a *GLI1*-mediated boost to *PI3K* activity [108]. | |
| | ← BLoc | RTK | Active *RTK*s recruit *PI3K* to the signaling complex they nucleate, where *PI3K* catalyzes the production of PtdIns(3,4,5)P3 (PIP3) [80]. |

| | | | |
|---|---|---|---|
| | ← Compl | Ras | *Ras* binds the catalytic subunit of *PI3K* and *Ras* knockdown / over expression decreases /increases the *PI3K*-dependent generation of PIP3 [103, 104]. |
| | ← PLoc | FAK | Attachment to the ECM activates *FAK* kinase, which promotes anchorage-dependent survival signaling via *PI3K* / *AKT* [106]. |
| | ← P | Src | *Src* kinases regulate *PI3K* signaling cascade by altering the function of the *PTEN* tumor suppressor via inhibitory phosphorylation [107]. |
| | ← Compl | TGF_bRII | *TGFβ* induces *PI3K* activation and *AKT* phosphorylatio, a pathway required for EMT [109]. This activation requires *TβRI* and *TβRII* kinase activity [105]. |
| | ← Compl | TGF_bRI | *TGFβ* induced *PI3K* activation requires *TβRI* and *TβRII* kinase activity [109, 105]. |
| | ← Compl | GLI | Overexpression of *GLI1* increases the phosphorylation of *PI3K* and *AKT*, while stable knockdown of *GLI1* attenuates it [108]. |
| PIP3 | **PIP3** = **PI3K_H** or **PI3K** | | |
| | Met | In our model, PIP3 is ON as a result of basal or high *PI3K* activity. | |
| | ← Cat | PI3K | Active *PI3K* recruited to the membrane catalyzes the production of membrane-bound PtdIns(3,4,5)P3 (PIP3) from PtdIns(4,5)P2 (PIP2) [80]. |
| | ← Cat | PI3K_H | Active *PI3K* recruited to the membrane catalyzes the production of membrane-bound PtdIns(3,4,5)P3 (PIP3) from PtdIns(4,5)P2 (PIP2) [80]. |
| PDK1 | **PDK1** = **PI3K** and **PIP3** | | |
| | K | *PDK1* enzyme activation requires active (at least basal) *PI3K* and *PIP3* [110]. | |
| | ← BLoc | PI3K | The *PDK1* kinase is recruited to the plasma membrane by *PIP3* at the sites of active *PI3K* activity [110]. |
| | ← BLoc | PIP3 | The *PDK1* kinase is recruited to the plasma membrane by *PIP3* at the sites of active *PI3K* activity [110]. |
| AKT_B | **AKT_B** = ((not **Casp3**) and **PIP3**) and (**PDK1** or **mTORC2**) | | |
| | K | Basal *AKT1* activity in our model requires the absence of *Caspase 3*, the availability of at least basal levels of *PIP3*, and phosphorylation by *PDK1* or *mTORC2*. In contrast, full mitogen-stimulated *AKT1* activation requires phosphorylation by both (see *AKT_H*) [110]. | |
| | ← P | mTORC2 | Maximal activation of *AKT1* requires phosphorylation of S473 by *mTORC2* [110]. |
| | ← BLoc | PIP3 | *PIP3* recruits *AKT1* to the plasma membrane and *PIP3* binding changes the conformation of *AKT1* such that it becomes accessible for T308 phosphorylation by *PDK1* [110]. |

**Table S1g: GF_Basal_MAPK module**

| | | | |
|---|---|---|---|
| | ← P | PDK1 | Membrane-recruited *PDK1* phosphorylates *AKT1* at T308, a critical step in its activation [110]. |
| | ⊢ Lysis | Casp3 | *AKT1* is cleaved and inhibited by *Caspase 3* [96]. |

**Table S1h: GF_PI3K module**

| Target Node | Node Gate<br>Node Type<br>Link Type | <br>Node Description<br>Input Node | <br><br>Link Description |
|---|---|---|---|
| p110_H | **p110_H** = **YAP** and ((**FoxO3** and (not **Nedd4L**)) or (**p110_H** and (**FoxO3** or (not **Nedd4L**)))) | | |
| | Prot | As *YAP* is a transcriptional inducer of *p110* subunits [111] and their expression is low in high cell density areas where *YAP* activity is suppressed [112], here we assume that high *p110* expression requires active *YAP*. In order to capture the cyclic dynamics of *p110* protein expression, we make the assumption that high *p110* protein levels can be induced by *FoxO3* in the absence of the growth factor-activated *Nedd4L* ubiquitin ligase. Once present, high *p110* can be maintained by *FoxO3* transcription, or the absence of activated *Nedd4L*. | |
| | ← Per | p110_H | Our model assumes that maintaining high *p110* levels is easier than driving the re-accumulation of the protein following its rapid destruction. |
| | ← TR | FoxO3 | *FoxO3* is a direct inducer *p110α* (*PIK3CA*), the catalytic subunit of *PI3K* [113]. |
| | ⊢ Ubiq | Nedd4L | *p110α* (*PIK3CA*) is polyubiquitinated by the E3 ligase *Nedd4L*, leading to its proteasomal degradation. Both free *p110α* and the regulatory subunit-bound protein is subject to ubiquitination by *Nedd4L* [114]. |
| | ← TR | YAP | *YAP* is a transcriptional inducer of both catalytic *p110* subunits of *PI3K*, *p110a* and *p110b*; *p110-H* albeit its effect on *p110a* expression requires raising *p110b* levels first. Moreover, *YAP* knockdown leads to downregulation of both subunits [111]. |
| PI3K_H | **PI3K_H** = ((((not **PTEN_c**) and **p110_H**) and (**RTK** or (**TGF_bRI** and **TGF_bRII**))) and **PI3K**) and (**Ras** or **GLI_H**) | | |
| | K | Full, peak-level activation of *PI3K* requires high levels of *p110* protein, basal *PI3K* activation, active *Ras*, and active *RTK*s or *TβRI-TβRII*. As the ON-state of *Ras* in our model represents strong *Ras* activation in the presence of proliferation-inducing (high) growth factors, *PI3K_H* activation can only occur with either high *Ras* or a *GLI1*-mediated boost to *PI3K* activity [108]. In addition, a reduction of cytoplasmic *PTEN* levels is also required for peak *PI3K* activity. | |

**Table S1h: GF_PI3K module**

| | | | |
|---|---|---|---|
| | ← BLoc | RTK | High levels of *PI3K* activation only occur at growth factor-bound *RTK*s, which recruit and activate *PI3K* at the plasma membrane [110]. |
| | ← Compl | Ras | *Ras* binds the catalytic subunit of *PI3K* and *Ras* knockdown / over expression decreases /increases the *PI3K*-dependent generation of PIP3 [103, 104]. |
| | ← Per | PI3K | In our model, high *PI3K* activation is contingent on the ON-state of the basal *PI3K* node. |
| | ← Per | p110_H | High levels of *PI3K* activity in response to strong growth factor stimulation only occur in cells that express high levels of *p110* protein [112]. |
| | ⊣ DP | PTEN_c | Cytoplasmic *PTEN* regulates *PI3K* signaling by dephosphorylating its lipid signaling intermediate *PIP3* [115]. |
| | ← Compl | TGF_bRII | *TGFβ* induced *PI3K* activation requires *TβRI* and *TβRII* kinase activity [109, 105]. |
| | ← Compl | TGF_bRI | *TGFβ* induced *PI3K* activation requires *TβRI* and *TβRII* kinase activity [109, 105]. |
| | ← Compl | GLI_H | Overexpression of *GLI1* increases the phosphorylation of *PI3K* and *AKT*, while stable knockdown of *GLI1* attenuates it [108]. |
| AKT_H | **AKT_H** = (((((**AKT_B** and **p110_H**) and (**PI3K_H** or (**PI3K** and **ILK_Rictor**))) and **PIP3**) and **PDK1**) and (**mTORC2** or **ILK_Rictor**)) and (**Ras** or **PAK1** or **GLI_H**) | | |
| | K | | In contact to basal *AKT1*, high *AKT1* activity in our model requires basal *AKT1* (*AKT_B*), the ongoing presence of high *p110* protein levels along with active *PI3K_H* and *PIP3*. In addition this maximal *AKT1* activation requires phosphorylation by both *PDK1* and *mTORC2*, , as well as either active *Ras* [110], active *PAK1* [116] or a *GLI1*-mediated boost to *PI3K/AKT* activity [108]. In response to *TGFβ*, high *AKT1* activity can also be induced by *PI3K* and the *ILK-Rictor* complex; the latter capable of replacing *mTORC2* to supply S473 phosphorylation of *AKT1* [42, 43]. |
| | ← Compl | GLI_H | Overexpression of *GLI1* increases the phosphorylation of *PI3K* and *AKT*, while stable knockdown of *GLI1* attenuates it [108]. |
| | ← Compl | Ras | *Ras* binding to the catalytic subunit of *PI3K* is required for its full potency in *PIP3* generation [103, 104]. Active *Ras* is thus required for inducing peak *AKT_H* activity. |
| | ← P | mTORC2 | Maximal activation of *AKT1* requires phosphorylation of S473 by *mTORC2* [110]. |
| | ← P | PI3K | As maximal *AKT1* activity also requires T308 phosphprylation by *PDK1* recruited to the membrane by *PI3K* [110], we assume that *ILK/Rictor* complexes can only dirve high *AKT* activity in the presence of at least basal *PI3K* activation. |
| | ← Compl | PIP3 | *PIP3* recruits *AKT1* to the plasma membrane and *PIP3* binding changes the conformation of *AKT1* such that it becomes accessible for T308 phosphorylation by *PDK1* [110]. |

**Table S1h: GF_PI3K module**

| | | | |
|---|---|---|---|
| | $\leftarrow$ P | PDK1 | Membrane-recruited *PDK1* phosphorylates *AKT1* at T308, a critical step in its activation [110]. |
| | $\leftarrow$ Per | AKT_B | In our model, high *AKT1* activation is contingent on the ON-state of basal *AKT1* (*AKT_B*). |
| | $\leftarrow$ P | p110_H | Ongoing high *p110* availability and *PI3K_H* activity are required to induce maximal activation of *AKT_H* [110]. |
| | $\leftarrow$ P | PI3K_H | Ongoing high *p110* availability and *PI3K_H* activity are required to induce maximal activation of *AKT_H* [110]. |
| | $\leftarrow$ P | PAK1 | *PAK1* interacts with and directly phosphorylates *AKT1* [117]. In addition, *PAK1* provides a scaffold to facilitate *Akt* stimulation by *PDK1* and to aid *AKT's* membrane recruitment [116]. |
| | $\leftarrow$ P | ILK_Rictor | Maximal activation of *AKT1* requires phosphorylation of S473. In response to *TGF-β*, *ILK* complexes with *Rictor* to phosphoprylate *AKT1* [42, 43]. |
| FoxO3 | **FoxO3** = (not((**AKT_B** or **AKT_H**) or **ERK**)) or ((not(**AKT_H** and (((**Plk1** or **Plk1_H**) or **AKT_B**) or **ERK**))) and (not((**Plk1** and **Plk1_H**) and **ERK**))) | | |
| | TF | | In order to account for all the influences on *FoxO3* activity, we used the following logic. In the absence of basal or high *AKT1* as well as *ERK*, *FoxO3* remains active. In addition, *FoxO3* can overcame peak (*AKT_H*) activation only if no other inhibitor is present and *AKT_B* is OFF (indicating that *AKT1* levels are falling). Finally, the joint activity of *ERK* and *Plk1* can also block *FoxO3*. |
| | $\vdash$ P | ERK | ERK downregulates *FoxO3* transcriptional activity by phosphorylating it at three Serines, inducing its *MDM2*-mediated ubiquitination and degradation [118]. |
| | $\vdash$ PLoc | AKT_B | *AKT1* mediates the translocation of the *FoxO3* our of the nucleus through direct phosphorylation of three conserved residues. These events create a recognition site for *14-3-3* family proteins, which export and sequester *FoxO3* in the cytosol [110]. |
| | $\vdash$ PLoc | AKT_H | *AKT1* mediates the translocation of the *FoxO3* our of the nucleus through direct phosphorylation of three conserved residues. These events create a recognition site for *14-3-3* family proteins, which export and sequester *FoxO3* in the cytosol [110]. |
| | $\vdash$ PLoc | Plk1 | *Plk1* binds *FoxO3*, induces its translocation to the cytosol, phosphorylates it and suppresses its activity through most of the the cell cycle, but most significantly during G2 and M [119]. |
| | $\vdash$ PLoc | Plk1_H | *Plk1* binds *FoxO3*, induces its translocation to the cytosol, phosphorylates it and suppresses its activity through most of the the cell cycle, but most significantly during G2 and M [119]. |

**Table S1h: GF_PI3K module**

PLCgamma — **PLCgamma** = ((((**RTK** and **Grb2**) and **GF_High**) and **p110_H**) and **PI3K_H**) and **PIP3**

| | | |
|---|---|---|
| Enz | Peak activation of $PLC\gamma$ requires active an *RTK* receptor node bound by active *Grb2*, as well as high *PI3K* activity (including high *p110* availability and the presence of PIP3). | |
| ← Compl | GF_High | We assume that high levels of *RTK* activity is required for tyrosine phosphorylation of $PLC\gamma$. |
| ← P | RTK | The SH2 domains of $PLC\gamma$ binds to active *RTK*s at tyrosine autophosphorylation sites, leading to tyrosine phosphorylation of $PLC\gamma$ and stimulation its enzymatic activity [120, 121]. |
| ← BLoc | Grb2 | *RTK* tyrosine autophosphorylation induces $PLC\gamma$ binding to the *Grb2* adaptor protein and likely aids the translocation of $PLC\gamma$ to the plasma membrane [122]. |
| ← BLoc | PIP3 | Membrane targeting of $PLC\gamma$ to growth receptor stimulation is mediated by *PIP3* binding of $PLC\gamma$ [123, 124]. |
| ← P | p110_H | Membrane targeting of $PLC\gamma$ to growth receptor stimulation requires *PI3K* activity and *PIP3* generation near growth receptors [123]. Thus, peak $PLC\gamma$ activity in our model requires high *p110* protein expression [124]. |
| ← P | PI3K_H | In addition to high *p110* protein levels, high *PI3K* activation is also required to fully activate $PLC\gamma$ [123, 124]. |

IP3 — **IP3** = **PLCgamma**

| | | |
|---|---|---|
| Met | Membrane-bound, active $PLC\gamma$ is responsible for converting phosphatidylinositol(4,5)P2 (*PIP2*) to the second messenger inositol(1,4,5)P3 (*IP3*) responsible for triggering a sudden $Ca^{2+}$ influx from the endoplasmic reticulum, along with *DAG* (diacylglycerol, another second messenger) [125]. | |
| ← Cat | PLCgamma | Membrane-bound, active $PLC\gamma$ is responsible for converting phosphatidylinositol(4,5)P2 (*PIP2*) to the second messenger inositol(1,4,5)P3 (*IP3*) responsible for triggering a sudden $Ca^{2+}$ influx from the endoplasmic reticulum, along with *DAG* (diacylglycerol, another second messenger) [125]. |

Ca2p — **Ca2p** = **IP3**

| | | |
|---|---|---|
| Met | *IP3* travels from the cell membrane to the endoplasmic reticulum where it opens *IP3*-sensitive $Ca^{2+}$ channels, releasing a sudden $Ca^{2+}$ efflux from the ER into the cytosol [126]. | |
| ← Loc | IP3 | *IP3* travels from the cell membrane to the endoplasmic reticulum where it opens *IP3*-sensitive $Ca^{2+}$ channels, releasing a sudden $Ca^{2+}$ efflux from the ER into the cytosol [126]. |

Nedd4L — **Nedd4L** = **Ca2p** and **IP3**

| | | |
|---|---|---|
| UbL | Activation of *Nedd4L* requires both $Ca^{2+}$ and *IP3* binding [127]. | |
| ← Compl | IP3 | In order to transition to its active form, the E3 ubiquitin ligase *Nedd4L* binds $Ca^{2+}$ and inositol 1,4,5-trisphosphate (*IP3*) [127]. |

**Table S1h: GF_PI3K module**

| | | | |
|---|---|---|---|
| | ← Compl | Ca2p | In order to transition to its active form, the E3 ubiquitin ligase *Nedd4L* binds $Ca^{2+}$ and inositol 1,4,5-trisphosphate (*IP3*) [127]. |

**Table S1i: GF_mTOR module**

| Target Node | Node Gate<br>Node Type<br>Link Type | <br>Node Description<br>Input Node | <br><br>Link Description |
|---|---|---|---|
| TSC2 | **TSC2** = (not **AKT_H**) or (not(**AKT_B** or **ERK**)) | | |
| | Prot | Blocking *TSC2* requires ongoing mitogen stimulation through *AKT* and/or *ERK*. In our model, *TSC2* inhibition requires high (peak) *AKT* activity, supported by either *ERK* or basal *AKT* (assuring that complete loss of *AKT* activity is not impending) [128]. | |
| | ⊢ P | ERK | *ERK* phosphorylates *TSC2* directly, causing dissociation of the complex and inhibition of its activity [129]. In addition, the *ERK* target *p90RSK* can also inactivate *TSC2* [130]. |
| | ⊢ P | AKT_B | *TSC2* is phosphorylated by *AKT1*, inhibiting it by dissociating *TSC2* from lysosomal membranes [131], where it stimulates GTP hydrolysis of the small GTPase *Rheb*, this inactivating it [128]. |
| | ⊢ P | AKT_H | *TSC2* is phosphorylated by *AKT1*, inhibiting it by dissociating *TSC2* from lysosomal membranes [131], where it stimulates GTP hydrolysis of the small GTPase *Rheb*, this inactivating it [128]. |
| PRAS40 | **PRAS40** = (not **AKT_H**) and ((not **mTORC1**) or (not **AKT_B**)) | | |
| | Prot | *PRAS40* is inhibited by peak *AKT1* activity aided by either basal *AKT1* (meaning *AKT_H* is on its way down), or ongoing *mTORC1* activation. Both *AKT1* and *mTORC1* phosphorylate *PRAS40*, leading to its dissociation from *mTORC1* [132]. | |
| | ⊢ P | AKT_B | *PRAS40* is phosphorylated by *AKT*, triggering its dissociation from *mTORC1* [133]. |
| | ⊢ P | AKT_H | *PRAS40* is an inhibitory component of the *mTORC1* complex. It is phosphorylated by *AKT*, triggering its dissociation from *mTORC1* and loss of *mTORC1* inhibition [133]. |
| | ⊢ P | mTORC1 | *PRAS40* is a substrate of the *mTORC1* kinase; its phosphorylation aids its dissociation from *mTORC1* and its sequestration by 14-3-3 proteins [134]. |
| DAG | **DAG** = **PLCgamma** | | |
| | Met | Membrane-bound, active *PLC*$\gamma$ is responsible for converting phosphatidylinositol(4,5)P2 (*PIP2*) to the second messenger diacylglycerol (*DAG*), along with *IP3* [125]. | |

**Table S1i: GF_mTOR module**

| Node | Type | Regulator | Description |
|---|---|---|---|
| | ← Cat | PLCgamma | Membrane-bound, active *PLC*$\gamma$ is responsible for converting phosphatidylinositol(4,5)P2 (*PIP2*) to the second messenger diacylglycerol (*DAG*), along with *IP3* [125]. |
| Rheb | **Rheb** = (not **TSC2**) and **DAG** | | |
| | GTPa | | *PKC* (and *DAG*)-dependent activation of *mTORC1* recruits *mTORC1* to the site of *Rheb* activity (to prenuclear lysosomes), while *AKT* and *ERK*-mediated *TSC2* inhibition guarantees that *Rheb* remains potent [135, 136, 137]. |
| | ⊢ GAP | TSC2 | *TSC2*, a key component of the heterotrimeric *TSC* complex, is a GTPase activating protein (GAP) that induces ATP hydrolysis and deactivation of the small GTPase *Rheb* [135]. |
| | ← Ind | DAG | The second messenger *DAG* activates both classical and novel *PKC*s. One of its targets, *PKC*$\eta$, is responsible for the translocation and accumulation of *mTORC1* to perinuclear lysosomes, where the majority of *Rheb* is anchored. Thus, *DAG* brings *Rheb* in proximity with its target, *mTORC1* [136]. |
| mTORC1 | **mTORC1** = not **Casp3** and (not **PRAS40** or **ERK**) and (((**Rheb** and not **Merlin**) or **E2F1**) or (**CyclinB** and **Cdk1** and **GSK3**)) | | |
| | PC | | *mTORC1* is activated by mitogenic signals via *Rheb* in the absence of *PRAS40*, counteracted by active *ERK* [138]. This, however, also requires inactivation of *Merlin* independently of *RTK*-induced signals. In addition, *E2F1* can promote *mTORC1* activity. Finally, the mitotic *Cyclin B* and its *Cdk1* kinase can also activate *mTORC1*, aided by *GSK3*. |
| | ⊢ IBind | PRAS40 | *PRAS40* is an inhibitory component of the *mTORC1* complex, removed by phosphorylation by *AKT* or *mTORC1* itself [128]. |
| | ← P | ERK | *ERK1/2* phosphorylate the *Raptor* component of mTORC1 to activate it downstream of *Ras* [138]. |
| | ← Compl | Rheb | The *Rheb* small GTPase binds *mTORC1* directly and activates the complex [139]. |
| | ⊢ ComplProc | Merlin | *Merlin* suppresses *mTORC1* activity via an unknown mechanism that appears to be independent of *PI3K/AKT* or of *TSC2* inhibition [140], and it suppression appears to be critical for integrin-mediated *mTORC1* activation [141]. |
| | ← Loc | E2F1 | *E2F1* induces *mTORC1* activity by inducing *mTORC1* translocation to late endosomes. This effect does not require *AKT* and is not blocked by high levels of *TSC2* [142]. |
| | ← Ind | CyclinB | During mitosis, *mTORC1* is activated by the G2/M-specific phosphorylation of *Raptor*, a component of *mTORC1*, by *CyclinB / Cdk1* complexes, aided by *GSK3* [143]. |
| | ← Ind | Cdk1 | During mitosis, *mTORC1* is activated by the G2/M-specific phosphorylation of *Raptor*, a component of *mTORC1*, by *CyclinB / Cdk1* complexes, aided by *GSK3* [143]. |
| | ← Ind | GSK3 | During mitosis, *mTORC1* is activated by the G2/M-specific phosphorylation of *Raptor*, a component of *mTORC1*, by *CyclinB / Cdk1* complexes, aided by *GSK3* [143]. |

**Table S1i: GF_mTOR module**

| | | | |
|---|---|---|---|
| | ⊢ Lysis | Casp3 | *Raptor*, a key component of the *mTORC1* complex, is cleaved and inhibited by *Caspase 3* [144]. |
| S6K | **S6K** = (not **Casp3**) and **mTORC1** | | |
| | K | *S6K* is activated by *mTORC1* in the absence of *Caspase 3*. | |
| | ← P | mTORC1 | *mTORC1* phosphorylates and activates 40S ribosomal S6 kinases (*S6K*s) [145]. |
| | ⊢ Lysis | Casp3 | *S6K* is cleaved and inhibited by *Caspase 3* [146]. |
| h_4EBP1 | **h_4EBP1** = (not **mTORC1**) or **SMAD2_3_4** | | |
| | Prot | *4EBP1*, or eukaryotic translation initiation factor 4E-binding protein 1, binds and sequesters the initiation factor *eIF4F* and thus limits translation. *mTORC1* phosphorylates and deactivates *4EBP* to upregulate protein biosynthesis [128]. While in our previous models this protein was implicit along the *mTORC1* → *eIF4F* link [147, 148], we include it here because *TGFβ* signaling induces *4EBP1* via *SMAD4* to block proliferative metabolism [149]. | |
| | ⊢ P | mTORC1 | *mTORC1* phosphorylates *4EBP*, triggering its dissociation from *eIF4F* [128]. |
| | ⊢ TR | SMAD2_3_4 | *4EBP1* is a transcriptional target of *SMAD4*, and its increase is critical for *TGF-β* mediated inhibition of proliferation [149]. |
| eIF4E | **eIF4E** = (not **Casp3**) and (not **h_4EBP1**) | | |
| | Prot | *eIF4E* is activated by *mTORC1*-mediated repression of *4EBP1*, which normally binds to it and blocks its activity [128]. *eIF4E* is cleaved and deactivated by *Caspase 3* [150]. | |
| | ⊢ IBind | h_4EBP1 | *mTORC1*-phosphorylated *4EBP* dissociates from *eIF4F*, allowing it to initiate translation [128]. |
| | ⊢ Lysis | Casp3 | *eIF4E* is cleaved and inhibited by *Caspase 3* [150]. |

**Table S1j: GF_connect module**

| Target Node | Node Gate<br>Node Type<br>Link Type | <br>Node Description<br>Input Node | <br><br>Link Description |
|---|---|---|---|
| GSK3 | **GSK3** = (not **AKT_H**) and (not(**S6K** and **ERK**)) | | |
| | K | *GSK3* activity can be completely blocked by peak *AKT* activation ( *AKT_H*), or by the joint action of *S6K* and *ERK*. | |
| | ⊢ P | ERK | *ERK* binds and phosphorylates *GSK3β* at Thr-43, which primes it for subsequent phosphorylation by the *ERK* target *p90RSK* at Ser-9, which inactivates *GSK3β* [151]. |

**Table S1j: GF_connect module**

| | | | |
|---|---|---|---|
| | ⊣ P | AKT_H | *AKT* blocks *GSK3* kinase activity via an inhibitory phosphorylation on the amino terminus, which blocks the substrate accessibility of *GSK3* [110]. |
| | ⊣ P | S6K | *GSK3* is a direct phosphorylation target of *S6K1*, resulting in its inhibition [101]. |
| FoxO1 | **FoxO1** = (not **Plk1**) and (not **AKT_H**) | | |
| | TF | | *FoxO1* is transcriptionally active in the absence of peak *AKT1* activation and *Plk1* activity. |
| | ⊣ PLoc | AKT_H | *AKT1* mediates the translocation of *FoxO1* our of the nucleus through direct phosphorylation of three conserved residues. These events create a recognition site for *14-3-3* family proteins, which export and sequester *FoxO1* in the cytosol [110]. |
| | ⊣ PLoc | Plk1 | *Plk1* interacts with and phosphorylates *FoxO1*, mainly at the G2/M phase of the cell cycle. *Plk1*-mediated phosphorylation leads to the impairment of *FoxO1*'s transcriptional activity in an *Akt*-independent manner. *Plk1*-induced *FoxO1* phosphorylation causes its nuclear exclusion [152]. |
| p21_mRNA | **p21_mRNA** = ((**FoxO1** and **FoxO3**) or (**SMAD2_3_4** or not **Myc**) and (**FoxO1** or **FoxO3**)) and not(**ZEB1_H** or **b_catenin_TCF4** or **N_bcatenin_H** or **HMGA1**) | | |
| | mRNA | | Our model requires both FoxOs to induce *p21*$^{Cip1}$ if *Myc* is active and one of the two if *Myc* is OFF or *FoxO* factors are aided by *Smad* activity [153]. This is based on data showing that both *FoxO3* and *FoxO1* bind and induce the *p21*$^{Cip1}$ promoter and that loss of *Myc* repression alone is not sufficient to induce *p21*$^{Cip1}$ [154]. Finally, high levels of *ZEB1*, *HMGA1*, or *β-catenin/TCF4* can all repress *p21*$^{Cip1}$ mRNA expression [155, 156, 157]. |
| | ← TR | FoxO3 | *p21*$^{Cip1}$ is a direct transcriptional target of *FoxO3* [154]. |
| | ← TR | FoxO1 | *p21*$^{Cip1}$ is a direct transcriptional target of *FoxO1* [154]. |
| | ← TR | SMAD2_3_4 | *Smad3–Smad4* complexes associate with *FoxO* transcription factors to induce *p21*$^{CIP1}$ [153]. |
| | ⊣ TR | ZEB1_H | *ZEB1* (old name δEF1) is a direct transcriptional repressor of the p21 promoter [155]. |
| | ⊣ TR | HMGA1 | *HMGA1* maintains cell proliferation through direct inhibition of *p21*$^{Cip1}$ [156]. |
| | ⊣ TR | N_bcatenin_H | *β-catenin/TCF4* are direct transcriptional repressors of *p21*$^{Cip1}$ [157]. |
| | ⊣ TR | b_catenin_TCF4 | *β-catenin/TCF4* are direct transcriptional repressors of *p21*$^{Cip1}$ [157]. |
| | ⊣ TR | Myc | *Myc* is a direct transcriptional repressor of the *p21*$^{Cip1}$ promoter (it is recruited by the DNA-binding *Miz-1*) [48, 158]. |
| IKKa | **IKKa** = **AKT_H** | | |

**Table S1j: GF_connect module**

| | | | |
|---|---|---|---|
| | K | *IKKα* is a subunit of the *IKK* protein complex composed of two catalytic subunits, *IKKα* and *IKKβ*, and the regulatory protein *NEMO*. Activation of the transcription factor *NF-κB* is mediated by the IKK complex, which phosphorylates and degrades the inhibitory *IκB* proteins [159]. | |
| | ← P | AKT_H | *IKKα* is phosphorylated by *AKT* at T23, and as subsequent *NF-κB* activation is induced when high *AKT* activity is observed, our model requires *AKT_H = ON* for this to occur [160]. |
| NfkB | **NfkB = IKKa** or **PAK1** | | |
| | TF | *NF-κB* is a transcription factor primarily known as a master regulator of inflammatory signaling and the immune system. Its role in cancer is partly due to its ability to aid EMT. It is activated via the destruction of its inhibitory binding partner *IκB*, which is phosphorylated by the *IKK* complex subsequently destroted [159]. | |
| | ← Loc | PAK1 | Active *PAK1* binds to with *NF-κB*-inducing kinase *NIK*, which induces degradation of *IκB* and thus activates *NF-κB* [161]. |
| | ← Ind | IKKa | *IKKα*, part of the *IKK* complex, phosphorylates and degrades the inhibitory *IκB* proteins [159]; an action that can be independent of the *IKKβ* subunit [162]. |
| c_Myb | **c_Myb = NfkB** or **E2F1** | | |
| | TF | *c-Myb* is a transcription factor that can induce the epithelial micro-RNA *miR-200* [163]. *c-Myb* is induced by *AKT*-mediated activation of *NF-κB* and/or *E2F1* [164]. | |
| | ← TR | NfkB | *NF-κB* is a direct transcriptional inducer of *c-Myb* [164]. |
| | ← TR | E2F1 | *E2F1* is a direct transcriptional inducer of *c-Myb* [164]. |

**Table S1k: Adhesion module**

| Target Node | Node Gate<br>Node Type<br>Link Type | Node Description<br>Input Node | Link Description |
|---|---|---|---|
| Integrin | **Integrin = ECM** | | |
| | Rec | Integrins are a superfamily of heterodimeric cell adhesion receptors that bind to extracellular matrix ligands, cell-surface ligands, and soluble ligands. Upon ligand binding, integrins transduce biomechanical information to the cell interior [165]. | |
| | ← Ligand | ECM | *Integrin* activation and signaling requires *integrin-ECM* attachment [166]. |
| ILK | **ILK = Integrin** | | |

**Table S1k: Adhesion module**

| | | | |
|---|---|---|---|
| | K | Integrin-linked kinase *ILK* interacts with the cytoplasmic domain of $\beta$1 integrins, and acrs as a scaffold connecting integrins to the actin cytoskeleton and other signalling pathways [167]. | |
| | ← BLoc | Integrin | Integrin-mediated adhesion to the ECM recruits and activates *ILK* [167]. |
| FAK | **FAK** = ((not **Casp3**) and (not(**Cdk1** and **CyclinB**))) and **Integrin** | | |
| | K | *FAK* is activated at integrin-ECM attachment sites in the absence of *Caspase 3*-mediated cleavage and *Cyclin B/Cdk1* activity [168, 169, 170]. | |
| | ← P | Integrin | *Integrin* activation leads to recruitment and phosphorylation of the Focal Adhesion Kinase (*FAK*) [168], one of its key signaling mediators. |
| | ⊢ Ind | CyclinB | During mitosis, cells detach most of their focal adhesions and round up. This process is *Cdk1/Cyclin B* dependent [171], and it leads to the dissociation of focal adhesion complex components including focal adhesion kinase (*FAK*), *paxillin*, and *CAS*. These proteins all change their phosphorylation status by losing active tyrosine and gaining inhibitory serine/threonine phosphorylation [170]. |
| | ⊢ Ind | Cdk1 | During mitosis, cells detach most of their focal adhesions and round up. This process is *Cdk1/Cyclin B* dependent [171], and it leads to the dissociation of focal adhesion complex components including focal adhesion kinase (*FAK*), *paxillin*, and *CAS*. These proteins all change their phosphorylation status by losing active tyrosine and gaining inhibitory serine/threonine phosphorylation [170]. |
| | ⊢ Lysis | Casp3 | Caspase 3 cleaves and deactivates *FAK* during apoptisis by separating its tyrosine kinase from its focal adhesion targeting domain. These fragments further suppress phosphorylation of intact *FAK* [169]. |
| Src | **Src** = (((**Integrin** or (**Nectin3** and **J_Ecadherin**)) and (**RTK** or **FAK**)) or (**Cdk1** and **CyclinB**)) or (**TGF_bRI** and **TGF_bRII**) | | |
| | K | *Src* is activated by *FAK* or *RTKs* at sites of integrin-ECM attachments [172, 173] aided by *Nectins* [174] and junctional *E cadherin* at cell cell contacts [175], or by *TGF$\beta$* signaling [176]. In addition, *Cyclin B / Cdk1* phosphorylate *Src* during mitosis [177]. | |
| | ← Loc | RTK | *RTKs* cooperate with *integrins* to recruit and activate *Src* kinases, which in turn help potentiate *RTK* signaling. Thus, in our model *Src* may be activated by basal *RTK* activity, and is, in turn, required for peak *RTK* activation [173]. |
| | ← PLoc | Integrin | *FAK* phosphorylation at Y397 at sites of *integrin-ECM* adhesion creates a high-affinity binding site for *Src*, which leads to the assembly of a *FAK–Src* signaling complex [172]. |
| | ← PLoc | FAK | *FAK* phosphorylation at Y397 at sites of *integrin-ECM* adhesion creates a high-affinity binding site for *Src*, which leads to the assembly of a *FAK–Src* signaling complex [172]. |

**Table S1k: Adhesion module**

| | | | |
|---|---|---|---|
| | ← PLoc | Nectin3 | *Nectin* was found to locally recruit and activate *Src*, leading to downstream phosphorylation of targets *Cdc42* and *Rac1*; downstream signaling was unable to be induced without *Nectins* [174]. |
| | ← PLoc | J _Ecadherin | *E_ cadherin* was found to upregulate *Src* signaling through localized phosphotryosine signaling in response to homophilic ligation; *Src* activity was also found to be increased around cell-cell contact sites [175]. |
| | ← BLoc | TGF_bRII | *TGFβ* activates *Src* via active *TβRI/TβRII* complexes [176]. |
| | ← BLoc | TGF_bRI | *TGFβ* activates *Src* via active *TβRI/TβRII* complexes [176]. |
| | ← P | CyclinB | Mitotic *Cyclin B/Cdk1* complexes phosphorylate and activate *c-Src* during mitosis [177]. |
| | ← P | Cdk1 | Mitotic *Cyclin B/Cdk1* complexes phosphorylate and activate *c-Src* during mitosis [177]. |
| Nectin3 | **Nectin3** = **CellDensity_Low** or **CellDensity_High** | | |
| | CAM | *Nectins* form weak adhesions between adjacent cells by binding to *Nectins* on other cells and promoting local membrane ruffling that is required for adherens junction formation [178]. As downstream effects of *Nectin3* - *Nectin* binding between two cells do not require tight junction formation and high cell density, *Nectin3* activation in our model only requires the presence of some neighbors. | |
| | ← Env | CellDensity _High | Nectins avtivate by forming weak adhesions between adjacent cells [178]. |
| | ← Env | CellDensity _Low | Nectins avtivate by forming weak adhesions between adjacent cells [178]. |
| Necl5 | **Necl5** = **FocalAdhesions** or (not(**Nectin3** and **CellDensity_High**)) | | |
| | Prot | *Necl-5* activity is controlled by co-localization with focal adhesions where it binds *Spry2* and aids receptor tyrosine kinase signaling [179]. We modeled this by turning the *Necl-5* node ON when the *Focal Adhesions* node is ON (indicating strong attachments that pull on the ECM and can form stress fibers), or in the absence of cell-cell adhesions at all sites of cell ECM-adhesion (i.e, fully surrounded with no free edge). | |
| | ⊢ Env | CellDensity _High | At high cell density, adherens junctions that surround the cell suppress integrin-mediated activation and recruitment of *Necl-5* to the cell surface across the entire cell, which releases its block on *Spry2* [179]. |
| | ⊢ Unbind | Nectin3 | *Necl-5* interacts with *Nectin3* on neighboring cells (*Nectin 3* in our model is a proxy for this, as it is activated by cell-cell contacts), which promotes downstream reorganization of the cytoskeleton to aid adherens junction formation, which, in turn releases *Necl-5* from these adhesions [179]. |

**Table S1k: Adhesion module**

| | | | |
|---|---|---|---|
| | ← Loc | FocalAdhesions | *Necl-5* is recruited to focal adhesions at the leading edge of cells by direct interactions with integrins [180]. This localization is important for its downstream effects. |
| SPRY2 | **SPRY2** = ((not **Necl5**) and **RTK**) and **Src** | | |
| | Prot | *SPRY2* (*Sprouty 2*) is a negative regulator of growth factor-induced signaling, especially *Ras/MAPK* [181]. | |
| | ← Loc | RTK | *Sprouty* proteins are activated by ligand-bound RTKs to modulate / inhibit downstream MAPK signaling [182]. |
| | ← P | Src | Growth factor-induced tyrosine phosphorylation of *Spry2* is mediated by a *Src*-like kinase [182]. |
| | ⊢ IBind | Necl5 | *Necl-5* localized to integrin clusters binds to and blocks the activity of *SPRY2* [181]. |
| J_Ecadherin | **J_Ecadherin** = (not **Casp3**) and not **MMP7** and (**Nectin3** and (**Ecadherin_mRNA_H** or **Ecadherin_mRNA**)) | | |
| | CAM | *E-cadherin* proteins are adhesion molecules required for adherens junction formation. The presence of juncitonal *E-cadherin* (J_Ecadherin = ON) in our model requires *E-cadherin* mRNA expression, *Nectin3*-mediated sensing of contact with neighboring cells (binding to nectins) [179] and inactive *MMP-7* [183] and/or Caspase 3 [184]. | |
| | ⊢ Lysis | MMP7 | Active *MMP-7* promotes EMT by cleaving *E-cadherin* protein [183]. |
| | ← BLoc | Nectin3 | Cadherins are recruited to cell-cell adhesions formed by *Nectin3-nectin* interactions between neighboring cells, where they bind to cadherins on adjacent cells to form AJs [179]. |
| | ← TL | Ecadherin_mRNA_H | *E-cadherin* mRNA expression is required for maintenance of *E-cadherin* protein. |
| | ← TL | Ecadherin_mRNA | *E-cadherin* mRNA expression is required for maintenance of *E-cadherin* protein. |
| | ⊢ Lysis | Casp3 | Caspase 3 cleaves junctional E-cadherin, dissociating it from the cell surface and blocking its ability to form adherens junctions [184]. |
| J_bcatenin | **J_bcatenin** = (not **Casp3**) and **J_Ecadherin** | | |
| | Prot | Junctional *E-cadherin* proteins binds to and recruits/sequesters *β-catenin* to adherens junctions. This negatively regulates *β*-catenin mediated transcription, but aids adherens junction formation [185]. Here we assume that *β-catenin* is localized to cell-cell junctions (*J_bcatenin* = ON) as long as there are *E-cadherin*-mediated attachments to neighboring cells (even if a cell is not fully surrounded) and *Caspase 3* is inactive [186]. | |
| | ← Per | J_Ecadherin | Junctional *E-cadherins* recruits/sequester *β-catenin* to adherens junctions [185]. |

| | | | |
|---|---|---|---|
| | ⊢ Lysis | Casp3 | Active *Caspase 3* cleaves *β-catenin* into several fragments that lose their transcriptional activity and become localized to the cytoplasm [186]. |
| J_acatenin | **J_acatenin** = **J_bcatenin** and **J_Ecadherin** | | |
| | Adap | Junctional *β-catenin* binds to and recruitcs *α-catenin* to adherens junctions. *α-catenin* links *β-catenin* to the actin cytoskeleton to stabilize adherens junctions [185]. | |
| | ← Compl | J_Ecadherin | Junctional *E-cadherin*-bound *β-catenin* binds to and recruitcs *α-catenin* to adherens junctions [185]. |
| | ← Compl | J_bcatenin | Junctional *β-catenin* binds to and recruitcs *α-catenin* to adherens junctions [185]. |

**Table S1l: CIP module**

| Target Node | Node Gate<br>Node Type<br>Link Type | Node Description<br>Input Node | Link Description |
|---|---|---|---|
| FocalAdhesions | **FocalAdhesions** = **Integrin** and **FAK** and **ECM** and (**Stiff_ECM** or **Basement_Membrane** or (**YAP** and (**Rac1** or **Rac1_H**) and **IQGAP1_LeadingE**)) | | |
| | MSt | *Focal adhesions* form at sites of *Integrin-ECM* attachment and clustering [187]. In order to take into account the effect of stiff ECM [188] as well as positive feedback between focal adhesion formation and horizontal cell polarization that creates an active leading edge, here we assume that force-generating Focal Adhesion formation requires ECM-Integrin attachments and *FAK* [189], and either strong traction force generation supported by a stiff ECM, adhesion to a basement membrane, or the existence of a leading edge with active *Rac1* [190] and *IQGAP1* [191, 192, 193] supported by *YAP*-mediated upregulation of adhesion and focal adhesion-associated proteins [194]. | |
| | ← Loc | Stiff_ECM | Adhesion to stiff ECM engages the *FAK/phosphopaxillin/vinculin* pathway, which generate a fluctuating "tugging" action on the ECM and probe ECM rigidity to aid *focal adhesion* formation and migration towards regions of stiffer ECM (durotaxis) [188]. |
| | ← Loc | Basement_Membrane | Adhesion to an intact basement membrane can also engage the *FAK/phosphopaxillin/vinculin* pathway, as it supports its "tugging" action to aid *focal adhesion* formation and migration towards regions of stiffer ECM (durotaxis) [188]. |
| | ← Loc | ECM | *Focal adhesions* form at sites of *Integrin-ECM* attachment and clustering [187]. |
| | ← Loc | Integrin | *Focal adhesions* form at sites of *Integrin-ECM* attachment and clustering [187]. |

**Table S1l: CIP module**

| | | | |
|---|---|---|---|
| | ← Loc | FAK | *FAK* activation at sites of cell-ECM attachments (driven by force generation) can increase paxillin phosphorylation and strengthen cytoskeletal linkage and vinculin recruitment to such adhesions, resulting in *focal adhesion* maturation [189]. |
| | ← TR | YAP | *YAP* is a transcriptional inducer of *f integrins* and *FA* docking proteins, and promotes *focal adhesion* formation by increasing cell spreading and *RhoA GTPase* activity [194]. |
| | ← Compl | IQGAP1 _LeadingE | In migrating cells *IQGAP1* localizes to lamellipodia at the leading edge, recruited by active *RTKs* synergistically activated here by *integrin-RTK* crosstalk. Here, *IQGAP1* supports *Rac1* activation and *focal adhesion* formation; it required for migration in response to growth signals such as *PDGF*, *VEGF*, ect [191, 192, 193]. |
| | ← Loc | Rac1 | Active *Rac1* promotes the association of nonmuscle myosin II (MIIA) with *focal adhesions* at the leading edge during cell migration, aiding the assembly of mini- filaments in *focal adhesions*. These promote further assembly of *focal adhesions* and modulation of the traction forces cells exert on the ECM [190]. |
| | ← Loc | Rac1_H | High *Rac1* activity promotes the association of nonmuscle myosin II (MIIA) with *focal adhesions* at the leading edge during cell migration, aiding the assembly of mini- filaments in *focal adhesions*. These promote further assembly of *focal adhesions* and modulation of the traction forces cells exert on the ECM [190]. |
| Stress _Fibers | **Stress_Fibers** = ((not **CellDensity_High**) and **Stiff_ECM**) and **FocalAdhesions** | | |
| | MSt | | In order to model the independent effects of cell density and matrix stiffness on stress fiber formation, we assume that they require both the absence of high cell density, and the presence of focal adhesions attached to a stiff ECM. |
| | ⊣ Ind | CellDensity _High | *High cell density* blocks stress fiber formation by forbidding cells access to a large enough area to spread and exert force on the *ECM* [195]. |
| | ← Ind | Stiff_ECM | In the absence of a sufficiently stiff *ECM*, *Focal Adhesions* are small and stress fibers are less abundant or fail to form, as cells cannot generate sufficient traction [196]. |
| | ← ComplProc | FocalAdhesions | *Stress Fibers* are anchored to the *ECM* via strong, stable *Focal Adhesions* [196]. |
| YAP | **YAP** = (**FocalAdhesions** and **Stress_Fibers**) and (not((((**ApicalBasal_Pol** and **J_acatenin**) and **AMOT**) and **Merlin**) and **Lats1_2**)) | | |

**Table S1l: CIP module**

| | | | |
|---|---|---|---|
| | TF | | YAP is a mechanosensitive transcriptional regulator of proliferation and migration, and its activation is controlled by both the cell's ability to spread on an ECM (*FocalAdhesions* and *Stress_Fibers*), and the lack of apical-basal polarity with mature adherens junctions that can sequester *YAP* in the cytoplasm by binding and inhibitory phosphorylation. Experimental evidence indicates that in cells that maintain apical-basal polarity, several junctional proteins ($\alpha$*-catenin*, *AMOT*, *Merlin*) and inhibitory kinases (*Lats1* and *Lats2* work together to sequester and block *YAP* [197, 198]. |
| | ⊣ Per | J_acatenin | Junctional *alpha-catenin* binds *YAP* and sequesters it in the cytoplasm [199] [200] [201]. This also concentrates *YAP* close proximity to junction-localized Hippo pathway components such as *Lats1/2*, *Merlin* and *Amot.* |
| | ← Ind | FocalAdhesions | *YAP* activation is abolished by the absence of *stress fibers* anchored to *focal adhesions*, even in the absence of inhibitory Hippo signaling [195, 202]. |
| | ← Ind | Stress _Fibers | *YAP* activation is abolished by the absence of *stress fibers* anchored to *focal adhesions*, even in the absence of inhibitory Hippo signaling [195, 202]. |
| | ⊣ Ind | ApicalBasal _Pol | Full inhibition of *YAP* by Hippo signaling linked to adherens and tight junction formation requires the establishment of *apical-basal polarity*, as cells at the edges of monolayers or in decreased cell density areas have active (nuclear) *YAP* in spite of strong remaining attachments to neighboring cells [202, 203]. |
| | ⊣ IBind | Lats1_2 | The *Lats1* and *Lats2* tumor suppressor kinases bind to and phosphorylate *YAP* in vitro and in vivo [204, 198]. |
| | ⊣ Compl | AMOT | *AMOT* localizes to tight junctions, where it suppresses *YAP* activity by direct binding and recruitment of *YAP* inhibitory kinase *LATS2* [197]. By binding both and *YAP/TAZ*, *AMOT* works as a scaffold that connects *LATS1/2* to both its activator *MST1* and its target *YAP/TAZ* [205]. |
| | ⊣ BLoc | Merlin | *Merlin* localizes to adherens junctions where it activates the *Hippo pathway* by binding to and recruiting *LATS1/2* kinases and *YAP/TAZ* to *adherens junctions* [206]. In the absence of *Merlin*, Hippo pathway components fail to block *YAP* activity [207]. *Merlin* - *YAP* binding requires active (phosphorylated) *AMOT* [208]. |
| TRIO | **TRIO = YAP** | | |
| | GEF | | *TRIO* is a *Rac1*-activating GTP-exchange factor induced by *YAP* [203, 209]. |
| | ← TR | YAP | *YAP* is a transcriptional inducer of *TRIO* [209]. |
| WT1 | **WT1 = YAP** | | |
| | TF | | The Wilms Tumor 1 (*WT1*) transcription factor is a repressor of *E-cadherin* expression [203]. Its nuclear localization is controlled by *YAP* binding [209]. |

**Table S1l: CIP module**

| | | | |
|---|---|---|---|
| | $\leftarrow$ BLoc | YAP | *YAP* binds to and controls nuclear localization of the Wilms Tumor 1 (*WT1*) transcription factor [209]. |
| TAZ | **TAZ** = **Stress_Fibers** and (not((((**ApicalBasal_Pol** and **J_acatenin**) and **AMOT**) and **Merlin**) and **Lats1_2**)) | | |
| | TF | | *TAZ* is a mechanosensitive transcriptional regulator of cell migration, and its activation is controlled by both the cell's ability to spread on stiff ECM (*Stress_Fibers* = ON) and the lack of apical-basal polarity with mature adherens junctions that can sequestered *TAZ* in the cytoplasm by binding and inhibitory phosphorylation [202, 203]. In cells that maintain apical-basal polarity, junctional proteins (*α-catenin*, *AMOT*, *Merlin*) and inhibitory kinases (*Lats1* and *Lats2*) work together to sequester and block *TAZ* [199, 200, 201, 210, 198]. |
| | $\dashv$ BLoc | J_acatenin | Junctional *alpha-catenin* binds *YAP/TAZ* and sequesters them in the cytoplasm [199, 200, 201, 210]. This also concentrates *YAP/TAZ* close proximity to junction-localized Hippo pathway components such as *Lats1/2*, *Merlin* and *Amot.* |
| | $\leftarrow$ Ind | Stress_Fibers | *TAZ* activation is abolished by the absence of *stress fibers* anchored to *focal adhesions*, even in the absence of inhibitory Hippo signaling [195, 202]. |
| | $\dashv$ Ind | ApicalBasal_Pol | Full inhibition of *TAZ* by Hippo signaling linked to adherens and tight junction formation requires the establishment of apical-basal polarity, as cells at the edges of monolayers or in decreased cell density areas have active (nuclear) *TAZ* in spite of strong remaining attachments to neighboring cells [202, 203]. |
| | $\dashv$ P | Lats1_2 | The *Lats1* and *Lats2* tumor suppressor kinases bind to and phosphorylate *YAP/TAZ* in vitro and in vivo [204, 198, 210]. |
| | $\dashv$ IBind | AMOT | *AMOT* localizes to tight junctions, where it suppresses *YAP/TAZ* activity by direct binding and recruitment of *YAP* inhibitory kinase *LATS2* [197, 210]. By binding both and *YAP/TAZ*, *AMOT* works as a scaffold that connects *LATS1/2* to both its activator *MST1* and its target *YAP/TAZ* [205]. |
| | $\dashv$ IBind | Merlin | *Merlin* localizes to adherens junctions where it activates the Hippo pathway by binding to and recruiting *LATS1/2* kinases and *YAP/TAZ* to adherens junctions [206]. In the absence of *Merlin*, Hippo pathway components fail to block *YAP/TAZ* activity [207, 210]. |
| Ecadherin_mRNA_H | **Ecadherin_mRNA_H** = **Ecadherin_mRNA** and (not(**YAP** and **WT1**)) | | |
| | mRNA | | Experiments show that *YAP* and *WT1* suppress but do not abolish *E-cadherin* protein expression in areas of lowered cell density [203]. To model this, we introduced a *Ecadherin_mRNA_H* node that is blocked by *YAP/WT1* repressor complexes (requiring their joint nuclear localization). *Ecadherin_mRNA_H*, in turn, must be ON to allow cells to establish a ring of adherens junctions sufficient for apical-basal polarity. |

**Table S1l: CIP module**

| | | | |
|---|---|---|---|
| | $\vdash$ Compl | YAP | Active *YAP* binds *WT1* and localizes it to the nucleus, where they form a complex a the *E-cadherin* promoter and reduce its transcription [203]. |
| | $\vdash$ Compl | WT1 | Active *YAP* binds *WT1* and localizes it to the nucleus, where they form a complex a the *E-cadherin* promoter and reduce its transcription [203]. |
| | $\leftarrow$ Per | Ecadherin _mRNA | High *E-cadherin* mRNA expression requires basal levels of *E-cadherin*. Our previously published epithelial model assumed this to be true [148], whereas in the current model this is only the case when *E-cadherin* expression is not fully inhibited by EMT-promoting repressors (modeled as acting on the *Ecadherin_ mRNA* node). |
| ApicalBasal _Pol | **ApicalBasal_Pol** = (**ECM** and ((((**CellDensity_High** and **Nectin3**) and **J_Ecadherin**) and **J_bcatenin**) and **J_acatenin**)) and (**Ecadherin_mRNA_H** or (not **Horizontal_Pol**)) | | |
| | MSt | | In addition to a need for high cell density and cell-cell adhesion proteins that help assemble adherens junctions (*Nectin3*, *J_Ecadherin*, *J_ bcatenin*, *J_ acatenin*), we assumed that either high (unimpeded) *E-cadherin* mRNA expression or lack of a horizontally polarized cell morphology are required for the establishment of apical-basal polarity. |
| | $\leftarrow$ Ind | CellDensity _High | Establishment of *apical-basal polarity* requires a ring of adherens and tight junctions that can only form in high cell density [211]. |
| | $\leftarrow$ ComplProc | ECM | Establishment of *apical-basal polarity* requires an underlying surface such as *ECM* to define a basal side. |
| | $\leftarrow$ Ind | Nectin3 | A key driver of *apical-basal polarization*, *Par-3*, is recruited to newly formed cell-cell adhesions by *Nectin-3* binding [211]. |
| | $\leftarrow$ Ind | J _Ecadherin | Formation of adherens junctions is a prerequisite for tight junction assembly, which is, in turn, required for *apical-basal polarity* [211]. |
| | $\leftarrow$ Ind | J_bcatenin | Formation of adherens junctions is a prerequisite for tight junction assembly, which is, in turn, required for *apical-basal polarity* [211]. |
| | $\leftarrow$ Ind | J_acatenin | Formation of adherens junctions is a prerequisite for tight junction assembly, which is, in turn, required for *apical-basal polarity* [211]. |
| | $\leftarrow$ Ind | Ecadherin _mRNA_H | Formation of adherens junctions in high concentration around the cell is required for *apical-basal polarity* [211], and thus aided by high *E-cadherin* protein expression. |
| | $\vdash$ Ind | Horizontal _Pol | Horizontal polarization and apical-basal (vertical) polarization are mutually exclusive; cells must first lose the asymmetry between their leading and trailing edge in order to establish a ring of adherens and tight junctions. |
| N_bcatenin | **N_bcatenin** = (not **Casp3**) and (not(**ApicalBasal_Pol** and **GSK3**)) | | |

**Table S1l: CIP module**

| | | | |
|---|---|---|---|
| | TF | When released from cell-cell junctions, *β-catenin* localizes to the nucleus and induces genes that promote proliferation and EMT [212]. | |
| | $\vdash$ Deg | GSK3 | The cytoplasmic pool of *β-catenin* not tied to junctions is highly unstable due to multiple phosphorylations promoting its proteasome-mediated degradation. This phosphorylations is maintained by the "*β-catenin* Destruction Complex" composed of the tumor suppressor *APC*, the scaffolding protein *Axin*, and the serine/threonine kinases *GSK3β* and *CK1* (casein kinase 1, which primes *GSK3*). When *GSK3* is inhibited, unphosphorylated *β-catenin* accumulates, translocates to the nucleus, and promotes transcription [212]. |
| | $\vdash$ Loc | ApicalBasal_Pol | Junctional *E-cadherins* recruits/sequesters *β-catenin* to adherens junctions and block *β-catenin* nuclear localization [185]. Here we assume that cells able to form a ring of adherens junctions and establish apical-basal polarity lack nuclear *β-catenin*. |
| | $\vdash$ Lysis | Casp3 | Active Caspase 3 cleaves *β-catenin* into several fragments that lose their transcriptional activity and become localized to the cytoplasm [186]. |
| Mst1_2 | **Mst1_2 = ApicalBasal_Pol** | | |
| | K | In cells that establish apical-basal cell polarity, *Mst1* and *2* activate the Hippo pathway by phosphorylating *Lats1/2* kinases to promote *YAP/TAZ* inhibition [213, 214, 215]. | |
| | $\leftarrow$ Ind | ApicalBasal_Pol | Activation of the Hippo pathway by *Mst1/2* requires apical-basal cell polarity [215]. |
| Lats1_2 | **Lats1_2 = Mst1_2** and **Merlin** | | |
| | K | *Mst1/2* activate *Lats1/2* kinases by phosphorylation, aided by their mutual binding to *Merlin*. | |
| | $\leftarrow$ P | Mst1_2 | *Mst1/2* phosphorylate and activate *Lats 1 / 2* kinases to promote *YAP/TAZ* inhibition [213, 214]. |
| | $\leftarrow$ BLoc | Merlin | *Merlin* binds to *Lats1/2* and recruits it to the plasma membrane near adherens and tight junctions, where *Mst1/2* can phosphorylate it [206]. |
| AMOT | **AMOT = Lats1_2** and **Merlin** | | |
| | Adap | In our model, *AMOT* = ON represents phosphorylated and tight junction localized *AMOT*, which requires *Merlin* binding and *Lats1/2* mediated phosphorylation [216, 217]. | |
| | $\leftarrow$ P | Lats1_2 | *Lats1/2* kinases phosphorylate the N-terminal regions of *Amot*, disrupting its interaction with F-actin [218]. As tight junction localized *AMOT* aids Hippo signaling whereas F-actin bound *AMOT* hinders it [216], *AMOT* = ON in our model represents the phosphorylated, TJ-bound protein. |

**Table S1l: CIP module**

| | | | |
|---|---|---|---|
| | ← BLoc | Merlin | *Merlin* binds to *AMOT* proteins and sequesters them to tight junctions [218], shifting its activity from cytoplasmic (where it binds *F-actin* and blocks the *Rac1* inhibitor *Rish1*) to junctional (where it forms a scaffold for Hippo signaling) [217]. |
| miR_29c | **miR_29c** = **YAP** | | |
| | miR | *YAP* induces the expression of microRNA *miR-29c* to target *PTEN* for degradation [219]. | |
| | ← TR | YAP | *YAP* is a direct inducer of *miR-29c* [219]. |
| PTEN_c | **PTEN_c** = (not **miR_29c**) and ((**S6K** and (not(**ERK** and **GSK3**))) or ((not **ERK**) and (not **GSK3**))) | | |
| | Ph | Cellular *PTEN* is a tumor supressor phosphatase that reverses PIP2 → PIP3 conversion carried out by *PI3K*, and thus supresses *PI3K / AKT* signaling [220]. The combinatorial effect of the distinct *PTEN* regulators included below is not well documented. We chose to model cytoplasmic *PTEN* availability as ON in the absence of *miR-29c* [219]. In addition, we combined the positive and negative effects of *S6K*, *ERK* and *GSK3* by assuming that *PTEN* is blocked by the joint action of *ERK* and *GSK3* in the presence of *S6K*, and by either inhibitor in its absence. | |
| | ⊢ TR | ERK | *ERK* activation suppresses *PTEN mRNA* and protein expression [221, 222]. |
| | ← P | S6K | *S6K* phosphorylates *PTEN*, which leads to *PTEN* deubiquitination and export from the nucleus to the cytoplasm [223]. |
| | ⊢ P | GSK3 | *GSK3* phosphorylates *PTEN* on Thr366 which leads to destabilization of the *PTEN* protein [224]. |
| | ⊢ Ind | miR_29c | *YAP* induces transcription of *miR-29*, which in turn binds to *PTEN mRNA* to block its translation [219]. |

**Table S1m: Migration_SW module**

| Target Node | Node Gate<br>Node Type<br>Link Type | <br>Node Description<br>Input Node | <br><br>Link Description |
|---|---|---|---|
| Merlin | **Merlin** = ((**J_bcatenin** and **J_acatenin**) and (not(**PAK1** and **ILK**))) and (not **AKT_H**) | | |
| | Prot | *Merlin* is functionally localized and able to mediate Hippo signaling when recruited by junctional *β*- and *α-catenin*, and not phosphorylated by high *AKT1* or the cooperative action of *PAK1* and *ILK*. Since the *AKT_B* node in our model represents basal *AKT* activity which co-occurs with *Merlin*-mediated contact inhibition, we choose the peak *AKT_H* node to mark the level of *AKT* signaling required to block *Merlin*. | |

#### Table S1m: Migration_SW module

| | | | |
|---|---|---|---|
| | ⊢ P | AKT_H | *Akt* directly binds to and phosphorylates *Merlin* at Thr230 and Ser315, blocking its ability to bind its regular partners and promoting its degradation [225]. |
| | ⊢ Ind | ILK | *ILK* suppresses the Hippo pathway via phosphorilation and inhibition of *MYPT1-PP1*, leading to inactivation of *Merlin* [226]. |
| | ← Loc | J_bcatenin | *Merlin* binds to *β-catenin* at adherens junctions, and disruption of *β-catenin* expression recoiples cell density sensing from Hippo signaling downstream *Merlin* [227]. |
| | ← BLoc | J_acatenin | *Merlin* binds to *α-catenin* to localize to adherens junctions, where it plays a role in their maturation, links AJ formation and the *Par3* polarity complex, and orchestrates Hippo signaling [227, 228]. |
| | ⊢ P | PAK1 | *Pak1* can directly phosphorylate *Merlin* at Ser518 [229]. This phosphorylation prevents it from binding to /textitAMOT and *Lats1/2* [230], and thus carrying out its contact inhibitory function. |
| IQGAP1 _LeadingE | **IQGAP1_LeadingE** = ((not **CellDensity_High**) and **FocalAdhesions**) and ((**Horizontal_Pol** and (**Rac1** or **Rac1_H**)) or **Grb2**) | | |
| | Adap | In our model, *IQGAP1_LeadingE* = ON represents *IQGAP1* localized near the leading edge of a horizontally polarized cell, where it links focal adhesion-mediated and *RTK*-mediated signaling [192]. Its recruitment is sustained by well-established horizontal polarity, or induced by *Rac1* [231] or *RTK-Grb2* binding [91, 232, 233]. | |
| | ⊢ Ind | CellDensity _High | *IQGAP1* localization to the leading edge is blocked by high cell density that leaves no free edge. |
| | ← BLoc | Grb2 | *Grb2* bound to active *RTK* binds and recruits *IQGAP1*, aiding its enrichment at the leading edge where the concentration of active *RTKs* is generally higher [91, 232, 233]. |
| | ← ComplProc | FocalAdhesions | *IQGAP1* interacts with focal adhesion proteins and tyrosine kinase receptors to link focal adhesion and *RTK* signaling. It is thus enriched at the leading edge by increased active FA formation [192]. |
| | ← ComplProc | Horizontal _Pol | *IQGAP1* localization to the leading edge is reinforced by horizontal polarization and the stabilization of a leading edge [192, 234]. |
| | ← Compl | Rac1 | Active *Rac1* forms a complex with *IQGAP1* and *CLIP-170* and recruits both to the base of the leading edge, where their aid cytoskeletal reorganization, horizontal polarization and directed migration [231]. |
| | ← Compl | Rac1_H | Active *Rac1* forms a complex with *IQGAP1* and *CLIP-170* and recruits both to the base of the leading edge, where their aid cytoskeletal reorganization, horizontal polarization and directed migration [231]. |
| Rassf1a | **Rassf1a** = not(**ZEB1** or (**TGF_bRI** and **TGF_bRII**)) | | |

**Table S1m: Migration_SW module**

| | | | |
|---|---|---|---|
| | Prot | | The cytoskeleton-associated protein *Rassf1a* serves as a scaffold to bring *Smurf1* (an E3 ubiquitin-protein ligase) to *RhoA*, marking it for degradation [235]. It is targeted for degradation by *TGF-β* signaling [41] and repressed by *ZEB1*. |
| | ⊣ Ind | TGF_bRII | In response to *TGF-β*, *Rassf1a* is recruited to *TGF-β receptor I* and targeted for degradation by the co-recruited E3 ubiquitin ligase *ITCH* [41]. |
| | ⊣ Ind | TGF_bRI | Activaton of *TGF-β receptor I* (which recruits *Rassf1a*) requires a complex with the ligand as well as *TGF-β receptor II* [236]. |
| | ⊣ TR | ZEB1 | *ZEB1* is a direct transcriptional repressor of *Rassf1a* [237]. |
| RhoA | **RhoA** = **AKT_H** or not **Rassf1a** or (**RhoA** and **Horizontal_Pol** and (**Myc** or not **miR_34**)) | | |
| | GTPa | | *RhoA* is a member of the Rho family of small GTPases; it promotes reorganization of the actin cytoskeleton. It's primarily role in mesenhymal of crawling cell migration is to promote actin stress fibers formation and actomyosin contractility at the rear of a horizontally polarized cell, where it aids detachment [238]. As our model focuses exlusively on mesenchymal migration, the active state of the *RhoA* node corresponds to its activity at the rear of the cell, and thus contributes to horizontal polarity. Thus, *RhoA* overexpression cannot be adequarly modeled by simply locking *RhoA* ON; changing its relationship to the *Horizontal_Pol* node would also be required. At the back of the cell, *RhoA* is activated by mitogen-drive *AKT* signaling [239] or the absence of its inhibitor *Rassf1a* [235]. Alternatively, *RhoA* can maintain its own activation at the back of the cell [240, 241] as long as its Myc-mediated transcription is not couteracted by *miR-34* [242, 243]. |
| | ← Ind | AKT_H | *AKT* phosphorylation of the *DLC1* tumor suppressor, a Rho GTPase-activating protein (RhoGAP), increases *RhoA-GTP* and thus boosts *RhoA* acticity [239]. |
| | ⊣ Ind | Rassf1a | The cytoskeleton-associated protein *Rassf1a* serves as a scaffold to bring *Smurf1* (an E3 ubiquitin-protein ligase) to *RhoA*, marking it for degradation [235]. |
| | ← Per | RhoA | Active *RhoA* binds to the PH domain of its GEF, locally increaseing GEF concentration leading and further activating of *RhoA* in a direct positive feedback [240, 241]. |
| | ← Per | Horizontal_Pol | We assume that the positive feedback auto-regulation of *RhoA* requires its concentration to the back of the cell, modeled via the ON state of the *Horizontal_Pol* node. |
| | ← Ind | Myc | *Myc*, together with *Skp2*, *Miz1* and *p300*, is a direct transcriptional inducer of *RhoA* [243]. |
| | ⊣ Ind | miR_34 | *miR-34* targets *c-Myc* mRNA for degradation, and was shown to lower its ability to induce *RhoA* transcription [242]. |
| Horizontal_Pol | **Horizontal_Pol** = not **ApicalBasal_Pol** and **ECM** and **IQGAP1_LeadingE** and **RhoA** and **FocalAdhesions** and **TAZ** and **FAK** and **RhoA** | | |

**Table S1m: Migration_SW module**

| | | | |
|---|---|---|---|
| | MSt | For cells to establish and maintain horizontal polarization, our model requires the lack of apical-basal polarization, the presence of an ECM, as well symmetric IQGAP1 and *RhoA* localization [244, 238], focal adhesions and spreading (FAK [244, 245] and TAZ [194]). | |
| | ← Ind | ECM | Horizontal polarization requires a leading edge with lamellipodia and a trailing edge linked to stress fibers; both of which require adhesions to an ECM. |
| | ← Ind | FAK | *FAK* activation at nascent adhesions at the leading edge is required for ongoing cell spreading, which is a prerequisite of ongoing *FA* maturation and maintenance of an active leading edge [244, 245]. |
| | ← Loc | FocalAdhesions | Horizontal polarization requires a leading edge with lamellipodia and a trailing edge linked to stress fibers, both of which depend on *Focal Adhesions* linked to the actin cytoskeleton. |
| | ← Ind | TAZ | *TAZ*-null cells lose their ability to spread on the ECM, indicating that *TAZ* transcriptional activity is required for the establishment of horizontal polarization [194]. (This is in contrast to *YAP*-null cells, which cannot even form *focal adhesions.*) |
| | ⊢ Ind | ApicalBasal_Pol | Apical-basal and horizontal polarization are mutually exclusive; cells must first lose their textitapical-basal polarity before they are able to establish horizontal polarization. |
| | ← Per | IQGAP1_LeadingE | Localization of *IQGAP1* to the leading edge is required for the establishment of horizontal polarization. Together with the adenomatous polyposis coli (*APC*) protein that is also recruited to the leading edge by active *Rac1* and *Cdc42*, *IQGAP1* links the actin cytoskeleton to microtubule dynamics to establish cell polarization [244]. |
| | ← Ind | RhoA | Localization of *RhoA* to the back of the cell and it's effect on cell contractility are required for horizontal polarization [238]. |
| Rac1 | **Rac1** = not **Casp3** and **FocalAdhesions** and **Necl5** and (**Horizontal_Pol** or **SMO**) and **TRIO** and **AKT_B** and (not(**Merlin** and **Nectin3** and **J_Ecadherin**) or **Stiff_ECM**) | | |
| | GTPa | In our model *Rac1* activation on stiff ECM requires the absence of *caspase 3*, Focal Adhesion formation, *Necl-5* leading edge localization, *TRIO* expression, at least basal *AKT* activity, and horizontal polarization or *SMO* activation. In addition, on soft ECM *Rac1* activity is inhibited by the cooperative action of *Merlin*, *Nectin3* and *E-cadherin* at adherens and tight junctions [246, 217]. | |
| | ← Per | Stiff_ECM | *Rac1* activated at *focal adhesions* by *FAK* is enhanced by force generation supported by stiff ECM, leading to increased intracellular stiffness [247]. |
| | ← Ind | SMO | *Shh* signaling stimulates *Rac1* (and RhoA) via *SMO* through a Gi protein [248]. |

**Table S1m: Migration_SW module**

| | | | |
|---|---|---|---|
| | ← Ind | AKT_B | *AKT* phosphorylates the *Rac1* GEF *Tiam1*, which boosts its interaction with *14-3-3* scaffold proteins and increase *Tiam1* stability. This results in increased *Rac1* activation and enhanced migration [249]. |
| | ⊢ Compl | Nectin3 | During initial cell-cell contact and adherens junciton initiation, cadherins and nectins cooperate to briefly induce, but then rapidly suppress Rac1 [246]. |
| | ⊢ Loc | Necl5 | *Necl-5* associates with integrins at the leading edge, where it promotes the activation of *Rac1* and *Cdc42*. *Necl-5* is required for serum-and *PDGF*-induced cell motility cell motility, an effect that does not require Nectin-3 binding on neighboring cells [250]. |
| | ⊢ Compl | J_Ecadherin | During initial cell-cell contact and adherens junciton initiation, cadherins and nectins cooperate to briefly induce, but then rapidly suppress Rac1 [246]. |
| | ← Per | FocalAdhesions | *Rac1* activated at *focal adhesions* by *FAK* is enhanced by force generation supported by stiff ECM, leading to increased intracellular stiffness [247]. |
| | ← GEF | TRIO | *TRIO* is a GEF that controls *Rac1* activation during migration [203] as well as proliferation [251]. |
| | ⊢ Ind | Merlin | A protein complex that includes *Merlin* sequesters *Angiomotin* to tight junctions, releasing it from binding the *Rac1*-inhibitor *Rich1* [217]. |
| | ← Per | Horizontal_Pol | Growth of the microtubule network at leading-edge lamellipodia activates *Rac1* to drive local actin polymerization and further lamellipodial protrusions, thus supporting the maintenance of horizontal polarization [252]. |
| | ⊢ Lysis | Casp3 | Caspase 3 cleaves *Rac1* at two sites, resulting in the inactivation of its GTPase activity and *PAK1* binding [253]. |
| Rac1_H | **Rac1_H** = **Rac1** and not **Casp3** and **FocalAdhesions** and **Necl5** and **Horizontal_Pol** and **TRIO** and **AKT_H** and **SMO** and (not(**miR_200_c** or **miR_200_b** or **miR_34**) and not(**Merlin** and **Nectin3** and **J_Ecadherin**) and **Stiff_ECM**) | | |
| | GTPa | | Full (high) *Rac1_H* activation requires requires the absence of *caspase 3*, a stiff ECM, Focal Adhesion formation, *Necl-5* leading edge localization, *TRIO* expression, at least basal *AKT* activity, horizontal polarization or *SMO* activation, the absence of *miR-200* [254] or *miR-34* [255] microRNAs, as well as the absence of inhibition by the cooperative action of *Merlin*, *Nectin3* and *E-cadherin* [246, 217]. |
| | ← Per | Rac1 | As the *Rac1* node represents moderate levels of *Rac1* activity, it's ON state is required for *Rac1_H* activation. |
| | ← Per | Stiff_ECM | *Rac1* activated at *focal adhesions* by *FAK* is enhanced by force generation supported by stiff ECM, leading to increased intracellular stiffness [247]. |
| | ← Ind | SMO | *Shh* signaling stimulates *Rac1* (and RhoA) via *SMO* through a Gi protein [248]. |

**Table S1m: Migration_SW module**

| | | | |
|---|---|---|---|
| | ← Ind | AKT_H | *AKT* phosphorylates the *Rac1* GEF *Tiam1*, which boosts its interaction with *14-3-3* scaffold proteins and increase *Tiam1* stability. This results in increased *Rac1* activation and enhanced migration [249]. |
| | ⊢ Compl | Nectin3 | During initial cell-cell contact and adherens junciton initiation, cadherins and nectins cooperate to briefly induce, but then rapidly suppress Rac1 [246]. |
| | ⊢ Loc | Necl5 | *Necl-5* associates with integrins at the leading edge, where it promotes the activation of *Rac1* and *Cdc42*. *Necl-5* is required for serum-and *PDGF*-induced cell motility cell motility, an effect that does not require Nectin-3 binding on neighboring cells [250]. |
| | ⊢ Compl | J _Ecadherin | During initial cell-cell contact and adherens junciton initiation, cadherins and nectins cooperate to briefly induce, but then rapidly suppress Rac1 [246]. |
| | ← Per | FocalAdhesions | *Rac1* activated at *focal adhesions* by *FAK* is enhanced by force generation supported by stiff ECM, leading to increased intracellular stiffness [247]. |
| | ← GEF | TRIO | *TRIO* is a GEF that controls *Rac1* activation during migration [203] as well as proliferation [251]. |
| | ← Ind | Merlin | A protein complex that includes *Merlin* sequesters *Angiomotin* to tight junctions, releasing it from binding the *Rac1*-inhibitor *Rich1* [217]. |
| | ← Per | Horizontal _Pol | Growth of the microtubule network at leading-edge lamellipodia activates *Rac1* to drive local actin polymerization and further lamellipodial protrusions, thus supporting the maintenance of horizontal polarization [252]. |
| | ⊢ Ind | miR_34 | Though *miR-34* does not appear to directly target *Rac1* mRNA [256], its overexpression blocks GTP-bound (active) *Rac1* [255]. |
| | ⊢ RNAi | miR_200 _b | *miR-200b/c-3p* represses *Rac1* mRNA by targeting its 3' UTR [254]. |
| | ⊢ RNAi | miR_200_c | *miR-200b/c-3p* represses *Rac1* mRNA by targeting its 3' UTR [254]. |
| | ⊢ Lysis | Casp3 | Caspase 3 cleaves *Rac1* at two sites, resulting in the inactivation of its GTPase activity and *PAK1* binding [253]. |
| PAK1 | **PAK1 = Rac1 or Rac1_H** | | |
| | K | The *p21*-Activated kinase 1 *PAK1*, a serine-threonine kinase interacts with the Rho GTPases *RAC1* and *CDC42* to drive migration, survival, cell cycle, EMT, stress response and inflammation [257]. It is activated by *Rac1*, which binds to *PAK1* and stimulates its kinase activity [258]. | |
| | ← Compl | Rac1 | *Rac1* binds to *PAK1* and stimulates its kinase activity [258]. |

**Table S1m: Migration_SW module**

| | | | |
|---|---|---|---|
| | ← Compl | Rac1_H | *Rac1* binds to *PAK1* and stimulates its kinase activity [258]. |
| Migration | **Migration** = **Horizontal_Pol** and **Stress_Fibers** and **FocalAdhesions** and **PAK1** and **Rac1** | | |
| | MSt | | In the current model, the *Migration* node represents migration by a polarized cell, whereas *Fast_Migration* tracks rapid mesenchymal migration typical of a cell that underwent EMT. *Migration* requires horizontal polarization, stress fiber maintenance, focal adhesion formation and the activity of *Rac1* and *PAK1* kinases [259, 260]. |
| | ← Ind | FocalAdhesions | Mesenchymal style migration requires force generation via stress fibers anchored by *focal adhesions* [259]. |
| | ← Ind | Stress_Fibers | Mesenchymal style migration requires force generation via stress fibers anchored by *focal adhesions* [259]. |
| | ← Ind | Horizontal_Pol | Directed mesenchymal style migration requires *horizontal cell polarization* with a well defined leading and trailing edge [259]. |
| | ← ComplProc | Rac1 | *Rac1* GTPase signals through *WRC* (WAVE regulatory complex) to drive *Arp2/3*-dependent lamellipodia formation at the leading edge of a migrating cell [238]. |
| | ← ComplProc | PAK1 | Active *PAK1* coordinates a series of cytoskeletal changes at the leading edge that are required for migration and invasion. These include: a) inhibition of myosin light chain kinase in order to decrease contractility if the leading lamellipodium and loss of established actin stress fibers and very strong *focal adhesions* at the leading edge; b) retraction of protrusions and the cell body at the sides and trailing edge with no active *PAK1*; c) suppressing actin filament turnover and promoting leading edge stabilization; d) promote membrane ruffle formation [259, 260]. |
| Fast_Migration | **Fast_Migration** = **Migration** and **Horizontal_Pol** and **Stress_Fibers** and **FocalAdhesions** and **PAK1** and **Rac1_H** | | |
| | MSt | | The *Fast_Migration* node represents sustained, rapid mesenchymal migration of a polarized cell. This requires horizontal polarization, stress fiber maintenance, focal adhesion formation, *PAK1* activity, and high levels of active *Rac1* [259, 260]. |
| | ← Per | Migration | As the *Migration* node represents moderate levels of movement, it's ON state is required for *Fast_Migration*. |
| | ← Ind | FocalAdhesions | Mesenchymal style migration requires force generation via stress fibers anchored by *focal adhesions* [259]. |
| | ← Ind | Stress_Fibers | Mesenchymal style migration requires force generation via stress fibers anchored by *focal adhesions* [259]. |
| | ← Ind | Horizontal_Pol | Directed mesenchymal style migration requires *horizontal cell polarization* with a well defined leading and trailing edge [259]. |

##### Table S1m: Migration_SW module

| | | | |
|---|---|---|---|
| | ← ComplProc | Rac1_H | *Rac1* GTPase signals through *WRC* (WAVE regulatory complex) to drive *Arp2/3*-dependent lamellipodia formation at the leading edge of a migrating cell [238]. |
| | ← ComplProc | PAK1 | Active *PAK1* coordinates a series of cytoskeletal changes at the leading edge that are required for migration and invasion. These include: a) inhibition of myosin light chain kinase in order to decrease contractility if the leading lamellipodium and loss of established actin stress fibers and very strong *focal adhesions* at the leading edge; b) retraction of protrusions and the cell body at the sides and trailing edge with no active *PAK1*; c) suppressing actin filament turnover and promoting leading edge stabilization; d) promote membrane ruffle formation [259, 260]. |

##### Table S1n: EMT module

| Target Node | Node Gate<br>Node Type<br>Link Type | Node Description<br>Input Node | Link Description |
|---|---|---|---|
| SNAI1 | **SNAI1** = (**PAK1** and (((not **GSK3**) or (**NfkB** and **GLI**)) or ((not **miR_34**) and (**ZEB1_H** or **ZEB1**)))) or ((**NfkB** and **GLI** and (not(**miR_34** and **GSK3**))) and (**ZEB1_H** or **ZEB1**)) or (**ZEB1_H** and **ZEB1** and ((not(**miR_34** or **GSK3**)) or (**NfkB** or **PAK1** or **HMGA2** or **HMGA1**))) or (**SMAD2_3_4** and not(**miR_34** or **GSK3**)) or **HMGA2** or **HMGA1** or **GLI_H** | | |
| | TF | *SNAI1*, also known as *Snail*, is a master inducer of EMT, responsible for starting the transcriptional cascade that locks in the EMT regulatory switch [37]. In epithelial cells *SNAI1* is repressed by RNA interference by *miR-34* [261] and protein degradation by *GSK3* [262]. Transcriptional *NF-κB* [263] and *GLI* induction along with loss of these repressive mechanism, and nuclear localization by *PAK1* [264] can activate *SNAI1*, starting EMT. In committed mesenhymal or hybrid E/M cells *ZEB1* lowers the barrier to maitnaining high levels of *SNAI1*. Downstream of *TGFβ*, *SMAD2/3/4*, *HMGA2* or *HMGA1* further aid its transcriptiopnal upregulation of [265]. Finally, high *GLI* activity downstream of *SHH* signaling also activates *SNAI1* [266]. | |
| | ← TR | GLI | *Snail/Snai1* is rapidly induced by *GLI1* [266]. |
| | ← TR | GLI_H | *Snail/Snai1* is rapidly induced by *GLI1* [266]. |
| | ⊢ Deg | GSK3 | *GSK3* both degrades and prevents the transcription of *SNAI1* [262]. |
| | ← PLoc | PAK1 | *PAK1* phosphorylation of *SNAI1* activates and localizes *SNAI1* the nucleus [264]. |
| | ← TR | NfkB | The transcription factor *NF-κB* promotes the expression of *SNAI1* [263], and *NF-κB* inhibition can lower *SNAI1* expression [263]. |

**Table S1n: EMT module**

| | | | |
|---|---|---|---|
| | ← TR | SMAD2_3_4 | Active, nuclear *Smad2/3/4* directly bind and activate the *SNAI1* promoter, though likely requires cooperation from other factors that recruit *HDACs* [267]. |
| | ← TR | HMGA2 | *HMGA2* binds to the *SNAI1* promoter to induce its expression, and cooperates with *Smads* by binding to them and increasing their affinity to the *SNAI1* promoter [265]. Thus, *TGFβ* activates the EMT transciptional program via *HMGA2* [44]. |
| | ← Ind | HMGA1 | siRNA against *HMGA1* strongly reduces *SNAI1* protein expression, though it is unclear whether this is the result of direct transcriptional repression similar to *HMGA2* [268]. |
| | ⊢ RNAi | miR_34 | *SNAI1* mRNA is a direct target of *miR-34* suppression [261]. |
| | ← Ind | ZEB1 | *ZEB1* is an indirect transcriptional inducer of *SNAI1* [269], likely by acting as a competitive inhibitor of microRNAs such as *miR-34* [270]. |
| | ← Ind | ZEB1_H | *ZEB1* is an indirect transcriptional inducer of *SNAI1* [269], likely by acting as a competitive inhibitor of microRNAs such as *miR-34* [270]. |
| LEF1 | **LEF1** = (**ZEB1** and not **miR_200_c**) or **ZEB1_H** or **NfkB** or **SMAD2_3_4** | | |
| | TF | | Lymphoid enhancer-binding factor 1, or *LEF1* is a high-mobility group transcription factor and mediator of *Wnt/β-catenin* signaling. In addition to promotong proliferation, *LEF1* helps induce EMT by activating the transcription *SNAI2* and *ZEB1* [271]. It is induced by *NF-κB* [272] or *Smad2/4* [273], its transcriptional potency is boosted by *ZEB1* [274], and it is targeted for degradation by *miR-200*. |
| | ← TR | NfkB | *LEF1* is a direct transcriptipnal target of *NF-κB* [272]. |
| | ← TR | SMAD2_3_4 | Phosphorilated *Smad2* and *Smad4* can induce *LEF1* gene expression [273]. |
| | ⊢ Ind | miR_200_c | *miR-200s* are indirect repressors of *LEF1*, limiting basal *Pitx2* and *β-catenin* complexes from inducing *LEF1* transcription [275]. Here we assume that medium *ZEB1* availability can aid *LEF1*-mediated transcription if *miR-200c* is repressed, while high levels of *ZEB1* can override *miR-200c*. |
| | ← Compl | ZEB1 | *ZEB1* can bind to and significantly boost *LEF1*-mediated transcription [274]. |
| | ← Compl | ZEB1_H | *ZEB1* can bind to and significantly boost *LEF1*-mediated transcription [274]. |
| Twist | **Twist** = (not **Casp3**) and ((**SNAI1** and (**NfkB** or (not **miR_34**))) or **HMGA2**) | | |
| | TF | | *Twist* is a master trancriptional regulator of the EMT program, induced by *NF-κB* [276] or *HMGA2* [277], stabilized and aided by *SNAI1*, and cleaved during apoptosis by *Caspase 3* [278]. |
| | ← TR | NfkB | Transcription of *Twist* is induced by *NF-κB* [276]. |

**Table S1n: EMT module**

| | | | |
|---|---|---|---|
| | ← TR | HMGA2 | *HMGA2* is a direct transcriptional inducer of *Twist*, and responsible for its increase in response to *TGFβ* [277]. |
| | ← Ind | SNAI1 | *SNAI1* is required a rapid inrease in *Twist* protein levels, and aids its subsequent transcription in response to *TGFβ* [279]. In addition, *SNAI1* potentiates *Twist*-mediated enhancer activation [280]. |
| | ⊢ RNAi | miR_34 | *Twist*'s 3'UTR is a direct target of *miR-34* [281]. |
| | ⊢ Lysis | Casp3 | *Twist* is a direct proteolytic target of *Caspase 3* [278]. |
| SNAI2 | **SNAI2** = (**Twist** and ((**SNAI2** or **N_bcatenin**) or (**N_bcatenin_H** and **LEF1**))) or **SMAD2_3_4** or **HMGA2** or **HMGA1** | | |
| | TF | | *SNAI2*, also known as *Slug*, is a prototypical EMT transcription factor that regulates tissue development and tumorigenesis [282]. It is induced / maintained in mesenhymal cells by *Twist* [283], *β-catenin* [284, 282], *LEF1* [285] as well as positive auto-regulation [286]. In response to *TGFβ* it is induced by *Smad3* [287], *HMGA2* [44] or *HMGA1* [288]. |
| | ← TR | N_bcatenin | The *Wnt/β-catenin* pathway promotes *SNAI2* transcription through nuclear *β-catenin* [284, 282]. |
| | ← TR | SMAD2_3_4 | *SMAD3* bind the cis-element in the SNAI2 promoter and recruits myocardin-related transcription factors to activate its transcription [287]. |
| | ← TR | HMGA2 | *HMGA2* is a direct transcriptional inducer of *SNAI2* [44]. |
| | ← TR | LEF1 | *LEF1* is a direct transcriptional inducer of *SNAI2* [285]. |
| | ← TR | SNAI2 | *SNAI2* is able to bind to its own promoter and induce transcription of its own mRNA [286]. |
| | ← TR | Twist | *Twist* is a direct transcriptional inducer of *SNAI2* [283]. |
| | ← Ind | HMGA1 | Loss of *HMGA1* strongly reduces *SNAI2* levels, though it is unclear whether this is the result of direct transcriptional repression [288]. |
| | ← TR | N_bcatenin_H | The *Wnt/β-catenin* pathway promotes *SNAI2* transcription through nuclear *β-catenin* [284, 282]. |
| ZEB1 | **ZEB1** = **SNAI2** or (**Twist** and **SMAD2_3_4**) or (**b_catenin_TCF4** and not(**miR_200_b** or **miR_200_c**)) or **GLI** or not **GLI3R** | | |

**Table S1n: EMT module**

| | | | |
|---|---|---|---|
| | TF | Zinc finger E-box binding homeobox 1 or *ZEB1* is one of the core regulators of the EMT transcriptional switch [289]. It is induced by *SNAI2* [290], nuclear *β-catenin/TCF4* [291], in the absence of the epithelial *miR-200* microRNAs [292, 293], *GLI2*, or the absence of the *GLI* repressor *GLIR*. As *ZEB1* has two distinct activation levels in hybrid E/M cells vs fully mesenchynmal ones [294, 295, 296], we modeled *ZEB1* activity with two nodes; this one represents at least medium *ZEB1* activity (characteristic of hybrid E/M cells and compatible with ongoing *miR-200* expression), while the *ZEB1_H* node representing maximal *ZEB1* activation seen in fully mesenchymal cells. | |
| | ← TR | GLI | *GLI2* is a direct transcriptional inducer of *ZEB1* [297]. |
| | ⊢ TR | GLI3R | As activator and repressor forms of the GLI transcription factors share the same DNA binding motifs, we assume that *GLI* competes with *GLI3R* for access to the GLI binding site on their target genes such as *ZEB1* [60]. |
| | ← TR | SMAD2_3_4 | The type I *TGF-β* receptor (*TβRI*) phosphorylates receptor-bound *Smad2* and *Smad3*, which triggers their nuclear translocation. Together with their activating partner *Smad4*, they aid (and are required for) *Twist*-induced transcription of *Zeb1* [35]. |
| | ← TR | SNAI2 | *SNAI2* promotes *ZEB1* transcription [290]. |
| | ← TR | Twist | *Twist* works together with *Smad2/3/4* active complexes to induce *Zeb1* expression [35]. |
| | ⊢ RNAi | miR_200_b | *miR_200* inhibits mRNA expression of *ZEB1* by targeting its mRNA for destruction [298, 292, 293]. |
| | ⊢ RNAi | miR_200_c | *miR_200* inhibits mRNA expression of *ZEB1* by targeting its mRNA for destruction [298, 292, 293]. |
| | ← TR | b_catenin_TCF4 | Nuclear *β-catenin/TCF4* are direct transcriptional inducers of the *ZEB1* promoter [291]. |
| N_bcatenin_H | **N_bcatenin_H** = **N_bcatenin** and (not **miR_34**) and (not **J_acatenin** or **SMAD2_3_4**) and not((**miR_200_c** or **miR_200_b**) and **GSK3**) and not((**CyclinE** or **CyclinA**) and **GSK3**) | | |
| | TF | The *N_bcatenin_H* node represents maximal nuclear *β-catenin* accumulation. This requires *N_bcatenin* = ON, a lack of *miR-34*, either complete absence of junctions (*J_acatenin* = OFF) or activation by *Smads*, and the lack of joint repression *GSK3* and either *miR-200* or *CyclinE/A*-bound *Cdk2*. | |
| | ⊢ Deg | GSK3 | When *GSK3* is inhibited, unphosphorylated *β-catenin* accumulates, translocates to the nucleus, and promotes transcription [212]. |
| | ⊢ Loc | J_acatenin | Junctional *E-cadherins* recruits/sequesters *β-catenin* to adherens junctions and block *β-catenin* nuclear localization [185]. We assume that cells unable to form any adherens junctions have high nuclear *β-catenin*. |

**Table S1n: EMT module**

| | | | |
|---|---|---|---|
| | ← Per | N_bcatenin | Reaching the *N_bcatenin_H* = ON state requires *N_bcatenin* = ON first. |
| | ← P | SMAD2_3_4 | *TGFβ* stimulation leads to increased binding of *SMAD3/4* with *β-catenin*, which in turn increases its nuclear accumulation and *β-catenin*-mediated transcription [299, 300]. |
| | ⊣ RNAi | miR_34 | *β-catenin*'s 3' UTR is a direct *miR-34* target [301]. |
| | ⊣ RNAi | miR_200_b | *β-catenin*'s 3' UTR is a direct *miR-200* target [302]. |
| | ⊣ RNAi | miR_200_c | *β-catenin*'s 3' UTR is a direct *miR-200* target [302]. |
| | ⊣ P | CyclinE | *cyclin E/Cdk2* phosphorylate *β-catenin* on Ser33, Ser37, Thr41, and Ser45, promoting its rapid proteasomal degradation [303]. |
| | ⊣ P | CyclinA | *cyclin A/Cdk2* phosphorylate *β-catenin* , promoting its rapid degradation [303]. |
| ZEB1_H | **ZEB1_H** = **ZEB1** and **LEF1** and (**SNAI2** or **GLI** or not **miR_200_c**) and (**N_bcatenin_H** or (**GLI_H** and not **GLI3R**) or **b_catenin_TCF4** or **HMGA1**) | | |
| | TF | | The *ZEB1_H* node represents maximal *ZEB1* activation seen in fully mesenchymal cells [294, 295, 296]. In our model this requires medium *ZEB1*, *LEF1* [304], either *SNAI2* [290], *GLI* [297] or the absence of *miR-200c* [292, 293], and one of the following: high nuclear *β-catenin* [291], high *GLI* without the inhibitory *GLI3R* [297, 60], *β-catenin/TCF4* complex activation, or *HMGA1* [305]. |
| | ← TR | GLI | *GLI2* is a direct transcriptional inducer of *ZEB1* [297]. |
| | ← TR | GLI_H | *GLI2* is a direct transcriptional inducer of *ZEB1* [297]. |
| | ⊣ TR | GLI3R | As activator and repressor forms of the GLI transcription factors share the same DNA binding motifs, we assume that *GLI* competes with *GLI3R* for access to the GLI binding site on their target genes such as *ZEB1* [60]. |
| | ← Ind | LEF1 | *LEF1* overexpression lead to a substantial increase in *ZEB1*, indicating that elevated levels of *LEF1* can help push *ZEB1* into its mesenchymal-specific high expression range (*LEF1_H* node) [304]. |
| | ← TR | SNAI2 | *SNAI2* promotes *ZEB1* transcription [290]. |
| | ⊣ RNAi | miR_200_c | *miR-200* reduces mRNA expression of *ZEB1* by targeting its mRNA for destruction [298, 292, 293]. |
| | ← Per | ZEB1 | As the *ZEB1* node represents moderate levels of this trascription factor, *ZEB1_H* = ON requires *ZEB1* = ON. |

**Table S1n: EMT module**

| | | | |
|---|---|---|---|
| | ← TR | N_bcatenin_H | Nuclear *β-catenin/TCF4* are direct transcriptional inducers of the *ZEB1* promoter [291]. We assume that high levels of nuclear *β-catenin* are required for turning on the *ZEB1_H* node. |
| | ← TR | b_catenin_TCF4 | Nuclear *β-catenin/TCF4* are direct transcriptional inducers of the *ZEB1* promoter [291]. We assume that high levels of nuclear *β-catenin* are required for turning on the *ZEB1_H* node. |
| | ← Ind | HMGA1 | *HMGA1* is a transcriptional inducer of *JMJD3*, which in turn directly demethylates the *ZEB1* promoter to increasde its transcription [305]. |
| b_catenin_TCF4 | **b_catenin_TCF4** = **N_bcatenin_H** and **SNAI1** and **SNAI2** and not **miR_200_c** | | |
| | PC | The *b_catenin_TCF4* node represents saturating levels of active nuclear *β-catenin/TCF4* transcriptional activity. In addition to the influences required to accumulate high nuclear *β-catenin*, this node's ON state also requires *SNAI1/2* expression (both factors promote the formation of active *β-catenin/TCF4* transcriptional complexes) [306] and the absence of inhibitory *miR-200c* [302]. | |
| | ← Ind | SNAI1 | *SNAI1/2* promote the formation of active *β-catenin/TCF4* transcriptional complexes [306]. |
| | ← Ind | SNAI2 | *SNAI1/2* promote the formation of active *β-catenin/TCF4* transcriptional complexes [306]. |
| | ← Compl | N_bcatenin_H | Reaching saturating levels of active nuclear *β-catenin/TCF4* complex formation requires maximal nuclear *β-catenin* accumularion (*N_bcatenin_H* = ON). |
| | ⊣ RNAi | miR_200_c | *β-catenin*'s 3' UTR is a direct *miR-200* target [302]. |
| ZEB2 | **ZEB2** = (not **miR_200_c**) and (**NfkB** or **SMAD2_3_4**) | | |
| | TF | Zinc finger E-box binding homeobox 2 or *ZEB2* is a core regulator of the EMT transcriptional switch. *ZEB2* is transcriptionally induced by *NF-κB* and/or *SMAD2/3/4* [307], and blocked by *miR_200c*-mediated mRNA degradation [298]. | |
| | ← TR | NfkB | *NF-κB* is a direct transcriptional inducer of *ZEB2* downstream of textitTNF-*α* or *IL1* signaling [307]. |
| | ← TR | SMAD2_3_4 | *SMAD2/3/4* complexes are direct transcriptional inducers of *ZEB2* downstream of textitTGF-*β* signaling [307]. |
| | ⊣ RNAi | miR_200_c | *miR_200c* inhibits mRNA expression of *ZEB2* by targeting its mRNA for destruction [298]. |
| HMGA1 | **HMGA1** = **ZEB2** or (**b_catenin_TCF4** and **N_bcatenin_H**) | | |

**Table S1n: EMT module**

| | | | |
|---|---|---|---|
| | TF | | *HMGA1* is a high mobility group (HMG) proteininvolved in transcriptional regulation via chromatin remodeling. It is directly induced by $\beta$*-catenin/TCF-4*-mediated transcription [308], and indirectly stabilized by *ZEB2* repression of the the *miR-637* mRNA (which targets its mRNA for degradation) [309]. |
| | ← Ind | ZEB2 | *ZEB2* is a direct transcriptional repressor of the *miR-637* promoter, a microRNA that targets *HMGA1* mRNA for destruction [309]. |
| | ← Ind | N_bcatenin _H | $\beta$*-catenin/TCF-4* complexes are direct transcriptional inducers of *HMGA1* [308]. |
| | ← Ind | b_catenin _TCF4 | $\beta$*-catenin/TCF-4* complexes are direct transcriptional inducers of *HMGA1* [308]. |
| TGFb_secr | **TGFb_secr** = ((**b_catenin_TCF4** or (**GLI_H** and not **GLI3R**)) and not(**miR_200_b** or **miR_200_c**)) and (not **Casp3**) | | |
| | Secr | | The *TGFb_secr* node represents the production and secretion of *TGF*$\beta$ by the modeled cell. *TGF*$\beta$ transcription in mesenchymal cells is driven by nuclear $\beta$*-catenin/TCF4*- or -*GLI2*-mediated transcription, while *TGF*$\beta$ mRNA is targeted for degradation by *miR-200* mircoRNAs [36]. Finally, apoptotic *Caspase* activation blocks the secretory pathway responsible for *TGF*$\beta$ release [59]. |
| | ← TR | GLI_H | *GLI2* is a direct transcriptional inducer of *TGF-beta1* transcription [310]. |
| | ⊣ TR | GLI3R | As activator and repressor forms of the GLI transcription factors share the same DNA binding motifs, we assume that *GLI* competes with *GLI3R* for access to the GLI binding site on their target genes such as *ZEB1* [60]. |
| | ⊣ Ind | miR_200 _b | colorOliveGreenOverexpression of *miR-200* significantly reduces mRNA levels of *TGF*$\beta$, with the strongest effect on *TGF*$\beta$*3* [36]. |
| | ⊣ Ind | miR_200_c | colorOliveGreenOverexpression of *miR-200* significantly reduces mRNA levels of *TGF*$\beta$, with the strongest effect on *TGF*$\beta$*3* [36]. |
| | ← TR | b_catenin _TCF4 | *TGF*$\beta$*3* is directly regulated by activated $\beta$*-catenin* [311], while *SNAI1/2* promote the formation of $\beta$*-catenin/TCF4* complexes, which bind to the *TGF*$\beta$*3* promoter induce its transcription [306]. |
| | ⊣ Ind | Casp3 | During apoptosis, caspases cleave key components vesicular transport between the ER and Golgi, resulting in a complete block in ER-mediated protein secretion [59]. |
| miR_34 | **miR_34** = (not **SNAI1**) or (not(**ZEB1** or **ZEB1_H**)) | | |
| | miR | | *miR-34* is an microRNA expressed in epithelial cells and central to blocking the accumulation of EMT-initiating transcription factors such as *SNAI1* and $\beta$*-catenin* [312]. *SNAI1*, together with *ZEB1*, feed back to repress *miR-34* expression in mesenhymal and hybrid E/M cells [313]. |

**Table S1n: EMT module**

| | | | |
|---|---|---|---|
| | ⊢ TR | SNAI1 | *SNAI1* is a direct inhibitor of *miR-34* transcription [313]. |
| | ⊢ TR | ZEB1 | *ZEB1* is a direct inhibitor of *miR-34* transcription [313]. |
| | ⊢ TR | ZEB1_H | *ZEB1* is a direct inhibitor of *miR-34* transcription [313]. |
| miR_200_b | **miR_200_b** = **p21**andnot((**Twist**and**SNAI1**and(**ZEB1**ornot **c_Myb**))or**ZEB1_H**or **ZEB2**) | | |
| | miR | | The *miR_200_b* node represents the miR-200b microRNA expressed in epithelial cells and central to blocking EMT transcription factors [314]. Its levels are increased by *p21* [315], it is directly induced by *c-Myb* [163] and repressed by *ZEB1/2* [316, 317]. This repression is indirectly supported by *SNAI1* [318, 319] and *Twist* [320]. |
| | ← TR | c_Myb | The proto-oncogene *c-Myb* induces the expression of the *miR-200* family, unless the locus is silenced by DNA methylation [163]. |
| | ⊢ Ind | SNAI1 | *SNAI1* induction reduces the expression of *miR-200* [318, 319]. |
| | ⊢ Ind | Twist | *Twist* upregulation is required to maintain high levels of *ZEB1* (*Twist* is a direct enhancer of *ZEB1*) [320]. Thus, we assume that *ZEB1* and *SNAI1* cannot fully repress the *miR-200* cluster in the absence of *Twist*. In addition, overexpression of *Twist* resulted in DNA methylation of the *miR-200* locus, though this effect is likely the indirect result of *ZEB1*-mediated repression [321]. |
| | ⊢ TR | ZEB1 | *ZEB1* is a direct transcriptional repressor of *miR-200* expression [316, 317]. |
| | ⊢ TR | ZEB1_H | *ZEB1* is a direct transcriptional repressor of *miR-200* expression [316, 317]. |
| | ⊢ TR | ZEB2 | Overexpression of *ZEB2* suppressed the activity of the *miR-200b-200a-429* promoter [322]; *ZEB2* is a direct transcriptional repressor of the *miR-200* cluster [316, 317, 323]. |
| | ← Ind | p21 | *p21* knockdown downregulates several EMT-blockign miRNAs, including *miR-200a*, *miR-200b*, *miR-200c* and the *miR-183-96-182* cluster. This inhibits EMT, migration and invasion [315]. While it is unclear if *p21* is a direct repressor of the *miR-200* cluster, it was shown to bind *ZEB1* and inhibit its transcriptipnal effets, relieving its ability to repress the *miR-183-96-182* cluster [315]. As this cluster, not explicitly accounted for in our model, further inhibits *ZEB1* expression [315], it is possible that *p21* increases *miR-200* levels indirectly via this cluster, directly by blocking *ZEB1*-mediated repression, or both. |
| miR_200_c | **miR_200_c** = **p21** or not(**Twist** and **SNAI1** and (**ZEB1_H** or **ZEB2** or (**ZEB1** and not(**miR_200_c** or **c_Myb**)))) | | |

**Table S1n: EMT module**

| | | |
|---|---|---|
| miR | The *miR_200_c* node represents miR-200c microRNA expressed in epithelial cells and central to blocking EMT transcription factors [314]. Its levels are increaased by *p21* [315], it is directly induced by *c-Myb* [163] and repressed by *ZEB1/2* [316, 317]. This repression is indirectly supported by *SNAI1* [318, 319] and *Twist* [320]. Here we assume that in addition to the absence of *p21* and the aid of *SNAI1* and *Twist*, either *ZEB2* or high levels of *ZEB1* (*ZEB_H* = ON) are required to silence the active *miR-200c* promoter. In contrast, medium *ZEB1* can maintain repression as long as *miR-200c* is silenced and its inducer *c-Myb* is off. | |
| ← TR | c_Myb | The proto-oncogene *c-Myb* induces the expression of the *miR-200* family, unless the locus is silenced by DNA methylation [163]. |
| ⊢ Ind | SNAI1 | *SNAI1* induction reduces the expression of *miR-200* [318, 319]. |
| ⊢ Ind | Twist | *Twist* upregulation is required to maintain high levels of *ZEB1* (*Twist* is a direct enhancer of *ZEB1*) [320]. Thus, we assume that *ZEB1* and *SNAI1* cannot fully repress the *miR-200* cluster in the absence of *Twist*. In addition, overexpression of *Twist* resulted in DNA methylation of the *miR-200* locus, though this effect is likely the indirect result of *ZEB1*-mediated repression [321]. |
| ← Epi | miR_200_c | As the *miR-200* promoter is subject of DNA methylaiton and epigenetic silencing during EMT [321], we assume that it is more difficult to turn off than to maintain its silenced state. |
| ⊢ TR | ZEB1 | *ZEB1* is a direct transcriptional repressor of *miR-200* expression [316, 317]. |
| ⊢ TR | ZEB1_H | *ZEB1* is a direct transcriptional repressor of *miR-200* expression [316, 317]. |
| ⊢ TR | ZEB2 | *ZEB2* is a direct transcriptional repressor of *miR-200* expression [316, 317, 323]. |
| ← Ind | p21 | *p21* knockdown downregulates several EMT-blockign miRNAs, including *miR-200a*, *miR-200b*, *miR-200c* and the *miR-183-96-182* cluster. This inhibits EMT, migration and invasion [315]. While it is unclear if *p21* is a direct repressor of the *miR-200* cluster, it was shown to bind *ZEB1* and inhibit its transcriptipnal effets, relieving its ability to repress the *miR-183-96-182* cluster [315]. As this cluster, not explicitly accounted for in our model, further inhibits *ZEB1* expression [315], it is possible that *p21* increases *miR-200* levels indirectly via this cluster, directly by blocking *ZEB1*-mediated repression, or both. |

| | |
|---|---|
| Ecadherin _mRNA | **Ecadherin_mRNA** = not(((**ZEB1_H** and **ZEB2**) or (not **MCRIP1** and (**ZEB1_H** or **ZEB2**))) and **ZEB1** and **SNAI1** and **SNAI2** and **Twist**) |

#### Table S1n: EMT module

| | | | |
|---|---|---|---|
| | mRNA | The *Ecadherin_mRNA* represents basal *E-cadherin* expression, required to make at least some adherens juncitons with neighbors. This basal expression is only blocked during full EMT by the joint action of *ZEB1* [324], *SNAI1* [325, 298, 313], *SNAI2* [325, 326, 298], *Twist* [327, 313] and *ZEB2*, protected from repression by *MCRIP1* [328]. | |
| | ← Ind | MCRIP1 | *MCRIP1* promotes transcription of *E-cadherin* by competitively binding to *ZEB1* co-repressor *CtBP*, thus inhibiting *ZEB1*-mediated transcriptional repression [328]. During EMT, *MCRIP1* is inhibited by *ERK* phosphorylation [328]. |
| | ⊢ TR | SNAI1 | *SNAI1* is a direct transcriptional inhibitor of *E-cadherin* [325, 298, 313]. |
| | ⊢ TR | SNAI2 | *SNAI2* is a direct transcriptional inhibitor of *E-cadherin* transcription [325, 326, 298]. |
| | ⊢ TR | Twist | *Twist* is a direct transcriptional inhibitor of *E-cadherin* transcription [327, 313]. |
| | ⊢ TR | ZEB1 | *ZEB1* is a direct transcriptional inhibitor of *E-cadherin* [324], either in partnership with its co-repressor *CtBP* [329], or by binding to the SWI/SNF chromatin-remodeling protein *BRG1* [330]. |
| | ⊢ TR | ZEB1_H | *ZEB1* is a direct transcriptional inhibitor of *E-cadherin* [324], either in partnership with its co-repressor *CtBP* [329], or by binding to the SWI/SNF chromatin-remodeling protein *BRG1* [330]. |
| | ⊢ TR | ZEB2 | *ZEB2* is a direct transcriptional inhibitor of *E-cadherin* [331]. |

#### Table S1o: Restriction_SW module

| Target Node | Node Gate<br>Node Type<br>Link Type | Node Description<br>Input Node | Link Description |
|---|---|---|---|
| p21 | **p21** = (**p21_mRNA** and (not **Casp3**)) and (not **CyclinE** or not **Myc**) | | |
| | Prot | In this model, the *p21* node corresponds to nuclear p21 in cells with relatively high basal *p21* activity. *p21* activity and or localization can be lowered by loss of *FoxO* mediated transcription (see $p21_{mRNA}$ node) and via feedback from *Cyclin E/Cdk2* [332] aided by ongoing *Myc*-mediated repression [333]. | |
| | ← TL | p21 _mRNA | *p21* protein activity requires the presence of *p21* transcription. |
| | ← TR | Myc | Induced *Myc* expression lead to decreased endogenous *p21* through *Myc* binding to its proximal promoter, preventing transcription. *Myc* is also suspected to sequester the *p21* activator *Sp1* [333]. |

**Table S1o: Restriction_SW module**

| | | | |
|---|---|---|---|
| | $\dashv$ Deg | CyclinE | *p21* and *Cyclin E/Cdk2* form a positive (double-negative) feedback loop in which *Cyclin E/Cdk2* activates the *SCF/Skp2* complex responsible for the degradation of *Cyclin E/Cdk2*-bound, phosphorylated *p21* [334]. *p21*, in turn, not only blocks *Cyclin E/Cdk2* activity, but it also inhibits *Cyclin D1*. Thus, *p21* interferes with the mitogen signal that turns on *Cyclin E* in the first place. In quiescent cells with high basal *p21* levels, this positive feedback renders cell cycle entry stochastic [332]. |
| | $\dashv$ Lysis | Casp3 | *Caspase 3* cleaves and deactivates *p21* [335]. |
| pRB | **pRB** = (((not **Casp3**) and (not **CyclinD1**)) and (not **CyclinA**)) and (**p27Kip1** or (not **CyclinE**)) | | |
| | TF | *pRB* is active in the absence of *Caspase 3*, *Cyclin D1*, *Cyclin A*, and *Cyclin E*. In addition, *pRB* maintains its activity when active $p27^{Kip1}$ counteracts the effects of *Cyclin E* [336, 337, 338, 339]. | |
| | $\leftarrow$ ComplProc | p27Kip1 | Active $p27^{Kip1}$ can counteract the inhibitory effects of active *CyclinE/Cdk2* complexes [336]. |
| | $\dashv$ P | CyclinD1 | *Cyclin D1/Cdk4,6* complexes bind and phosphorylate *RB*, inhibiting its activity [340, 341, 338]. |
| | $\dashv$ P | CyclinE | *Cyclin E/Cdk2* complexes bind and phosphorylate *RB*, inhibiting its activity [342, 338]. |
| | $\dashv$ P | CyclinA | *Cyclin A/Cdk1,2* complexes phosphorylate and deactivate *RB* [337]. |
| | $\dashv$ Lysis | Casp3 | *Caspase 3* cleaves *RB*, generating fragments that do not associate with *E2F1*, rendering *RB* inactive [343]. |
| p27Kip1 | **p27Kip1** = (((not **Casp3**) and (not **CyclinD1**)) and (not(**Cdk1** and **CyclinB**))) and ((((not((**CyclinA** and **Necl5**) and **CyclinE**)) and (**FoxO3** and **FoxO1**)) or (((not **CyclinA**) or (not(**Necl5** or **CyclinE**))) and (**FoxO3** or **FoxO1**))) or ((not **CyclinA**) and (not(**Necl5** and **CyclinE**)))) | | |
| | Prot | Active $p27^{Kip1}$ is cleaved by *Caspase 3* and inhibited (sequestered) by *Cyclin D1/Cdk4,6* [336] or *Cyclin B/Cdk1* [344]. In addition, maintenance of $p27^{Kip1}$ requires one or both *FoxO* factors when sequestered by *Cyclin E/Cdk2* (one *FoxO* factor) or *Cyclin A/Cdk2* (both *FoxO* factors), but it cannot keep pace with the simultaneous activity of *Cyclin E/Cdk2* and *Cyclin A/Cdk2* [345]. | |
| | $\leftarrow$ TR | FoxO3 | *FoxO* factors are direct inducers of $p27^{Kip1}$ expression [346]. |
| | $\leftarrow$ TR | FoxO1 | *FoxO* factors are direct inducers of $p27^{Kip1}$ expression [346]. |
| | $\dashv$ TR | Necl5 | *Necl5* downregulates the transcription of *p27Kip1* in response to growth factor stimulation [347]. |
| | $\dashv$ IBind | CyclinD1 | Active *Cyclin D/Cdk4,6* complexes competitively bind to *p27Kip1* and progressively inhibit its ability to keep *Cyclin-E/Cdk2* inactive, thereby inducing cdk2 activity and cell-cycle progression [336]. |

**Table S1o: Restriction_SW module**

| | | | |
|---|---|---|---|
| | $\vdash$ Deg | CyclinE | Active *Cyclin-E/Cdk2* phosphorylate *p27Kip1* at threonine 187 (Thr187) [348], which marks it for degradation by the $SCF^{\mathrm{SKP2}}$ complex at the onset of S-phase [349]. (*Cyclin-E/Cdk2* complexes remain active in the presence of *p27Kip1* and promote its degradation when *Cyclin-A* is also active.) |
| | $\vdash$ Deg | CyclinA | *Cyclin A/Cdk2* complexes bind and inactivate $p27^{Kip1}$ by sequestration, phosphorylate it, and promote its degradation [350]. |
| | $\vdash$ PLoc | CyclinB | *Cyclin B/Cdk1* complexes phosphorylate $p27^{Kip1}$ [350], and although they do not promote its degradation, phosphorylated $p27^{Kip1}$ is exported from the nuclear compartment and looses its ability to inhibit *Cdk* activity [344]. |
| | $\vdash$ PLoc | Cdk1 | *Cyclin B/Cdk1* complexes phosphorylate $p27^{Kip1}$ [350], and although they do not promote its degradation, phosphorylated $p27^{Kip1}$ is exported from the nuclear compartment and looses its ability to inhibit *Cdk* activity [344]. |
| | $\vdash$ Lysis | Casp3 | *Caspase 3* cleaves *p27Kip1* [351]; the cleaved fragments can no longer associate with *Cdk2 / Cyclin* complexes [352]. |
| Myc | **Myc** = (((**ERK** and **YAP**) and (not **SMAD2_3_4**)) or ((**ERK** or (**YAP** and (not **SMAD2_3_4**))) and (**eIF4E** and (not **GSK3**)))) or ((**E2F1** and (not **pRB**)) and ((**eIF4E** or **ERK**) or (not **GSK3**))) | | |
| | TF | | *Myc* activity is turned on by stabilization of the protein via *ERK* phosphorylation, aided by *YAP*-mediated transcription [353, 354] in the absence of inhibitory *SMADs* [355]. To take into account theincrease in translation initiated by *eIF4E* and loss of degradation-promoting phosphorylation when *GSK3β* is off [356], we assumed that they can compensate for the lack of *ERK*, absence of *YAP*, or interference from *SMADs*. Alternatively, increased transcription by *E2F1* can also promote *Myc* accumulation in the absence of active *pRB* [357], provided that the protein is stabilized by *ERK*, *eIF4E*, or the absence of *GSK3β*. |
| | $\leftarrow$ P | ERK | Ser-62 phosphorylation by *ERK* increases its half life, leading to *Myc* accumulation [356, 358]. |
| | $\leftarrow$ TR | eIF4E | Increased translational initiation in the presence of activated *eIF4E* leads to an increase in *Myc* protein levels [359]. |
| | $\vdash$ P | GSK3 | Thr-58 phosphorylation by *GSK-3* promotes *Myc* degradation [356, 360]. |
| | $\leftarrow$ TR | YAP | *YAP* is a transcriptional inducer of /textitc-Myc [353, 354]. |
| | $\vdash$ TR | SMAD2_3_4 | *TGFβ* stimulation induces *Smad* complex formation that recognizes a *TGFβ* inhibitory element in the *c-Myc* promoter [355]. This inhibition may be partially rescued by *YAP*, as *YAP/Smad7* complexes were shown to interfere with *Smad*-dependent gene expterssion control [361]. |
| | $\vdash$ TR | pRB | *E2F1*'s ability to induce *Myc* is blocked by active (hypo-phosphorylated *pRB*) [362]. |

**Table S1o: Restriction_SW module**

| | | | |
|---|---|---|---|
| | ← TR | E2F1 | *E2F1* binds and activates the *c-Myc* promoter [363, 364]. |
| CyclinD1 | **CyclinD1** = ((not **p15**) and (not **CHK1**)) and (((not **p21**) and ((((not **GSK3**) and **YAP**) and (**Myc** or **E2F1**)) or (((**CyclinD1** and **YAP**) and (**Myc** or **E2F1**)) or (**Myc** and **E2F1**)))) or (((not **pRB**) and **E2F1**) and (((**Myc** and **CyclinD1**) or (**Myc** and (not **GSK3**))) or ((**YAP** and **CyclinD1**) and (not **GSK3**))))) | | |
| | PC | Ongoing DNA synthesis keeps the *CHK1* kinase active, which inhibits *Cyclin D1*. Similarly, the CDKI *p15* also binds and blocks active *Cdk/Cyclin D* complexes. The precise regulatory logic of *Cyclin D1* as a function of transcriptional control by *Myc* and *E2F1*, combined with the regulation of its protein stability / activity by *GSK3β* / basal *p21* is not known. Here, we assume that in the absence of *p21* (once *p21* levels drop due to growth factor signals and/or *Cdk2* activation), *Cyclin D1* can be activated by *YAP* and either *Myc* or *E2F1* – as long as *GSK3β is OFF*. In the presence of *GSK3β*, we assume that *Cyclin D1* can be induced by the combined action of both *Myc* and *E2F1* [365], but sustained in an ON state by either. In the presence of basal (normal quiescent) levels of *p21*, we assume that *Cyclin D1* transcription requires *E2F1* unencumbered by *pRB*, as well as any two of the following: *Myc*, already active *Cyclin D1*, sustained by *YAP* and not blocked by *GSK3β*. | |
| | ⊢ P | GSK3 | *GSK-3β* phosphorylates *Cyclin D1* on Thr-286, promoting its ubiquitination and degradation [366]. |
| | ← TR | YAP | *YAP* is a direct transcriptional inducer of *Cyclin D* [367]. |
| | ⊢ IBind | p15 | *p15* is a cyclin-dependent kinase inhibitor of *cdk4 / cdk6*. As it displaces *Cyclin D*, it blocks G1/S progression [45]. |
| | ⊢ IBind | p21 | *p21*$^{Cip1}$ is a Cyclin Dependent kinase inhibitor which binds to and blocks the activity of *Cdk2*, *Cdk3*, *Cdk4* and *Cdk6* kinases [368] and thus inhibits *CyclinD1/Cdk4,6* [369]. |
| | ⊢ TR | pRB | *E2F1*'s ability to induce *Cyclin D1* is blocked by active (hypo-phosphorylated) *RB* protein [370]. |
| | ← TR | Myc | Extracellular growth signals activate the MAPK pathway, leading to transcriptional activation of *Cyclin D1* by *Myc* [371, 372]. *Myc* overexpression leads to rapid *Cyclin D1* induction and subsequent cell cycle entry [373], while its absence halves *Cyclin D1* levels [374]. In addition, *Myc* induces *Cdk4*, aiding the assembly of active *Cyclin D1 / Cdk4,6* complexes [374, 375]. |
| | ← Per | CyclinD1 | In order to take into account both production and stability of *Cyclin D1*, we assumed that the presence of active *CyclinD/Cdk2,4* complexes renders transcriptional maintenance of their levels easier. |

**Table S1o: Restriction_SW module**

| | | | |
|---|---|---|---|
| | ← TR | E2F1 | The *Cyclin D1* promoter is bound by *E2F* factors including *E2F1* [376], and *E2F1* overexpression can increase *Cyclin D1* (though its effects are context-dependent, as *E2F1* overexpression can also lead to apoptosis) [376]. Dominant negative *E2F1* overexpression results in a 2-3 fold decrease in *Cyclin D* expression and *Cyclin D/Cdk4,6* activity [377]. |
| | ⊣ P | CHK1 | During replication, checkpoint kinases such as *CHK1* (active during normal DNA synthesis) suppress *Cyclin D1* [378], which has a very short half-life (∼ 24 min) [379]. |
| E2F1 | **E2F1** = (not((**CAD** or **CyclinA**) or **pRB**)) and ((**YAP** and (**E2F1** or **Myc**)) or (**E2F1** and **Myc**)) | | |
| | TF | In the absence of both *CyclinA* and *pRB*, *E2F1* transcription can be induced by *YAP* and *Myc* or maintained by active *E2F1*. *CAD* deactivates *E2F1* as it destroys the cell's DNA. | |
| | ← TR | YAP | *YAP* is a direct transcriptional inducer of *E2F1* [380]. |
| | ⊣ TR | pRB | *RB* binds to *E2F/DP1* complexes and switches their DNA binding activity from activation to repression [370, 381]. |
| | ← TR | Myc | *Myc* is required for growth-factor mediated induction of *E2F1* [382, 374]. It binds to and remodels the *E2F1* promoter, facilitating *E2F1* transcription [365]. In addition, *Myc* augments protein expression of *E2F1* [383]. Single-cell experiments show that *Myc* is a critical modulator of the amplitude of *E2F* activation [384]. |
| | ← TR | E2F1 | *E2F1* binds to its own promoter and up regulates transcription (as long as *Cyclin D/E* activity blocks *RB-E2F1* binding) [385]. |
| | ⊣ P | CyclinA | The phosphorylation of the *E2F1*-binding *DP-1* protein by *Cyclin A*, which binds directly to *E2F-1* (as well as *E2F-2,3*) downregulates *E2F1* transcriptional activity in S phase [386, 387, 388]. |
| | ⊣ Deg | CAD | This link from Caspase-activated DNase (*CAD*) to *E2F1* ensures that apoptotic cells settle into an *E2F1*-negative attractor regardless of their initial state. The rationale for this is that *E2F1* cannot maintain its activity if DNA is fragmented. |
| CyclinE | **CyclinE** = **E2F1** and **Cdc6** and **Pre_RC** and not(**pRB** or (**p27Kip1** and not **CyclinA**) or **CHK1** or **Casp3**) | | |
| | PC | In our model, the ON state of *Cyclin E* represents active *Cyclin E/Cdk2* complexes. Thus, its full activation requires transcription via *E2F1* not blocked by active *pRB*, binding to *Cdc6* and *pre-RC* complexes, and the absence of its inhibitors $p27^{Kip1}$ in the absence of *Cyclin A/Cdk2* binding partners, *CHK1* and *Caspase 3*. | |
| | ⊣ TR | pRB | *Cyclin E* transcription by *E2F1* requires the absence of active, un-phosphorylated *RB* [388]. |

**Table S1o: Restriction_SW module**

| | | |
|---|---|---|
| ⊣ IBind | p27Kip1 | *p27Kip1* binds to and prevents the activation of *Cyclin E/Cdk2* complexes [336]. |
| ← TR | E2F1 | *E2F1* is a potent transcriptional activator of *Cyclin E* [389]. |
| ← Compl | Cdc6 | Chromatin association and full activation of *Cyclin E/Cdk2* requires *Cdc6* [390]. |
| ← Compl | Pre_RC | At the G1/S transition, *Cyclin E* is loaded onto chromatin by *pre-RC* complexes (*Cdc6* and *Cdt1* binding), where it is required for *MCM2* loading, origin firing and the start of DNA synthesis [391]. In addition, activation of its partner *Cdk2* by *Cdc6* is contingent on this localization [390]. |
| ⊣ Ind | CyclinA | *Cyclin A/Cdk2* complexes bind and inactivate $p27^{Kip1}$ by sequestration, phosphorylate it, and promote its degradation [350]. Here we assume that full potency of $p27^{Kip1}$ requires a lack of *Cyclin A/Cdk* activity. |
| ⊣ P | CHK1 | *Chk1* activation during normal S-phase progression keeps *Cdk2* activity in a physiological range by binding to both *Cdk2* and *Cdc25A*, aiding the loss of *Cyclin E/Cdk1* activity [392]. |
| ⊣ Lysis | Casp3 | *Caspase 3* cleaves and deactivates *Cyclin E*, which is then rapidly degraded [393]. |

**Table S1p: Origin_Licensing module**

| Target Node | Node Gate<br>Node Type<br>Link Type | Node Description<br>Input Node | Link Description |
|---|---|---|---|
| ORC | **ORC** = **E2F1** or ((**Pre_RC** and **Cdt1**) and **Cdc6**) | | |
| | PC | *ORC* proteins can bind at origins of replication when transcribed by *E2F1* or as part of a fully assembled and licensed *Pre-RC* complex (including active *Cdc6* and *Cdt1*). | |
| | ← TR | E2F1 | Expression of the *ORC1* gene is regulated by *E2F1* [394]. |
| | ← Compl | Cdc6 | Availability of stable (unphosphorylated) *Cdc6* in the *Pre-RC* is necessary for the maintenance of licensed origins [395]. |
| | ← Compl | Cdt1 | Active (unphosphorylated and not geminin-bound) *Cdt1* bound to the *Pre-RC* is necessary for the maintenance of licensed origins [395]. |
| | ← Compl | Pre_RC | Licensed but not yet fired replication complexes (*Pre-RC*s containing *ORC*, *Cdc6*, *Cdt1* and inactive *MCM*s) remain stable at sites of replication origin until fired by the activation of the *MCM* helicase [395]. |

#### Table S1p: Origin_Licensing module

Cdc6

**Cdc6** = ((not **Casp3**) and (not(**f4N_DNA** and **CyclinA**))) and (((**E2F1** and **ORC**) and (not **Plk1**)) or (((**Pre_RC** and **ORC**) and **Cdc6**) and **Cdt1**))

Prot — In our model the *Cdc6* node represents nuclear, chromatin-bound *Cdc6.* Thus, the node is only active during the assembly of pre-replication complexes, or their ongoing presence during DNA replication. *Cdc6* is ON in the absence of *Caspase 3* or *CyclinA / Cdk2* phosphorylation of *Cdc6* in all origins required for the completion of DNA replication (thus, its inhibition by *Cyclin A* also requires 4N DNA). In addition, active *Cdc6* requires either transcription by *E2F1* and recruitment by origin-bound *ORC* proteins in the absence of mitotic *Plk1* or maintenance of *Pre-RCs* by the presence of all of its components.

| | | |
|---|---|---|
| $\leftarrow$ TR | E2F1 | Transcription of Cdc6 is directly induced by E2F1 [396]. |
| $\leftarrow$ Compl | ORC | *ORC* recruits *Cdc6* to origins of replication [395]. |
| $\leftarrow$ Per | Cdc6 | Stable (unphosphorylated) *Cdc6* in the *Pre-RC* is necessary for the maintenance of licensed origins [395]. |
| $\leftarrow$ Compl | Cdt1 | Active (unphosphorylated and not geminin-bound) *Cdt1* bound to the *Pre-RC* is necessary for the maintenance of licensed origins [395]. |
| $\leftarrow$ Compl | Pre_RC | Licensed but not yet fired replication complexes (*Pre-RC*s containing *ORC*, *Cdc6*, *Cdt1* and inactive *MCM*s) remain stable and *Cdc6*-bound until fired by the activation of the *MCM* helicase [395]. |
| $\dashv$ P | Plk1 | *Plk1* binds, phosphorylated and strongly recruits Cdc6 to the spindle pole during metaphase, then to the central spindle in anaphase, leading to its exclusion from chromosomes until telophase, when the majority of *Plk1* is degraded by $APC/C^{Cdh1}$ [397]. |
| $\dashv$ P | CyclinA | Phosphorylation of *CDC6* by *Cyclin A/Cdk2* during DNA replication leads to its re-localization to the cytoplasm [398]. |
| $\dashv$ Ind | f4N_DNA | In our model, full deactivation of Cdc6 represents the firing of all ORCs as DNA replication is completed. Thus *Cyclin A*'s inhibitory action takes full effect once the cell reaches 4N DNA content [398]. |
| $\dashv$ Lysis | Casp3 | *Caspase 3* cleaves and deactivates *Cdc6* [399]. |

Cdt1

**Cdt1** = ((((not **geminin**) and **ORC**) and **Cdc6**) and (not((**CyclinE** and **CyclinA**) and **Cdc25A**))) and ((**Pre_RC** and (**E2F1** or **Myc**)) or (**E2F1** and (**Myc** or (not **pRB**))))

**Table S1p: Origin_Licensing module**

| | | | |
|---|---|---|---|
| | Prot | Replication-origin bound *Cdt1* requires the absence of *geminin*, the presence of origin-bound *ORC* and *Cdc6*, and the absence of sustained *Cdk2* activity responsible for the firing of all origins during DNA synthesis (modeled as simultaneous *Cyclin E*, *Cyclin A* and *Cdc25A* activity). Bound into a licensed pre-replication complex (*Pre-RC*), *Cdt1* remains stable as long as it is transcribed by *E2F1* [400] or *Myc* [401] (this guarantees that *Pre-RC* complexes cannot persist indefinitely in the absence of de novo transcription). Alternatively, it can be turned on by *E2F1*, aided by *Myc* or the absence of *RB*, and *FoxO3* in cells with 4N DNA. | |
| | ⊢ TR | pRB | *E2F1*-mediated transcription of *Cdt1* is blocked by hypophosphorylated (active) *pRB* [400]. |
| | ← TR | Myc | *Cdt1* is a direct transcriptional target of the *Myc-Max* complex [401], ensuring its availability for *Pre-RC* formation and maintenance. |
| | ← TR | E2F1 | *Cdt1* is a direct transcriptional target of *E2F1* [400], ensuring its availability for *Pre-RC* formation and maintenance. |
| | ⊢ P | CyclinE | Sustained *Cdk2* activity during S-phase (modeled as simultaneous *Cyclin E*, *Cyclin A* and *Cdc25A* activity) is responsible for the firing of all origins required to complete DNA synthesis; it also leads to the phosphorylation and proteasomal degradation of *Cdt1* [395]. |
| | ← Compl | ORC | *ORC*-bound origin of replication sites are the point of pre-replication complex assembly, where *Cdt1* is recruited by *ORC*-bound *Cdc6* [395]. |
| | ← Compl | Cdc6 | *ORC*-bound *Cdc6* recruits *Cdt1* to *pre-RC* complexes [395]. |
| | ← Compl | Pre_RC | Licensed but not yet fired replication complexes (*Pre-RC*) remain stable until fired during DNA replication [395]. |
| | ⊢ IBind | geminin | *Geminin* binds to *Cdt1* at pre-replication complexes, where it blocks *Cdt1* binding to DNA, sequestering it away from *Pre-RC*s [402]. |
| | ⊢ P | Cdc25A | Sustained *Cdk2* activity leads to phosphorylation and degradation of *Cdt1* [395]. |
| | ⊢ P | CyclinA | Sustained *Cdk2* activity leads to phosphorylation and degradation of *Cdt1* [395]. |
| Pre_RC | **Pre_RC** = ((**ORC** and **Cdc6**) and **Cdt1**) and (not(**Replication** and **f4N_DNA**)) | | |
| | PC | *Pre-RC* complexes assemble when *ORC*, *Cdc6*, and *Cdt1* are all bound to sites of replication origin along the DNA. The node denoting their licensing turns OFF at the moment of transition from ongoing *Replication* to *f4N_DNA* (it is blocked in the one time-point when both of these nodes are ON). | |
| | ← Compl | ORC | *Pre-RC* complexes assemble when *ORC*, *Cdc6*, and *Cdt1* are all bound to sites of replication origin along the DNA [395]. |
| | ← Compl | Cdc6 | *Pre-RC* complexes assemble when *ORC*, *Cdc6*, and *Cdt1* are all bound to sites of replication origin along the DNA [395]. |

**Table S1p: Origin_Licensing module**

| | | | |
|---|---|---|---|
| | ← Compl | Cdt1 | *Pre-RC* complexes assemble when *ORC*, *Cdc6*, and *Cdt1* are all bound to sites of replication origin along the DNA, leading to the recruitment of the *MCM* helicase [395]. |
| | ⊢ Unbind | Replication | *Pre-RC*s fire and fall apart during DNA replication [395]. |
| | ⊢ Ind | f4N_DNA | In our model the *Pre-RC* node turns OFF when *Replication* is completed, marked by the time-point when both *Replication* and *f4N_DNA* are ON. |
| geminin | **geminin** = (**E2F1** and (not **Cdh1**)) and (not(**pAPC** and **Cdc20**)) | | |
| | Prot | *Geminin* is present when transcribed by *E2F1* and not targeted for degradation by $APC/C^{Cdh1}$ or $APC/C^{Cdc20}$ [403]. | |
| | ← TR | E2F1 | *Geminin* is a direct transcriptional target of *E2F1* [400]. |
| | ⊢ Ubiq | pAPC | *Geminin* is a target of $APC/C^{Cdc20}$ at the metaphase/ anaphase transition [403]. |
| | ⊢ Ubiq | Cdc20 | *Geminin* is a target of $APC/C^{Cdc20}$ at the metaphase/ anaphase transition [403]. |
| | ⊢ Ubiq | Cdh1 | *Geminin* is a target of $APC/C^{Cdh1}$ ubiquitin ligase [404]. |

**Table S1q: Phase_SW module**

| Target Node | Node Gate<br>Node Type<br>Link Type | <br>Node Description<br>Input Node | <br><br>Link Description |
|---|---|---|---|
| CyclinA _mRNA | **CyclinA_mRNA** = (not **CAD**) and ((**E2F1** and (not **pRB**)) or **FoxM1**) | | |
| | mRNA | In non-apoptotic cells (no *CAD*), *Cyclin A* is transcribed by *E2F1* in the absence of active *RB* or by *FoxM1*. | |
| | ⊢ TR | pRB | Active *RB* blocks *E2F1*'s ability to transcribe *Cyclin A* [405]. |
| | ← TR | E2F1 | *Cyclin A* is transcriptionally activated by *E2F* factors [388]. |
| | ← TR | FoxM1 | Depletion of *FoxM1* results in reduced *Cyclin A2* expression (it is not clear whether *FoxM1* is a direct transcriptional inducer of *Cyclin A*) [406, 407, 408, 409]. |
| | ⊢ Ind | CAD | This link from Caspase-activated DNase (*CAD*) to *Cyclin A* mRNA ensures that apoptotic cells settle into a G0-like attractor regardless of their initial state. The rationale for this is that no mRNA synthesis can be maintained if DNA is fragmented (we only use these links from *CAD* if needed). |

**Table S1q: Phase_SW module**

| | | | |
|---|---|---|---|
| Emi1 | **Emi1** = ((**E2F1** or (not **pRB**)) or (not **p21**)) and (not(((**Plk1** and **CyclinB**) and **Cdk1**) and (**U_Kinetochores** or **A_Kinetochores**))) | | |
| | Prot | Our model allows the sustained presence of *Emi1* protein when it is either actively transcribed by *E2F1* [410, 411] or lacks joint inhibition by *pRB* [411] and *p21* [412]. Degradation of *Emi1* is mediated by *Plk1* and *CyclinB/Cdk1* complexes; initiation of this degradation requires at least temporary co-localization of *Emi1* with *Plk1* at mitotic spindle poles [413]. | |
| | ⊣ Ind | p21 | *p21* activation during DNA damage lead to a substantial decrease of *Emi1* levels, not observed in *p21*-null cells [412]. |
| | ⊣ TR | pRB | Active retinoblastoma protein can block *Emi1* transcription mediated by *E2F1* [411]. |
| | ← TR | E2F1 | *Emi1* is a direct transactional target of *E2F1* [410, 411]. |
| | ⊣ P | Plk1 | *Plk1* phosphorylates *Emi1* at mitotic spindle poles, stimulating its *βTrCP* binding and ubiquitination [413]. |
| | ⊣ Ind | CyclinB | *Cyclin B/Cdk1* enhances the ability of *Plk1* to mediate *Emi1* destruction [413]. |
| | ⊣ Ind | Cdk1 | *Cyclin B/Cdk1* enhances the ability of *Plk1* to mediate *Emi1* destruction [413]. |
| | ⊣ Ind | U_Kinetochores | As *Plk1*-mediated phosphorylation of *Emi1* occurs at mitotic spindle poles, our model requires ongoing mitosis for this interaction [413]. |
| | ⊣ Ind | A_Kinetochores | *Plk1* phosphorylates *Emi1* at mitotic spindle poles, stimulating its *βTrCP* binding and ubiquitination [413]. |

| | | | |
|---|---|---|---|
| FoxM1 | **FoxM1** = (((**Myc** or **YAP**) and **CyclinE**) or ((**CyclinA** and **Cdc25A**) and **Cdc25B**)) or ((**Plk1** and **CyclinB**) and **Cdk1**) | | |
| | TF | In our model, *FoxM1* activity requires increased expression by *Myc* [414] or *YAP* [367] and activating phosphorylation by *Cyclin E/Cdk2*. Alternatively, *FoxM1* activity can be sustained by potent *Cdk2 / Cdk1* activity in G2 (supported by *Cdc25A* or *Cdc25B*), or a serial phosphorylation by *Cyclin B/Cdk1* and *Plk1* during mitosis. | |
| | ← TR | YAP | *FoxM1* is a direct transcriptional target of *YAP* [367]. |
| | ← TR | Myc | *FoxM1* is a direct transcriptional target of *c-Myc* [414]. |
| | ← P | CyclinE | *Cyclin E/Cdk2* complexes bind and phosphorylate FoxM1, potently inducing its transcriptional activity, which starts during S-phase [415]. |
| | ← Compl | Cdc25A | Active *Cdc25A* binds to and enhances the transcriptional activity of *FoxM1*, potentually by bridging FoxM1 and active cyclin-*Cdk2* complexes [416]. |
| | ← Ind | Cdc25B | *Cdc25B* overexpression can increase *FoxM1*-dependent transcription, likely via aiding *Cdk1* activity [417]. |

**Table S1q: Phase_SW module**

| | | | |
|---|---|---|---|
| | ← P | Plk1 | *Plk1* binds and phosphorylates *FoxM1*, which activates *FoxM1*-mediated transcription in early mitosis [418]. |
| | ← P | CyclinA | In addition to *Cyclin E/Cdk2*, *Cyclin A/Cdk2* complexes can also keep *FoxM1* transcriptionally active by phosphorylating its autoinhibitory N-terminal region [419]. |
| | ← P | CyclinB | *FoxM1* binds *Plk1*, and phosphorylation of two key residues at this binding domain by *Cyclin B/Cdk1* primes it for *Plk1* binding [418]. |
| | ← P | Cdk1 | *FoxM1* binds *Plk1*, and phosphorylation of two key residues at this binding domain by *Cyclin B/Cdk1* primes it for *Plk1* binding [418]. |
| Cdc25A | **Cdc25A** = (((**FoxM1** and **E2F1**) and (not **pRB**)) or ((not **Cdh1**) and (**FoxM1** or (**E2F1** and (not **pRB**))))) and ((((not(**GSK3** or **CHK1**)) or **CyclinE**) or **CyclinA**) or (**CyclinB** and **Cdk1**)) | | |
| | Ph | | As the precise combinatorial regulation of *Cdc25A* throughput the cell cycle is unknown, our model assumes that accumulation of the *Cdc25A* protein requires transcriptional activation by both *E2F1* in the absence of *pRB*, and *FoxM1* to override destruction by $APC/C^{Cdh1}$. Alternatively, one of the two transcription factors can drive *Cdc25A* accumulation in the absence of $APC/C^{Cdh1}$. In addition, stabilization of *Cdc25A* either requires the absence of *GSK3β* and *CHK1* (both of which promote its degradation), or stabilization by *Cdk* activity. |
| | ⊢ P | GSK3 | *GSK3β* phosphorylates *Cdc25A*, promoting its proteolysis [420]. |
| | ⊢ TR | pRB | Active (hypo-phosphorylated) *pRB* blocks *E2F1*'s ability to drive *Cdc25A* transcription [421, 422]. |
| | ← TR | E2F1 | *E2F1* is a direct transcriptional inducer of *Cdc25A* [421]. |
| | ← P | CyclinE | *Cdc25A* protein levels are stabilized during S-phase by *CyclinE/Cdk2*-dependent phosphorylation [423]. |
| | ← TR | FoxM1 | *FoxM1* is a direct transcriptional inducer of *Cdc25A* [416]. |
| | ← P | CyclinA | *Cdc25A* protein levels are stabilized during S and G2 by *Cdk2*-dependent phosphorylation. *Cdk2* first partners with *Cyclin E* [423], then continues to stabilize *Cdc25A* past the point of *Cyclin E* expression by partnering with *Cyclin A* [424]. |
| | ← P | CyclinB | During mitosis, *Cdc25A* is stabilizes by *Cyclin B/Cdk1* phosphorylation, which protects it from the proteasome [425]. |
| | ← P | Cdk1 | During mitosis, *Cdc25A* is stabilizes by *Cyclin B/Cdk1* phosphorylation, which protects it from the proteasome [425]. |
| | ⊢ Deg | Cdh1 | The $APC/C^{Cdh1}$ complex degrades *Cdc25A* at mitotic exit [426, 427]. |

**Table S1q: Phase_SW module**

| | | | |
|---|---|---|---|
| | ⊣ P | CHK1 | *CHK1* phosphorylates *Cdc25A*, promoting its proteolysis and inhibiting its interaction with *Cyclin B/Cdk1* [428]. |
| CyclinA | **CyclinA** = (**CyclinA_mRNA** and (not **pAPC**)) and ((**Cdc25A** and ((not **Cdh1**) or **Emi1**)) or (**CyclinA** and (((not **Cdh1**) and (**Emi1** or (not **UbcH10**))) or (**Emi1** and (not **UbcH10**))))) | | |
| | PC | | *Cyclin A* activity requires transcription (*Cyclin A* mRNA) and the absence of degradation by phosphorylated (mitotic) *pAPC*. In addition, turning ON inactive *Cyclin A* requires activation of *Cdk2* by *Cdc25A* [429] and the absence / *Emi1*-mediated inhibition of $APC/C^{Cdh1}$. Once active, *Cyclin A* maintains its activity in the absence of overpowering influences driving its degradation. Namely, *Cyclin A* relies on either *Emi1* or the absence of *UbcH10* for its ability to keep inactive $APC/C^{Cdh1}$ in check. To overpower active $APC/C^{Cdh1}$, *Cyclin A* requires both *Emi1* and no *UbcH10*. The precise combinatorial regulation of *Cyclin A* is not known; the above logic is consistent with *Cyclin A* activity pattern during cell cycle progression. |
| | ← TL | CyclinA _mRNA | Sustained availability of *Cyclin A* requires ongoing translation from *CyclinA* mRNA. |
| | ← Compl | Emi1 | *Emi1* binding to *Cdh1* is required to stabilize *Cyclin A* levels at the G1/S transition, allowing *Cyclin A/Cdk2* to block $APC/C^{Cdh1}$ [430, 431, 432]. |
| | ← DP | Cdc25A | *Cdc25A* promotes active *Cyclin A/Cdk2* complex formation by removing inhibitory phosphorylation of *Cdk2* [429, 433]. |
| | ← Per | CyclinA | We assume that once activated, *Cyclin A/Cdk2,1* complexes can sustain their activity until *Cyclin A* is degraded. |
| | ⊣ Deg | UbcH10 | *Cyclin A* degradation by $APC/C^{Cdh1}$ requires *UbcH10* [434]. |
| | ⊣ Deg | pAPC | *Cyclin A* is degraded by the $APC/C^{Cdc20}$ in prometaphase (as soon as the APC/C components are phosphorylated by Cdk1) [435, 436], before the full activation of the complex at SAC passage [437]. In our model, this stage of mitotic $APC/C^{Cdc20}$ activation is represented by *Cdk1*-phosphorylated *APC/C* (*pAPC*). |
| | ⊣ Deg | Cdh1 | *Cyclin A* is degraded by $APC/C^{Cdh1}$ in the presence of the *UbcH10* protein [438, 434, 345]. |
| Wee1 | **Wee1** = (((not **Casp3**) and (**Replication** or **CHK1**)) and (not(**Cdk1** and **CyclinB**))) and (**CHK1** or (not((**Cdk1** and **CyclinA**) and **Plk1**))) | | |
| | K | | *Wee1* is active during *Replication*, unless its activity is blocked by *CyclinA/Cdk1* OR *CyclinB/Cdk1* [439]. |
| | ⊣ P | Plk1 | *Plk1* phosphorylation at S53 promotes *Wee1* degradation [440]. This event is primed by *Cdk1* phosphorylation of *Wee1* at S123 [440]. As the main partner of *Cdk1* in mitosis is *Cyclin B*, we assume that assistance from *Plk1* to block *Wee1* is more relevant when paired with *Cyclin A/Cdk1* complexes. |

**Table S1q: Phase_SW module**

| | | | |
|---|---|---|---|
| | $\vdash$ P | CyclinA | *Cyclin A/Cdk1* is a strong inducer of *Wee1* phosphorylation and deactivation [439]. |
| | $\vdash$ P | CyclinB | *Cyclin B/Cdk1* is a strong inducer of *Wee1* phosphorylation and deactivation [439]. |
| | $\vdash$ P | Cdk1 | The somatic *Wee1* protein is an order of magnitude more sensitive to *Cdk1* activity than *Cdc25C*. Thus, both *Cyclin A/Cdk1* and *Cyclin B/Cdk1* strongly induce *Wee1* phosphorylation and deactivation [439]. |
| | $\leftarrow$ ComplProc | Replication | To model the sensitivity of *Wee1* activation to ongoing DNA synthesis even in the absence of damage, our model turns on *Wee1* immediately upon the start of DNA replication and maintains it until both Replication and the checkpoint kinase *Chk1* is OFF [441]. In addition, *Wee1* activity has been implicated in maintaining normal replication fork procession, linking its activity directly to ongoing replication [442]. |
| | $\leftarrow$ P | CHK1 | During DNA replication *Wee1* is activated by the checkpoint kinase *CHK1* [441]. |
| | $\vdash$ Lysis | Casp3 | *Caspase 3* cleaves and deactivates *Wee1* [443]. |
| UbcH10 | **UbcH10** = (not **Cdh1**) or (**UbcH10** and ((**Cdc20** or **CyclinA**) or **CyclinB**)) | | |
| | UbL | | The ubiquitin-conjugating enzyme (E2) *UbcH10* is active in the absence of *Cdh1*. Alternatively, active *UbcH10* is maintained in the presence of *Cdh1* when some of its targets are present: *Cdc20* OR *CyclinA* OR *CyclinB* [434]. |
| | $\leftarrow$ PBind | CyclinA | The presence of $APC/C^{Cdh1}$ substrates, including *Cyclin A*, inhibit the autoubiquitination of *UbcH10* but not its function, thus preserving APC activity [434]. |
| | $\leftarrow$ PBind | CyclinB | The presence of $APC/C^{Cdh1}$ substrates, including *CyclinB*, inhibit the autoubiquitination of *UbcH10* but not its function, thus preserving APC activity [434]. |
| | $\leftarrow$ Per | UbcH10 | Active *UbcH10* cannot be autoubiquitinated in the presence of $APC/C^{Cdh1}$ substrates and thus remains active [434]. |
| | $\leftarrow$ PBind | Cdc20 | The presence of $APC/C^{Cdh1}$ substrates, including *Cdc20*, inhibit the autoubiquitination of *UbcH10* but not its function, thus preserving APC activity [434]. |
| | $\vdash$ Deg | Cdh1 | *UbcH10* is degraded by $APC/C^{Cdh1}$. |
| CyclinB | **CyclinB** = (**FoxM1** or (**FoxO3** and **CyclinB**)) and (not(**Cdh1** or (**pAPC** and **Cdc20**))) | | |
| | PC | | *Cyclin B* node is ON when the concentration of *Cyclin B* proteins is high (does not represent the activity of *CyclinB/Cdk1* complexes). This occurs when *Cyclin B* is transcribed by *FoxM1*, maintained by *FoxO3* transcription, and not undergoing *APC*-mediated degradation by $APC/C^{Cdc20}$ or $APC/C^{Cdh1}$. |
| | $\leftarrow$ TR | FoxO3 | *FoxO3* is a direct transcriptional regulator of *Cyclin B*; its activation in G2 helps increase/maintain *Cyclin B* levels [444]. |

**Table S1q: Phase_SW module**

| | | | |
|---|---|---|---|
| | $\underset{TR}{\leftarrow}$ | FoxM1 | *FoxM1* is a direct transcriptional regulator of *Cyclin B1* [409, 445]. |
| | $\underset{Per}{\leftarrow}$ | CyclinB | Here we assume that FoxO3 alone can only maintain, but not independently induce *Cyclin B1* expression. |
| | $\underset{Deg}{\vdash}$ | pAPC | *Cyclin B* is degraded by $APC/C^{Cdc20}$ [438]. |
| | $\underset{Deg}{\vdash}$ | Cdc20 | *Cyclin B* is degraded by $APC/C^{Cdc20}$ [438]. |
| | $\underset{Deg}{\vdash}$ | Cdh1 | *Cyclin B* is degraded by $APC/C^{Cdh1}$ [438]. |
| Cdc25B | **Cdc25B** = **FoxM1** and **f4N_DNA** | | |
| | Ph | | *Cdc25B* activation requires transcription by *FoxM1*, centrosomal localization, and activation by *Aurora A* kinase on replicated centrosomes. |
| | $\underset{TR}{\leftarrow}$ | FoxM1 | *FoxM1* is an essential inducer or *Cdc25B* [446]. |
| | $\underset{Loc}{\leftarrow}$ | f4N_DNA | *Cdc25B* is localized at centrosomes, where it is activated by *Aurora A* kinase [447]. As *Aurora A* itself is only recruited to duplicated, centrosomes before their separation [448], *Cdc25B* activation requires duplicated centrosomes. As our model does not directly account for centrosome dynamics, we account for this by requiring the completion of S-phase (4N DNA). |
| Plk1 | **Plk1** = ((not **Cdh1**) and (**FoxM1** or **Plk1_H**)) and ((**CyclinB** and **Cdk1**) or ((**CyclinA** and (not **Wee1**)) and **Cdc25A**)) | | |
| | K | | *Plk1* activity requires the absence of $APC/C^{Cdh1}$, transcription by *FoxM1*, or high *Plk1* levels transcribed earlier by both *FoxM1* and *FoxO3* (see *Plk1_H* below) [444]. In addition, *Plk1* activation requires phosphorylation by either *CyclinB/Cdk1* during mitosis or *Cyclin A/Cdk2* (aided by lack of *Wee1* and *Cdc25A*) at the G2/M boundary [449]. |
| | $\underset{TR}{\leftarrow}$ | FoxM1 | *Plk1* is a direct transcriptional target of *FoxM1* [418]. |
| | $\underset{Ind}{\leftarrow}$ | Cdc25A | As we do not include a separate *Cdk2* node in our model, strong *Cyclin A/Cdk2* activity requires ongoing dephosphorylation of *Cdk2* by *Cdc25A* [450]. |
| | $\underset{Ind}{\vdash}$ | Wee1 | *Cyclin A*-mediated induction of *Plk1* is blocked by *Wee1* kinase, which specifically inhibits *Cdk2* activity [449]. |
| | $\underset{P}{\leftarrow}$ | CyclinA | *Plk1* activation at the G2/M boundary, before *Cdk1/Cuclin B* complexes are activated, requires active *Cyclin A/Cdk* [449]. |
| | $\underset{P}{\leftarrow}$ | CyclinB | *Plk1* is activated by *Cyclin B/Cdk1* phosphorylation [451, 452, 453]. |
| | $\underset{P}{\leftarrow}$ | Cdk1 | *Plk1* is activated by *Cyclin B/Cdk1* phosphorylation [451, 452, 453]. |

**Table S1q: Phase_SW module**

| | | | |
|---|---|---|---|
| | $\vdash$ Ubiq | Cdh1 | The majority of *Plk1* is degraded in anaphase by the $APC/C^{Cdh1}$ complex [454]. |
| | $\leftarrow$ Per | Plk1_H | Our model tracks the accumulauton of high-enough levels of *Plk1* to survive $APC/C^{Cdh1}$ mediated destruction into telophase via the *Plk1_H* node. Its ON state represents strong prior *Plk1* activation. Thus, it sustains the *Plk1* node in the absence of *FoxM1*-mediated transcription until *Plk1_H* itself is lost as *Plk1* levels fall. |
| Cdc25C | **Cdc25C** = (**f4N_DNA** and **Plk1**) and ((**Cdc25B** and (not **CHK1**)) or (**CyclinB** and **Cdk1**)) | | |
| | Ph | | In our model, *Cdc25C* is active in cells with replicated DNA (see *f4N_DNA* $\rightarrow$ *Cdc25C* link). Its activation is initiated by a small, initially cytoplasmic pool of *Cyclin B/Cdk1* activated by *Cdc25B* (not directly represented in our model) and further increased by *Cdc25B* itself, which translocates to the nucleus with the aid of *Plk1*. During mitosis, *Plk1* potentiates the ability of *Cyclin B/Cdk1* to maintain *Cdc25C* activity. |
| | $\leftarrow$ Ind | Cdc25B | *CDC25B* starts the cascade leading to mitotic entry by activating a small centrosomal pool of *Cyclin B/Cdk1*, leading to their nuclear translocation where they trigger the activation of *Cdc25C* and eventually the larger nuclear *Cyclin B/Cdk1* pool [455, 456, 457]. |
| | $\leftarrow$ P | Plk1 | In addition, *Plk1* induces nuclear transport of *CDC25B*, where it contributes to the initiation of *Cdk1* activity [458]. During mitosis, *Plk1* helps maintain strong *Cdc25C* activation by phosphorylating it on the same site as *Cyclin B/Cdk1* [459], as indicated by the profound decrease of Cdc25C activity in Plk1-inhibited mitotic cells [452, 460]. |
| | $\leftarrow$ P | CyclinB | *Cyclin B/Cdk1* complexes are potent activators of *Cdc25C*, creating positive feedback that causes switch-like mitotic entry [461, 462]. |
| | $\leftarrow$ P | Cdk1 | *Cyclin B/Cdk1* complexes are potent activators of *Cdc25C*, creating positive feedback that causes switch-like mitotic entry [461, 462]. |
| | $\vdash$ P | CHK1 | *CHK1* phosphorylates *Cdc25C*, leading to its nuclear exclusion, loss of access to its main target, *Cdk1* [463]. In addition, *CHK1* blocks the ability of *Cdc25B* to activate *Cdc25C* at the centrosomes by phosphorylating it and blocking its *Cdk1* activity [464, 465]. |
| | $\leftarrow$ Ind | f4N_DNA | The nature and localization of the signals responsible for the onset and maintenance of *Cdc25C* activity require replicated DNA (*f4N_DNA*) [458, 457]. Namely, *Cdc25C* is initially activated by a small pool of *Cyclin B/Cdk1* (below the ON-threshold of *Cdk1* in our model) which starts our at the replicated centrosome. Moreover, the pool of mitotic *Cdc25C* co-localized with active *Chk1/Cyclin B* is found on condensed chromosomes, again requiring the presence of *f4N_DNA* [463]. |
| Cdk1 | **Cdk1** = (**CyclinB** and **Cdc25C**) and ((not **CHK1**) or ((not **Wee1**) and **Cdk1**)) | | |

**Table S1q: Phase_SW module**

| | | | |
|---|---|---|---|
| | K | | Full *Cdk1* kinase activation requires its binding partner *Cyclin B* and the *Cdc25C* phosphatase, which maintains *Cdk1* in an active dephosphorylated state. *Cdk1* is inhibited by the checkpoint kinase *CHK1*, unless it is already full active and *Wee1* kinase is inhibited. |
| | $\vdash$ P | Wee1 | *Wee1* is a nuclear protein that ensures the completion of DNA replication prior to mitosis by blocking nuclear *Cdk1* activation [466]. |
| | $\leftarrow$ DP | Cdc25C | *Cdk1* is subject to inhibitory phosphorylation by *Wee1* or *Myt1*, and its dephosphorylation is carried out by activated *Cdc25C* [461, 467, 462]. |
| | $\leftarrow$ Compl | CyclinB | Full kinase activation of *Cdk1* in our model requires it to complex with *Cyclin B* [467]. |
| | $\leftarrow$ Per | Cdk1 | We assume that the presence of fully activated, nuclear *Cdk1* is able to overcome the effect of active *Wee1*, given that *Wee1* is very sensitive to *Cdk1*-mediated inhibitory phosphorylation [439]. |
| | $\vdash$ P | CHK1 | In the absence of *CHK1* kinase, a small cytosolic (centrosomal) pool of *Cyclin B/Cdk1* can be activated by *Cdc25B*, the nuclear translocation of which can trigger a positive feedback loop that activates the full *Cdk1* pool (assuming nuclear *Wee1* is also inactive). Thus, *CHK1* can maintain the OFF state of inactive *Cdk1* [449]. |
| pAPC | **pAPC** = (((**CyclinB** and **Cdk1**) and **Plk1**) or ((**CyclinB** and **Cdk1**) and **pAPC**)) or (**pAPC** and **Cdc20**) | | |
| | PC | | In line with evidence that *Plk1* can aid full activation of *APC/C*, but *Cdk1* appears to be the more potent inducer, our model requires both *Cyclin B/Cdk1* and *Plk1* to activate *APC/C* from an OFF state, but only *Cdk1* activity to maintain it. In addition, ongoing phosphorylation of the functional $APC/C^{Cdc20}$ complex is no longer required. |
| | $\leftarrow$ P | Plk1 | In addition to *Cyclin B/Cdk1* phosphorylation, full activation of the $APC/C^{Cdc20}$ complex also requires the kinase activity of *Plk1* [468]. |
| | $\leftarrow$ P | CyclinB | *CyclinB/Cdk1* activation triggers mitotic entry and promotes $APC/C^{Cdc20}$ activity via APC/C subunit phosphorylation [469, 470]. |
| | $\leftarrow$ P | Cdk1 | *CyclinB/Cdk1* activation triggers mitotic entry and promotes $APC/C^{Cdc20}$ activity via APC/C subunit phosphorylation [469, 470]. |
| | $\leftarrow$ Per | pAPC | Activated $APC/C^{Cdc20}$ initiates the Metaphase / Anaphase transition by degrading *Cyclin B* and securin [403]. Once active, $APC/C^{Cdc20}$ no longer requires sustained *CyclinB/Cdk1* or *Plk1* phosphorylation. |
| | $\leftarrow$ Compl | Cdc20 | Once active, $APC/C^{Cdc20}$ no longer requires sustained *CyclinB/Cdk1* or *Plk1* phosphorylation. |

**Table S1q: Phase_SW module**

| | | | |
|---|---|---|---|
| Cdc20 | **Cdc20** = ((**pAPC**and(not **Emi1**))and(not **Cdh1**))and((not **Mad2**)or((not **CyclinA**)and (not(**CyclinB** and **Cdk1**)))) | | |
| | Prot | | In our model, *APC/C*$^{Cdc20}$ complex formation is represented by the joint activity of *Cdc20* and phosphorylated *APC/C* (*pAPC*). *Cdc20* is thus ON in the presence of *pAPC* when both *Emi1* and *Cdh1* are absent (*APC/C*$^{Cdh1}$ is represented by the *Cdh1* node, see below). In addition, *Cdc20* activity requires either the absence of *Mad2* at unattached kinetochores, or the absence of *Cdc20* phosphorylation by *Cyclin B/Cdk1* or by *Cyclin A/Cdk2* complexes to potentiate the interaction between *Mad2* and *Cdc20*, and *pAPC* is ON (present and phosphorylated) [471]. |
| | $\dashv$ IBind | Emi1 | *Emi1* binds *Cdc20* and inhibits the ubiquitin ligase activity of $APC/C^{Cdc20}$ [431]. |
| | $\dashv$ P | CyclinA | *Cyclin A/Cdk2* complexes phosphorylate *Cdc20* and inactivate the $APC/C^{Cdc20}$ complex during S and G2 [472]. |
| | $\dashv$ P | CyclinB | *Cyclin B* partners with *Cdk1* to keep *Cdc20* phosphorylated, increasing its interaction with *Mad2* rather than *APC/C* [473]. |
| | $\dashv$ P | Cdk1 | *Cdk1*-phosphorylated *Cdc20* interacts with *Mad2* rather than *APC/C*, resulting in a block on $APC/C^{Cdc20}$ activation until completion of spindle assembly [471]. |
| | $\leftarrow$ Compl | pAPC | *Cdc20* becomes active in early mitosis by binding to *APC/C*, an event that requires *Cyclin B/Cdk1*-mediated phosphorylation of several core *APC/C* subunits [427, 474]. |
| | $\dashv$ Deg | Cdh1 | $APC/C^{Cdh1}$ complexes degrade *Cdc20*, leading to a complete switch from $APC/C^{Cdc20}$ to $APC/C^{Cdh1}$ during mitotic exit [427, 475]. |
| | $\dashv$ IBind | Mad2 | Eukaryotic cells do not separate their replicated genome until they pass the Spindle Assembly Checkpoint (SAC). Namely, all their chromosomes need to be are aligned with respect to the metaphase plane and the two copies of each chromosome need to be attached to opposite poles of the mitotic spindle [470]. This physical alignment is monitored via *Mad2*: kinetochores that remain unattached to microtubules catalyze the sequestration of *Cdc20* and thus inhibit $APC/C^{Cdc20}$ [476, 477]. |
| Cdh1 | **Cdh1** = (not(**CyclinB** and **Cdk1**)) and (not(**CyclinA** and (**Emi1** or **Cdc25A**))) | | |
| | PC | | *APC/C*$^{Cdh1}$ activity requires the absence of Cyclin Dependent kinase phosphorylation by *Cyclin B/Cdk1*, or *Cyclin A/Cdk2* aided by further inhibition of *Cdh1* by *Emi1*, or ongoing *Cdk2* activation by *Cdc25A* in the absence of *Emi1*. |
| | $\dashv$ IBind | Emi1 | *Emi1* blocks *APC/C*$^{Cdh1}$ binding to its substrates [432], as well as its ability to add ubiquitin chains to them [478]. |

**Table S1q: Phase_SW module**

| | | | |
|---|---|---|---|
| | $\vdash$ Ind | Cdc25A | As we do not include a separate *Cdk2* node in our model, strong *Cyclin A/Cdk2* activity capable of overriding *Cdh1* activity even in the presence of *Emi1* requires ongoing dephosphorylation of *Cdk2* by *Cdc25A* [461]. |
| | $\vdash$ P | CyclinA | Active *Cyclin A/Cdk1,2* complexes phosphorylate *Cdh1* during S, G2 and early mitosis, impairing its interaction with *APC/C* until late stages of mitosis when *Cdk1/2* activity falls [427, 438]. |
| | $\vdash$ P | CyclinB | *Cyclin B/Cdk1* phosphorylates *Cdh1* during mitosis, impairing its interaction with *APC/C* [427, 438]. |
| | $\vdash$ P | Cdk1 | *Cyclin B/Cdk1* phosphorylates *Cdh1* during mitosis, impairing its interaction with *APC/C* [427, 438]. |

**Table S1r: Cell_Cycle_Process module**

| Target Node | Node Gate<br>Node Type<br>Link Type | Node Description<br>Input Node | Link Description |
|---|---|---|---|
| Replication | **Replication** = ((not **CAD**) and **Pre_RC**) and (((**E2F1** and **CyclinE**) and **Cdc25A**) or (((**Replication** and **CyclinA**) and **Cdc25A**) and (**E2F1** or (not **f4N_DNA**)))) | | |
| | Proc | The *Replication* node represents ongoing DNA synthesis. This requires a non-apoptotic cell, licensed pre-replication complexes (*Pre_RC*). The start of DNA synthesis requires *E2F1*-meditated transcription of the genes that help execute it, as well as the firing of the first round of replication origins by *Cyclin E/Cdk2*. Once ongoing, replication is sustained by *Cyclin A/Cdk2* and *Cdc25A*, aided by *E2F1* and terminated by completion of a full round of synthesis (*4N_DNA*). | |
| | $\leftarrow$ ComplProc | E2F1 | In addition to *E2F1* target genes directly included in our model, *E2F1* transcribes an array of critical S-phase genes responsible for carrying out DNA synthesis (e.g, *POLA1*, *POLA2*, *MCM3*, *MCM5*, *MCM6*, *PCNA*, *TOP2A*, *RFC2*, *TK1*) [479, 480]. |
| | $\leftarrow$ ComplProc | CyclinE | DNA replication is initiated by fully active *Cyclin E/Cdk2* [481]. |
| | $\leftarrow$ ComplProc | Pre_RC | Ongoing DNA replication requires licensed replication origins, which fire throughout DNA synthesis [482]. |
| | $\leftarrow$ ComplProc | Cdc25A | Active *Cdc25A* is required for onset as well as progression through S-phase [483, 484]. |
| | $\leftarrow$ ComplProc | CyclinA | *Cyclin A/Cdk1* complexes regulate the origin firing program in mammalian cells and are required for the completion of DNA replication [484, 433]. |

**Table S1r: Cell_Cycle_Process module**

| | | | |
|---|---|---|---|
| | $\leftarrow_{\text{Per}}$ | Replication | Once ongoing, DNA synthesis continues in the presence of active *Cyclin A/Cdk2*, only ending when DNA content is doubled. |
| | $\vdash_{\text{ComplProc}}$ | f4N_DNA | Complete duplication of a cell's DNA, represented in our model by *f4N_DNA* = ON, marks the end of active *Replication*. |
| | $\vdash_{\text{ComplProc}}$ | CAD | Caspase-activated DNase (*CAD*) destroys DNA, preventing ongoing replication. |
| ATR | **ATR = Replication** | | |
| | K | | *ATR* accumulates at replication forks during unperturbed DNA synthesis [485]. |
| | $\leftarrow_{\text{Loc}}$ | Replication | *ATR* accumulates at replication forks during unperturbed DNA synthesis [485]. |
| CHK1 | **CHK1 = ATR** | | |
| | K | | *ATR* kinase activates *CHK1* at replication forks, which not only blocks premature mitosis but also regulates the rate of origin firing by keeping *Cdc25* protein levels from increasing above their physiological range [485]. |
| | $\leftarrow_{\text{P}}$ | ATR | *ATR* kinase activates *CHK1* at replication forks (by phosphorylation of serines 317 and 345), which not only blocks premature mitosis but also regulates the rate of origin firing by keeping *Cdc25* protein levels from increasing above their physiological range [485]. |
| f4N_DNA | **f4N_DNA** = (not **CAD**) and ((**Replication** and ((**Pre_RC** and **CyclinA**) or **f4N_DNA**)) or (**f4N_DNA** and (not **Cytokinesis**))) | | |
| | MSt | | 4N DNA content in our model is reached via the completion of *Replication* (via the firing of the last round of replication origins by *Cyclin A/Cdk* complexes) and maintained in non-apoptotic cells the absence of a contractile ring driving cytokinesis. |
| | $\leftarrow_{\text{ComplProc}}$ | Pre_RC | *Replication* can only complete DNA synthesis and produce double DNA content if the availability of licensed replication origins is not blocked [482]. |
| | $\leftarrow_{\text{ComplProc}}$ | CyclinA | *Cyclin A/Cdk1* complexes regulate the origin firing program in mammalian cells and are required for the completion of DNA replication [481, 433]. |
| | $\leftarrow_{\text{ComplProc}}$ | Replication | DNA content is doubled by the process of *Replication*. |
| | $\vdash_{\text{Per}}$ | f4N_DNA | Once achieved, a cell's 4N DNA content is sustained up to the point of cytokinesis. |
| | $\vdash_{\text{ComplProc}}$ | Cytokinesis | The process of cytokinesis separates the replicated sister chromatids and resets the DNA content of each daughter cell to a diploid 2N. |
| | $\vdash_{\text{Deg}}$ | CAD | Caspase-activated DNase (*CAD*) destroys DNA, preventing maintenance of a double DNA content. |

**Table S1r: Cell_Cycle_Process module**

| | | | |
|---|---|---|---|
| U_Kinetochores | **U_Kinetochores** = ((**f4N_DNA** and (not **Cdh1**)) and (not **A_Kinetochores**)) and ((**CyclinB** and **Cdk1**) or **U_Kinetochores**) | | |
| | MSt | The *U_Kinetochores* node in our model is on from the moment the nuclear envelope is dissolved in prometaphase and the mitotic spindle starts to form, until all kinetochores are properly attached. In addition to the presence of unattached kinetochores, *U_Kinetochores* = ON requires attached sister chromatics, the absence of $APC/C^{Cdh1}$ activity. It is turned on my *Cyclin B/Cdk1* and remains on until the spindle is complete (or it is destroyed by $APC/C^{Cdh1}$). | |
| | ← ComplProc | CyclinB | The start of mitotic spindle assembly is initiated by active *Cyclin B/Cdk1* [486]. |
| | ← ComplProc | Cdk1 | The start of mitotic spindle assembly is initiated by active *Cyclin B/Cdk1* [486]. |
| | ⊣ ComplProc | Cdh1 | Premature activation of $APC/C^{Cdh1}$ destroys the incomplete spindle by triggering premature, aberrant anaphase. This occurs due to premature degradation of *APC/C* targets including *Securin* (responsible for keeping sister chromatids attached [487]), *Cyclin B*, *Cdc20*, and Aurora kinase A (*AURKA*) [488]. |
| | ← ComplProc | f4N_DNA | Metaphase requires replicated sister chromatids (*f4N_DNA*), held together by their kinetochores, face in opposing directions and can be attached to opposite poles of the mitotic spindle. |
| | ← Per | U_Kinetochores | Once metaphase starts, the mitotic spindle remains incomplete as long as some of the kinetochores remain unattached. |
| | ⊣ ComplProc | A_Kinetochores | In our model, the transition from unattached to all attached kinetochores (*U_Kinetochores* → *A_Kinetochores*) marks the completion of the mitotic spindle and Spindle Assembly Checkpoint (SAC) passage. |
| Mad2 | **Mad2** = **U_Kinetochores** and (not **A_Kinetochores**) | | |
| | Prot | Our model represents the SAC via the *Mad2* kinetochore-binding protein. *Mad2* is active as long as the cell has at least one unattached kinetochore and it is responsible for keeping *Cdc20* sequestered from *APC/C*. By keeping *APC* at bay until the spindle is complete, *Mad2* is required for the proper timing of anaphase [477]. | |
| | ← Compl | U_Kinetochores | The *Mad2* SAC protein is active and potent in the presence of even a single unattached kinetochore [477]. |
| | ⊣ ComplProc | A_Kinetochores | *Mad2* is inhibited by SAC passage, marked by the completion of the spindle and proper attachment of all kinetochore [477]. |
| A_Kinetochores | **A_Kinetochores** = ((**f4N_DNA** and (not **Cdh1**)) and (not(**pAPC** and **Cdc20**))) and (**A_Kinetochores** or ((((**U_Kinetochores** and **Src**) and **Plk1**) and **CyclinB**) and **Cdk1**)) | | |

**Table S1r: Cell_Cycle_Process module**

| | | | |
|---|---|---|---|
| | MSt | The completed spindle, represented by the *A_ Kinetochores* node, requires replicated and attached sister chromatids (*f4N_ DNA*) and the absence of *APC/C* activity. It turns on when the process of spindle assembly (*U_ Kinetochores*) is completed by active *Src*, active *Plk1* localized to unattached kinetochores in the presence of ongoing *Cyclin B/Cdk1* activity, and it remains on until anaphase (*APC/C* activation). | |
| | ← ComplProc | Src | *Src* promotes correct spindle orientation [489]. Moreover, absence of *c-Src* leads to severely reduced astral microtubules [490]. Finally, *Src*-mediated phosphorylation of the *Eg5* motor domain is required for the formation of a bipolar spindle and correct chromosome segregation [491]. |
| | ← ComplProc | Plk1 | *Plk1* activity at unattached kinetochores is required for promoting their attachment [492]. In its absence, kinetochores remain unattached and cells eventually undergo mitotic catastrophe and apoptosis [493]. |
| | ← ComplProc | CyclinB | Ongoing *Cyclin B/Cdk1* at unattached kinetochores is necessary to keep *Plk1* active and allow the completion of mitosis [494]. |
| | ← ComplProc | Cdk1 | Ongoing *Cyclin B/Cdk1* at unattached kinetochores is necessary to keep *Plk1* active and allow the completion of mitosis [494]. |
| | ⊢ Deg | pAPC | During normal mitosis, the completed spindle is pulled apart in response to $APC/C^{Cdc20}$-mediated degradation of *Securin*, which normally blocks *Separate* from severing the *Cohesin* rings keeping sister chromatids attached [487]. |
| | ⊢ Deg | Cdc20 | During normal mitosis, the completed spindle is pulled apart in response to $APC/C^{Cdc20}$-mediated degradation of *Securin*, which normally blocks *Separate* from severing the *Cohesin* rings keeping sister chromatids attached [487]. |
| | ⊢ Deg | Cdh1 | $APC/C^{Cdh1}$ destroys the spindle by triggering anaphase via the degradation of *APC/C* targets, including *Securin* [488]. |
| | ← ComplProc | f4N_DNA | Completion of the mitotic spindle requires replicated and attached sister chromatids (*f4N_ DNA*). |
| | ← ComplProc | U _Kinetochores | The mitotic spindle is assembled gradually, as the number of unattached kinetochores gradually decreased by the formation of microtubule attachments. |
| | ← Per | A _Kinetochores | Once assembled, separation of the mitotic spindle requires *APC/C* activity to promote the destruction of sister chromatid cohesion [487]. |
| Plk1_H | **Plk1_H** = (**Plk1** and **FoxM1**) and ((**Plk1_H** or **FoxO3**) or **FoxO1**) | | |
| | K | The ON state of *Plk1_ H* encodes the short-lived memory of a sufficiently large active *Plk1* pool to temporarily survive *Plk1* destruction by $APC/C^{Cdh1}$ [454], recruit *Ect2* to the central spindle, and thus aid the completion of cytokinesis [495]. Thus, *Plk1_ H* requires ongoing *Plk1* activation and transcription by *FoxM1*, and either induction by *FoxO3* or *FoxO1*, or prior accumulation. | |

#### Table S1r: Cell_Cycle_Process module

| | | | |
|---|---|---|---|
| | ← TR | FoxO3 | *Plk1* is a direct transcriptional target of *FoxO3*, but *Plk1* appears to be sufficiently induced in the absence of *FoxO* preteens to aid its G2/M and mitotic functions. In contrast, accumulation if a large enough *Plk1* pool to briefly outlast $APC/C^{Cdh1}$ activation (modeled by the *Plk1_H* node), requires *FoxO* activity in G2 [444]. |
| | ← TR | FoxO1 | In addition to *FoxO3*, *FoxO1* also binds the *Plk1* promoter, potentially aiding its accumulation during G2 [496]. |
| | ← TR | FoxM1 | *Plk1* is a direct transcriptional target of *FoxM1*; loss of *FoxM1* severely reduces *Plk1* protein levels [408, 418]. |
| | ← Per | Plk1 | Active mitotic *Plk1* is a prerequisite for the accumulation of the larger active *Plk1* pool denoted by *Plk1_H*. |
| | ← Per | Plk1_H | Once accumulated, we assume that the *Plk1_H* pool of active *Plk1* remains stable in the absence of *FoxO*-mediated transcription. This is supported by negative feedback regulation of *FoxO* proteins by *Plk1* [119], indicating that ongoing high FoxO activity is likely not required for the maintenance of *Plk1_H*. |
| Ect2 | **Ect2** = (((**f4N_DNA** and **Plk1_H**) and **Cdh1**) and (not **U_Kinetochores**)) and (not **A_Kinetochores**) | | |
| | GEF | | *Ect2* activaiton at the spindle midzone represents the step of cytokinesis in our model. Thus, *Ect2* requires *f4N_DNA*, high *Plk1* activity, as well as *Cdh1* for the assembly of a normal spindle midzone. Finally, *Ect2* cannot be recruited to the mid zone before anaphase is completed. |
| | ← Ind | Cdh1 | $APC/C^{Cdh1}$-mediated destruction of Aurora kinase is required for the assembly of a robust spindle midzone at anaphase and for the normal timing of cytokinesis [497]. |
| | ← Ind | f4N_DNA | Formation of a spindle midzone, where *Ect2* accumulates in preparation of cytokinesis requires recently separated sister chromatids (4N DNA content). |
| | ⊣ ComplProc | U _Kinetochores | Formation of a spindle midzone requires the separation of sister chromatids; thus it cannot occur before anaphase. |
| | ⊣ ComplProc | A _Kinetochores | Formation of a spindle midzone requires the separation of sister chromatids; thus it cannot occur before anaphase. |
| | ← Ind | Plk1_H | *Plk1* activity in telophase (*Plk1_H*) is required for the recruitment of *Ect2* to the central spindle [454, 498]. |
| Cytokinesis | **Cytokinesis** = (**Ect2** and **FAK**) and **Src** | | |
| | Proc | | In contrast to our previous model in [147] where *Ect2* recruitment to the central spindle marked the start of cytokinesis, in [148] we introduced a separate *Cytokinesis* node to mark cytokinesis and the subsequent resetting of daughter cell DNA content to 2N by a separate node. In addition to *Ect2* recruitment, completion of cytokinesis also requires ECM attachments able to activate *FAK* and *Src* kinases. |

**Table S1r: Cell_Cycle_Process module**

| | | | |
|---|---|---|---|
| | ← ComplProc | FAK | Integrin-activated *FAK* and *Src* control cytokinetic abscission by decelerating *PLK1* degradation at aiding *CEP55* in recruiting abscission process proteins to the midbody [172, 499]. |
| | ← ComplProc | Src | Integrin-activated *FAK* and *Src* control cytokinetic abscission by decelerating *PLK1* degradation at aiding *CEP55* in recruiting abscission process proteins to the midbody [172, 499]. |
| | ← Loc | Ect2 | At the start of cytokinesis, the *Ect2 RhoGEF* is recruited to the central spindle [495]. *Ect2* aids the accumulation of GTP-bound *RhoA* [500, 495] and the formation of the contractile ring. |

**Table S1s: TRAIL module**

| Target Node | Node Gate<br>Node Type<br>Link Type | Node Description<br>Input Node | Link Description |
|---|---|---|---|
| Trail | **Trail = Trail** | | |
| | Env | The *Trail* node represents environmental availability of the *Trail* protein outside the cell. | |
| | ← Env | Trail | The *Trail* input node remains on/off if set ON/OFF in the absence of *in silico* perturbation. |

**Table S1t: Apoptotic_SW module**

| Target Node | Node Gate<br>Node Type<br>Link Type | Node Description<br>Input Node | Link Description |
|---|---|---|---|
| DR4_5 | **DR4_5 = Trail** | | |
| | Rec | The *DR4* and *DR5* death receptors, represented by the *DR4_5* node, are activated by extracellular *Trail* [501]. | |
| | ← Ligand | Trail | *DR4* and *DR5* death receptors are activated by extracellular *Trail* [501]. |
| Casp8 | **Casp8 = DR4_5** or **Casp3** | | |
| | PTase | *Pro-Caspase 8* may be cleaved independently by *DISC* (not directly reprsented; an adaptor protein for *DR4_5*) or *Caspase 3*. | |
| | ← Compl | DR4_5 | *Trail*-bound (active) *DR4* and *DR5* receptors trigger the assembly of the pro-apoptotic death-inducing signaling complex (*DISC*), which binds a cluster of *pro-Caspase 8* proteins and initiates their cleavage into active *Caspase 8* [502]. |

**Table S1t: Apoptotic_SW module**

| | | | |
|---|---|---|---|
| | ← Ind | Casp3 | *Caspase 3* indirectly activates *Caspase 8* by cleaving *Caspase 6* [503], which, in turn, cleaves *Caspase 8* [504]. |
| Casp2 | **Casp2** = **Casp3** or ((**U_Kinetochores** and **Mad2**) and (not(**CyclinB** and **Cdk1**))) | | |
| | PTase | | *Pro-caspase 2* is cleaved and activated by *Caspase 3*, or by failed cytokinesis marked by the presence of unattached kinetochores, an active SAC, and the absence of active *Cyclin B/Cdk1* complexes to phosphorylate and inhibit *Caspase 2*. |
| | ⊣ P | CyclinB | *Cyclin B1/Cdk1* phosphorylate *caspase-2* at Ser 340, preventing its activation [505]. |
| | ⊣ P | Cdk1 | *Cyclin B1/Cdk1* phosphorylate *caspase-2* at Ser 340, preventing its activation [505]. |
| | ← Ind | Mad2 | A functional spindle assembly checkpoint is required for mitotic cell death upon prolonged mitotic arrest [506] or spindle damage [507]. |
| | ← Ind | U _Kinetochores | Although the precise molecular mechanism by which *Caspase 2* is activated during prolonged or stalled mitosis is unclear, its activation platform, the *PIDDosome*, has been localized to unattached kinetochores [508]. Even though a checkpoint protein keeps the *PIDDosome* unresponsive to DNA damage signals, the loss of protective *Cyclin B/Cdk1* phosphorylation only leads to *Caspase 2* activation in the presence of a partially assembled mitotic spindle, and requires active SAC. |
| | ← Lysis | Casp3 | *Caspase 2* is a target of *Caspase 3*, as its inhibition severely limits *Caspase 2* cleavage during apoptosis [509, 510]. |
| MCL_1 | **MCL_1** = (((not **Casp3**) and (not **Casp2**)) and ((not **GSK3**) or (**AKT_B** and (**ERK** or (not **E2F1**))))) and (not((**Cdk1** and **CyclinB**) and **U_Kinetochores**)) | | |
| | Prot | | *Caspase 3* or *2*-mediated destruction of *MCL-1* must be absent for *MCL-1* to be ON. Avoiding degradation via textitGSK3 requires the *GSK3*-weakenjng presence of basal *AKT* activity (*AKT_B*) [511] and either *ERK*-mediated stabilization, or the absence of its repressor *E2F1*. Finally, during mitotic arrest (*U_Kinetochores*), *MCL-1* is deactivated by *Cyclin B/Cdk1* phosphorylation, which shields it from the *PPA2*-mediated dephosphorylation of its degradation-targeting sites [512]. |
| | ← P | ERK | *ERK* phosphorylates *MCL-1*, promoting its interaction with *Pin1*, which stabilizes it [513, 514]. |
| | ← Ind | AKT_B | In order to account for the loss of *MCL-1* in the complete absence of growth factors versus its presence in low growth factor environments, we required basal *AKT* to modulate the strength of *GSK3* inhibition [511]. |
| | ⊣ P | GSK3 | *MCL-1* is phosphorylated by *GSK3*, leading to ubiquitinylation and degradation of Phosphorylation [511]. |
| | ⊣ TR | E2F1 | *E2F1* is a direct transcriptional repressor of *MCL-1* [515]. |

**Table S1t: Apoptotic_SW module**

| | | | |
|---|---|---|---|
| | ⊢ P | CyclinB | In cells arrested in mitosis, phosphorylation by *Cyclin B/Cdk1* on T92 initiates *MCL-1* degradation [516]. |
| | ⊢ P | Cdk1 | Phosphorylation by *Cyclin B/Cdk1* in cells arrested in mitosis initiates *MCL-1* degradation [516]. |
| | ⊢ Ind | U _Kinetochores | During prolonged mitotic arrest (*U_kinetochores*), *MCL-1* levels drop steadily due to phosphorylation by *JNK*, *p38* and/or *CKII* and its subsequent degradation by the E3 ubiquitin ligase *SCF* (*FBW7*) [512]. |
| | ⊢ Lysis | Casp2 | *Caspase 2* activation destabilizes the *MCL-1* protein [517]. |
| | ⊢ Lysis | Casp3 | *Caspase 3* cleaves and deactivated *MCL-1* [518]. |
| BCLXL | **BCLXL** = ((not **Casp3**) and ((**BCL2** and (not **SMAD2_3_4**)) and (not **BAD**))) and (((not **U_Kinetochores**) or (**Plk1** and ((not(**CyclinB** and **Cdk1**)) or (**BCL2** and **MCL_1**)))) or ((**BCL2** and **MCL_1**) and (not(**CyclinB** and **Cdk1**)))) | | |
| | Prot | | *Bcl-$x_L$* activity requires the absence of *Caspase 3*. In addition, *BAD* can block *Bcl-$x_L$*, as it preferentially binds to it rather than *BCL2* (meaning in the absence of the latter *Bcl-$x_L$* is more likely to be sequestered by basal levels of *BAD*) [519]. Similarly, *TFG$\beta$* can also repress *Bcl-$x_L$* [520]. Lastly, mitotic *Bcl-$x_L$* can be inhibited by *Cdk1* activity if either *BCL2*, *MCL-1*, or *Plk1* are OFF. In the absence of *Plk1*, loss of either *BCL2* or *MCL-1* can result in *Bcl-$x_L$* inhibition (even without *Cdk1* phosphorylation), as we assume its targets are no longer competitively bound by its family members. |
| | ⊢ Ind | SMAD2_3 _4 | *BCL-$X_L$* is repressed by *TFG$\beta$* signaling, an event that mediates *TFG$\beta$*-induced apoptosis [520]. |
| | ← Ind | Plk1 | In addition to other effects of prolonged mitotic arrest on *BCL-2* proteins, *Plk1* inhibition synergistically enhances the inhibitory phosphorylation of *BCL-2* and *BCL-xL*, as well as downregulation of *MCL-1* [521]. |
| | ⊢ P | CyclinB | During normal mitosis, *Cyclin B/Cdk1* only transiently phosphorylates part of the *BCL-xL* pool. Prolonged mitosis, however, results in high levels of *BCL-xL* (and *Bcl-2*) phosphorylation, priming the system for *Caspase 2*-mediated apoptosis [522, 523]. |
| | ⊢ P | Cdk1 | During normal mitosis, *Cyclin B/Cdk1* only transiently phosphorylates part of the *BCL-xL* pool. Prolonged mitosis, however, results in high levels of *BCL-xL* (and *Bcl-2*) phosphorylation, priming the system for *Caspase 2*-mediated apoptosis [522, 523]. |
| | ⊢ Ind | U _Kinetochores | Prolonged mitosis is required for the accumulation of *BCL-xL* phosphorylation, weakening its interaction with *Bax* [524]. |
| | ← Ind | MCL_1 | *MCL-1* competes with *BCL-xL* for *BAK* binding; the presence of *MCL-1* can keep part of the *BCL-xL* pool active [525]. |

#### Table S1t: Apoptotic_SW module

| | | | |
|---|---|---|---|
| | ← Ind | BCL2 | *BCL2* competes with *BCL-xL* for *BAD* binding. As *BCL-xL* is a stronger binding partner of *BAD* [519], here we assume that loss of *BCL2* or *BAD* can both result in low *BCL-xL* activity. |
| | ⊢ IBind | BAD | *Bad* can bind *BCL-xL* and displace it from *BAX*, thus deactivating it [519]. |
| | ⊢ Lysis | Casp3 | *BCL2* is cleaved and deactivated by *Caspase 3* [526]. |
| BCL2 | **BCL2** = not(**Casp3** or **BAD** or **BIM** or **BIK**) and ((not **U_Kinetochores** or (**MCL_1** and **BCLXL**)) or (**Plk1** and (**BCLXL** or **MCL_1** or not(**Cdk1** and **CyclinB**))) or **GLI** or **GLI_H**) | | |
| | Prot | | While the precise combinatorial logic governing *BCL2* activity is not clear from literature, we modeled *BCL2* as ON in the absence of *Caspase 3*, *BAD*, *BIM* or *BIK*. This choice makes *BCL2* the most sensitive of the three family members to activation of its three inhibitors. In addition, mitotic *BCL2* is blocked by *Cdk1* if both *BCL-xL* and *MCL-1* are OFF. In the absence of *Plk1*, loss of either *BCL2* or *MCL-1* can result in *BCL-2* inhibition (even without *Cdk1* phosphorylation), as we assume its targets are no longer competitively bound by its family members. This effect can be counteracted by *GLI2*-mediated *BCL2* induction [527]. |
| | ← TR | GLI | *GLI2* is a direct transcriptional inducer of *BCL2* [527]. |
| | ← TR | GLI_H | *GLI2* is a direct transcriptional inducer of *BCL2* [527]. |
| | ← Ind | Plk1 | In addition to other effects of prolonged mitotic arrest on *BCL2* proteins, *Plk1* inhibition synergistically enhances the inhibitory phosphorylation of *BCL2* and *BCL-xL*, as well as downregulation of *MCL-1* [521]. |
| | ⊢ P | CyclinB | *Cyclin B/Cdk1* phosphorylates *BCL2* (and *BCL-xL*) during mitosis [522, 523, 526]. |
| | ⊢ P | Cdk1 | Prolonged mitosis results in high levels of *BCL-xL* and *BCL2* phosphorylation, priming the system for *Caspase 2*-mediated apoptosis [522, 523, 526]. |
| | ⊢ Ind | U _Kinetochores | Prolonged mitosis is required for the accumulation of *BCL2* phosphorylation [522, 523, 526]. |
| | ← Ind | MCL_1 | *MCL-1* competes with *BCL-xL* for binding most of their apoptotic partners, including *BIK*, *BIM*, *BID*, *BAX* and *BAK*. |
| | ← Ind | BCLXL | *BCL2* competes with *BCL-xL* for binding most of their apoptotic partners, including *BIK*, *BIM*, *BID*, *BAX* and *BAK*. |
| | ⊢ IBind | BAD | *BCL2* competes with *BCL-xL* for *BAD* binding. *BAD* displaces *BCL2* from its inhibitory binding of *Bax/Bak*. Although *BCL-xL* is a stronger binding partner, we assume that *BAD* alone cannot fully block *BCL-xL* in the presence of *BCL2* [519]. |

**Table S1t: Apoptotic_SW module**

| | | | |
|---|---|---|---|
| | $\vdash$ IBind | BIK | *BIK* binds *BCL2* and they mutually inhibit each other's activity [528]. |
| | $\vdash$ IBind | BIM | *BIM* binds *BCL2* and they mutually inhibit each other's ability to activate further targets [529]. |
| | $\vdash$ Lysis | Casp3 | *BCL2* is cleaved and deactivated by *Caspase 3* [526]. |
| BAD | **BAD** = (**Casp3** or (not(((**AKT_H** or **AKT_B**) or **ERK**) or **S6K**))) or (**Casp8** and ((not((**AKT_B** and **ERK**) and **S6K**)) and (not(**AKT_H** and (**AKT_B** or **ERK**))))) | | |
| | Prot | *BAD* in our model is ON when cleaved by *Caspase 3*, or in the complete absence of survival signals (*AKT*, *ERK* or *S6K*). Alternatively, *BAD* can be cleaved and activated by *Caspase 8* in the absence of strong survival signaling. We modeled this inhibitory survival signal as either the combined activity of *ERK*, *S6K* and (at least) basal *AKT*, or high *AKT* in the joint presence of ERK and basal *AKT* (indicating that *AKT_H* will not drop by the next time-step). | |
| | $\vdash$ P | ERK | *ERK* phosphorylates *BAD* at Ser-112, inducing its sequestration away from the mitochondrial membrane where its BCL-2 family targets are located (e.g., *BCL-2*, *BCL-xL*)[530]. |
| | $\vdash$ P | AKT_B | *Akt* phosphorylates *BAD* at Ser-136, inducing its sequestration away from the mitochondrial membrane where its BCL-2 family targets are located (e.g., *BCL2*, *BCL-xL*) [531]. |
| | $\vdash$ P | AKT_H | *Akt* phosphorylates *BAD* at Ser-136, inducing its sequestration away from the mitochondrial membrane where its BCL-2 family targets are located (e.g., *BCL2*, *BCL-xL*) [531]. |
| | $\vdash$ P | S6K | *S6K1* phosphorylates *BAD* at Ser-155, directly blocking its binding to *BCL-xL* [532]. |
| | $\leftarrow$ Lysis | Casp8 | *Caspase 8* is also able to cleave *BAD*, generating a more potently apoptotic fragment [533]. In addition, *TRAIL*-mediated apoptosis results in *BAD* cleavage by a Caspase upstream of MOMP, creating a potent apoptotic inducer before full *Caspase 3* activation [534]. |
| | $\leftarrow$ Lysis | Casp3 | *Caspase 3* cleaves *BAD*, generating a more potently apoptotic fragment [533]. |
| BIK | **BIK** = (not((**MCL_1** or **BCLXL**) or **BCL2**)) and (**SMAD2_3_4** or (not **AKT_H**)) | | |
| | Prot | *BIK* is free to activate its target, *BAX*, only when it is not sequestered by any of the three *BCL-2* family proteins [525] and it is either induced by *SMADs* [520] or protected by no more than basal *AKT* activity, which lowers the level of both *BCL2* and *BCL-$x_L$* [535]. | |
| | $\vdash$ Ind | AKT_H | *AKT_H* elevates the level of both *BCL2* [536] and *BCL-$x_L$* [535], thus making it easier for these inhibitors to sequester *BIK*. |
| | $\leftarrow$ TR | SMAD2_3_4 | *Smad2/3/4* are direct transciprional inducers of *BIK* [520]. |

#### Table S1t: Apoptotic_SW module

| | | | |
|---|---|---|---|
| | ⊢ IBind | MCL_1 | *MCL-1* binds *BIK*; they mutually inhibit each other [537]. |
| | ⊢ IBind | BCLXL | *BCL-xL* binds *BIK*; they mutually inhibit each other [538]. |
| | ⊢ IBind | BCL2 | *BCL2* binds *BIK*; they mutually inhibit each other [528]. |
| BIM | **BIM** = **FoxO3** and ((**GSK3** and not(**ERK** or **MCL_1** or **ZEB1** or **BCLXL** or **BCL2**)) or ((**Runx1** and not **ERK** and not **NfkB**) and (not((**MCL_1** and **BCLXL**) and **BCL2**)))) | | |
| | Prot | | *BIM*'s pro-apoptotic activity requires expression driven by *FoxO3* and aided by *GSK3*, as well as the absence of *ERK*, any of the three inhibitory *BCL2* family proteins, or *ZEB1*. Alternatively, *Runx1* can induce BIM to apoptotic levels in the absence of *ERK* and *NF-κB*, assuming the three inhibitory *BCL2* family proteins are not all fully active. |
| | ⊢ Ind | ERK | The *MEK/ERK* pathway represses *BIM* protein levels, likely via transcriptional repression [539]. |
| | ← TR | FoxO3 | *FoxO3* is a transcriptional activator of *BIM* [540]. |
| | ← Ind | GSK3 | *GSK3* kinase is likely required for the *AP1*-dependent expression of *BIM* [541]. |
| | ⊢ TR | NfkB | *NF-κB* is a transcriptional repressor of *BIM* [542, 543]. |
| | ← TR | Runx1 | *TGF-β* stimulation leads to increased protein levels of pro-apoptotic *BIM* [544] via *Runx1* induction and subsequent *Foxo3/Runx1* driven transcription [545]. |
| | ⊢ TR | ZEB1 | *ZEB1* is a direct transcriptional repressor of the *BIM* promoter [546]. |
| | ⊢ IBind | MCL_1 | *MCL-1* binds *BIM* and inhibits its apoptotic activity [547]. |
| | ⊢ IBind | BCLXL | *BCL-xL* binds *BIM* and inhibits its apoptotic activity [529]. |
| | ⊢ IBind | BCL2 | *BLC2* binds *BIM* and inhibits its apoptotic activity [529]. |
| BID | **BID** = **Casp8** or (**Casp2** and (not((**BCL2** or **BCLXL**) or **MCL_1**))) | | |
| | Prot | | *BID* is truncated in response to *Caspase 8* activation. In addition, *Caspase 2* can also promote *BID* activation once all three pro-apoptotic *BCL2* family proteins are blocked. |
| | ← Lysis | Casp8 | In response to *TRAIL* (or *FAS* ligand), the initiator *Caspase 8* cleaves *BID* to its active truncated form [548, 549, 550]. |
| | ← Lysis | Casp2 | *Caspase 2* cleaves *BID* to its active truncated form [551]. |

**Table S1t: Apoptotic_SW module**

| | | | |
|---|---|---|---|
| | ⊣ IBind | MCL_1 | All three anti-apoptotic BCL2 proteins (*BCL2*, *BCL-xL* and *MCL-1*) sequesters *BID* into stable complexes, preventing them from activating *BAX* or *BAK* [552]. |
| | ⊣ IBind | BCLXL | All three anti-apoptotic BCL2 proteins (*BCL2*, *BCL-xL* and *MCL-1*) sequesters *BID* into stable complexes, preventing them from activating *BAX* or *BAK* [552]. |
| | ⊣ IBind | BCL2 | All three anti-apoptotic BCL2 proteins (*BCL2*, *BCL-xL* and *MCL-1*) sequesters *BID* into stable complexes, preventing them from activating *BAX* or *BAK* [552]. |
| BAK | **BAK** = (**BID**and((**BIM**or**BIK**)or(not((**BCL2**and**BCLXL**)and**MCL_1**))))or((**BIM**or **BIK**) and (not(**BCLXL** or **MCL_1**))) | | |
| | Prot | | Given that *BAK* is preferentially activated by *BID* compared to *BIM* [553] and that it is less responsive to sequestration by *BCL2* than the other two anti-apoptotic *BCL2* family proteins [554, 555], *BAK* in our model turns on when stimulated by *BID* if one or more *BCL2* family proteins are absent, or if *BIM* or *BIK* are also present. In contrast, *BIM* or *BIK* only activate *BAK* if *BCL-xL* and *MCL-1* are absent (*BCL-2* alone cannot block them). |
| | ⊣ IBind | MCL_1 | *MCL-1* binds *BAK* and prevent its oligomerization in the mitochondrial membrane [554, 555]. |
| | ⊣ IBind | BCLXL | *BCL-xL* binds *BAK* and prevent its oligomerization in the mitochondrial membrane [556, 554, 555]. |
| | ⊣ IBind | BCL2 | *BCL2* can also bind *BAK* to prevent its oligomerization, but it does so less potently than the other two BCL-2 family members [554, 555, 557]. |
| | ← Compl | BIK | *BIK* can aid the activation of both *BAK* and *BAX* by triggering *BAK* oligomerization on the ER membrane and promoting a $Ca^{2+}$ efflux required for the fragmentation of hyper fused mitochondrial tubules, aiding *BAK* and *BAX* activation [558]. |
| | ← Compl | BIM | *BAK* is preferentially activated by *BID* compared to *BIM*, but *BIM* can also promote BAK oligomerization [553]. |
| | ← Compl | BID | Activated (truncated) *BID* binds to mitochondrial *BAK*, resulting in its activation and oligomerization in the mitochondrial membrane, followed by *cytochrome c* release [559]. |
| BAX | **BAX** = (**BIM**and((**BID**or**BIK**)or(not((**BCL2**and**BCLXL**)and**MCL_1**))))or((**BID**or **BIK**) and (not(**BCL2** or **BCLXL**))) | | |
| | Prot | | In contrast to *BAK*, *BAX* is preferentially activated by *BIM* compared to *BID* [553] and it is less responsive to sequestration by *MCL-1* than the other two anti-apoptotic BCL2 family proteins [554, 555]. *BAX* in our model turns on when stimulated by *BIM* if one or more *BCL2* family proteins are absent, or if *BID* or *BIK* are also present. In contrast, *BID* or *BIK* only activate *BAK* if *BCL2* and *BCL-xL* are both absent (*MCL-1* alone cannot block them). |
| | ⊣ IBind | MCL_1 | *MCL-1* can also bind *BAK* to prevent its oligomerization, but it does so less potently than the other two BCL-2 family members [554, 555, 560]. |

**Table S1t: Apoptotic_SW module**

| | | | |
|---|---|---|---|
| | $\vdash$ IBind | BCLXL | *BCL-xL* binds *BAX* and prevent its oligomerization in the mitochondrial membrane [554, 555]. |
| | $\vdash$ IBind | BCL2 | *BCL2* binds *BAX* and prevent its oligomerization in the mitochondrial membrane [554, 555]. |
| | $\leftarrow$ Compl | BIK | *BIK* can aid the activation of both *BAK* and *BAX* by triggering *BAK* oligomerization on the ER membrane and promoting a $Ca^{2+}$ efflux required for the fragmentation of hyper fused mitochondrial tubules, aiding *BAK* and *BAX* activation [558]. |
| | $\leftarrow$ Compl | BIM | Activated *BIM* binds to mitochondrial *BAX*, resulting in its allosteric activation and oligomerization in the mitochondrial membrane, leading to *cytochrome c* release [553]. |
| | $\leftarrow$ Compl | BID | *BAX* is preferentially activated by *BIM* compared to *BID*, but *BID* can also promote *BAK* oligomerization [553]. |
| Cyto_C | **Cyto_C** = **BAX** or **BAK** | | |
| | Prot | | *Cytochrome C* release from mitochondria requires the oligomerization of either *BAK* or *BAX* [561]. |
| | $\leftarrow$ Loc | BAK | *BAK* oligomerization at the mitochondrial membrane triggers MOMP, which results in the release of *cytochrome C* from mitochondria [561]. |
| | $\leftarrow$ Loc | BAX | *BAX* oligomerization at the mitochondrial membrane triggers MOMP, which results in the release of *cytochrome C* from mitochondria [562, 563]. |
| SMAC | **SMAC** = **BAX** or **BAK** | | |
| | Prot | | *SMAC/Diablo* release from mitochondria requires the oligomerization of either *BAK* or *BAX* [561, 564]. |
| | $\leftarrow$ Loc | BAK | *BAK* oligomerization at the mitochondrial membrane triggers MOMP, which results in the release of *SMAC/Diablo* from mitochondria [561, 564]. |
| | $\leftarrow$ Loc | BAX | *BAX* oligomerization at the mitochondrial membrane triggers MOMP, which results in the release of *SMAC/Diablo* from mitochondria [561, 564]. |
| IAPs | **IAPs** = (not **SMAC**) or **AKT_H** | | |
| | Prot | | Inhibitor of Apoptosis Proteins (*IAP*s) are active in the absence of *SMAC* inhibition, or following *AKT_H* mediated upregulation (this protection from *SMAC* requires peak or oncogenic *AKT* activity). |
| | $\leftarrow$ Ind | AKT_H | *cIAP-2* and *XIAP* are both transcriptionally up-regulated in response to strong *PI3K/AKT1* activation [565]. |
| | $\vdash$ IBind | SMAC | *SMAC/Diablo* binds tightly to *IAP* proteins and blocks their ability to inhibit *Caspase 3* [566]. |
| Casp9 | **Casp9** = **Casp3** or ((not **IAPs**) and **Cyto_C**) | | |

**Table S1t: Apoptotic_SW module**

| | | | |
|---|---|---|---|
| | PTase | *Procaspase 9* is cleaved into active *Caspase 9* by *Caspase 3*, or by the apoptosome (which relies on *cytochrome C* for its assembly) in the absence of *IAP* proteins. | |
| | ← Compl | Cyto_C | *Cytochrome c* binds to *APAF-1* proteins, promoting their assembly into the apoptosome, a platform for *procaspase 9* binding and cleavage into its active form [567]. |
| | ⊢ IBind | IAPs | *XIAP*, *cIAP1* and *cIAP2* inhibit the *cytochrome c*-induced activation of *procaspase-9* [568]. |
| | ← Lysis | Casp3 | *Procaspase 9* is a direct cleavage target of *Caspase 3* [510]. |
| Casp3 | **Casp3** = ((**Casp9** and **Casp8**) or (**Casp3** and (**Casp9** or **Casp8**))) or ((not **IAPs**) and ((**Casp9** or **Casp8**) or **Casp3**)) | | |
| | PTase | Activation of *Caspase 3* requires proteolytic cleavage of *procaspase-3* by initiator caspases such as *Caspase 9* or *Caspase 8*. In our model, cooperation of two of the three caspases (*Casp9*, *Casp8*, *Casp3*) is required in the presence of *IAP*s, which inhibit the proteolytic activity of *Caspase 3* by bind tightly to its active site. In the absence of *IAP*s, either of the three caspases can cleave and activate *Caspase 3*. | |
| | ← Lysis | Casp8 | *Caspase 8* can cleave *Caspase 3* [569], but full *Caspase 3* activation also requires MOMP (potentially due to a need for *IAP* inhibition) [570]. |
| | ⊢ IBind | IAPs | *IAP*s bind tightly to the active site of *Caspase 3*, keeping its activity in check [568, 571]. |
| | ← Lysis | Casp9 | Active *Caspase 9* cleaves *procaspase 3* [572]. |
| | ← Per | Casp3 | Once activated, *Caspase 3* helps sustain its own activation by cleaving *procaspase 8* and *6*. *Caspase 6*, in turn, generates additional active *caspase 8* and *9*. Together they all sustains a continuing active pool of Caspase 3. |

**Table S1u: DNA_Fragmentation module**

| Target Node | Node Gate<br>Node Type<br>Link Type | Node Description<br>Input Node | Link Description |
|---|---|---|---|
| CAD | **CAD** = **Casp3** and **Casp9** | | |
| | DNase | Caspase-activated DNase (*CAD*) is activated when its inhibition is released via the cleavage of *ICAD* (inhibitor of caspase-activated DNase). While *Capsase 3* and *7* (a direct target of *Caspase 9*) can inhibit *ICAD* [573], in our model they are both required, as *CAD* = ON is represents terminal, irreversible apoptotic commitment, which is fully locked in when both *Caspase 3* and *9* are on. | |

**Table S1u: DNA_Fragmentation module**

| | | | |
|---|---|---|---|
| | $\underset{\text{Ind}}{\leftarrow}$ | Casp9 | In addition of *Caspase 3*, *CAD* inhibition can also be relieved by *ICAD* cleavage by *Caspase 7*, which is a direct target of *Caspase 9* [573]. |
| | $\underset{\text{Ind}}{\leftarrow}$ | Casp3 | *Caspase 3* relives *CAD* inhibition by cleaving its inhibitor *ICAD* [573]. |

**Table S2a: Key to Node Type Symbols**

| Symbol | Node Type | Description |
|---|---|---|
| Cell | Cell | Node type used when a node's state is used to represent discrete phenotypes of an entire cell; appropriate for multi models. |
| DM | DM_Switch | Multi-stable regulatory module (Dynamically Modular Switch) in a higher-level model layer. |
| Conn | Connector | Grouping of nodes that are either mono-stable, or do not form a switch-like circuit that controls discrete phenotype transitions. For example, signaling input layers without feedback, or multi-step links between distinct switches can be represented as connector nodes in coarse-grained (switch-level) models. |
| Env | Environment | Nodes or modules that represent the extracellular environment of a single cell or cell collective. These are self-sustaining nodes or node groups that maintain their initial states and receive no feedback from the rest of the network (they act as inputs). |
| Proc | Process | Nodes or modules that stand in for complex cellular processes not modeled in detail (e.g., DNA replication or the process of aligning chromosomes at the metaphase plane during mitosis). |
| MSt | Macro_Structure | Nodes or modules that represent the state of large, complex cellular structures such as DNA content, cytoskeletal features, junctions or mitochondria. |
| Met | Metabolite | Regulatory node representing a metabolite (not protein, gene product or complex structure). |
| mRNA | MRNA | mRNA. |
| miR | MicroRNA | microRNA. |
| PC | Protein_Complex | Protein complex represented by a single node or via a key member of the complex. |
| Rec | Receptor | Cell surface receptor protein or complex. |
| Adap | Adaptor_Protein | Protein that helps scaffold a signaling complex or other large assembly of proteins. |
| Secr | Secreted_Protein | Protein secreted into the extracellular environment, such that the state of the node tagged with this type represents the availability fo this protein outside the cell. |
| TF | TF_Protein | Transcription factor. |
| K | Kinase | Kinase (enzyme that catalyzes the phosphorylation of its target). |
| Ph | Phosphatase | Phosphatase (enzyme that catalyzes the removal of phosphorylation from its target). |
| UbL | Ubiquitin_Ligase | Ubiquitin ligase (protein that recruits an ubiquitin-conjugating enzyme that has been loaded with ubiquitin to a target protein and assists or directly catalyzes the transfer of ubiquitin from the ubiquitin-conjugating enzyme to the target). |
| PTase | Protease | Protease (enzyme that catalyzes the breakdown of proteins into smaller fragments). |
| DNase | DNase | Deoxyribonuclease (DNase, for short); endonuclease that catalyzes the hydrolytic cleavage of the DNA backbone. |

**Table S2a: Key to Node Type Symbols**

| | | |
|---|---|---|
| CAM | CAM | Cell adhesion proteins located on the cell surface. |
| CDK | CDK | Cyclin-dependent kinase. |
| CDKI | CDKI | Cyclin-dependent kinase inhibitor. |
| GEF | GEF | Guanine nucleotide exchange factor. |
| GAP | GAP | GTPase-activating protein (also called GTPase-accelerating protein). |
| GTPa | GTPase | GTPase enzymes that hydrolyze ATP to ADP. |
| Enz | Enzyme | Enzyme that does not fit the more specific enzyme categories listed above. |
| Prot | Protein | Regulatory protein that does not fit any of the more specific classifications listed above. |
| MP | Membrane_Potential | A relative measure of membrane potential across a biological membrane, generally indicating whether this potential is within the normal range, or abnormally low / high in a way that affects other regulatory processes. |
| lncRNA | LncRNA | Long intervening noncoding RNA |
| SLig | Cell_Surgace_Ligand | Membrane-bound signaling molecule that serves as a ligand to receptors on neighboring cells. |

**Table S2b: Key to Link Type Symbols**

| Symbol | Link Type | Description |
|---|---|---|
| Env | Enforced_Env | This link type represents self-loops on Environment nodes, which guarantee that these nodes maintain their initial state throughout a time-course simulation unless they are explicitly altered by the simulation's settings. |
| Ind | Indirect | Regulatory influence that does not involve direct binding, processing, or enzyme activity. |
| ComplProc | Complex_Process | Regulatory influence that is not modeled in detail, but involves more than one molecule or a macrostructure. For example, the physical need for kinetochores on replicated sister chromatids for the assembly of certain protein complexes can be represented as a link from the node representing kinetochores to the regulatory proteins, with a Complex_Process link type. |
| Per | Persistence | This link type represents self-loops that alter the ability of a node to stay in a particular state depending on its own current state. For example, if transcription of a protein is easier to maintain than to induce de novo, this may be encoded by a logic gate that includes the node itself and creates a self-loop. The link type of this loop is "Persistence". |
| TR | Transcription | Action of a transcription factor to alter the expression of the target node (mRNA or protein). Link type should be used for induction as well as repression (the link effect contains this information). |

**Table S2b: Key to Link Type Symbols**

| | | |
|---|---|---|
| TL | Translation | Regulatory influence that controls the translation of mRNA into protein; should be used for induction as well as repression of translation. |
| Ligand | Ligand_Binding | Binding of extracellular ligand to its receptor. |
| Compl | Complex_Formation | Binding even that leads to a regulatory protein complex. |
| IBind | Inhibitory_Binding | Binding even that represses the target node's level or activity. |
| Loc | Localization | Regulatory influence that alters the localization of a molecule. |
| BLoc | Binding_Localization | Binding even that alters the localization of a molecule. |
| PBind | Protective_Binding | Binding even that increases / protects the target node's activity. |
| Unbind | Unbinding | A regulatory influence that causes the target node to be released from a protein complex and change its activity (increase or decrease) as a result. |
| P | Phosphorylation | Phosphorylation. |
| DP | Dephosphorylation | Dehosphorylation. |
| PLoc | Phosphorylation_Localization | Phosphorylation resulting in altered protein localization. |
| Ubiq | Ubiquitination | Ubiquitination, usually leading to protein degradation. |
| Deg | Degradation | Regulatory influence leading to the degradation of the target molecule (more general than Ubiquitination; the latter link type should be used when appropriate). |
| GEF | GEF_Activity | Action of a Guanine nucleotide exchange factor (GEF) leading to GTP loading onto (and usually the activation of) a GTPase. |
| GAP | GAP_Activity | Actions of a GTPase-activating protein (GAP) leading to the hydrolysis of GTP by (and usually de-activation of) a GTPase. |
| Lysis | Proteolysis | Protein cleavage. |
| Cat | Catalysis | Increasing the rate of metabolite production by an enzyme. |
| Epi | Epigenetic | Process that alters gene expression via modifying chromatin condensation or altering DNA methylation. |
| TrConf | Transcription_Conflict | |
| Secr | Secretion | Secretion or shedding of a protein or other regulatory molecule to the extracellular environment. |
| RNAi | RNAi | This process represents inhibitory binding of cytoplasmic mRNAs by RISC-bound microRNAs that block translation and/or enhance mRNA degradation. |
| Acet | Acetylation | Acetylation |
| Deacet | Deacetylation | Deacetylation |
| -OH | Hydroxylation | Hydroxylation |

**Table S2c: Key to Link Effect Symbols**

| Symbol | Link Effect | Description |
|---|---|---|

**Table S2c: Key to Link Effect Symbols**

| | | |
|---|---|---|
| $\leftarrow$ | Activation | Link in which the input node aids the expression, activity, persistence or localization of the target such that the target is easier to turn/keep in an ON state. It can be used for multi-level nodes as long as these levels represent increasing intervals of activity. |
| $\vdash$ | Repression | Link in which the input node hinders the expression, activity, persistence or localization of the target such that the target is easier to turn/keep in an OFF state. It can be used for multi-level nodes as long as these levels represent increasing intervals of activity. |
| $\bullet\!-$ | Context_Dependent | Link acting on a node in such a way that it activates it under certain conditions (e.g., when another input is OFF), but represses it when this condition is not met. The XOR gate is a good example of a context-dependent link effect. |
| $\perp$ | Inapt | This link-type refers to connections between complex regulatory switches (rather than molecules), where categorizing the effect of an input as Activation or Repression, or even context-depended activation or repression does not apply. This is usually the case with multi-state switches, where the 3 or more phenotypes represented by the discrete states of these switches do not have a meaningful ordering. Thus, stating that this switch is "activated" by another one is not appropriate. |

[26] Mathew J. Joseph, Surabhi Dangi-Garimella, Mario A. Shields, Michelle E. Diamond, Limin Sun, Jennifer E. Koblinski, and Hidayatullah G. Munshi. Slug is a downstream mediator of transforming

growth factor-beta1-induced matrix metalloproteinase-9 expression and invasion of oral cancer cells. *Journal of Cellular Biochemistry*, 108(3):726–736, October 2009.

[27] Ming-Zhe Li, Jing-Jing Wang, Shi-Bin Yang, Wen-Feng Li, Long-Bin Xiao, Yu-Long He, and Xin-Ming Song. ZEB2 promotes tumor metastasis and correlates with poor prognosis of human colorectal cancer. *American Journal of Translational Research*, 9(6):2838–2851, 2017.

[28] Debashis Sarker, Ruth Plummer, Tim Meyer, Mikael H. Sodergren, Bristi Basu, Cheng Ean Chee, Kai-Wen Huang, Daniel H. Palmer, Yuk Ting Ma, T. R. Jeff Evans, Duncan R. C. Spalding, Madhava Pai, Rohini Sharma, David J. Pinato, James Spicer, Sarah Hunter, Vineet Kwatra, Joanna P. Nicholls, David Collin, Robert Nutbrown, Helen Glenny, Sonia Fairbairn, Vikash Reebye, Jon Voutila, Stephanie Dorman, Pinelopi Andrikakou, Peter Lloyd, Steve Felstead, Jenni Vasara, Robert Habib, Chris Wood, Pal Saetrom, Hans E. Huber, David C. Blakey, John J. Rossi, and Nagy Habib. MTL-CEBPA, a Small Activating RNA Therapeutic Upregulating C/EBP-$\alpha$, in Patients with Advanced Liver Cancer: A First-in-Human, Multicenter, Open-Label, Phase I Trial. *Clinical Cancer Research: An Official Journal of the American Association for Cancer Research*, 26(15):3936–3946, August 2020.

[209] Sagar R. Shah, Nathaniel D. Tippens, JinSeok Park, Ahmed Mohyeldin, Shuyan Wang, Guillermo Vela, Juan C. Martinez-Gutierrez, Seth S. Margolis, Susanne Schmidt, Shuli Xia, Andre Levchenko, and Alfredo Quiñones-Hinojosa. YAP controls cell migration and invasion through a Rho-GTPase switch. Preprint, Cancer Biology, April 2019.

[302] Chunsheng Kang. MicroRNA-200a suppresses the Wnt/$\beta$-catenin signaling pathway by interacting with $\beta$-catenin. *International Journal of Oncology*, December 2011.
