## Supplementary figures and images for "HHIP’s Dynamic Role in Epithelial Wound Healing Reveals a Potential Mechanism of COPD Susceptibility"

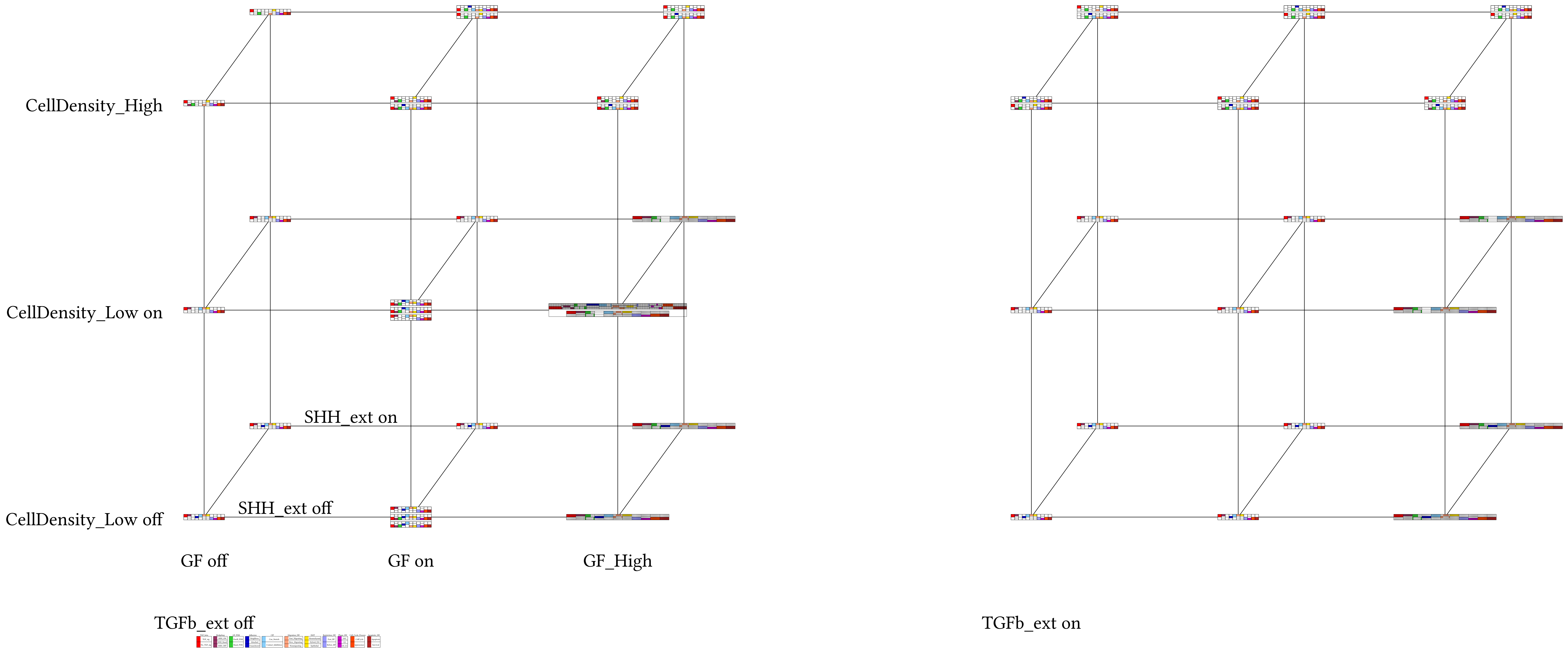

Supplement: Supplementary Material [file 611545_file02.zip › supplementary material/Supplementary File 2 - package to run dynmod - FINAL/COPD_EMT_CellCycle_Apoptosis_Attrs_Env_Space.pdf]

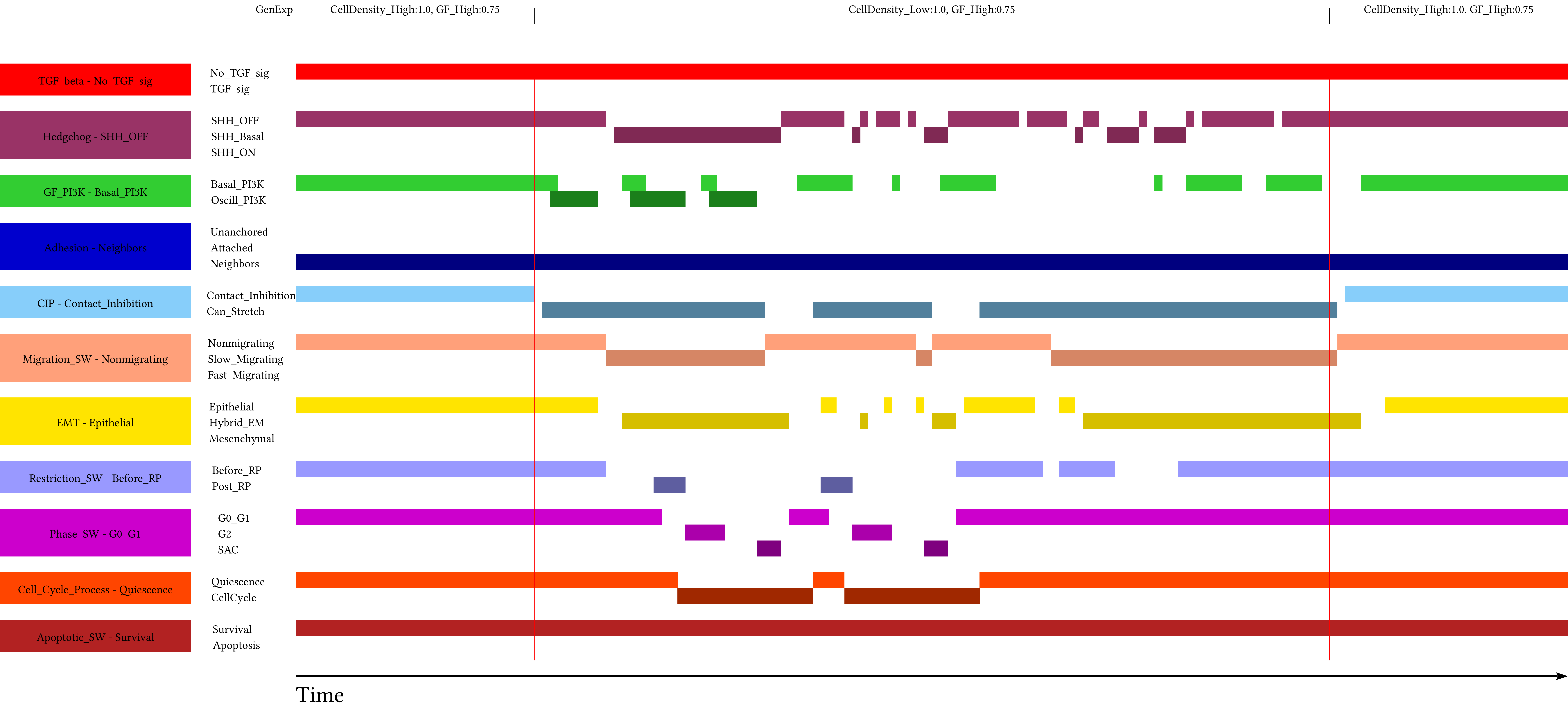

Supplement: Supplementary Material [file 611545_file02.zip › supplementary material/Supplementary File 2 - package to run dynmod - FINAL/_EXP/General_Time_Series/Fig_4A_WT_AV_short/PHTC/bc12101221211_Fig_4A_WT_AV_short.pdf]

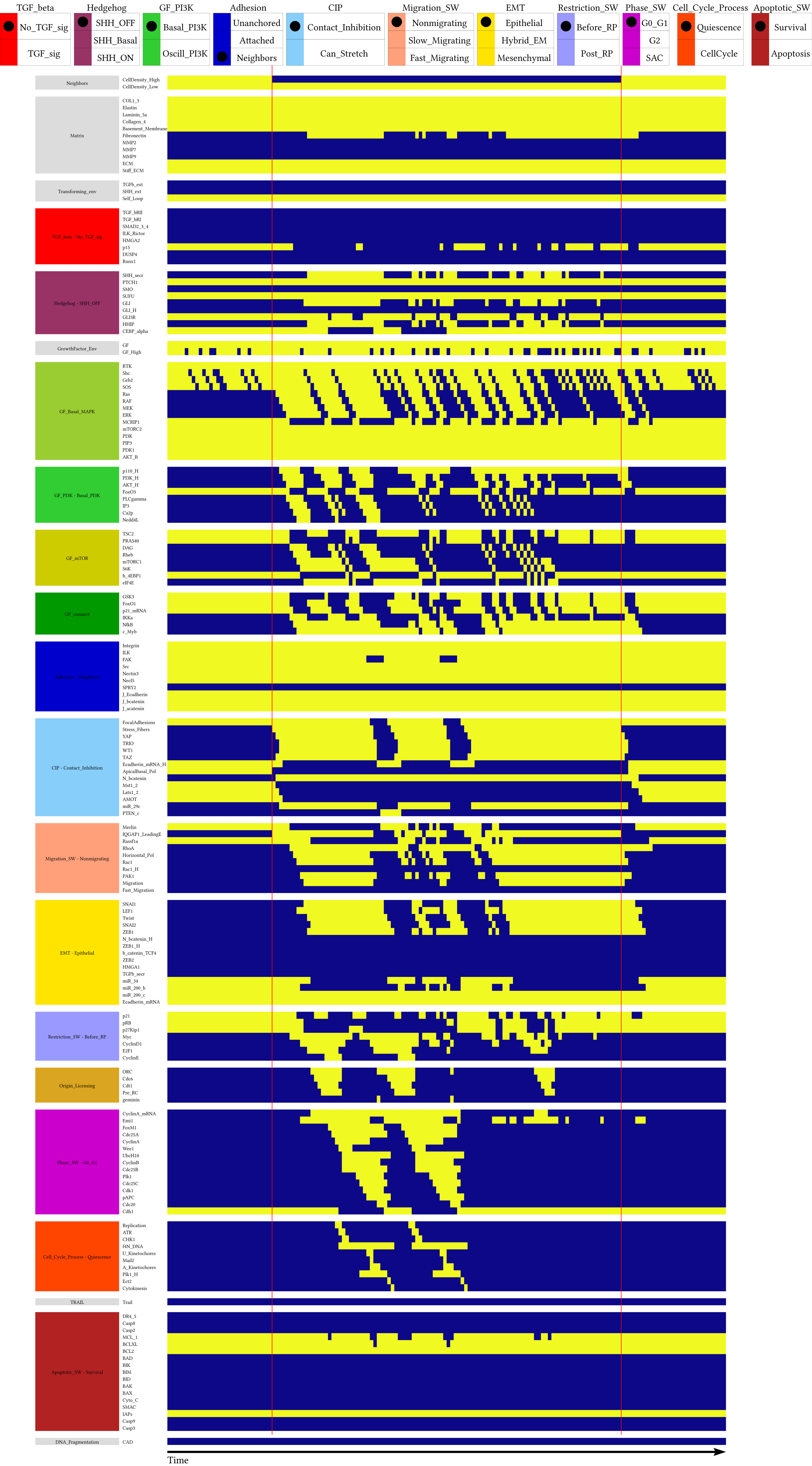

Supplement: Supplementary Material [file 611545_file02.zip › supplementary material/Supplementary File 2 - package to run dynmod - FINAL/_EXP/General_Time_Series/Fig_4A_WT_AV_short/NodeTC/bc12101221211_Fig_4A_WT_AV_short.pdf]

### Fig_4A_WT_AV_short

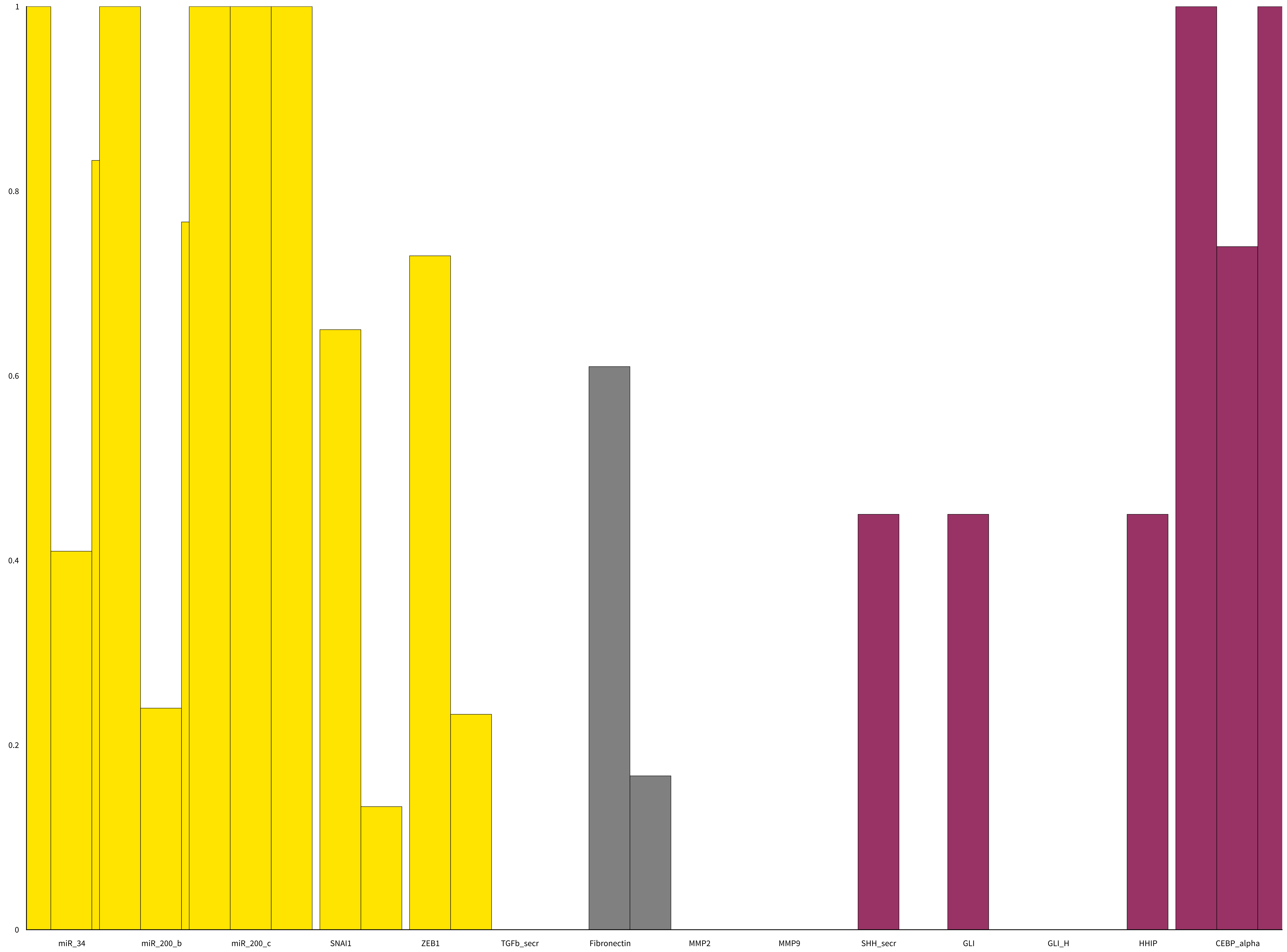

Supplement: Supplementary Material [file 611545_file02.zip › supplementary material/Supplementary File 2 - package to run dynmod - FINAL/_EXP/General_Time_Series/Fig_4A_WT_AV_short/NodeBCh/bc12101221211_Fig_4A_WT_AV_short.pdf]

### Fig_4A_WT_AV_short

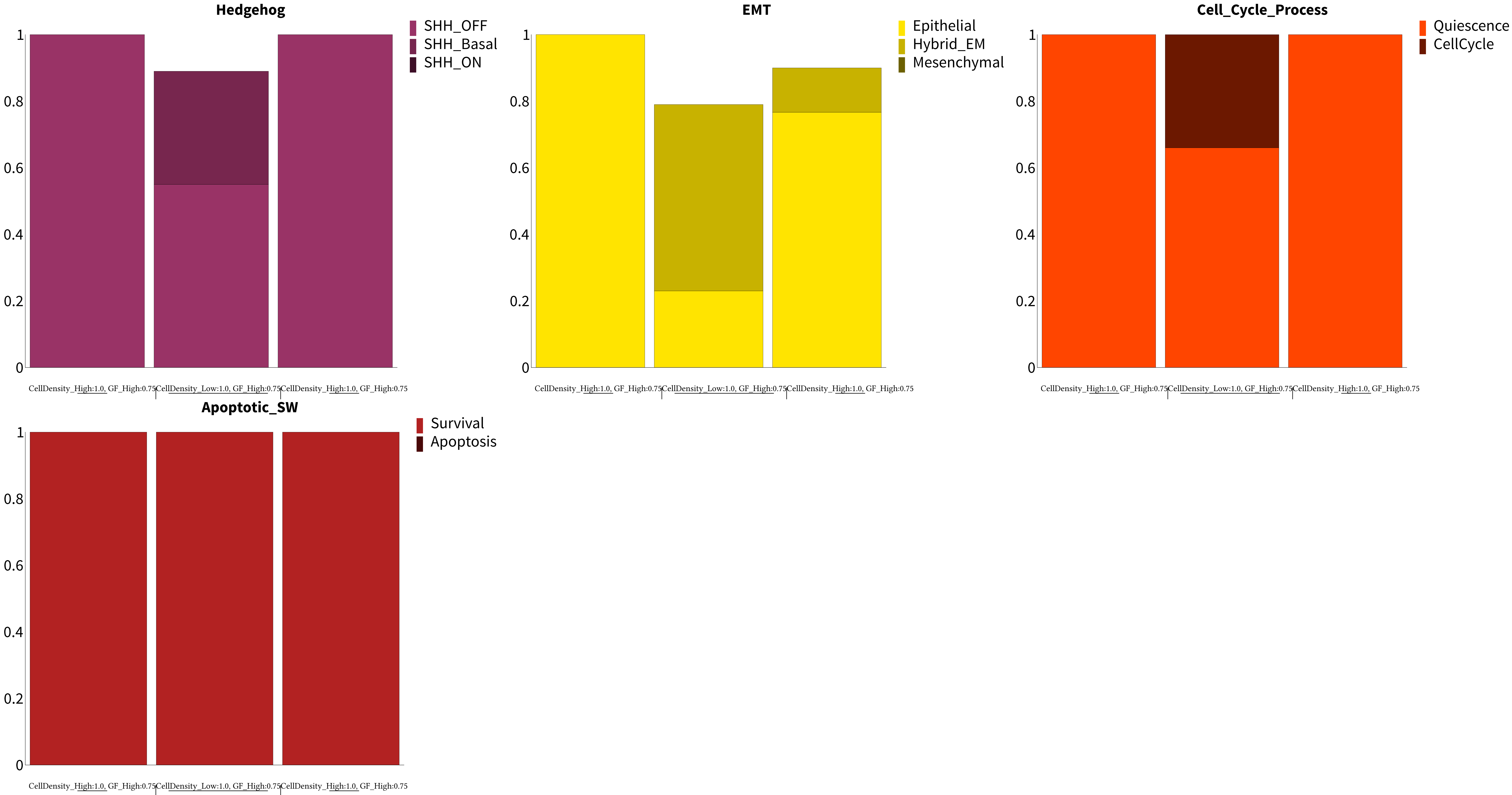

Supplement: Supplementary Material [file 611545_file02.zip › supplementary material/Supplementary File 2 - package to run dynmod - FINAL/_EXP/General_Time_Series/Fig_4A_WT_AV_short/PhBCh/bc12101221211_Fig_4A_WT_AV_short.pdf]

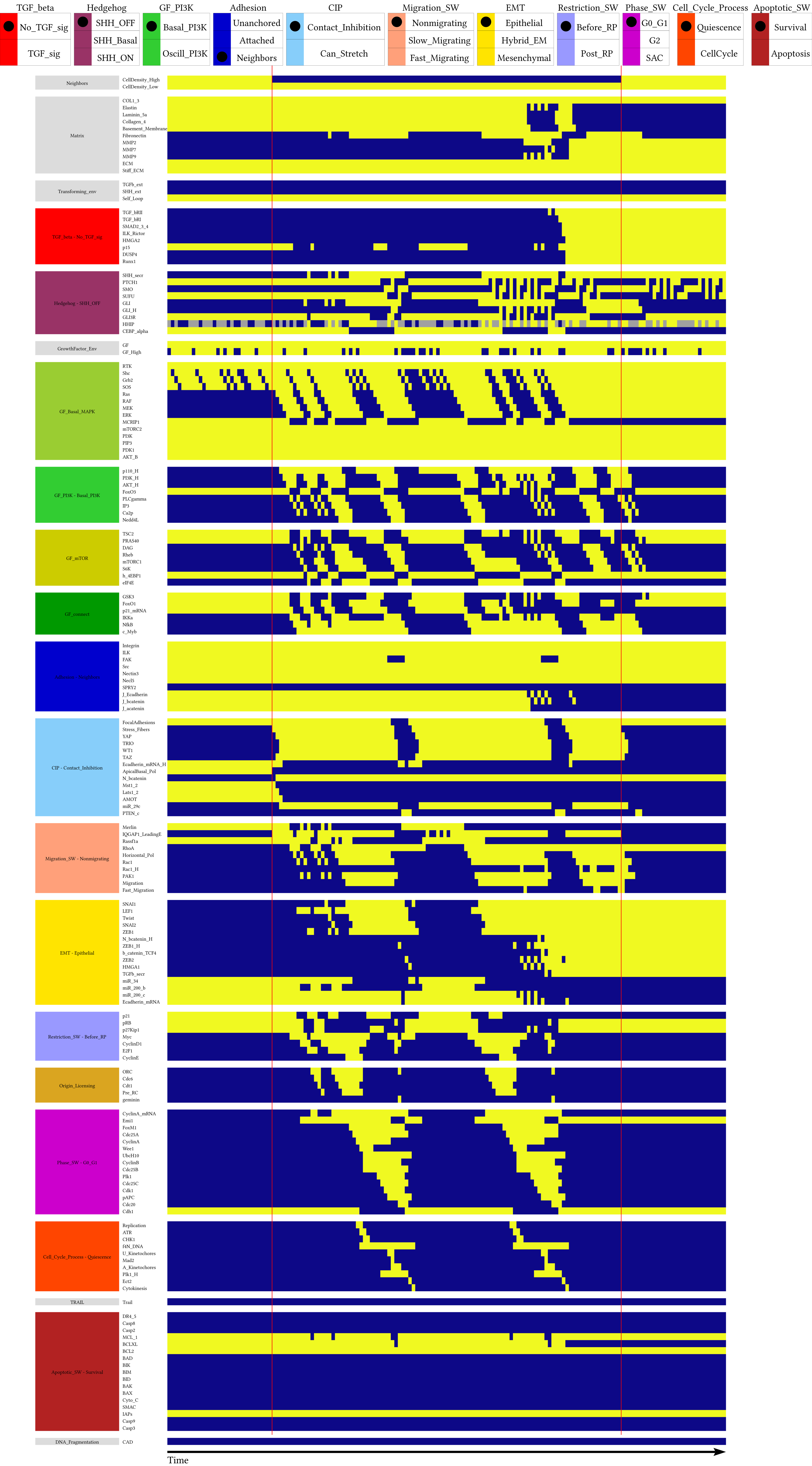

Supplement: Supplementary Material [file 611545_file02.zip › supplementary material/Supplementary File 2 - package to run dynmod - FINAL/_EXP/General_Time_Series/Fig_4B_HHIP_Haplo_AV_short/NodeTC/bc12101221211_Fig_4B_HHIP_Haplo_AV_short.pdf]

### Fig_4B_HHIP_Haplo_AV_short

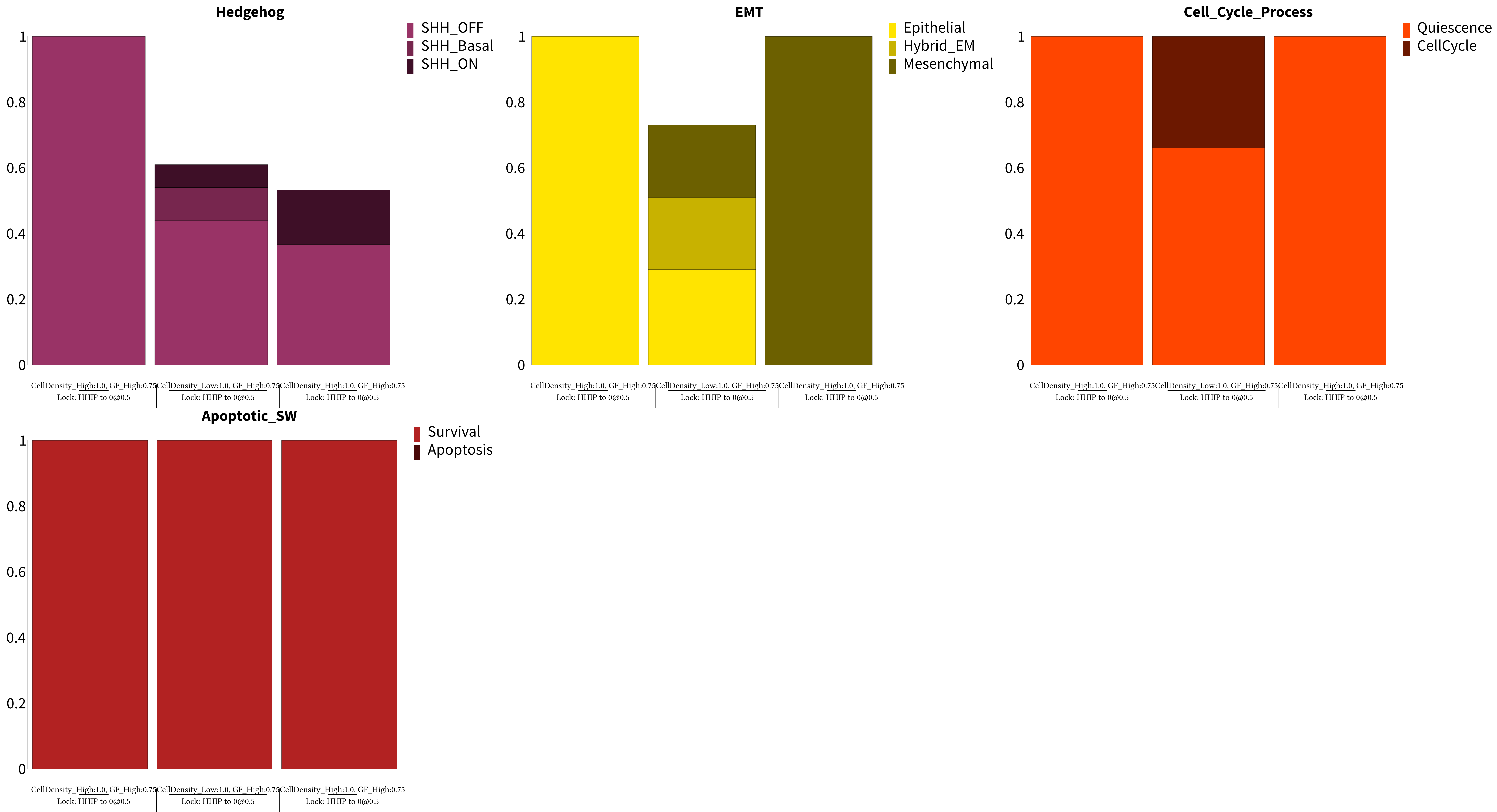

Supplement: Supplementary Material [file 611545_file02.zip › supplementary material/Supplementary File 2 - package to run dynmod - FINAL/_EXP/General_Time_Series/Fig_4B_HHIP_Haplo_AV_short/PhBCh/bc12101221211_Fig_4B_HHIP_Haplo_AV_short.pdf]

### Fig_4B_HHIP_Haplo_AV_short

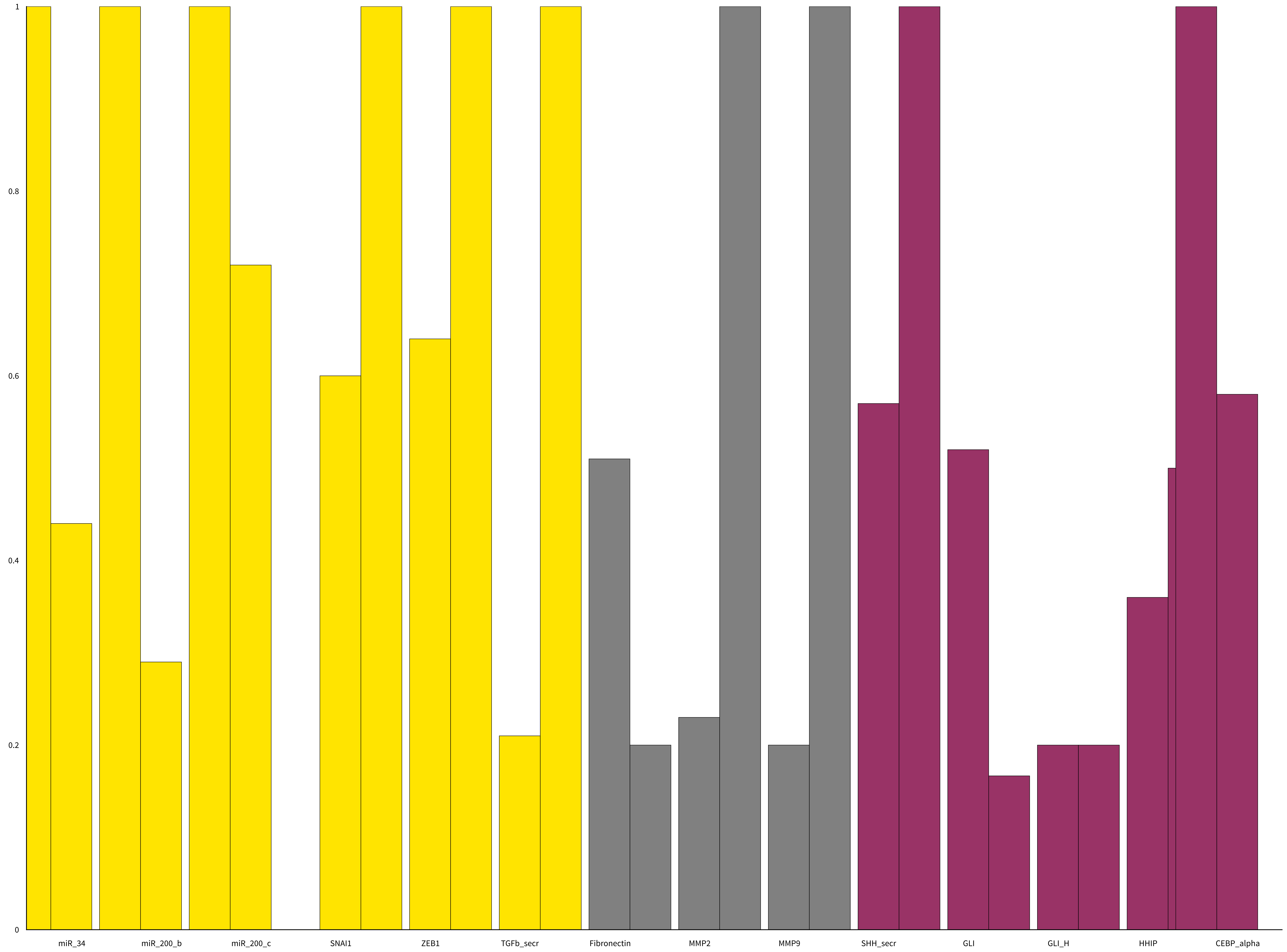

Supplement: Supplementary Material [file 611545_file02.zip › supplementary material/Supplementary File 2 - package to run dynmod - FINAL/_EXP/General_Time_Series/Fig_4B_HHIP_Haplo_AV_short/NodeBCh/bc12101221211_Fig_4B_HHIP_Haplo_AV_short.pdf]

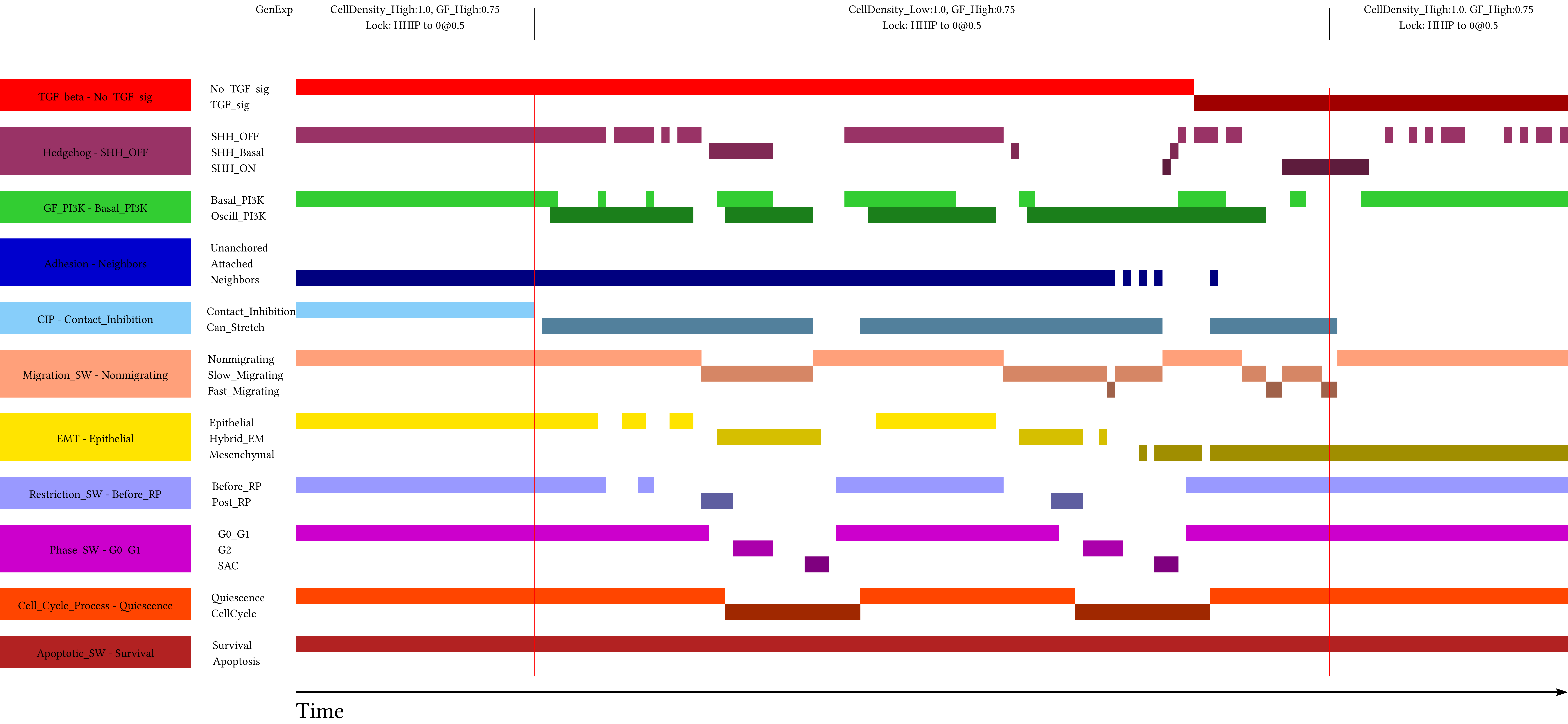

Supplement: Supplementary Material [file 611545_file02.zip › supplementary material/Supplementary File 2 - package to run dynmod - FINAL/_EXP/General_Time_Series/Fig_4B_HHIP_Haplo_AV_short/PHTC/bc12101221211_Fig_4B_HHIP_Haplo_AV_short.pdf]

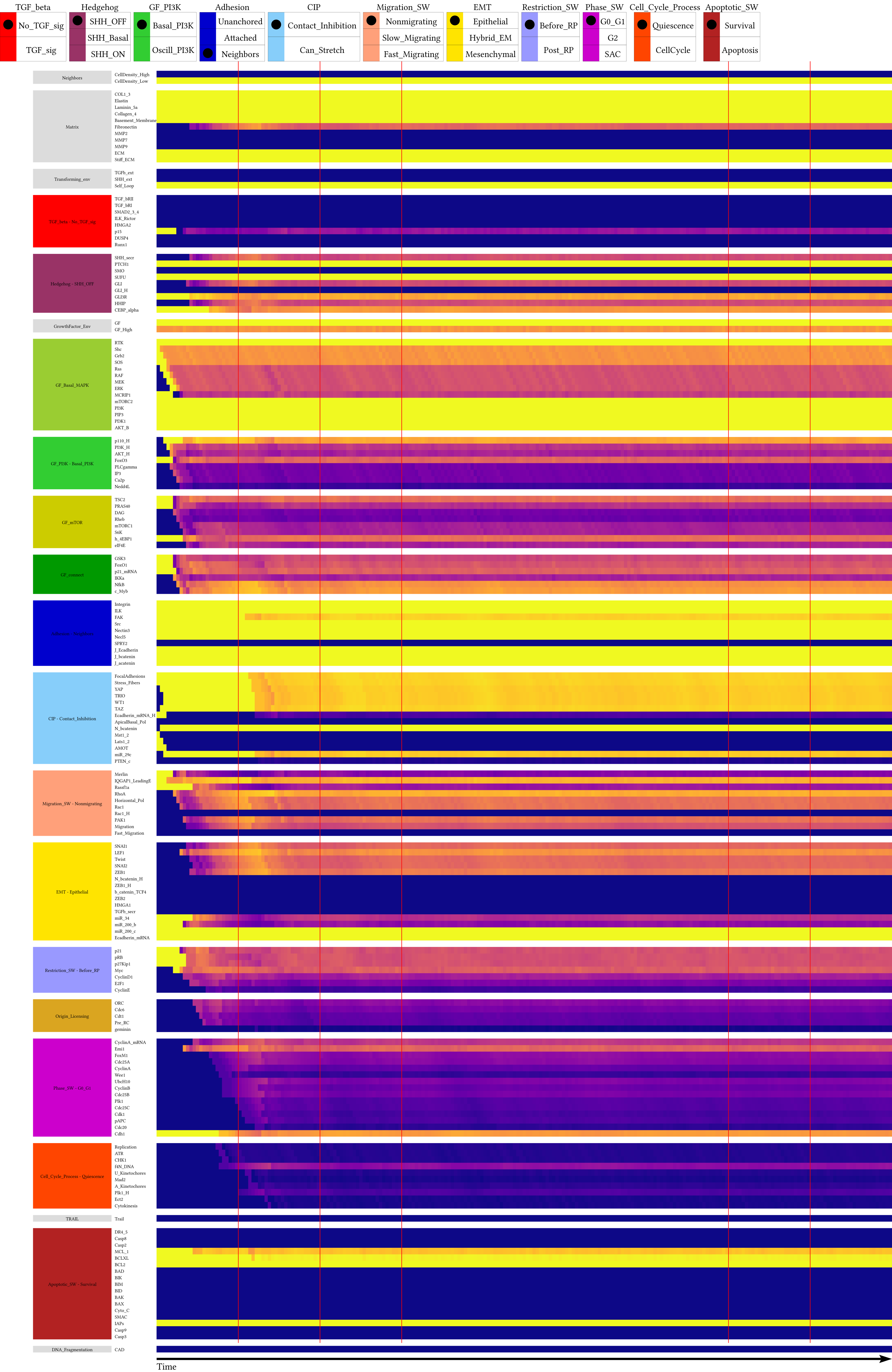

Supplement: Supplementary Material [file 611545_file02.zip › supplementary material/Supplementary File 2 - package to run dynmod - FINAL/_EXP/General_Time_Series/Fig_4C_WT_Time/NodeTC/bc12101221211_Fig_4C_WT_Time.pdf]

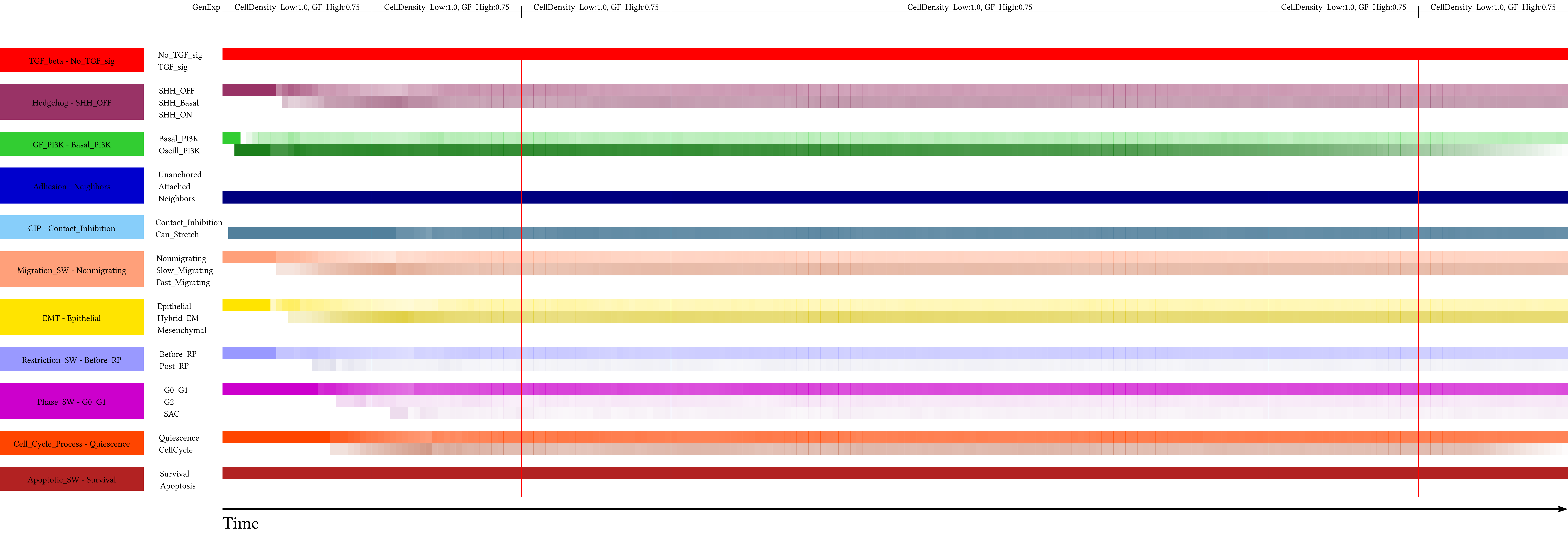

Supplement: Supplementary Material [file 611545_file02.zip › supplementary material/Supplementary File 2 - package to run dynmod - FINAL/_EXP/General_Time_Series/Fig_4C_WT_Time/PHTC/bc12101221211_Fig_4C_WT_Time.pdf]

### Fig_4C_WT_Time

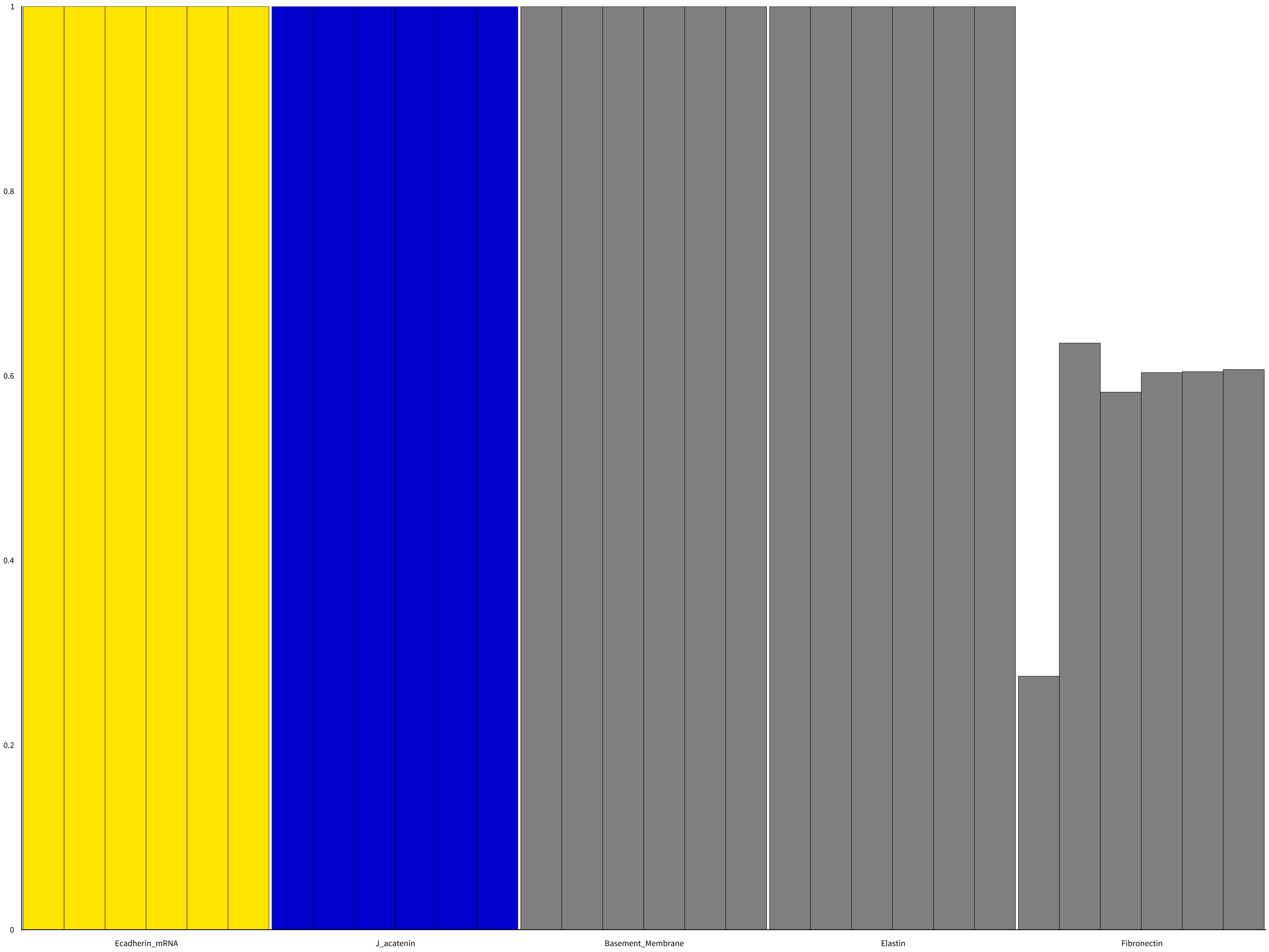

Supplement: Supplementary Material [file 611545_file02.zip › supplementary material/Supplementary File 2 - package to run dynmod - FINAL/_EXP/General_Time_Series/Fig_4C_WT_Time/NodeBCh/bc12101221211_Fig_4C_WT_Time.pdf]

### Fig_4C_WT_Time

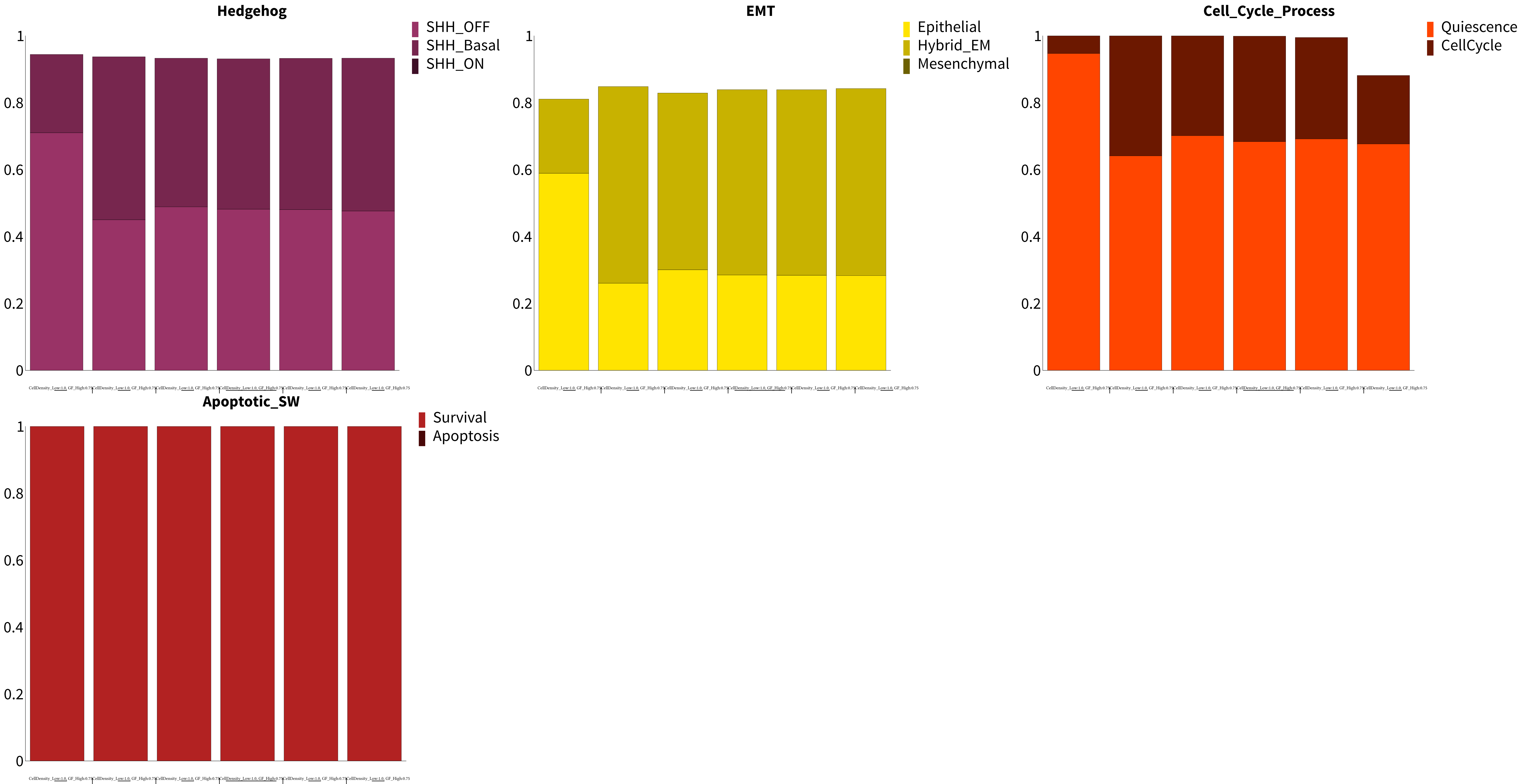

Supplement: Supplementary Material [file 611545_file02.zip › supplementary material/Supplementary File 2 - package to run dynmod - FINAL/_EXP/General_Time_Series/Fig_4C_WT_Time/PhBCh/bc12101221211_Fig_4C_WT_Time.pdf]

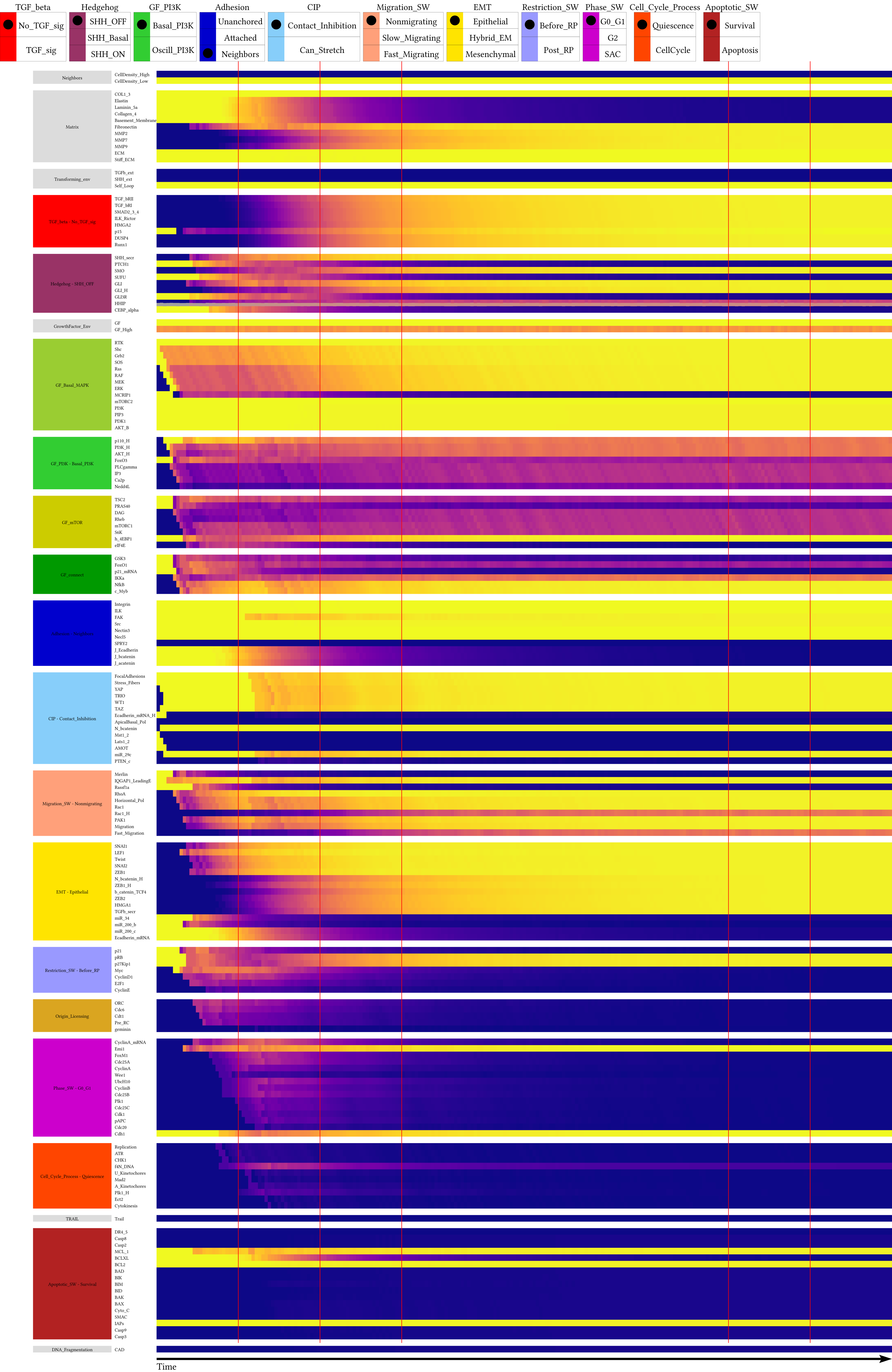

Supplement: Supplementary Material [file 611545_file02.zip › supplementary material/Supplementary File 2 - package to run dynmod - FINAL/_EXP/General_Time_Series/Fig_4C_HHIP_Haplo_Time/NodeTC/bc12101221211_Fig_4C_HHIP_Haplo_Time.pdf]

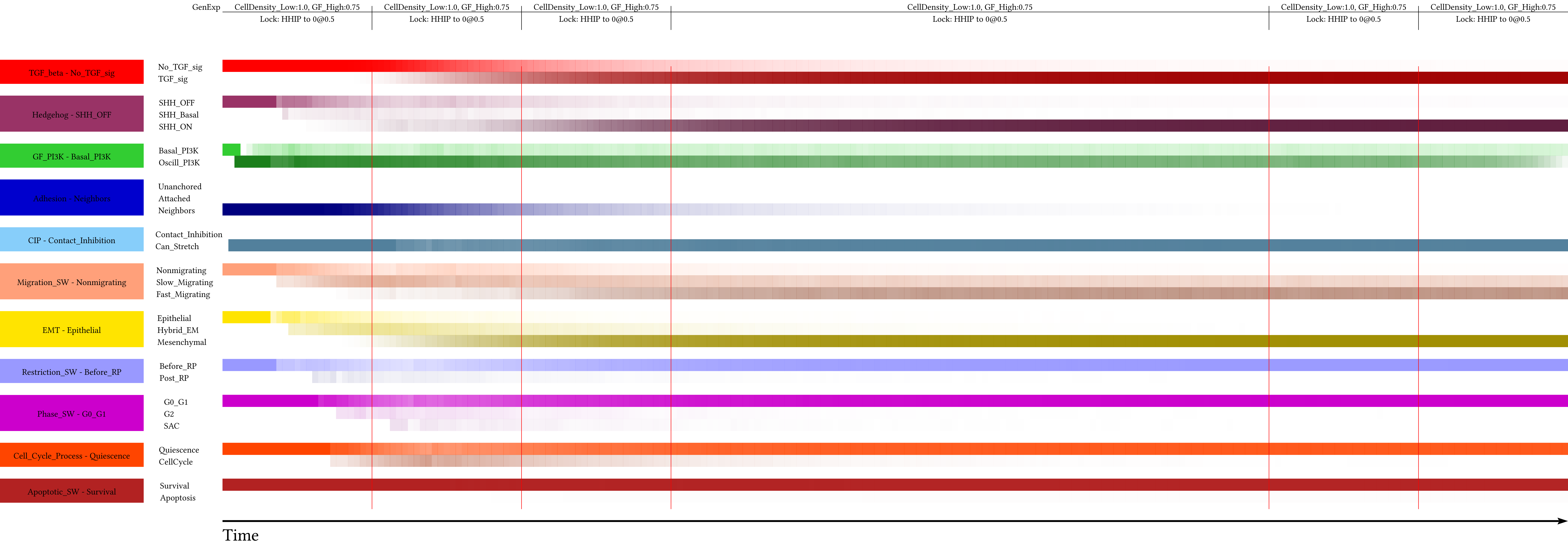

Supplement: Supplementary Material [file 611545_file02.zip › supplementary material/Supplementary File 2 - package to run dynmod - FINAL/_EXP/General_Time_Series/Fig_4C_HHIP_Haplo_Time/PHTC/bc12101221211_Fig_4C_HHIP_Haplo_Time.pdf]

### Fig_4C_HHIP_Haplo_Time

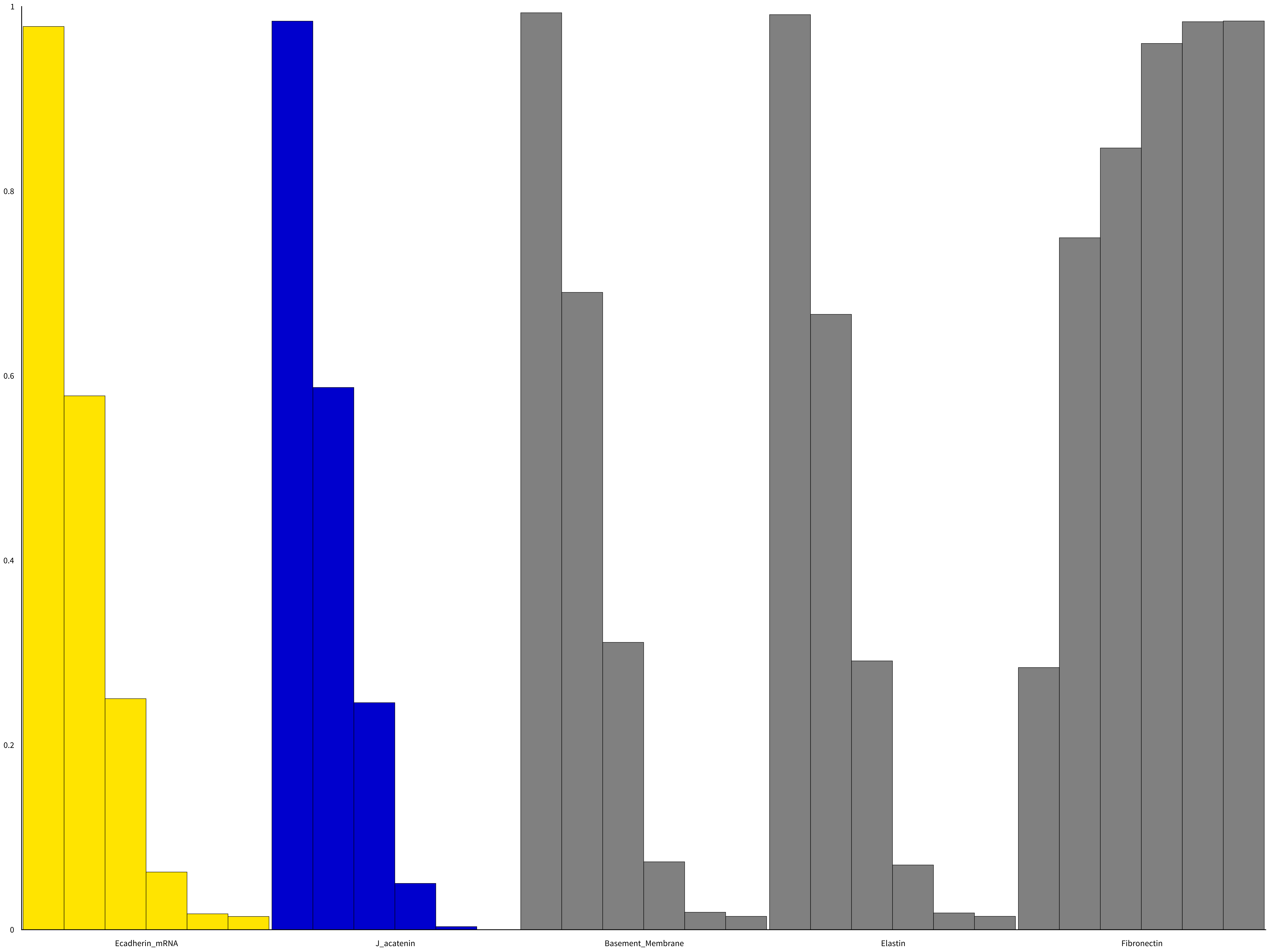

Supplement: Supplementary Material [file 611545_file02.zip › supplementary material/Supplementary File 2 - package to run dynmod - FINAL/_EXP/General_Time_Series/Fig_4C_HHIP_Haplo_Time/NodeBCh/bc12101221211_Fig_4C_HHIP_Haplo_Time.pdf]

### Fig_4C_HHIP_Haplo_Time

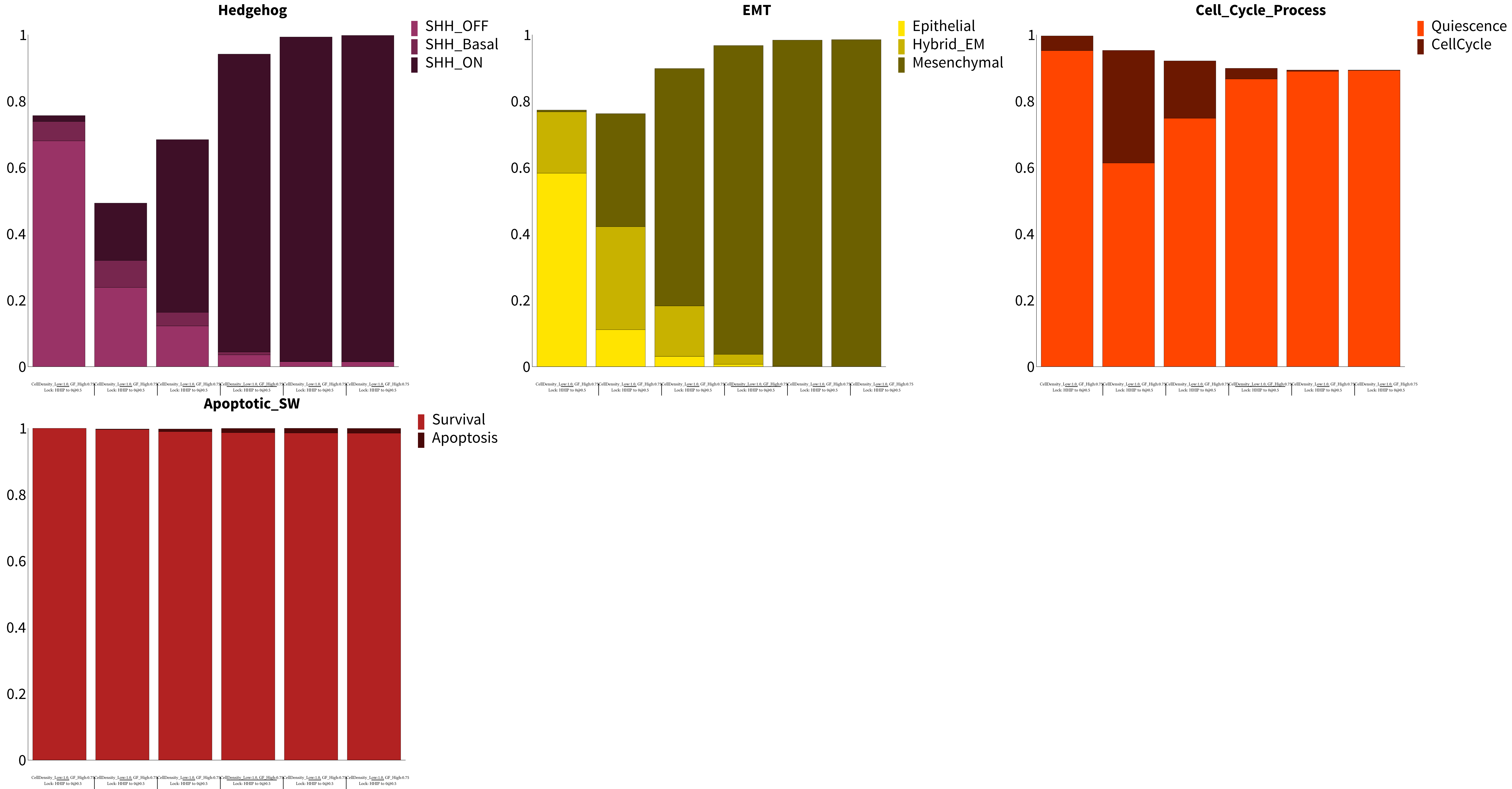

Supplement: Supplementary Material [file 611545_file02.zip › supplementary material/Supplementary File 2 - package to run dynmod - FINAL/_EXP/General_Time_Series/Fig_4C_HHIP_Haplo_Time/PhBCh/bc12101221211_Fig_4C_HHIP_Haplo_Time.pdf]

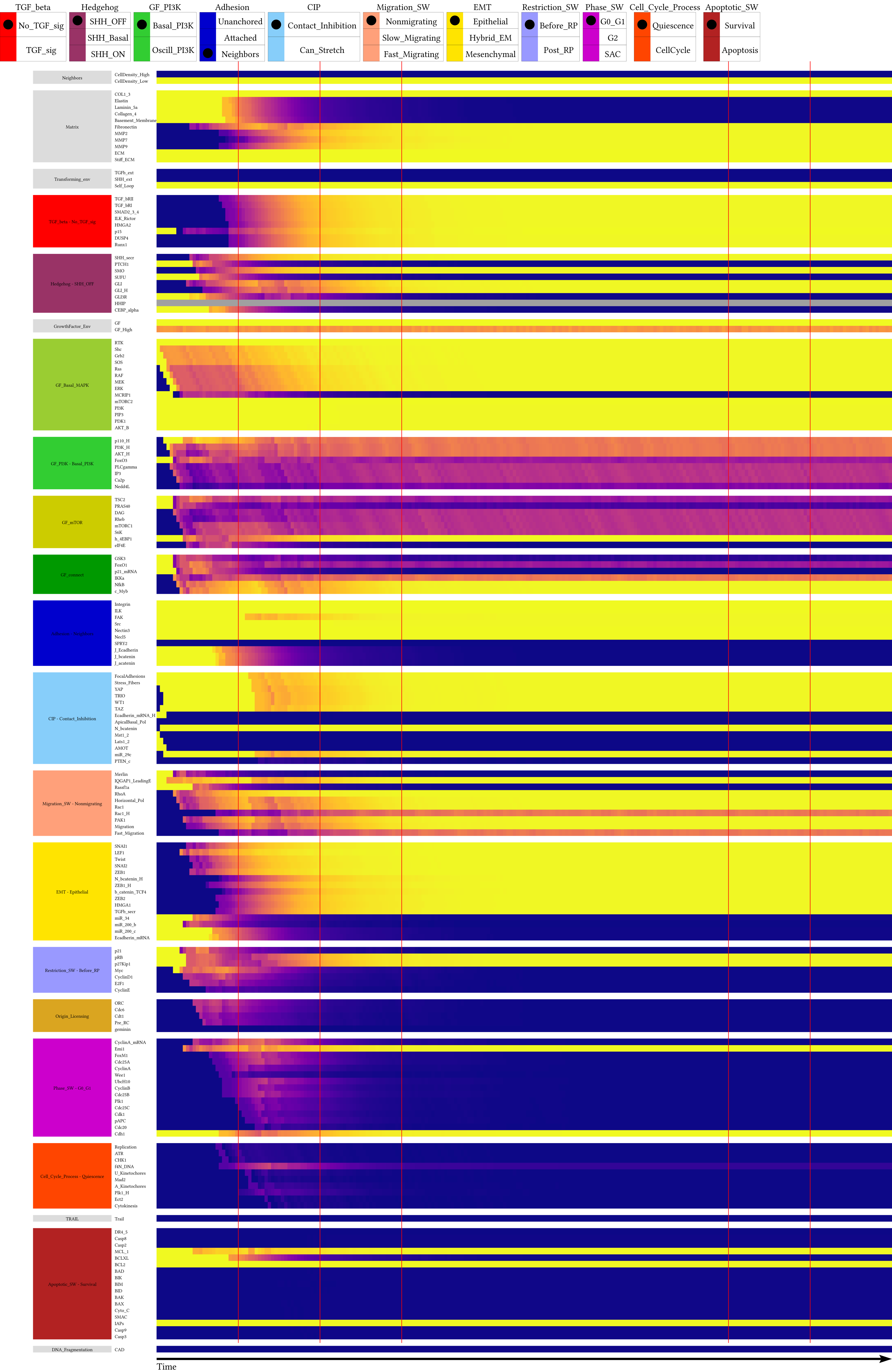

Supplement: Supplementary Material [file 611545_file02.zip › supplementary material/Supplementary File 2 - package to run dynmod - FINAL/_EXP/General_Time_Series/Fig_4C_HHIP_KO_Time/NodeTC/bc12101221211_Fig_4C_HHIP_KO_Time.pdf]

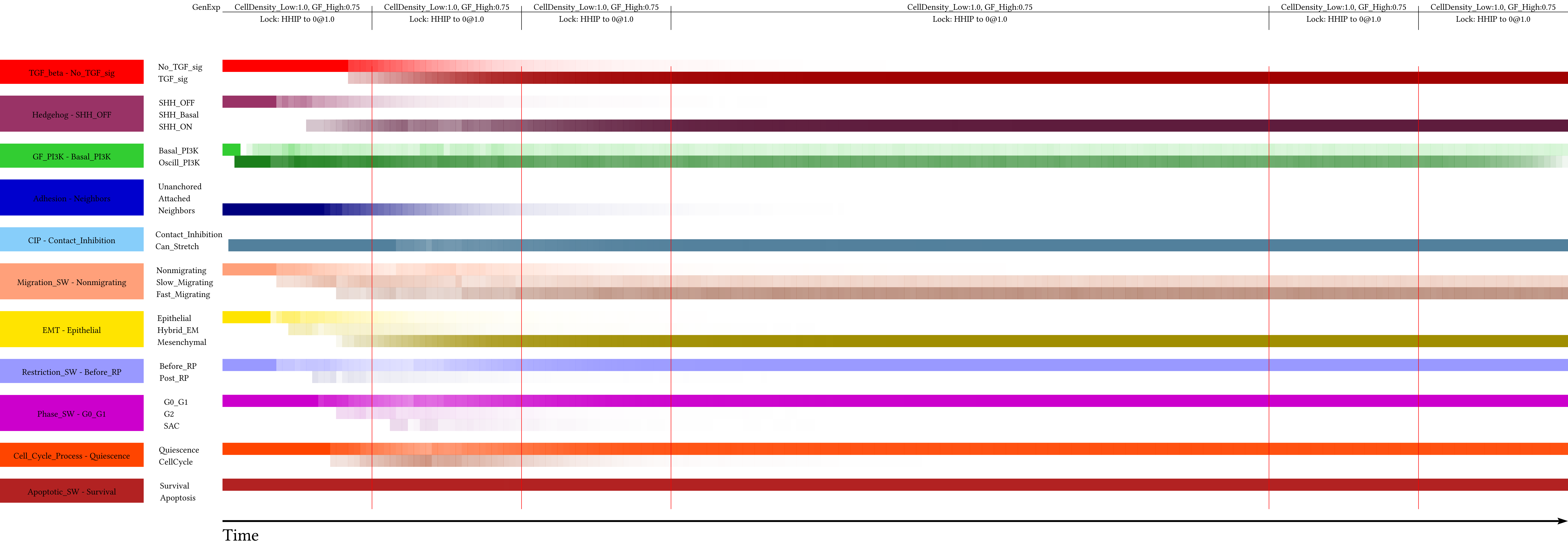

Supplement: Supplementary Material [file 611545_file02.zip › supplementary material/Supplementary File 2 - package to run dynmod - FINAL/_EXP/General_Time_Series/Fig_4C_HHIP_KO_Time/PHTC/bc12101221211_Fig_4C_HHIP_KO_Time.pdf]

### Fig_4C_HHIP_KO_Time

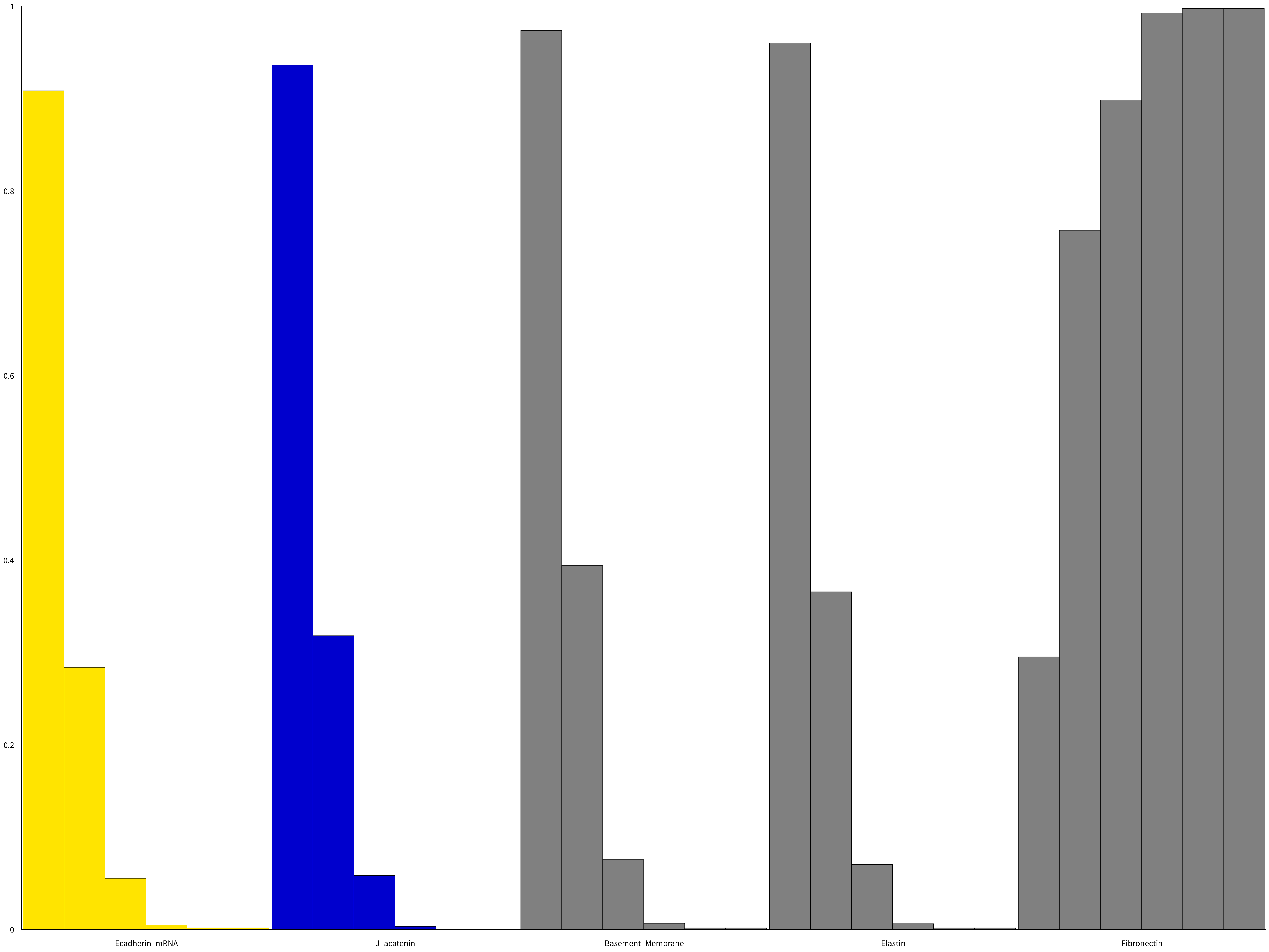

Supplement: Supplementary Material [file 611545_file02.zip › supplementary material/Supplementary File 2 - package to run dynmod - FINAL/_EXP/General_Time_Series/Fig_4C_HHIP_KO_Time/NodeBCh/bc12101221211_Fig_4C_HHIP_KO_Time.pdf]

### Fig_4C_HHIP_KO_Time

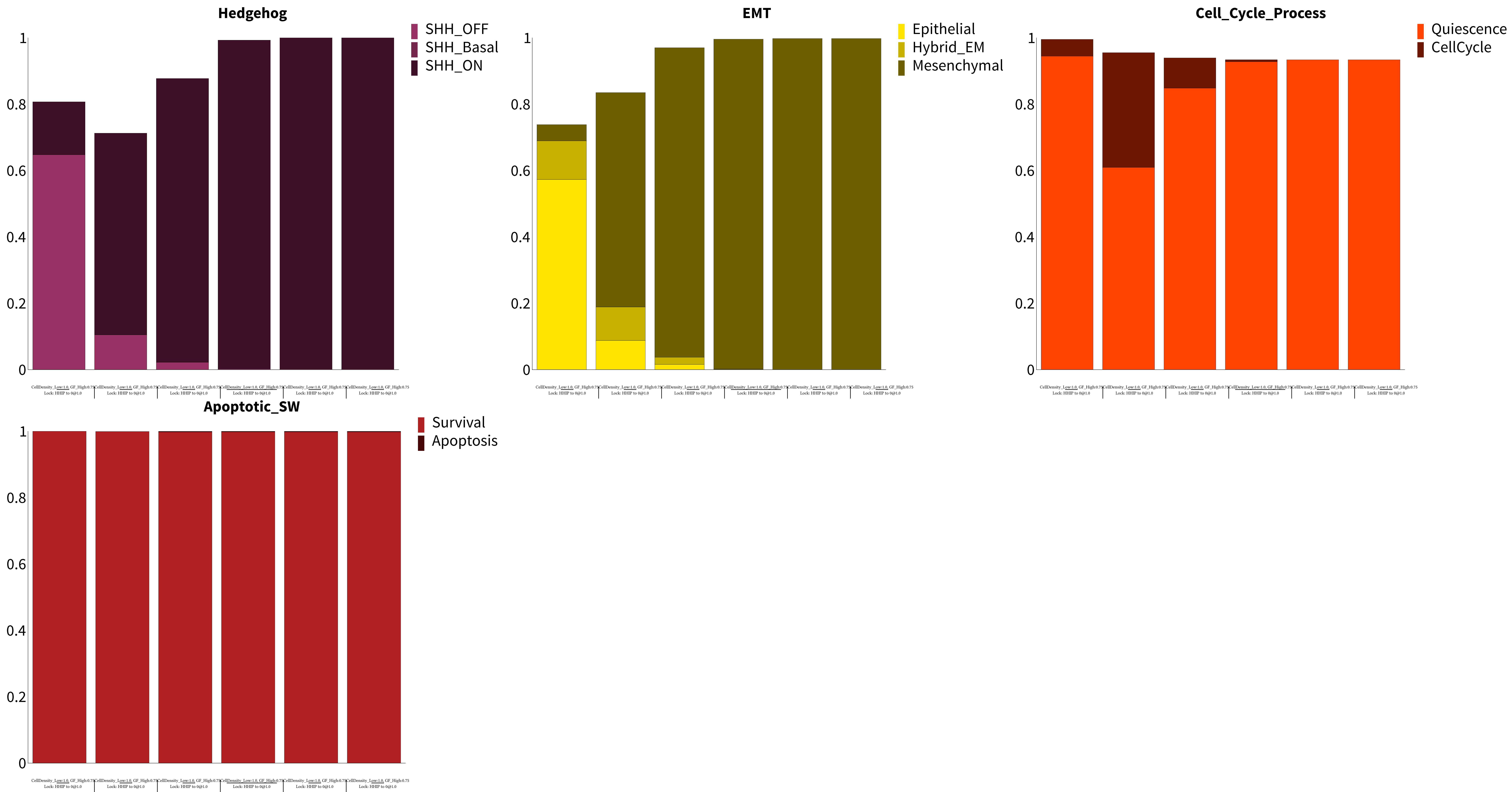

Supplement: Supplementary Material [file 611545_file02.zip › supplementary material/Supplementary File 2 - package to run dynmod - FINAL/_EXP/General_Time_Series/Fig_4C_HHIP_KO_Time/PhBCh/bc12101221211_Fig_4C_HHIP_KO_Time.pdf]

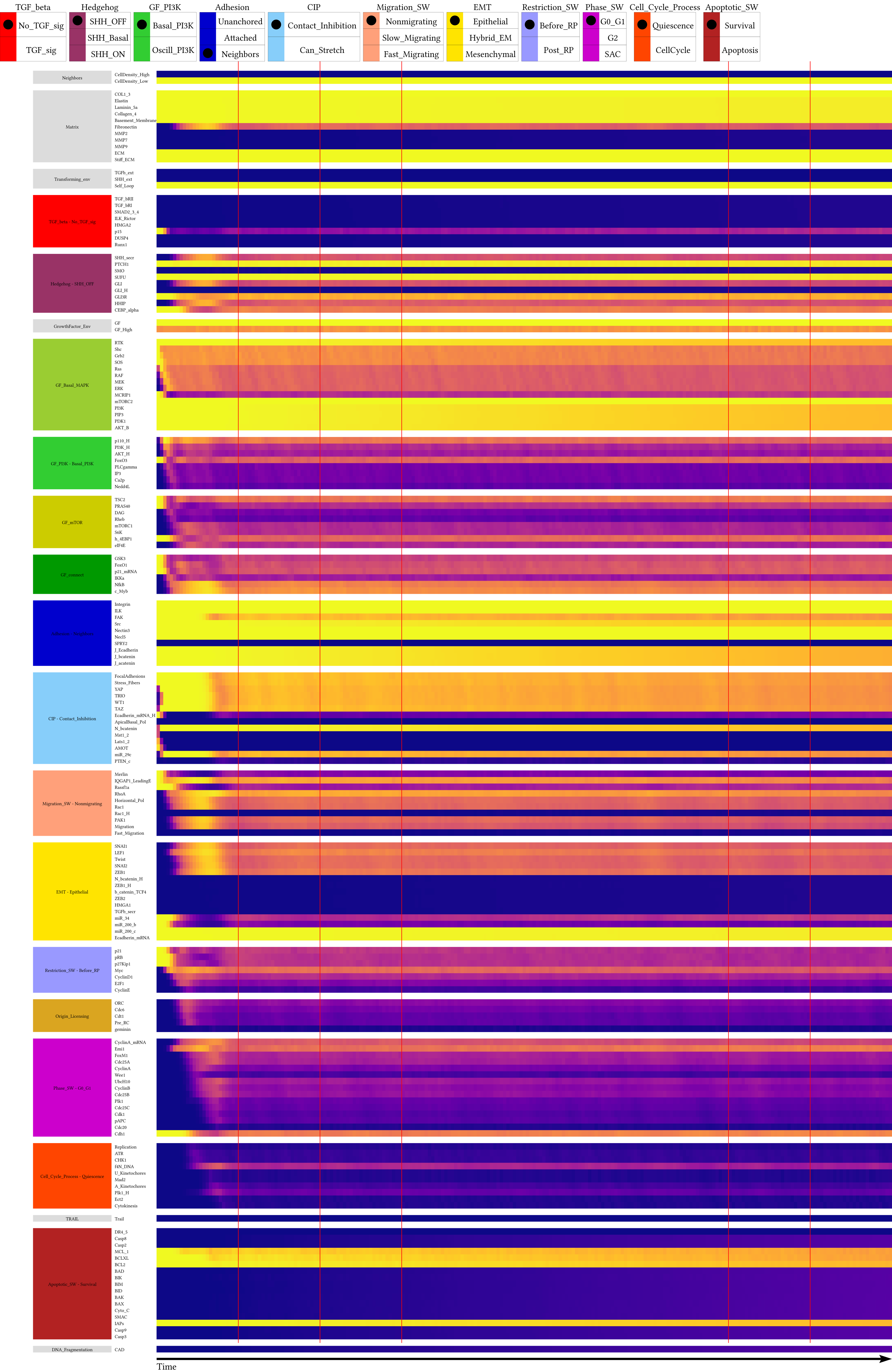

Supplement: Supplementary Material [file 611545_file02.zip › supplementary material/Supplementary File 2 - package to run dynmod - FINAL/_EXP/General_Time_Series/SM_Fig_4C_WT_Time_BAsync/NodeTC/bc12101221211_SM_Fig_4C_WT_Time_BAsync.pdf]

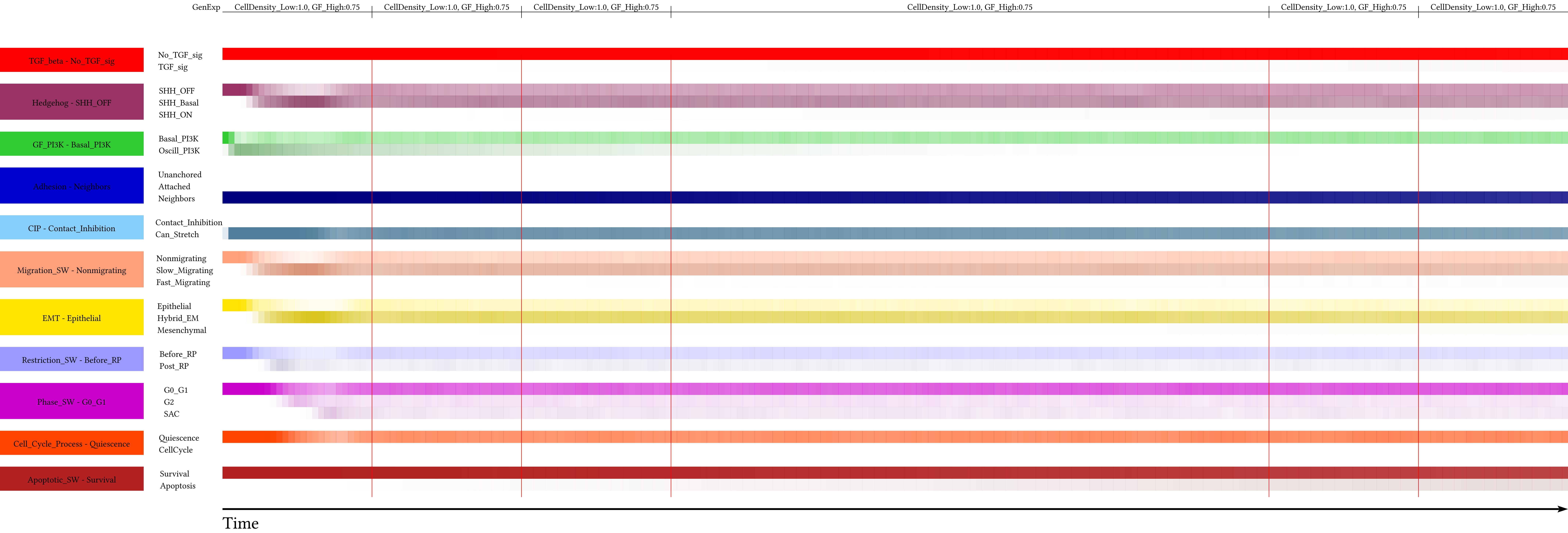

Supplement: Supplementary Material [file 611545_file02.zip › supplementary material/Supplementary File 2 - package to run dynmod - FINAL/_EXP/General_Time_Series/SM_Fig_4C_WT_Time_BAsync/PHTC/bc12101221211_SM_Fig_4C_WT_Time_BAsync.pdf]

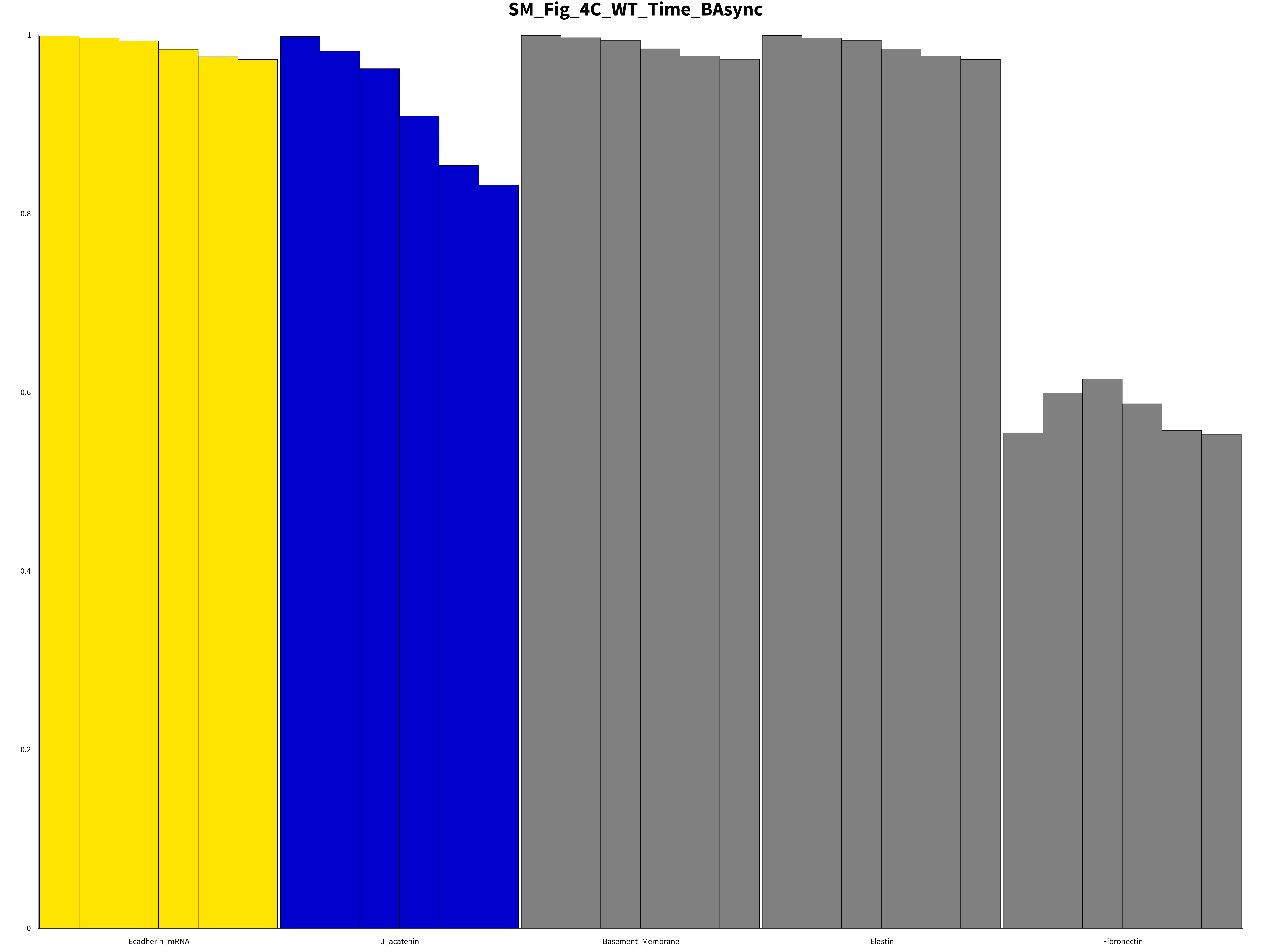

Supplement: Supplementary Material [file 611545_file02.zip › supplementary material/Supplementary File 2 - package to run dynmod - FINAL/_EXP/General_Time_Series/SM_Fig_4C_WT_Time_BAsync/NodeBCh/bc12101221211_SM_Fig_4C_WT_Time_BAsync.pdf]

### SM_Fig_4C_WT_Time_BAsync

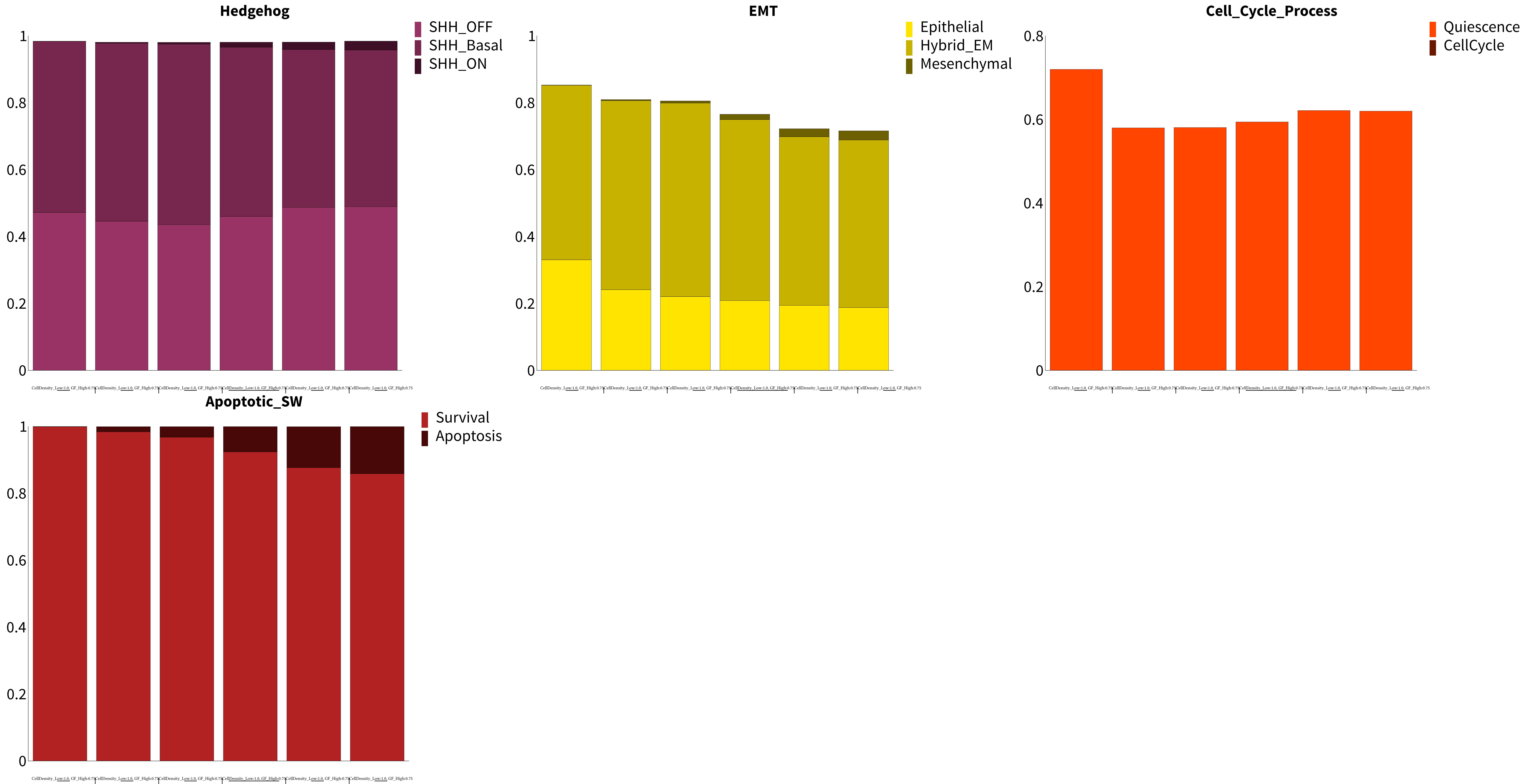

Supplement: Supplementary Material [file 611545_file02.zip › supplementary material/Supplementary File 2 - package to run dynmod - FINAL/_EXP/General_Time_Series/SM_Fig_4C_WT_Time_BAsync/PhBCh/bc12101221211_SM_Fig_4C_WT_Time_BAsync.pdf]

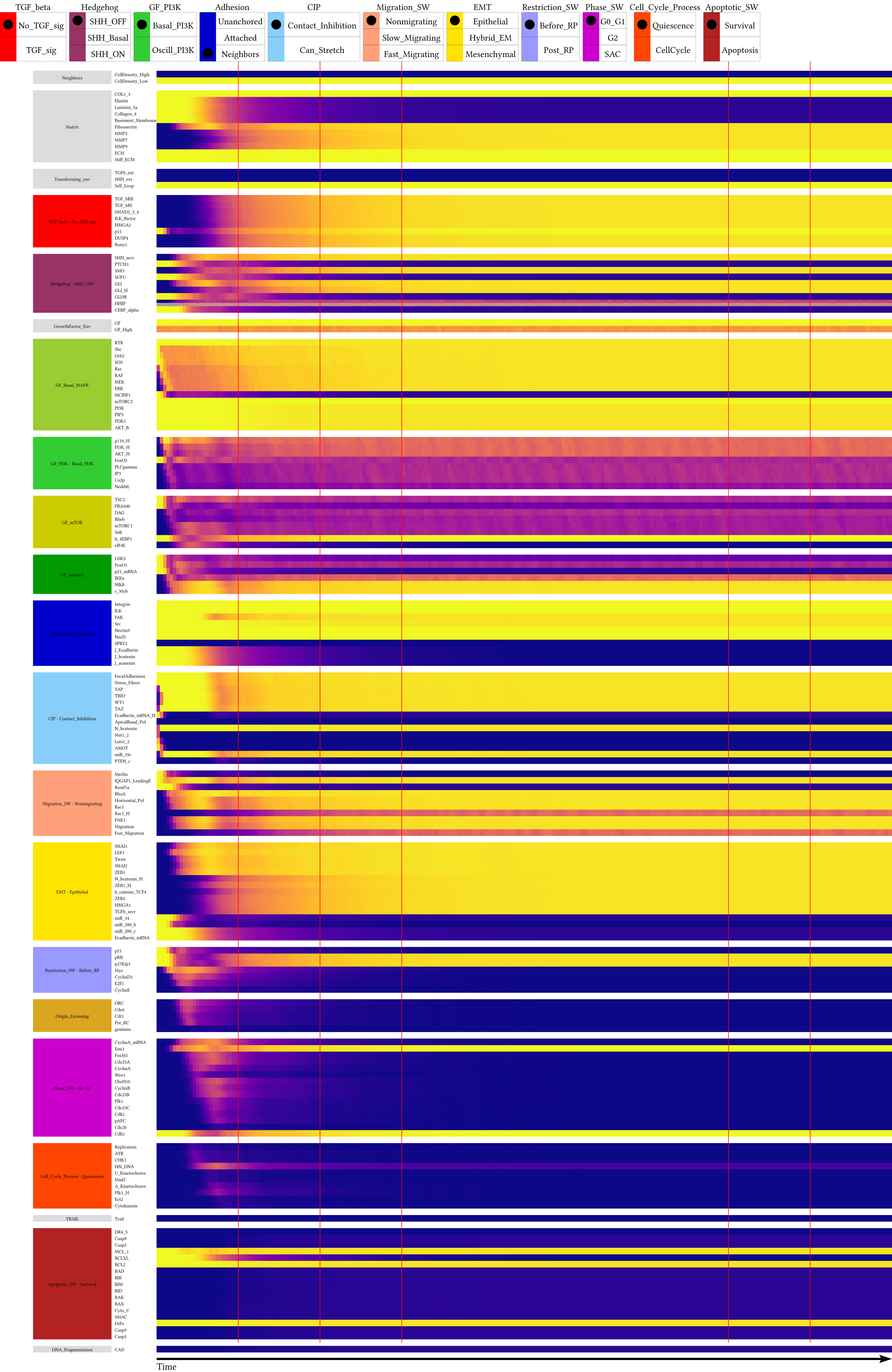

Supplement: Supplementary Material [file 611545_file02.zip › supplementary material/Supplementary File 2 - package to run dynmod - FINAL/_EXP/General_Time_Series/SM_Fig_4C_HHIP_Haplo_Time_BAsync/NodeTC/bc12101221211_SM_Fig_4C_HHIP_Haplo_Time_BAsync.pdf]

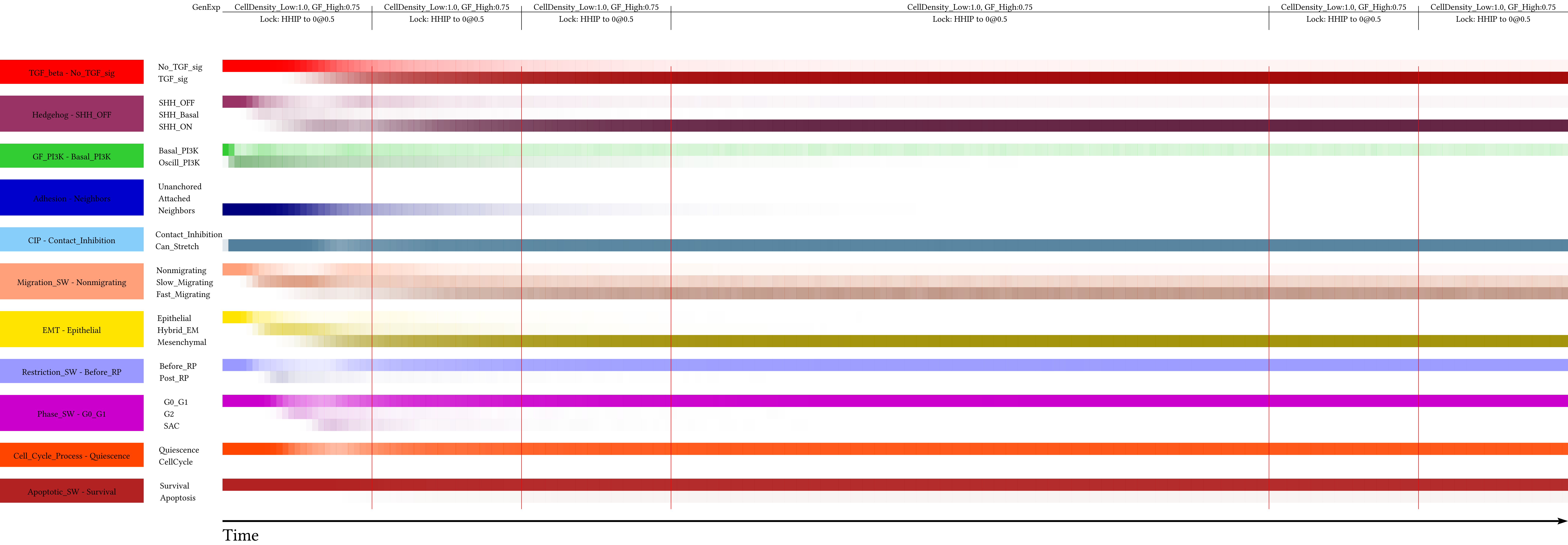

Supplement: Supplementary Material [file 611545_file02.zip › supplementary material/Supplementary File 2 - package to run dynmod - FINAL/_EXP/General_Time_Series/SM_Fig_4C_HHIP_Haplo_Time_BAsync/PHTC/bc12101221211_SM_Fig_4C_HHIP_Haplo_Time_BAsync.pdf]

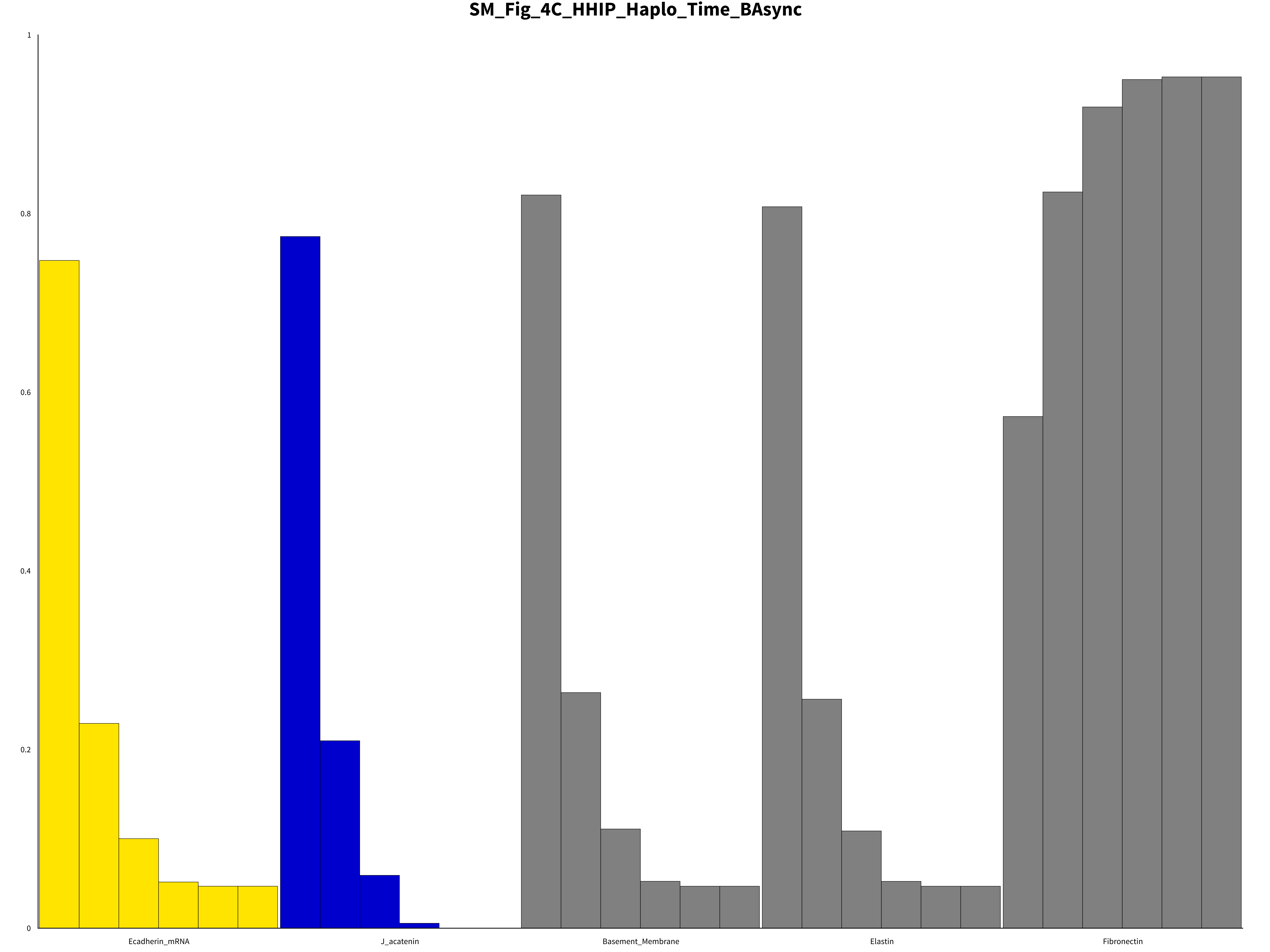

Supplement: Supplementary Material [file 611545_file02.zip › supplementary material/Supplementary File 2 - package to run dynmod - FINAL/_EXP/General_Time_Series/SM_Fig_4C_HHIP_Haplo_Time_BAsync/NodeBCh/bc12101221211_SM_Fig_4C_HHIP_Haplo_Time_BAsync.pdf]

### SM_Fig_4C_HHIP_Haplo_Time_BAsync

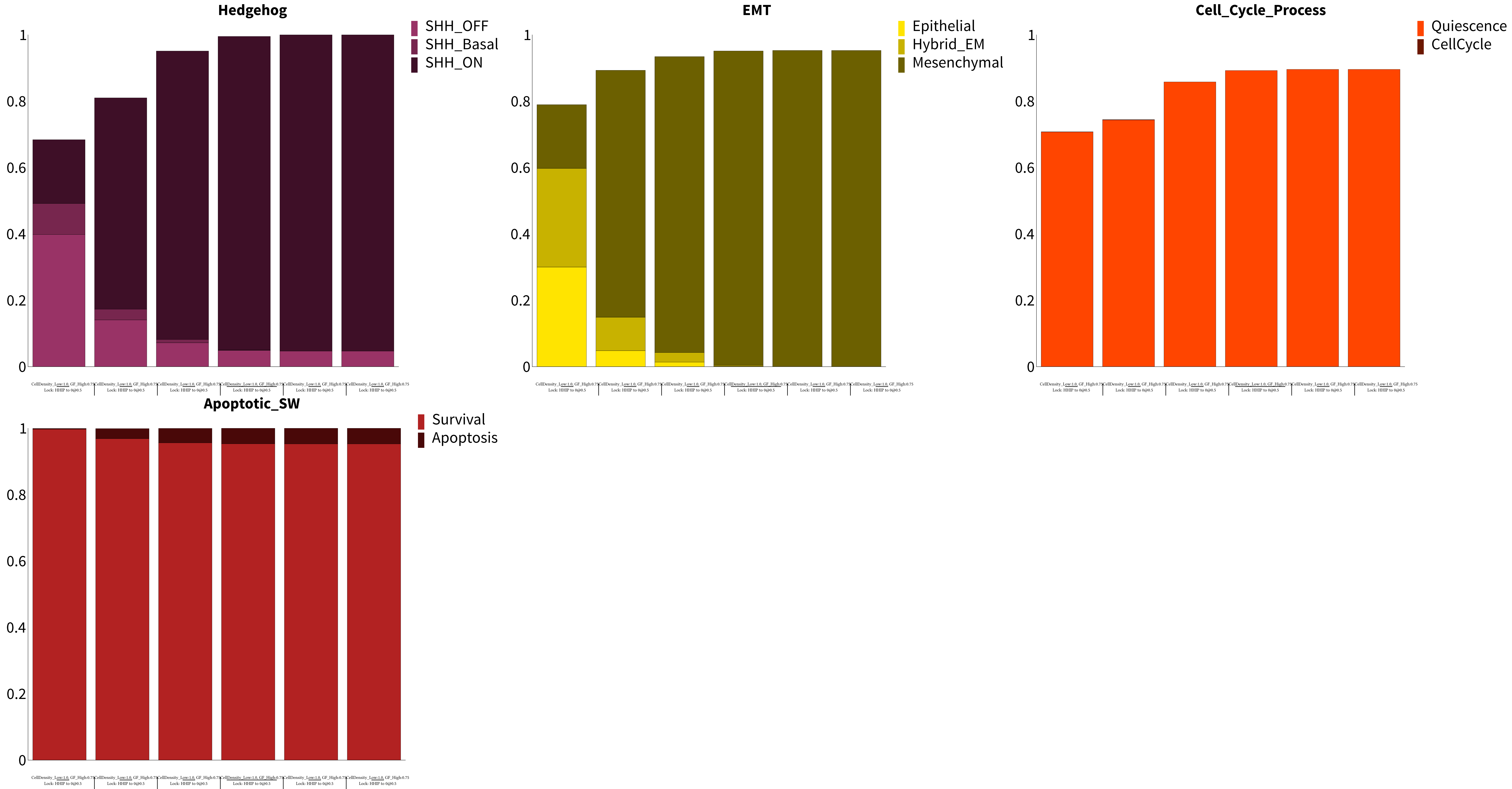

Supplement: Supplementary Material [file 611545_file02.zip › supplementary material/Supplementary File 2 - package to run dynmod - FINAL/_EXP/General_Time_Series/SM_Fig_4C_HHIP_Haplo_Time_BAsync/PhBCh/bc12101221211_SM_Fig_4C_HHIP_Haplo_Time_BAsync.pdf]

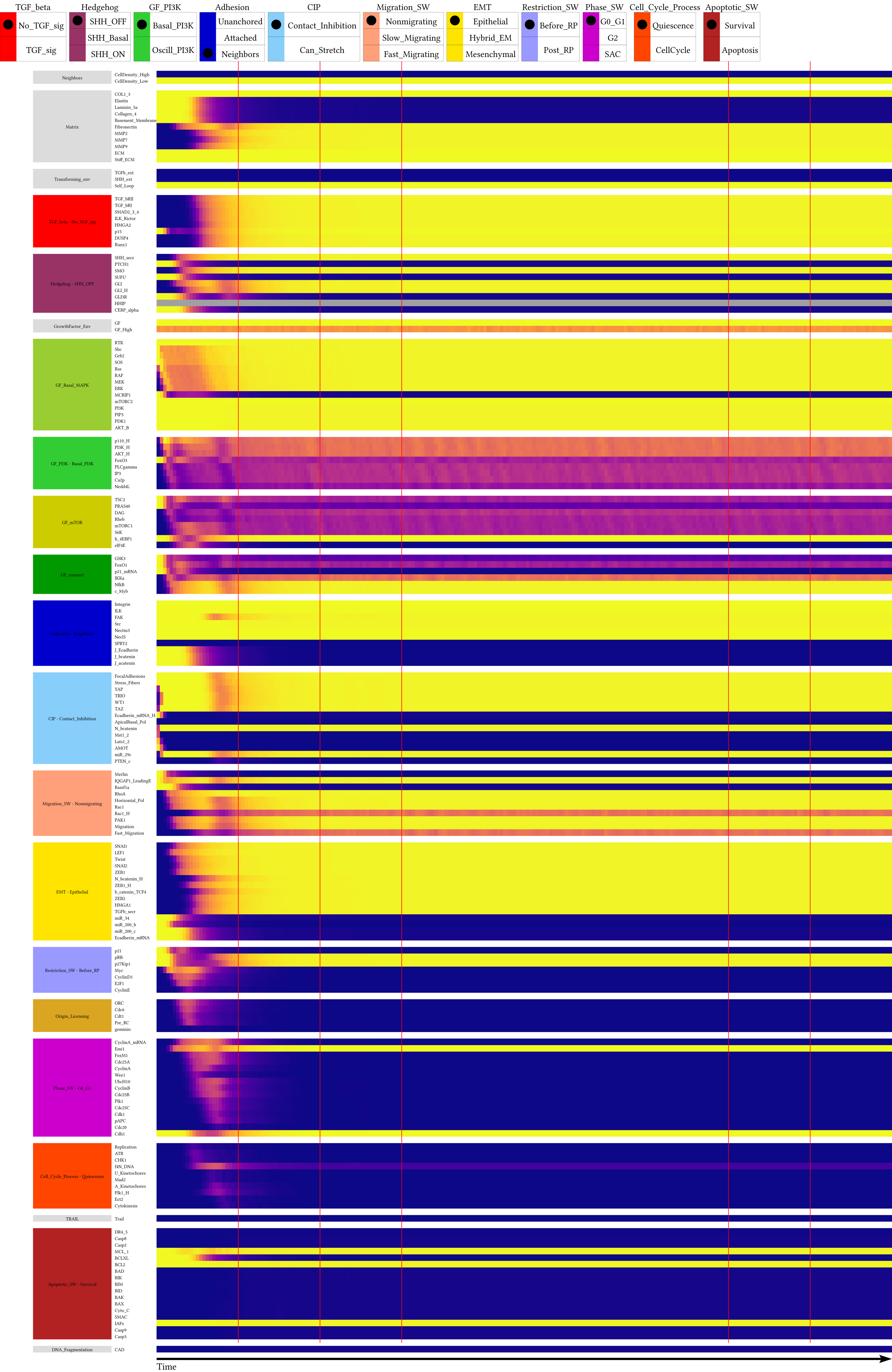

Supplement: Supplementary Material [file 611545_file02.zip › supplementary material/Supplementary File 2 - package to run dynmod - FINAL/_EXP/General_Time_Series/SM_Fig_4C_HHIP_KO_Time_BAsync/NodeTC/bc12101221211_SM_Fig_4C_HHIP_KO_Time_BAsync.pdf]

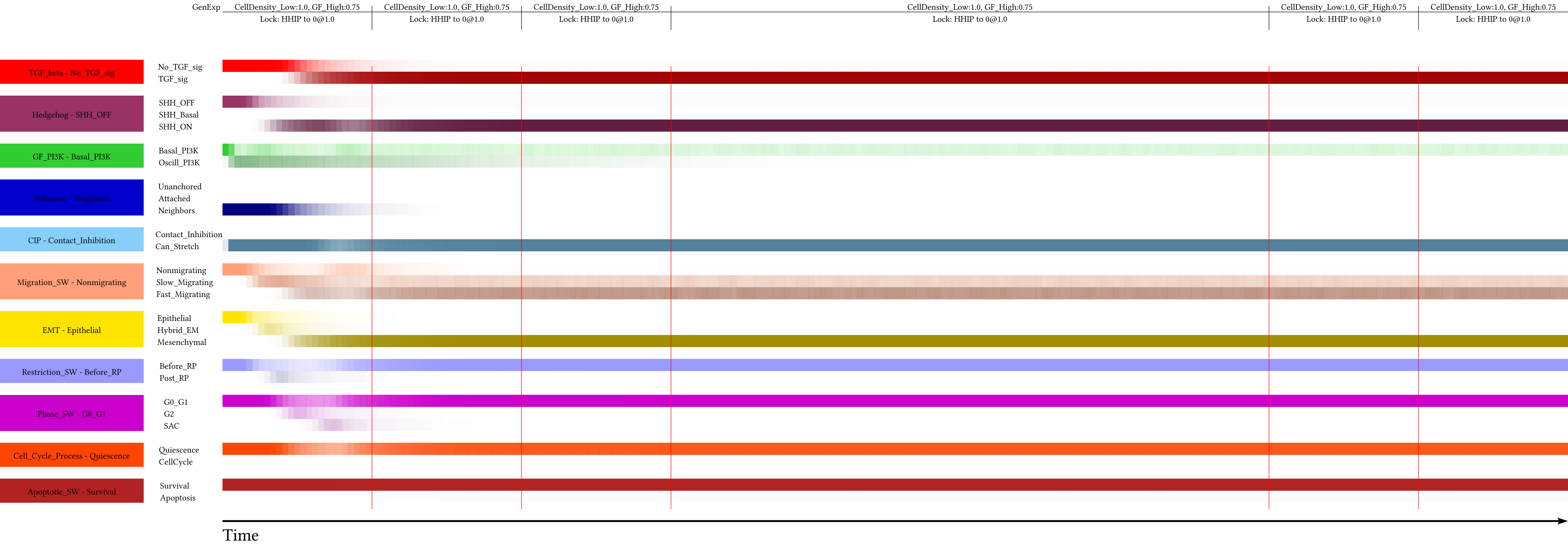

Supplement: Supplementary Material [file 611545_file02.zip › supplementary material/Supplementary File 2 - package to run dynmod - FINAL/_EXP/General_Time_Series/SM_Fig_4C_HHIP_KO_Time_BAsync/PHTC/bc12101221211_SM_Fig_4C_HHIP_KO_Time_BAsync.pdf]

### SM_Fig_4C_HHIP_KO_Time_BAsync

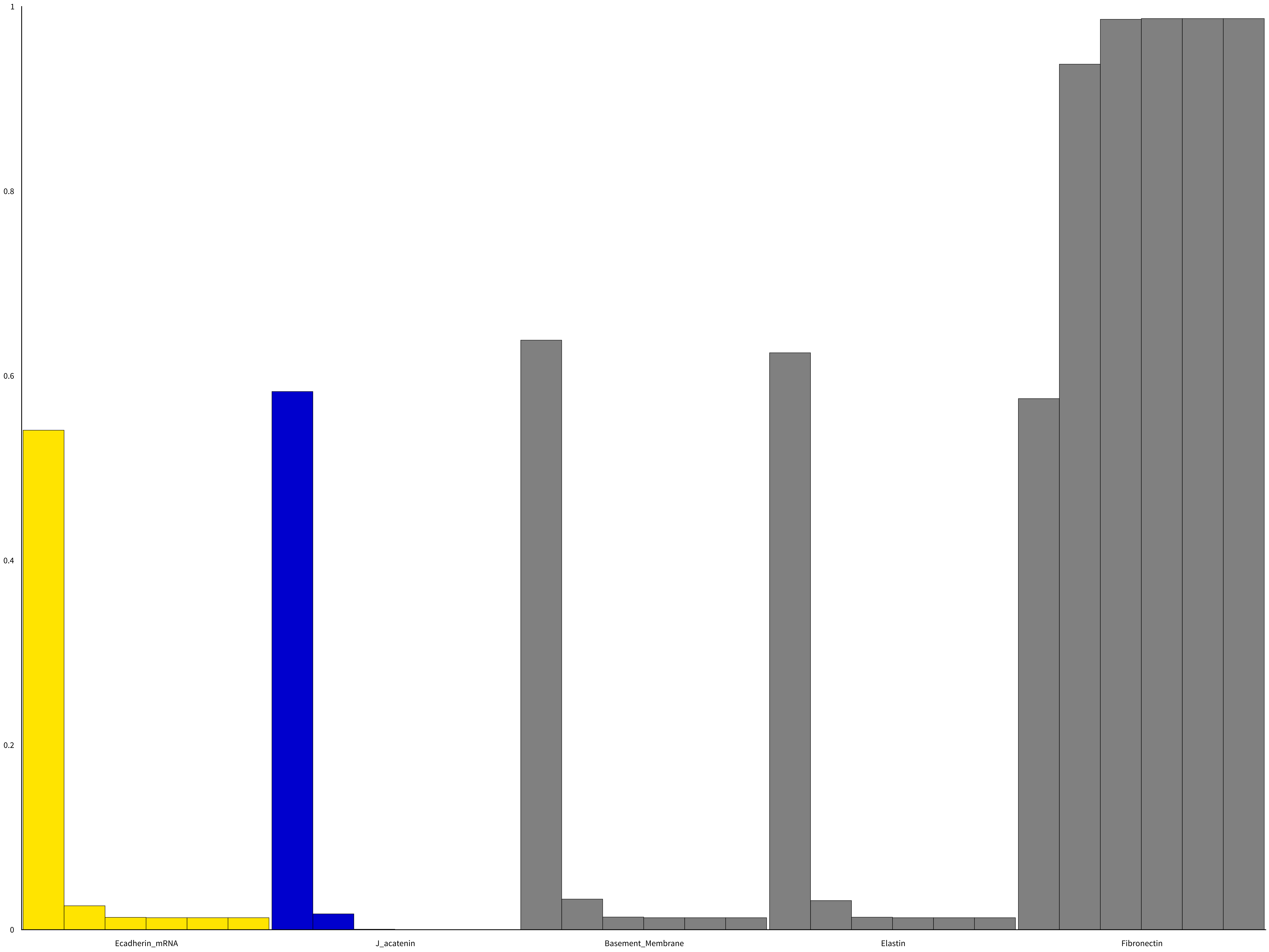

Supplement: Supplementary Material [file 611545_file02.zip › supplementary material/Supplementary File 2 - package to run dynmod - FINAL/_EXP/General_Time_Series/SM_Fig_4C_HHIP_KO_Time_BAsync/NodeBCh/bc12101221211_SM_Fig_4C_HHIP_KO_Time_BAsync.pdf]

### SM_Fig_4C_HHIP_KO_Time_BAsync

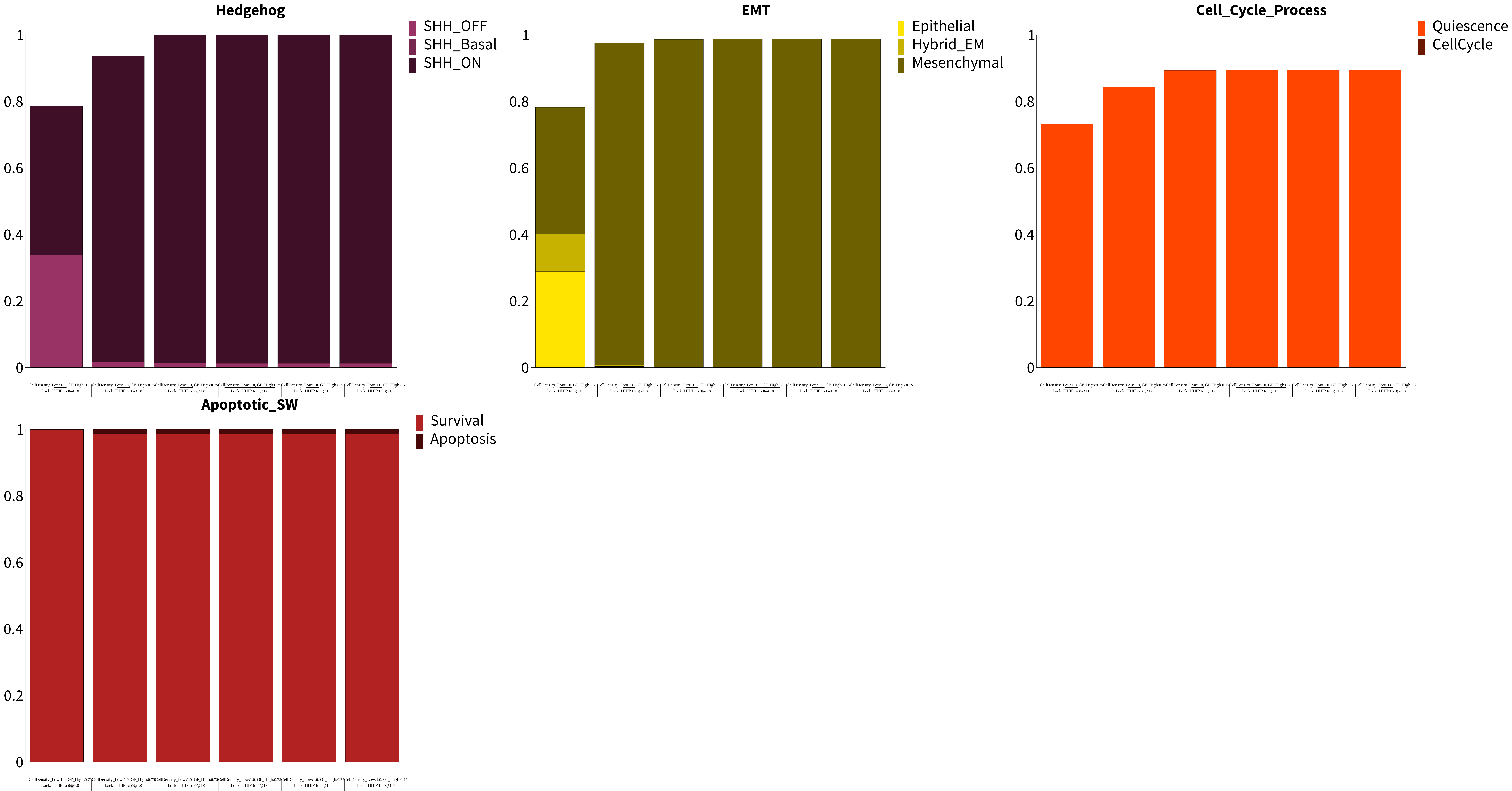

Supplement: Supplementary Material [file 611545_file02.zip › supplementary material/Supplementary File 2 - package to run dynmod - FINAL/_EXP/General_Time_Series/SM_Fig_4C_HHIP_KO_Time_BAsync/PhBCh/bc12101221211_SM_Fig_4C_HHIP_KO_Time_BAsync.pdf]

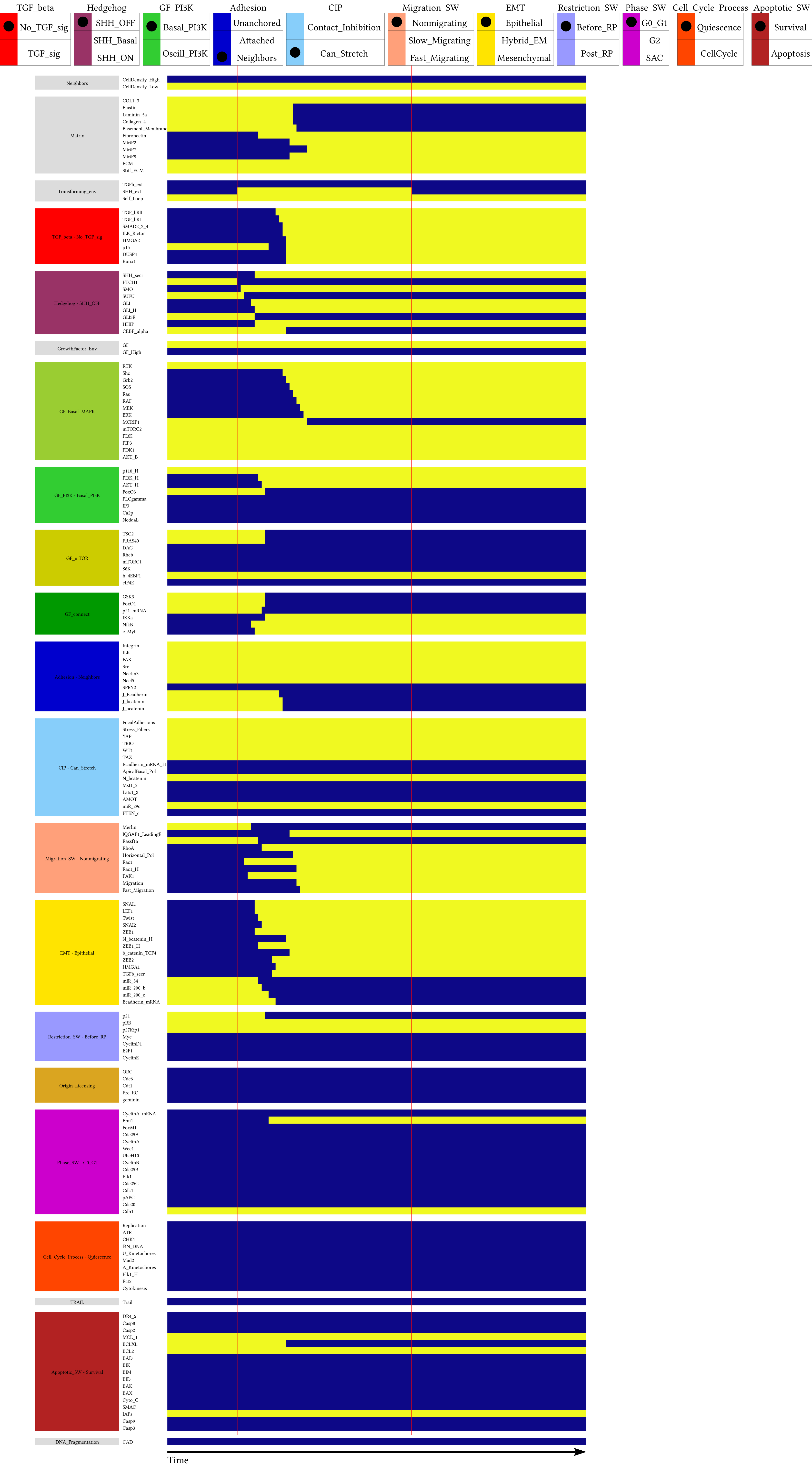

Supplement: Supplementary Material [file 611545_file02.zip › supplementary material/Supplementary File 2 - package to run dynmod - FINAL/_EXP/Pulse1/pulse1_SHH_ext-1.0-UserD 50_wInputs_GF-1_CellDensity_Low-1_SHH_ext-0_TGFb_ext-0_Self_Loop-1_Trail-0_COL1_3-1/NodeTC/bc12100221211_pulse1_SHH_ext-1.0-UserD 50_wInputs_GF-1_CellDensity_Low-1_SHH_ext-0_TGFb_ext-0_Se]
